# Cysteine dependence in *Lactobacillus iners* constitutes a novel therapeutic target to modify the vaginal microbiota

**DOI:** 10.1101/2021.06.12.448098

**Authors:** Seth M. Bloom, Nomfuneko A. Mafunda, Benjamin M. Woolston, Matthew R. Hayward, Josephine F. Frempong, Aaron B. Abai, Jiawu Xu, Alissa J. Mitchell, Xavier Westergaard, Fatima A. Hussain, Nondumiso Xulu, Mary Dong, Krista L. Dong, Thandeka Gumbi, Xolisile Ceasar, Justin K. Rice, Namit Choksi, Nasreen Ismail, Thumbi Ndung’u, Musie S. Ghebremichael, Emily P. Balskus, Caroline M. Mitchell, Douglas S. Kwon

## Abstract

Vaginal microbiota composition affects several important reproductive health outcomes. *Lactobacillus crispatus*-dominant bacterial communities have favorable associations whereas anaerobe-dominant communities deficient of lactobacilli are linked to poor outcomes, including bacterial vaginosis (BV). *Lactobacillus iners*, the most abundant vaginal species worldwide, has adverse associations compared to *L. crispatus*, but standard metronidazole treatment for BV promotes *L. iners*-dominance, likely contributing to post-treatment relapse. *L. iners* is under-studied because it fails to grow in standard *Lactobacillus* media *in vitro*. Here we trace this *in vitro* phenotype to a species-specific cysteine requirement associated with limitations in cysteine-related transport mechanisms and show that vaginal cysteine concentrations correlate with *Lactobacillus* abundance *in vivo*. We demonstrate that cystine uptake inhibitors selectively impede *L. iners* growth and that combining an inhibitor with metronidazole thus promotes *L. crispatus* dominance of defined BV-like communities. These findings identify a novel target for therapeutic vaginal microbiota modulation to improve reproductive health.

## MAIN

Female genital tract (FGT) microbiota composition is linked to numerous important women’s health and reproductive outcomes including HIV risk^1, 2^, preterm birth^3^, mucosal inflammation^4–6^, high-risk human papilloma virus (hrHPV) infection^7^, and cervical dysplasia^7^. Diverse, anaerobe-dominant bacterial communities are associated with negative sequalae and are a hallmark of bacterial vaginosis (BV)^8, 9^, a syndrome involving vaginal discharge and mucosal inflammation that affects up to 58% of women worldwide, with disproportionately high impact in sub-Saharan Africa^10^. By contrast, bacterial communities dominated by *Lactobacillus crispatus* and most other FGT *Lactobacillus* species are generally considered optimal for health^9^. One notable exception is *Lactobacillus iners*-dominant communities, which are associated with many of the same unfavorable outcomes as BV^1, 7, 11^ and have higher risk of transitioning to BV and BV-like communities^3, 12–15^. *L. iners* is the most prevalent and abundant vaginal *Lactobacillus* species worldwide, but factors influencing the balance between *L. iners* and more health-associated lactobacilli are incompletely understood^1, 16–19^.

The close associations between vaginal microbiota and disease make microbiota-targeted therapies a key component of interventions to improve women’s health and reproductive outcomes^8^. Therapeutic microbiota modulation has primarily been studied in relation to BV, but current BV treatments exhibit low efficacy and high recurrence rates. Over 50% of women with symptomatic BV who receive standard therapy with the antibiotic metronidazole (MTZ) experience recurrence by 12 months, while up to 80% of women with a prior history of recurrent BV experience relapse within 16 weeks^20–22^. MTZ typically shifts BV-associated microbiota towards *L. iners*-dominant communities^23–27^, which have high probability of transition back to BV-like communities^12, 14, 15^. Novel strategies to shift the FGT microbiota toward *L. crispatus* instead of *L. iners* during BV treatment are thus a key priority to improve clinical outcomes^15, 21^.

Despite its ubiquity and epidemiologic importance, *L. iners*’ biology remains incompletely characterized^28^. It is known to differ from other *Lactobacillus* species through several key features including secretion of the putative virulence factor inerolysin^29^ and failure to produce D-lactic acid and hydrogen peroxide^30–32^. *L. iners* has a smaller genome with reduced metabolic potential, suggestive of niche adaptation and dependence on exogenous nutrients^31^. However, two widely recognized technical challenges have significantly limited *L. iners* research^18, 28^. The first has been *L. iners*’ failure to grow in standard *Lactobacillus* MRS media^33^, a species-defining phenotype that complicates efforts to isolate and characterize it *in vitro* and which delayed its identification as a species until 1999^28, 33^. Consequently, efficient tools for experimental genetic manipulation of *L. iners* do not currently exist and efforts to develop them have thus far proved unsuccessful^34^, The second challenge has been the paucity and limited diversity of experimental *L. iners* strains and genomes, with many studies reporting little or no success isolating *L. iners* even when culturing from hundreds of clinical samples^35–37^.

In this study we show that *L. iners*’ unique *in vitro* growth limitation relates to restricted capacity to exploit exogenous cysteine (Cys) sources, making it susceptible to small molecule inhibitors that spare *L. crispatus* and other lactobacilli. Using functional experiments and analysis of a novel collection of >1200 diverse *Lactobacillus* genomes, we demonstrate that major vaginal *Lactobacillus* species lack canonical Cys biosynthesis pathways. We find that vaginal Cys concentrations are higher among women without BV and that *in vivo* abundances of both *L. iners* and *L. crispatus* significantly correlate with Cys availability. However, *L. iners* lacks key transport mechanisms for Cys and Cys-containing molecules that are present in other *Lactobacillus* species, rendering it uniquely susceptible to growth inhibition by cystine uptake inhibitors. Combining an inhibitor with MTZ thus promotes *L. crispatus* dominance in mock, BV-like bacterial communities by suppressing *L. iners*. Our findings elucidate important new aspects of *L. iners* biology, resolve key technical barriers that have historically impeded *L. iners* research, and identify a novel target for therapeutically modulating vaginal microbiota to promote women’s health.

## RESULTS

### L-cysteine supplementation supports growth of geographically and genetically diverse *L. iners* strains

We investigated the nutritional dependencies underlying *L. iners*’ species-specific failure to grow in standard *Lactobacillus* MRS media^33^ by testing various nutrient additives. Supplementing MRS broth with IsoVitaleX*™* (a defined mixture of 12 micronutrients) supported robust *L. iners* growth while retaining selectivity against *G. vaginalis* (Fig. 1a). Testing components of IsoVitaleX*™* individually and in combination revealed that L-cysteine (L-Cys) was both necessary and sufficient for this growth phenotype (Extended Data Fig. 1a-f; reagent details in Supplementary Table 1 and 2). L-glutamine (L-Gln) supplementation was neither necessary nor sufficient, but augmented growth in presence of L-Cys, so subsequent experiments were performed using a base of L-Gln supplemented MRS broth (“MRSQ”).

**Fig. 1.**
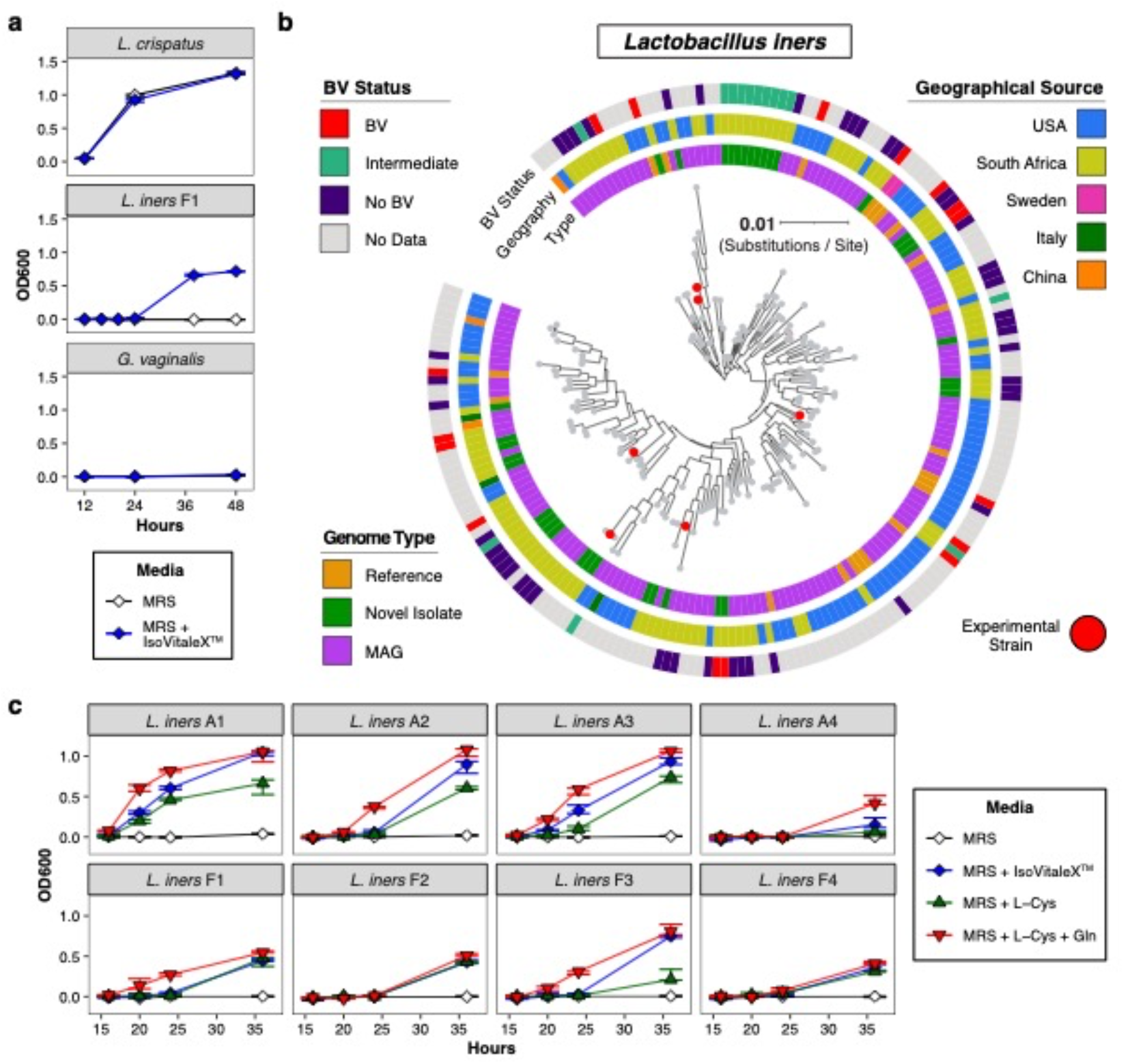
*Lactobacillus iners* requires L-cysteine supplementation to grow in conditions that support other vaginal lactobacilli. (**a**) Growth of *Lactobacillus crispatus*, *L. iners* (strain F1), and *Gardnerella vaginalis* in *Lactobacillus* MRS broth ± supplementation with 2% IsoVitaleX*™*. (**b**) Unrooted phylogenetic tree of 198 *L. iners* genomes from novel and reference bacterial isolates and culture-independent novel metagenomically assembled genomes (MAGs) derived from geographically diverse populations. Red dots indicate isolates experimentally studied in this work. Genome type, geographic origin, and BV status of the source samples are shown. The tree depicts only genome assemblies exceeding certain quality thresholds to ensure robustness of the phylogenetic reconstruction (see **Methods**); additional strains and genomes were included in other analyses (**Supplementary Table 3** and Extended Data Fig. 2). (**c**) Growth of 8 representative US (prefix “A”) or South African (prefix “F”) *L. iners* strains cultured in MRS broth supplemented as indicated with IsoVitaleX*™* (2%), L-cysteine (L-Cys, 4 mM), and/or L-glutamine (L-Gln, 1.1 mM). In **1a,c**, and subsequent figures, bacterial growth was assessed by optical density at 600 nm (OD600) and plotted as median (± range) for 3 replicates from 1 of *≥*2 independent experiments per strain and media condition.

In order to assess species-level generalizability of the Cys-dependent growth phenotype, we established a large, geographically diverse *L. iners* isolate collection and genome catalog of strains from women both with and without BV. The isolate collection was assembled from culture repositories as well as novel US and South African strains isolated on L-Cys-supplemented MRS agar or blood agar. The associated genome catalog consists of 327 genomes from disparate human populations, including genomes from the novel isolates and “reference” isolate genomes retrieved from RefSeq^38, 39^. To enhance catalog diversity and minimize risk of isolation bias, we also included culture-independent metagenome-assembled genomes (MAGs) generated from FGT shotgun metagenomic sequencing studies of U.S., South African, Italian, and Chinese populations^39^. A species-level phylogenetic reconstruction based on this catalog revealed that *L. iners* genomic diversity was largely independent of geographic source, genome type, or clinical context (Fig. 1b). South African and US *L. iners* strains broadly representative of species diversity were selected for further experimental characterization (Fig. 1b and Supplementary Table 3). Supplementing MRSQ broth with L-Cys permitted growth of all strains from this expanded panel (representative examples are shown in Fig. 1c), confirming Cys-dependent growth as a species-level phenotype.

### Vaginal *Lactobacillus* species lack canonical Cys biosynthesis pathways

We next investigated whether *L. iners’* species-specific *in vitro* Cys-dependent growth phenotype was due to differences in Cys biosynthetic capacity as compared to other common FGT lactobacilli. Bacteria canonically synthesize Cys *de novo* from serine (Ser) via either the Cys synthase pathway (*cysE*, followed by *cysK*, *cysM*, or *cysO*) or reverse transsulfuration pathway (*cbs* or *mccA*, followed by *mccB*; Fig. 2a)^40^. We assessed for presence of these pathways in our catalog of 327 *L. iners* genomes, along with similarly constructed catalogs for *L. crispatus* (n = 508 genomes), *L. gasseri* (n = 216), *L. jensenii* (n = 137), and *L. vaginalis* (n = 30) (Extended Data Fig. 2a,b)^39^. These catalogs represent substantial expansion of known genomic diversity for all species, most notably for *L. iners* with a >10-fold increase in the number of genomes and a 73% increase in pan-genome size compared to prior RefSeq-derived genomes (Extended Data Fig. 2b,c). No genomes from any of the species were predicted to possess intact Cys synthase or reverse transsulfuration pathways based on gene annotations performed using eggNOG^41^, although interestingly all non-*iners* species had predicted *mccB* orthologs (Fig. 2b and Supplementary Table 4). Absence of intact Cys synthase and reverse transsulfuration pathways in these species was further supported by results of BLAST^42^ searches against the genome collections using gene sequences of interest from related species (Supplementary File 1) and by analysis of publicly available *Lactobacillus* genomes in the online Integrated Microbial Genomes and Microbiomes (IMG/M) system^43^ (not shown). Thus, all common FGT *Lactobacillus* species, including *L. iners*, are predicted to lack canonical mechanisms for *de novo* Cys biosynthesis.

**Fig. 2.**
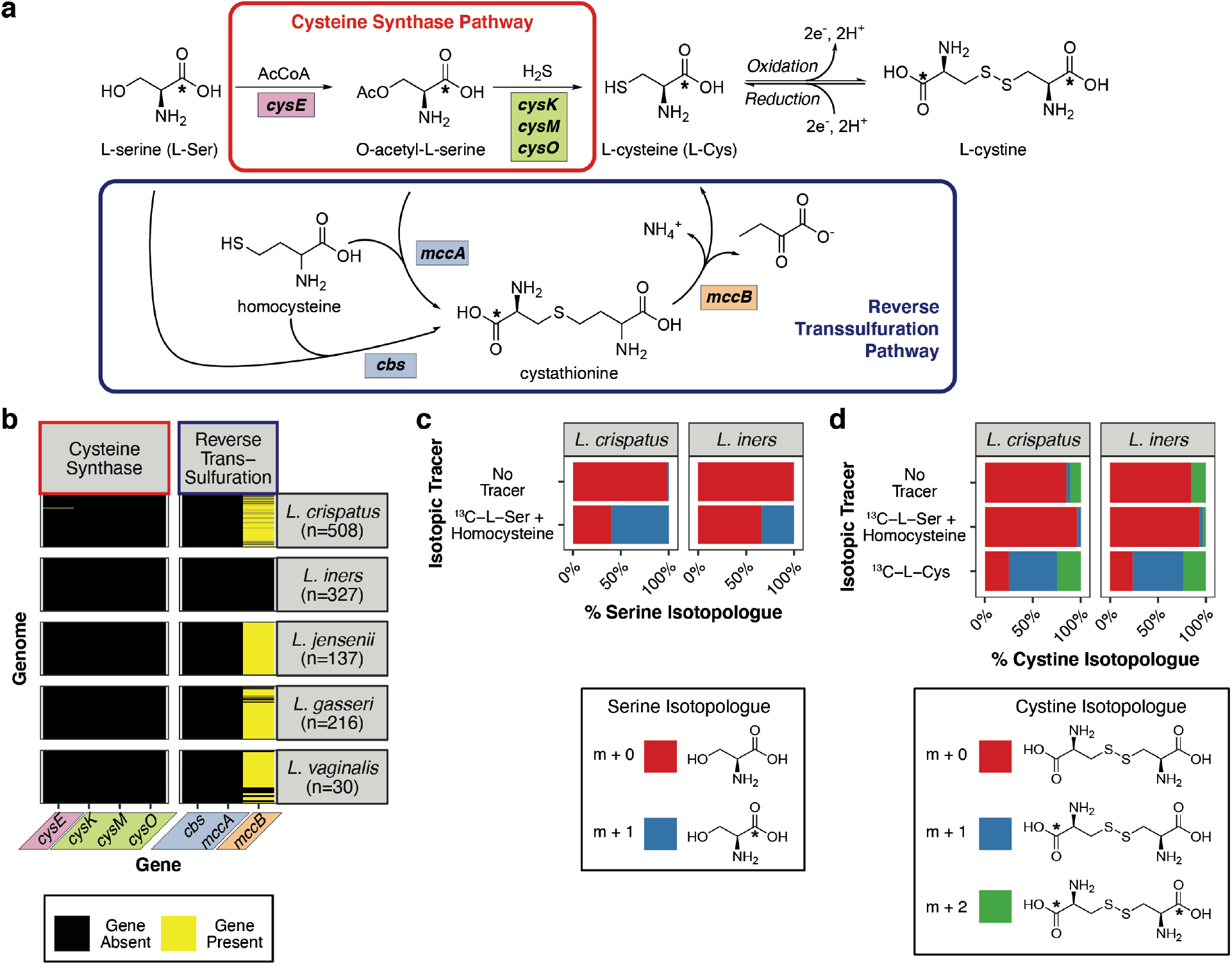
Vaginal lactobacilli lack canonical Cys biosynthesis pathways. **(a)** Canonical bacterial Cys biosynthesis pathways. Cysteine synthase pathway: serine *O*-acetyltransferase (encoded by *cysE*), then cysteine synthase (*cysK*, *cysM*, or *cysO*). Reverse transsulfuration pathway: cystathionine β-synthase (*cbs*) or *O*-acetylserine-dependent cystathionine β-synthase (*mccA*), then cystathionine *γ*-lyase (*mccB*). Asterisks indicate position of ^13^C label in isotopic tracing experiments. **(b)** Predicted presence of genes encoding cysteine biosynthesis enzymes within the cysteine synthase or reverse transsulfuration pathways in isolate genomes and MAGs of the indicated FGT *Lactobacillus* species (n = number of genomes; detailed results in Supplementary Table 4). **(c)** Serine isotopic enrichment in *L. crispatus* and *L. iners* grown for 24 hours in L-Gln-supplemented MRS broth (“MRSQ”) containing no isotopic tracer (supplemented with unlabeled L-Cys 4 mM) or supplemented with 1-^13^C-L-serine (^13^C-L-Ser) + homocysteine (both 4 mM). Plot depicts fractional abundance of unlabeled (m+0, Ser) and labeled (m+1, ^13^C-Ser) isotopologues in cellular hydrolysates. **(d)** Cystine isotopic enrichment in lysates of *L. crispatus* or *L. iners* cultured as shown in **(c)** or in broth supplemented with 1-^13^C-L-cysteine (^13^C-L-Cys; 4 mM). An oxidation step during sample processing converts Cys to cystine for quantification. Plot depicts fractional abundance of cystine isotopologues: unlabeled (m+0; cystine), partially labeled (m+1; ^13^C_1_-L-cystine), and fully labeled (m+2; ^13^C_2_-L-cystine) isotopologues. Data in 2**c**,**d** depict median values of 3 replicates per group (full data in Supplementary Tables 5 and 6).

We phenotypically tested this result by measuring the ability of *L. iners* and *L. crispatus* to convert Ser into Cys *in vitro*. Bacteria were grown in MRSQ broth supplemented with isotopically labeled Ser (^13^C-L-Ser) plus homocysteine or with labeled L-Cys (^13^C-L-Cys; labeling scheme as in Fig. 2a). Bacteria were lysed and hydrolyzed, then cellular amino acid isotopic labeling patterns were analyzed via liquid chromatography-mass spectrometry (LC-MS). Comparison to bacteria grown without isotopic tracers revealed that both *L. iners* and *L. crispatus* took up labeled Ser but failed to convert it to Cys (Fig. 2c,d), consistent with genomic predictions. By contrast, both strains readily incorporated labeled Cys, supporting the genomic prediction that despite their different growth phenotypes in MRS broth, both utilize exogenous Cys.

### Vaginal Cys concentrations are lower in women with BV and strongly correlate with *in vivo Lactobacillus* abundance

Given the prediction that vaginal lactobacilli lack canonical Cys biosynthesis pathways, we investigated *in vivo* relationships between vaginal Cys concentrations, BV, and *Lactobacillus* colonization. We measured Cys concentrations in cervicovaginal lavage (CVL) fluid from 142 participants in a cohort of South African women aged 18-23 years^1, 4, 44^, 53 of whom were also evaluated for BV using the Nugent scoring method^45^. Cys concentrations differed strikingly by BV status (p = 6.3 x 10^-9^), with significantly higher concentrations among women without BV compared to those with BV (Fig. 3a). To explore underlying microbiota composition in this cohort, we performed bacterial 16S rRNA gene sequencing of cervical swab samples from all participants. Employing previously established criteria, individual bacterial communities were classified into four “cervicotypes” (CTs): CT1 (*L. crispatus*-dominant), CT2 (*L. iners*-dominant), CT3 (*Gardnerella*-dominant), and CT4 (non*-Lactobacillus*-dominant, diverse communities typically featuring high *Prevotella* abundance)^1^ (Fig. 3b). As expected^9^, CT was strongly related to BV status (p = 1.902 x 10^-11^, two-sided Fisher’s Exact Test; Extended Fig 3a). Cys concentration differed markedly between CTs (p = 2.7 x 10^-14^ overall), with significantly higher concentrations in *Lactobacillus*-dominant CTs compared to CT3 and CT4 (Fig. 3c).

**Fig. 3.**
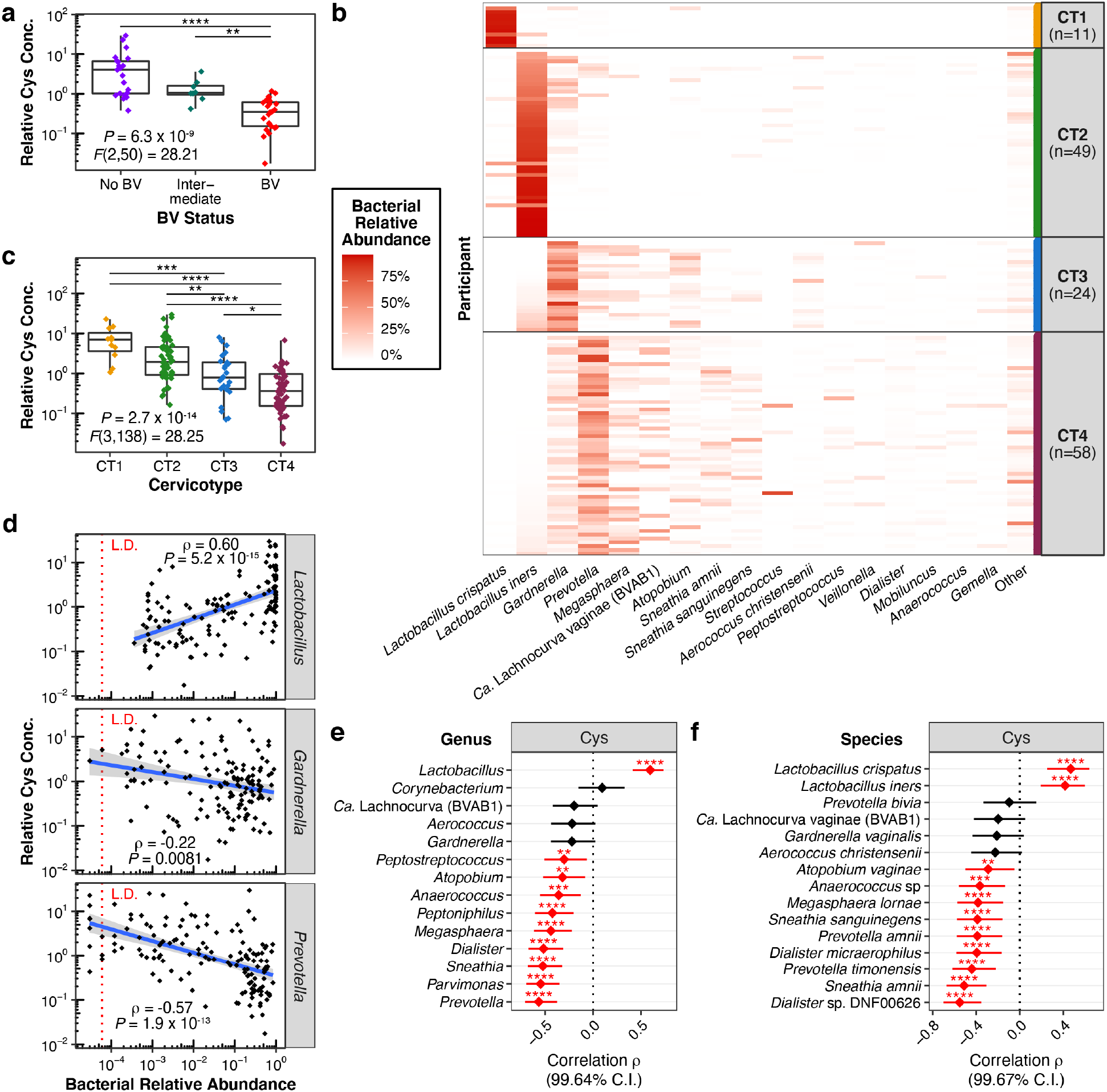
Vaginal Cys concentrations are higher in women without BV and correlate with *Lactobacillus*-dominant microbiota. **(a)** Relative Cys concentration by BV status in cervicovaginal lavage (CVL) fluid from 53 South African women (21 without BV, 24 with BV, and 8 intermediate by Nugent method^45^). **(b)** FGT bacterial microbiota composition among 142 HIV-uninfected South African women (including the 53 from **a**), determined by bacterial 16S rRNA gene sequencing (taxonomy assignments as in Supplementary Table 7). Bacterial communities were classified into “cervicotypes” (CTs) using previously defined criteria^1^. **(c)** Relative Cys concentrations per CT in CVL fluid from women in **(b)**. In **(a & c)** between-group differences of log-transformed concentrations were determined by one-way analysis of variance (ANOVA) with post-hoc Tukey’s test; all significant pairwise differences are shown. **(d)** Spearman rank correlation between Cys concentrations and bacterial relative abundances of the CT-associated genera *Lactobacillus*, *Gardnerella*, and *Prevotella*, showing correlation coefficients (*ρ*) with unadjusted p-values. A linear regression line (solid blue) with 95% confidence interval is added to assist visualization. Red dotted line represents limit of detection (L.D.). **(e & f)** Spearman correlation coefficients (*ρ*) with adjusted confidence intervals between Cys concentrations and relative abundances of each genus **(e)** or species **(f)** detected at >50% prevalence in cohort. P-values and confidence intervals were adjusted for multiple comparisons using the Bonferroni method (full statistic results in Supplementary Tables 8 & 9). Significance of adjusted p-values depicted as * p *≤* 0.05, ** p *≤* 0.01, *** p *≤* 0.001, **** p *≤* 0.0001.

We next examined correlations between Cys concentrations and relative abundance of individual taxa within the microbiota. Analysis of key CT-associated genera revealed that *Lactobacillus* relative abundance had a strong positive correlation with Cys concentrations (*ρ* = 0.6, p = 5.2 x 10^-15^ by Spearman correlation), while *Prevotella* abundance had equally strong negative correlation (*ρ* = −0.57, p = 1.9 x 10^-13^) and *Gardnerella* abundance was weakly negatively correlated to Cys concentrations(Fig. 3d). *L. crispatus* and *L. iners* each positively correlated with Cys when analyzed individually (Extended Data Fig. 3b). To assess whether lactobacilli were the only bacteria correlated with Cys, we calculated correlations for each taxon detected in *≥*50% of samples at either the genus or species level (Extended Fig 3c,d), adjusting for multiple comparisons. *Lactobacillus* (genus-level) and *L. crispatus* and *L. iners* (species-level) were the only taxa positively correlated with Cys (Fig. 3e,f; full statistical results in Supplementary Tables 8 & 9). Other taxa, including various BV-associated bacteria, showed no correlation or had significant negative correlations with Cys concentrations. Results were similar for the Cys-containing peptides reduced glutathione (GSH) and cysteinylglycine (Extended Fig. 4 and Supplementary Tables 8 & 9). Thus, vaginal Cys concentration is higher among women without BV and preferentially correlates with abundance of both *L. iners* and *L. crispatus*, consistent with the hypothesis that Cys availability is important for *Lactobacillus* colonization success *in vivo*.

### *L. iners* utilizes L-cystine but has limited capacity to exploit more complex L-Cys sources

The species-specific nature of *in vitro* Cys-dependence for *L. iners* appeared discordant with the finding that other *Lactobacillus* species also lack canonical Cys biosynthesis pathways and that abundance of both *L. iners* and *L. crispatus in vivo* correlates with high vaginal Cys concentrations. We therefore hypothesized that *L. iners’* unique *in vitro* growth phenotype was due to restricted capacity to utilize exogenous Cys sources in MRS broth compared to other lactobacilli. Alternatively, Cys supplementation might support *L. iners* growth by acting as a chemical reducing agent rather than a direct nutritional supplement. Measurement of concentrations of free molecular Cys and cystine (its oxidized counterpart) in MRSQ broth revealed that they were ≤4.6 μM and 0.7 μM respectively, below levels required for growth by Cys-auxotrophic *E. coli* strains^46^. Supplementation with L-cystine produced similar *L. iners* growth as L-Cys supplementation (Fig 4a), even in presence of the oxidizing agent hydrogen peroxide (H_2_O_2_; Extended Fig 5b), demonstrating that L-cystine supports growth by acting as a direct nutritional supplement rather than causing chemical reduction of the media. Thus *L. iners* does not require a chemically reduced environment to grow when it has access to a bioavailable source of L-Cys (including L-cystine), but L-Cys bioavailability in un-supplemented MRSQ broth is inadequate for *L. iners*.

**Fig. 4.**
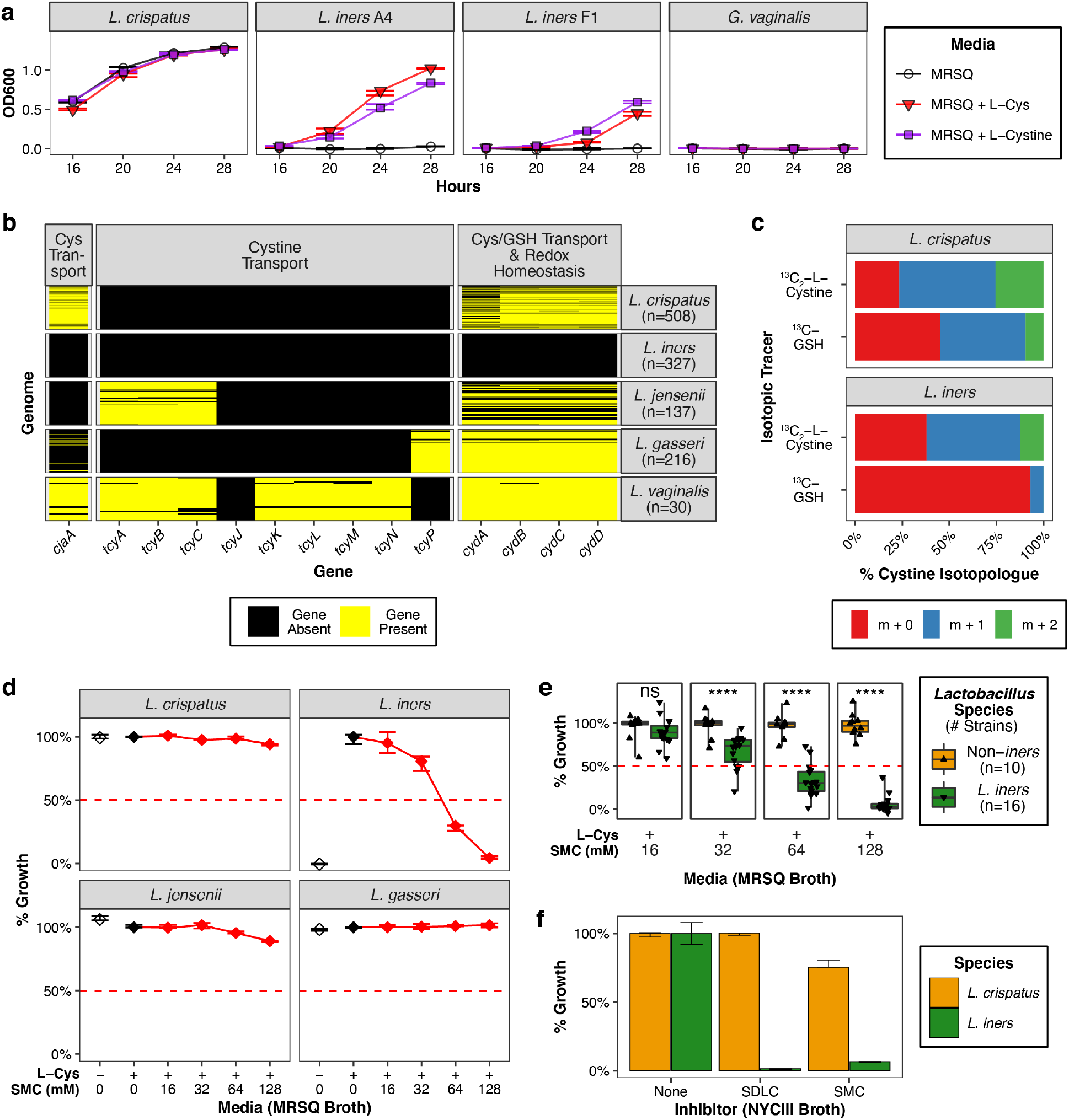
*L. iners* lacks Cys-related transport mechanisms present in other lactobacilli and is selectively inhibited by cystine uptake inhibitors. (**a**) Growth of *L. crispatus*, *G. vaginalis*, and representative *L. iners* strains in MRSQ broth ± L-Cys (4 mM) or L-cystine (2 mM). **(b)** Predicted presence of putative Cys transport-related gene *cjaA*, cystine transport loci *tcyABC*, *tcyJKLMN*, and *tcyP*, and the Cys/GSH transport/redox homeostasis locus *cydABCD* locus in FGT *Lactobacillus* species genomes and MAGs (n = number genomes; detailed results in Supplementary Table 4). TcyBC takes up glutathione in some species. **(c)** Cystine isotopic enrichment in *L. crispatus* or *L. iners* grown in MRSQ broth containing labeled L-cystine (^13^C_2_-L-cystine; 2 mM) or GSH (^13^C-GSH; 4 mM; synthesis described in Extended Data Fig. 7). Plot depicts isotopologue median fractional abundance. (Full data in Supplementary Tables 5 & 6). **(d)** Growth of representative *Lactobacillus* strains in L-Cys-supplemented MRSQ broth containing the cystine uptake inhibitor *S*-methyl-L-cysteine (SMC). Percentages calculated relative to median OD600 measurement in L-Cys-containing no-inhibitor control. Plots depict median (± range) for 3 replicates per condition and are representative of *≥*2 independent experiments per condition. **(e)** Median growth inhibition of non-*iners Lactobacillus* strains (*L. crispatus*, n = 7; *L. jensenii*, n = 2; *L. gasseri*, n = 1) or *L. iners* (n = 16) by SMC in L-Cys-supplemented MRSQ broth. Two-sided t-test: ns, p *≥* 0.05; ****, p < 0.0001. **(f)** Growth inhibition of *L. crispatus* and *L. iners* by SMC (128 mM) or SDLC (2 mM) in NYCIII broth.

The ability of non-*iners* lactobacilli to grow in un-supplemented MRS broth implied presence of other Cys sources that *L. iners* failed to utilize (MRS contains yeast and beef extracts, which contribute protein-derived peptides in addition to monomeric amino acids). Adding chemical reducing agents to MRSQ broth substantially raised Cys and GSH concentrations (Extended Data Fig. 5a), demonstrating that most Cys and Cys-containing molecules in un-supplemented media exist as mixed disulfide compounds. Addition of chemical reducing agents other than L-Cys also supported *L. iners* growth (Extended Data Fig. 5b,c), but in contrast to L-cystine, the oxidized counterparts of these reducing agents did not. Thus MRS broth contains complex sources of L-Cys, including mixed disulfides, that *L. iners* has restricted capacity to utilize.

### *L. iners* lacks Cys-related transport mechanisms possessed by other vaginal lactobacilli and its growth is selectively inhibited by cystine uptake inhibitors

We hypothesized that *L. iners*’s restricted ability to use exogenous Cys sources might reflect a limited repertoire of transport mechanisms for Cys and Cys-containing molecules in comparison to other lactobacilli. Cys transporters are not well-characterized in bacteria compared to transporters for other amino acids^47^. Uptake of Cys and Cys-containing molecules is described as occurring through multiple known or putative mechanisms. These include the well-characterized cystine transport systems TcyABC, TcyJKLMN, and TcyP^48^, a putative Cys ABC transporter associated with the Cys-binding protein CjaA^47, 49^, and the heterodimeric ABC transporter CydDC encoded by the redox-regulating locus *cydABCD*^50^, a conserved system that exports both Cys and GSH to the periplasm in *E. coli*^50^ and is proposed to perform glutathione uptake in lactobacilli^51^. Analysis of our expanded *Lactobacillus* genome catalogs annotated using eggNOG revealed that genomes of all non-*iners* species encoded *cydABCD*, while *L. crispatus*, *L. vaginalis*, and some *L. gasseri* encoded one or more predicted *cjaA* orthologs and *L. jensenii*, *L. gasseri*, and *L. vaginalis* each encoded one or more cystine transport systems (Fig. 4b and Supplementary Table 4). Low-affinity Cys uptake can also occur via the branched-chain amino acid importer LivJ/LivKHMGF^47^, but no species possessed *livKHMGF* orthologs except 2 *L. iners* MAGs (out of 327 genomes), nor did any species possess the full GsiABCD glutathione transport system (Extended Data Fig 6a and Supplementary Table 4)^52^. The findings were supported by results of BLAST^42^ searches against the genome collections using gene sequences of interest from related species (Supplementary File 1). Thus, each of these FGT *Lactobacillus* species except *L. iners* encodes multiple, mechanistically distinct known or putative systems for transporting Cys or Cys-containing molecules, while *L. iners*’ Cys-related uptake occurs via one or more currently unrecognized mechanisms.

*L. iners* has thus far proved refractory to genetic manipulation and experimental tools to perform genetic screens are lacking^34^, so we used phenotypic approaches to characterize the breadth of its Cys-related transport mechanisms. Isotopic tracing (Fig. 4c) revealed that *L. iners* incorporated labeled L-cystine (^13^C_2_-L-cystine) at levels similar to *L. crispatus* but failed to take up labeled GSH (^13^C-GSH; synthesis detailed in Extended Data Fig. 7), consistent with predicted absence of GSH transport activity (Fig. 4b and Extended Data Fig. 6a). We hypothesized that if *L. iners* differed from other lactobacilli in relying on a more limited repertoire of transport mechanisms, its growth might be uniquely impacted by uptake inhibition. We therefore tested effects of the known cystine uptake inhibitors *S*-methyl-L-cysteine (SMC) and seleno-DL-cystine (SDLC)^48^ on *L. iners* growth in L-Cys-supplemented MRSQ broth. SMC caused species-specific, dose-dependent growth inhibition of a diverse collection of *L. iners* strains at concentrations of 32-64 mM, while SDLC caused selective inhibition at a concentration of 0.25 mM (Fig. 4d,e and Extended Data Fig. 6b,c). Inhibitor potency is known to vary for different transporters, but these concentrations appear higher for SMC and slightly higher for SDLC than reported inhibitory concentrations for the TcyJKLMN and TcyP transporters in *Bacillus subtilis*^48^. To confirm that growth inhibition was not an artefact specific to MRS media, we assessed inhibition in NYCIII broth, a serum-containing, nutrient-rich medium that supports growth of diverse fastidious FGT bacteria^53^. Both SMC and SDLC inhibited *L. iners* growth in NYCIII broth, but not growth of *L. crispatus* (Fig. 4f). They also selectively inhibited *L. iners* growth in MRSQ broth treated with chemical reducing agents, supporting the conclusion that reducing agents promote *L. iners* growth by enhancing Cys bioavailability (Extended Data Fig. 6d). Collectively these results confirm the genomic prediction that *L. iners* possesses a uniquely restricted repertoire of Cys-related transport mechanisms, rendering it susceptible to selective growth inhibition.

### SMC provides selective advantage to *L. crispatus* in direct competition with *L. iners*

To assess the functional significance of *L. iners* growth inhibition by cystine uptake inhibitors, we queried whether an inhibitor could shift the balance between *L. iners* and *L. crispatus* in mixed culture competition assays. We tested pairwise strain combinations of *L. crispatus* and *L. iners* in L-Cys-supplemented MRSQ broth at varying SMC concentrations, then measured ratios of *L. iners* to *L. crispatus* in the mixed cultures by 16S rRNA gene sequencing. Experimental controls were included to confirm expected growth patterns and absence of significant contamination during sample processing (Extended Data Fig. 8a-c). SMC significantly suppressed *L. iners* relative to *L. crispatus* in a dose-dependent fashion (representative experiments in Fig. 5a), confirming that it offers selective advantage to *L. crispatus* during direct competition.

**Fig. 5.**
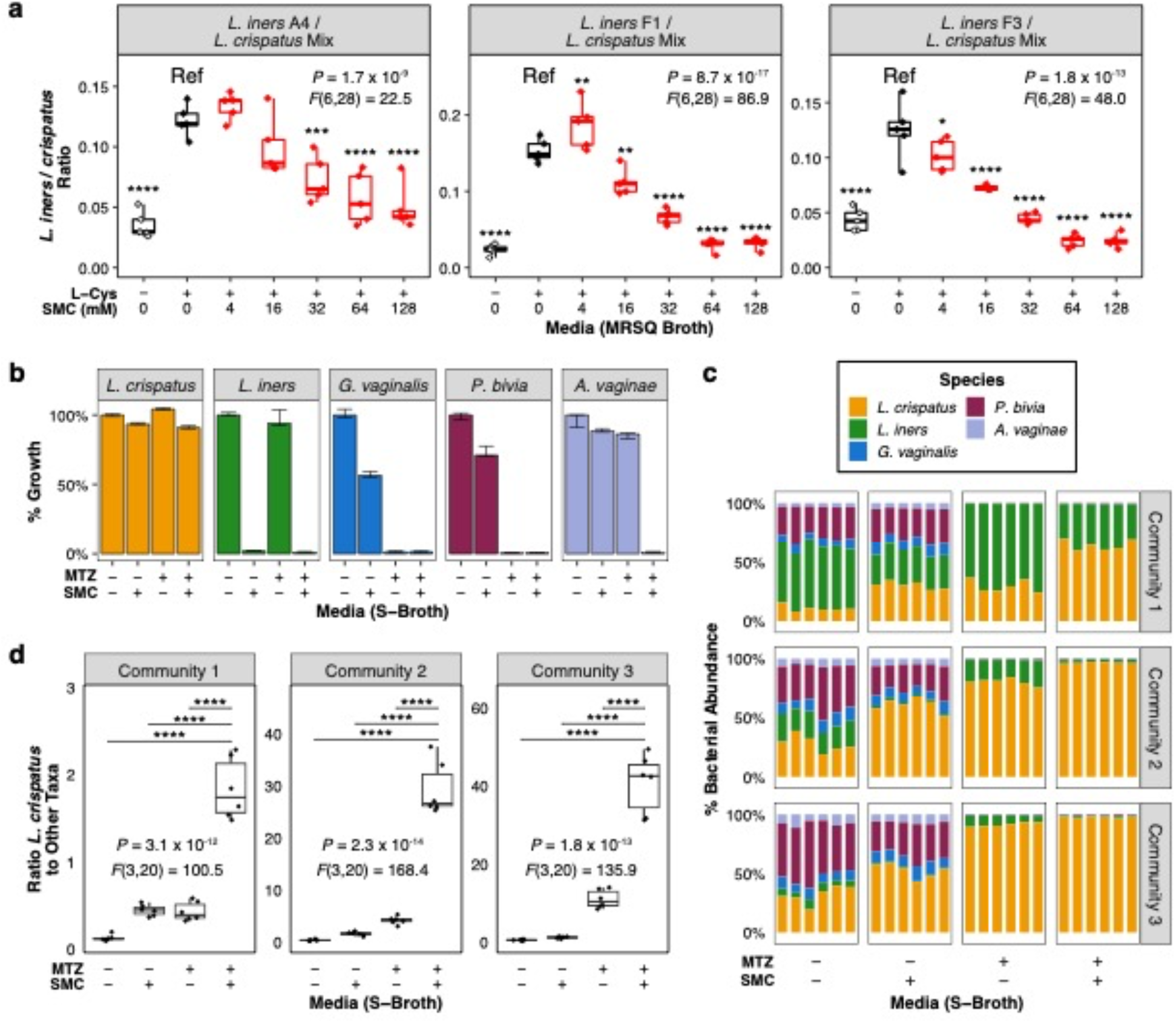
SMC inhibits *L. iners* in competition with *L. crispatus* and combining SMC with metronidazole enhances *L. crispatus* dominance of mock BV-like communities. (**a**) Ratios of *L. iners* to *L. crispatus* in representative mixed culture competition assays after 28 hours incubation in L-Cys-supplemented MRSQ broth with varying concentrations of SMC. Ratios were determined via sequencing the bacterial 16S rRNA gene in DNA isolated from the mixed cultures; plots depict results for 5 replicates per condition. Between-group differences were determined by one-way ANOVA. Pairwise comparisons to the reference condition (“Ref”: no-inhibitor, L-Cys-supplemented control) were calculated using Dunnett’s test. All significant comparisons are shown (full statistical results in Supplementary Table 10). **(b)** Growth inhibition of *L. crispatus*, *L. iners*, and three BV-associated species at 48 hours incubation in S-broth with or without SMC and/or metronidazole (MTZ). **(c)** Relative abundance of bacterial species in three representative mock BV-like communities (each with a different *L. iners* strain) grown in S-broth with or without SMC and/or MTZ at 28 hours incubation. Composition was determined by bacterial 16S rRNA gene sequencing. **(d)** Ratios of *L. crispatus* to the sum of all other taxa in the mock communities shown in **(c)**. **(c,d)** depict 6 replicates per condition. Between-group differences in **(d)** were determined via one-way ANOVA with post-hoc Tukey test; selected pairwise differences are shown (full statistical results in Supplementary Table 11). In 5a,e significance is depicted as * p *≤* 0.05, ** p *≤* 0.01, *** p *≤* 0.001, **** p *≤* 0.0001

### Combining SMC with MTZ enhances *L. crispatus* dominance in mock BV-like communities

To assess whether cystine uptake inhibitors could augment BV therapy by preferentially promoting *L. crispatus* expansion, we investigated SMC’s effects in combination with MTZ on mock BV-like bacterial communities grown in a rich nutritional milieu. Multiple South African *L. crispatus* isolates unexpectedly failed to grow in NYCIII broth, so we developed a novel serum-containing formulation named “S-broth” that supported the South African *L. crispatus* strains as well as diverse BV-associated bacteria (Extended Data Fig. 9a). Using S-broth, we tested effects of SMC and MTZ in pure cultures of *L. crispatus*, *L. iners*, *G. vaginalis*, *Prevotella bivia*, and *Atopobium* (*Fannyhessea*) *vaginae*. Neither MTZ nor SMC blocked *L. crispatus* growth, whereas SMC inhibited *L. iners* and MTZ suppressed both *G. vaginalis* and *P. bivia* (Fig. 5b). Our *A. vaginae* strain was relatively MTZ-resistant (a known phenotype of some strains from this species), but interestingly susceptibility was enhanced by addition of SMC.

We cultured defined communities of these species in S-broth and assessed whether adding SMC to MTZ enhanced *L. crispatus* community dominance. Each community contained a different pairwise combination of *L. crispatus* and *L. iners* strains (input ratios as in Extended Data Fig. 9b). We used 16S sequencing to quantify mock community composition, including technical controls to confirm input strain identity and absence of significant contamination during processing (Extended Data Fig. 9c,d). Cultivation in media without inhibitors produced diverse mixtures of all 5 species (representative examples in Fig. 5c). As hypothesized, SMC alone diminished *L. iners* relative abundance, MTZ alone suppressed BV-associated species, and MTZ combined with SMC preferentially favored *L. crispatus*. To quantify the competitive advantage for *L. crispatus*, we calculated ratios between *L. crispatus* relative abundance and the summed abundances of all other taxa (Fig. 5d). The combination of MTZ and SMC increased this ratio significantly more than MTZ alone, supporting the concept that cystine uptake inhibitors could complement MTZ in promoting *L. crispatus* dominance during BV treatment.

## DISCUSSION

From its initial species description in 1999, *L. iners* was recognized to possess diminished metabolic potential compared to other lactobacilli – indeed, the name “*iners*” refers to its relative inertness in biochemical assays^33^. But despite being the most abundant FGT species worldwide and having several adverse health associations compared to *L. crispatus*^1, 3, 7, 11, 15, 16, 18, 19, 28, 54, 55–57^, *L. iners*’ biology has remained poorly characterized because it fails to grow in standard *Lactobacillus* MRS media, limiting laboratory-based experiments and resulting in a paucity of isolates and genomes^18, 28^. Here we resolve this unique growth defect by identifying a species-specific requirement for exogenous Cys supplementation in *L. iners*, which we show corresponds to a uniquely limited repertoire of mechanisms for transporting exogenous Cys and Cys-containing molecules compared to other FGT lactobacilli. Employing functional assays along with a novel collection of isolates and >1200 *Lactobacillus* genomes derived from diverse geographical and clinical contexts, we extend and experimentally confirm predictions from smaller-scale genomic studies suggesting FGT lactobacilli lack Cys biosynthesis pathways^58^. We further demonstrate that vaginal Cys concentrations strongly correlate with *Lactobacillus* relative abundance *in vivo* in a South African cohort. However, compared to other lactobacilli, *L. iners* has a restricted repertoire of transport mechanisms for Cys and Cys-containing molecules, rendering it selectively susceptible to cystine uptake inhibitors. Combining a cystine uptake inhibitor with MTZ (a standard antibiotic used for BV treatment) enhances *L. crispatus* dominance in mock, BV-like bacterial communities by suppressing growth of *L. iners*. These advances establish an essential foundation for future characterization of *L. iners*’ biology, diversity, and effects on human health – including its role in predisposition to BV and other adverse outcomes^1, 7, 11, 12, 14, 15^ –while simultaneously identifying a novel target for therapeutic vaginal microbiota modulation.

The close *in vivo* correlation between Cys and *L. iners* and *L. crispatus* abundance is consistent with the hypothesis that Cys availability may influence FGT microbiota composition. A smaller US-based case-control study comparing metabolomes of women with and without BV found that vaginal Cys concentrations positively correlated with *L. crispatus* and *L. jensenii* and negatively correlated with various BV-associated species^59^. Neither strength nor statistical significance of individual species-metabolite correlations were reported in that study, but it appeared to show a weaker correlation between *L. iners* and Cys than the correlation we identify here^59^, which may reflect the different study designs or may indicate differences in species abundance and prevalence between the two study populations. Since FGT lactobacilli appear to lack *de novo* Cys biosynthetic capacity, we hypothesize that the high Cys concentrations observed in *Lactobacillus*-dominant states are likely to be primarily host-derived. Cys’s positive correlation with lactobacilli and negative correlation with many BV-associated bacteria could thus be due to mechanisms including preferential *Lactobacillus* colonization of hosts with high mucosal Cys secretion, induction of host Cys secretion by lactobacilli, and/or degradation of Cys by BV-associated bacteria^59^ as a means to outcompete lactobacilli. However other metabolites in addition to Cys also likely influence microbiota composition. Further investigation of FGT microbiome-metabolome relationships and *in vitro* exploration of bacterial metabolism and interactions with host epithelium will be required to fully assess these possibilities.

A striking aspect of our findings is the paucity of transport mechanisms for Cys and Cys-containing molecules in *L. iners* compared to other FGT lactobacilli. Unlike other amino acids, transporters for Cys are not well characterized in bacteria due to technical challenges inherent to redox chemistry that complicate Cys quantification and efforts to distinguish between transport of reduced Cys and transport of its oxidized counterpart, cystine^47^. The relatively few functional studies of putative bacterial Cys transporters have focused predominantly on Gram-negative bacteria from the phylum Proteobacteria rather than lactobacilli or related Gram-positive species^47, 49^. In contrast to Cys transporters, several bacterial cystine transporters are better characterized, including among Gram-positive species^48^. We found that genomes of non-*iners* FGT *Lactobacillus* species were each predicted to encode multiple, mechanistically distinct known or putative transporters for Cys, cystine, and/or Cys-containing molecules, but our *L. iners* genome analysis failed to identify predicted transporters for any of these molecules. However both L-Cys and L-cystine supported *L. iners* growth, and isotopic tracing experiments demonstrated uptake when *L. iners* was grown in presence of either labeled L-Cys or L-cystine, indicating existence of one or more currently unrecognized uptake mechanisms. Genetic tools to definitively identify the unknown transporter(s) do not currently exist for *L. iners*^34^, but our finding that *L.. iners* growth is selectively inhibited by known cystine uptake inhibitors provides further evidence that it lacks the multiple, mechanistically distinct Cys-related transport mechanisms present in other FGT *Lactobacillus* species.

In addition to elucidating *L. iners* biology and function within the microbiota, our results have important implications for BV treatment. Standard BV therapy with MTZ often provides initial relief, but disease recurs in up to 75% of cases^20–22^. High post-treatment recurrence rates may be partially explained by the fact that MTZ typically promotes *L. iners*-dominant FGT bacterial communities^23–27^, which are more prone than *L. crispatus*-dominant communities to transition back to BV-like states^3, 13–15^. In a recent study of longitudinal FGT microbiota dynamics in a South African population, we demonstrated that *L. iners*-dominant communities constitute a “gateway” for transition to more diverse, BV-like communities. Modelling of microbiota transition probabilities predicted that interventions which specifically shift transition probabilities in favor of *L. crispatus*-dominance over *L. iners*-dominance would have the greatest effect in increasing the fraction of women with *L. crispatus*-dominant communities^15^. Therapies to promote *L. crispatus* dominance during BV treatment are under active investigation. In a randomized trial of a live *L. crispatus* biotherapeutic administered after MTZ treatment, BV recurred less frequently among biotherapeutic recipients than placebo recipients^21^. However recurrence rates in the treatment arm remained high (30% recurrence within 3 months) and post-treatment microbiota composition was not reported^21^. A small Israeli study without a control group reported achieving *L. crispatus*-dominance and long-term remission in four of five women treated for refractory BV with MTZ plus vaginal microbiome transplants from healthy donors^60^, but most patients required multiple rounds of therapy. Interestingly, that study did not report detecting *L. iners* in any donors or recipients – a notable difference from typical FGT microbiota composition in North and South America, Europe, Asia, Africa, and Australia^1, 2, 3, 4, 5, 11, 12, 16, 17, 23, 24–27, 53–57, 59, 61^. Thus live biotherapeutic approaches for BV show promise but appear unlikely to be a panacea. In this study, we show that combining MTZ with a novel growth inhibitor for *L. iners* was superior to MTZ alone in promoting *L. crispatus* community dominance of defined, BV-like bacterial communities. The specific *L. iners* inhibitors identified here have potential limitations for direct therapeutic use in humans including low potency (SMC) and potential host toxicity (SDLC), but show proof of principle for a novel therapeutic approach. Further development of this approach as an adjunct to antibiotics has potential to improve therapies for adverse microbiota-linked reproductive health conditions worldwide, including for women in low and middle income countries, where the burden of these adverse outcomes is highest.

## METHODS

### Clinical cohorts and samples

This study includes novel specimens from participants in the Females Rising through Education, Support, and Health (FRESH) cohort, an ongoing prospective observational study based in Umlazi, South Africa that enrolls 18-23-year-old, HIV-uninfected, non-pregnant, otherwise healthy women. Study characteristics and inclusion and exclusion criteria have been described elsewhere in greater detail^1, 4, 44^. The study protocol was approved by the Massachusetts General Hospital Institutional Review Board (2012P001812/MGH) and the Biomedical Research Ethics Committee of the University of KwaZulu-Natal (UKZN). All participants provided informed consent. Multiple cervical swab samples (Puritan 6” Sterile Standard Foam Swab w/Polystyrene Handle) were collected by swabbing the ectocervix in two full revolutions under direct visualization during speculum exam. The swabs were then used to make a slide preparation for Gram stain analysis, were cryopreserved in thioglycolate broth with 20% glycerol for bacterial isolation, or were frozen without cryopreservatives for bacterial isolation or nucleic acid extraction and microbiome profiling. Cervicovaginal lavage (CVL) samples were collected using a flexible plastic bulb pipette to dispense 5 mL of normal saline into the vaginal vault, wash the cervix four times, then re-aspirate the fluid into a 15 mL conical tube. CVL and swab samples were stored on ice for 1-4 hours during transport to the processing laboratory at the Nelson R. Mandela School of Medicine at UKZN, where swabs were stored at −80°C and CVL samples were centrifuged at 700xg for 10 minutes at 4°C, then supernatants were transferred to cryovials and stored at −80°C.

BV status was determined using the Gram-stain-based Nugent scoring method ^45^ for a subset of 53 previously uncharacterized, HIV-uninfected FRESH participants sampled consecutively in September and October 2017. Scoring was performed at Global Laboratories (now Neuberg Global Laboratories, Durban, South Africa), an accredited commercial laboratory diagnostics company, by trained laboratory technologists. Paired FGT metabolite and 16S rRNA gene-based microbiome profiling were performed on samples from an expanded group of 143 HIV-uninfected FRESH participants (including the above 53 participants) who were sampled consecutively between May and October 2017. Nugent scoring for BV assessment among these study participants was performed only sporadically until a protocol change that instituted universal Nugent scoring in September 2017. We therefore restricted analysis of BV-metabolite associations to samples collected under the universal testing protocol to minimize risk of bias. One participant was excluded from analysis of vaginal metabolite concentrations based on the pre-specified criterion of low 16S sequence read count (<10,000 reads). No *a priori* sample size calculation was performed in relation to analysis of microbiota-metabolite correlations because this is (to our knowledge) the first analysis of FGT microbiota-metabolite relationships in a sub-Saharan African cohort and we therefore lacked sufficient information upon which to make informed *a priori* estimates about metabolite distributions for power calculations. Metagenomic whole genome shotgun sequencing (WGS) for metagenomic genome assemblies was performed on samples from the above 143 participants and on samples from additional FRESH study participants for whom 16S rRNA gene-based microbiome profiling has previously been reported^1, 4^. Primary bacterial isolates from a subset of study participants were generated from cryopreserved swabs as described below.

Additional metagenomics data and bacterial isolates were generated by the Vaginal Microbiome Research Consortium (VMRC; https://vmrc4health.org/) as detailed more fully elsewhere^39, 62^. These included samples originating from two longitudinal studies of US women of reproductive age. One cohort enrolled pregnant women and collected vaginal swab specimens during pregnancy and in the post-partum period^13^. The second cohort enrolled non-pregnant women and collected vaginal swabs longitudinally over 10 weeks^25^.

### Bacterial culture conditions and additives for bacterial growth media

Bacterial cultivation was performed at 35-37°C under anaerobic conditions using an AS-580 anaerobic chamber (Anaerobe Systems) in an atmosphere of 5% carbon dioxide, 5% hydrogen, and 90% nitrogen (Airgas^®^). All media and other culture reagents were pre-reduced (deoxygenated) by placement overnight in the anaerobic chamber prior to use. Nutritional additives or inhibitors (reagents listed in Supplementary Table 1; additives prepared as in Supplementary Table 2) were added to autoclave-sterilized broth media after cooling to room temperature (broth media), then broth was re-sterilized by passage through a 0.22 μm vacuum filter before being transferred to the anaerobic chamber to pre-reduce (deoxygenate) for use in experiments. For testing of nutrient pools (Extended Data Figure 1), a single concentrated stock comprising all pool constituents was added to media as per Supplementary Table 2 rather than stocks of each additive being added separately. In contrast to other additives, stocks of hydrogen peroxide (H_2_O_2_) and metronidazole (MTZ) were added to pre-reduced broth media as freshly prepared, filter-sterilized stock solutions immediately prior to inoculation of bacterial cultures due to potential volatility or chemical instability. For preparation of solid media, filter-sterilized additives were added to autoclave-sterilized agar media after cooling to 50°C, mixed using a magnetic stir/hot plate, then used to pour agar plates.

### Bacterial broth culture media

Growth characteristics and nutritional requirements distinguishing *L. iners* from other *Lactobacillus* species were investigated using *Lactobacillus* MRS broth. We tested two different commercial MRS formulations (BD Difco*™* and Hardy Criterion). The BD Difco*™*-formulated media is referred to as “MRS” while the Hardy Criterion-formulated media is referred to as “HMRS” throughout this paper. The broth was prepared by autoclaving according to manufacturer instructions and allowed to cool to room temperature. We confirmed that *L. crispatus*, *L. jensenii*, and *L. gasseri* grew rapidly and robustly in both formulations whereas *L. iners* failed to grow even after >10 days in HMRS and grew only after a prolonged delay of ∼48-72 hours in BD Difco*™*-formulated MRS. Supplementation of MRS broth (BD Difco*™*) with IsoVitaleX™ (2% v/v), L-Cys or L-Cys-containing mixtures (augmented by L-Gln) enabled robust growth of *L. iners* by 20-36 hours, depending on the experimental strain (Fig. 1a,c, and Extended Data Fig. 1a-e), while supplementation of HMRS broth had similar pro-growth effects after a more prolonged lag phase (Extended Data Fig. 1f). Unless otherwise indicated, figures depict growth in MRS (BD Difco*™*) broth; key growth results were reproduced where shown using HMRS broth. MRS broth supplemented with L-Gln (1.1 mM) is abbreviated “MRSQ” (or “HMRSQ” for L-Gln-supplemented HMRS broth) in figures and text.

NYCIII broth (ATCC medium 1685) was prepared using a slightly modified version of the standard ATCC protocol. Pre-media consisted of 4 g/L HEPES, 15 g/L Proteose Peptone No.3 and 5 g/L Sodium Chloride in 875 mL distilled water, was pH-adjusted to 7.3 and autoclaved. Prior to use, complete NYCIII broth was prepared from autoclaved, cooled pre-media by adding dextrose (3g/45mL) at 7.5% v/v, yeast extract solution at 2.5% v/v, and heat inactivated horse serum at 10% v/v, then sterilized by passage through a 0.22 μm vacuum filter. Where indicated, NYCIII broth was supplemented with IsoVitaleX™ (2% v/v), Vitamin K1-Hemin Solution (5% v/v), and/or Tween-80 (1 g/L); volumes of distilled water in the pre-media were decreased accordingly to ensure all other components were present at standard concentrations in the final solution.

“S-broth” pre-media consisting of 37 g/L BHI Broth, yeast extract (powder) 10 g/L, and dextrose 1 g/L was brought to a boil in 880 mL distilled water, then autoclaved (121°C for 15 min) and cooled, followed by addition of fetal bovine serum 5% v/v, Vitamin K1-Hemin Solution 5% v/v, and IsoVitaleX™ Enrichment 2% v/v to a final volume of 1L. The complete broth was then sterilized by passage through a 0.22 micron vacuum filter. Unless otherwise indicated, S-broth was supplemented with 1g/L Tween-80.

Except where otherwise indicated, lactobacilli were propagated in MRSQ broth with L-Cys (4 mM), *Prevotella* species were propagated in Wilkins-Chalgren Anaerobe Broth (Thermo Scientific™), and other species including *Gardnerella vaginalis* were propagated in NYCIII broth.

### Bacterial solid culture media

Non-*iners Lactobacillus* isolates were maintained and quantitatively titered on *Lactobacillus* MRS Agar plates (purchased as prepared media from Hardy Diagnostics). Most other species, including *L. iners*, were maintained and titered on Columbia Blood Agar (“CBA”; purchased as prepared media from Hardy Diagnostics). *Prevotella* species were maintained on CBA agar or CDC Anaerobe Laked Sheep Blood Agar with Kanamycin and Vancomycin (“LKV”; purchased as prepared media from BD BBL™). For bacterial isolations from primary clinical samples, we used commercial CBA and LKV agar plates and prepared in-house MRS agar plates by autoclaving *Lactobacillus* MRS Broth (BD DIFCO™) in Noble Agar (BD DIFCO™, 2% w/v), cooling to 50°C with agitation, then supplementing with IsoVitaleX™ Enrichment (BD BBL™; 2% v/v) with or without Vitamin K1-Hemin Solution (BD BBL™; 5% v/v), or with freshly filter-sterilized stocks of L-Cys and L-Gln (at final concentrations of 4 mM and 1.1 mM respectively). All solid media was stored at 4°C in the dark after purchase or preparation until being introduced to the anaerobic chamber for deoxygenation and use.

### Bacterial isolate sources and isolation protocols

Experiments were performed using both previously described strains from culture repositories and novel bacterial isolates from South African and US cervicovaginal samples (See Supplementary Table 3). Isolations of South African strains from FRESH cohort samples were performed at the Ragon Institute. Cervicovaginal swabs were thawed in the anaerobic chamber. Swabs that had been frozen dry (without cryopreservatives) were immersed in 500uL of room temperature pre-reduced PBS. Swabs that had been frozen in cryopreservative solution were rapidly thawed directly without additional dilution. Thawed swabs were agitated vigorously within the solution for 30 seconds using sterile forceps to dislodge bacteria, then removed from the cryovial using a sterile micropipette tip to strain excess fluid from the swab. The solution was then serially diluted in 10-fold dilutions in sterile, pre-reduced PBS and dilutions were plated on *Lactobacillus* MRS agar plates (with or without nutritional additives), CBA, and LKV agar. After 2-5 days incubation (depending on growth rate), colonies were selected based on distinct morphology, and individual representative colonies were sub-cultured onto reduced agar plates, then further clonally sub-cultured into broth media. Aliquots from the purified broth cultures were frozen at −80°C in 20% glycerol, with additional aliquots reserved for nucleic acid extraction and sequence-based taxonomic identification as described below. *L. iners* colonies were preferentially identified for isolation from primary swab cultures on supplemented MRS or CBA plates based on characteristic morphology after 3-4 days incubation (circular, white-translucent colonies between 0.5 and 2 mm in diameter with entire vs slightly irregular edges, smooth surface, and convex/umbonate contour that were non-hemolytic on CBA after ∼4 days). In experienced hands (S.M.B. and N.A.M.), morphology-based identification of L. iners colonies for isolation from CBA plates resulted in equally high or higher rates of L. iners recovery from clinical samples compared to isolation using plates with supplemented MRS media (which had lower CFU yield than CBA), enabling us to perform targeted isolations of *L. iners* from >80% of attempted clinical samples, including BV-associated, anaerobe-dominated samples known to have <5% *L. iners* relative abundance based on 16S sequencing (not shown), Isolated bacteria were identified based on Illumina-based sequencing of genomic DNA and/or Sanger sequencing of the full-length bacterial 16S rRNA gene as described below. BV status of the samples from which isolates were derived was determined based on Nugent scoring when available (Supplementary Table 3).

The following experimental strains were obtained through BEI Resources, NIAID, NIH as part of the Human Microbiome Project: Lactobacillus iners, Strain LEAF 2053A-b, HM-705; Lactobacillus iners, Strain LEAF 3008A-a, HM-708; Lactobacillus iners, Strain UPII 143-D, HM-126. Other strains were obtained from the indicated isolate collections (Supplementary Table 3). Genomes corresponding to these strains were obtained from RefSeq. Information about the BV status of the samples from which these isolates and other isolate genomes were derived was obtained (when available) from metadata accompanying the isolate genome entries in RefSeq or the Genomes OnLine Database (GOLD) or from associated information from the reference strain repositories from which they were obtained. Additional novel US-based strains were obtained from Jacques Ravel and the VMRC, isolated from the sources detailed above^39^, and non-iners Lactobacillus strains previously isolated from a clinical cohort in Seattle, Washington, (a kind gift from Dr. Jeanne Marrazzo^63^) were genome-sequenced and analyzed for genomic content as described below, but not otherwise studied experimentally.

Unless otherwise indicated, all growth, growth inhibition, isotopic tracing, and mock community experiments employed *L. crispatus* strain 233, *L. iners* strain F1, and *G. vaginalis* strain ATCC 14018. Reference bacterial strains used experimentally in this study are available either from the indicated culture repositories (Supplementary table 3) (for reference strains) and novel strains used in these experiments are available upon reasonable request to the corresponding author.

### Nucleic acid extraction

Total nucleic acids from cervicovaginal swab samples were extracted with a phenol-chloroform method, which includes a bead beating process to disrupt bacteria as previously described^4^. Genomic DNA (gDNA) from bacterial isolates or mock communities cultured *in vitro* was extracted using a plate-based protocol that included a bead beating process and combined phenol-chloroform isolation with Qiagen QIAamp 96 DNA QIAcube HT kit (Qiagen) procedures.

### Bacterial full-length 16S rRNA gene PCR and Sanger sequencing

The near-full-length bacterial 16S rRNA gene was PCR-amplified from gDNA of individual bacterial isolates using the broad-range primers Bact-8F (5’-AGAGTTTGATCCTGGCTCAG-3’) and Bact-1510R (5’-CGGTTACCTTGTTACGACTT-3’; Integrated DNA Technologies, Inc.). PCR amplicons were confirmed via agarose gel electrophoresis, then purified and directly Sanger-sequenced on an ABI3730XL DNA Analyzer at the MGH Center for Computational and Integrative Biology DNA Core using the Bact-8F and Bact-1510R PCR primers as sequencing primers in separate forward and reverse sequencing reactions. Isolates identities were determined by BLAST searches against the NCBI nucleotide collection (nt) database and Bacterial 16S Ribosomal RNA RefSeq Targeted Loci Project database (NCBI accession PRJNA33175), and further confirmed in most cases by whole genome sequencing as detailed below.

### Shotgun library preparation

Shotgun sequencing libraries for cultured bacterial isolates or culture-independent swab samples were prepared following a modified protocol of Baym et. al^64^ using the Nextera DNA Library Preparation Kit (Illumina) and KAPA HiFi Library Amplification Kit (Kapa Biosystems). Briefly, DNA concentration of each sample was standardized to 0.6 ng/μL after quantification with SYBR Green I. Simultaneous fragmentation and sequencing adaptor incorporation was performed by mixing 0.6 ng DNA with 1.25 μL TD buffer and 0.25 μL TDE1 (Tagment DNA Enzyem, Nextera) and incubating for 10 min at 55°C. Tagmented DNA fragments were PCR-amplified using KAPA high fidelity library amplification reagents and primers incorporating Illumina adaptor sequences and sample barcodes. Products were pooled, purified with magnetic beads, and paired-end sequenced on an Illumina NextSeq with a 300-cycle kit.

### Genome and meta-genome sequence processing and assembly

We constructed *Lactobacillus* genome catalogs from reference isolate genomes as well as from genome sequences of novel bacterial isolates and metagenomically assembled genomes (MAGs) assembled from culture-independent shotgun metagenomic sequencing of genital tract samples, described in greater detail elsewhere^39^. Briefly, we retrieved all reference genomes annotated as *Lactobacillus crispatus*, *Lactobacillus iners*, *Lactobacillus jensenii*, *Lactobacillus gasseri*, or *Lactobacillus vaginalis* that were deposited in the NCBI RefSeq database^38^ as of February 2020. Genomes of novel FGT isolates from the South African FRESH cohort (isolated as described above) and of non-*iners Lactobacillus* strains isolated from a US-based cohort^63^ were sequenced as described above at the Ragon Institute. Additional US FGT *Lactobacillus* isolate genomes were provided by the VMRC from the sources detailed above^39^. A total of >1000 FGT shotgun metagenomic samples from studies in South Africa, the United States of America, Italy, and China were used to generate MAGS^39^. Sequence reads from isolate genomes and shotgun metagenomic samples were trimmed and filtered to high quality reads. Human reads from shotgun metagenomic samples were removed by mapping to GRCh38 (Genbank accession GCA_000001405.15). We assembled genomes and MAGs from high quality reads, then binned contigs and removed contamination. Bin completeness, contamination, and strain heterogeneity metrics were determined, then genomes and MAGs were assigned quality scores based on criteria from the Genome Standards Consortium^65^. Species genome bins (SGBs) were determined based on 95% pairwise absolute nucleotide identity (ANI).

### Lactobacillus genome taxonomy assignment

We assigned taxonomy to SGBs with FGT *Lactobacillus* genomes based on presence of reference genomes with defined NCBI taxonomy, confirming that each contained reference genomes only from a single species. Of note, for both *L. jensenii* and *L. gasseri*, publicly available reference genomes classified in RefSeq as belonging to these species actually segregate into two separate SGBs per species, indicating that *L. jensenii* and *L. gasseri* (as traditionally defined) each comprise two distinct genomic species^39, 66^. However we analyzed *L. jensenii* and *L. gasseri* as single species units in this study to correspond with prevailing paradigms in literature on 16S gene-based FGT microbiota profiling^67^.

### Cervicovaginal *Lactobacillus* pan-genome construction and analysis

We constructed a cervicovaginal *Lactobacillus* pan-genome using genes from catalogs of *L. iners*, *L. crispatus*, *L. jensenii*, *L. gasseri*, and *L. vaginalis* genomes and MAGs. To maximize comprehensiveness of the pan-genome, we included genomes and MAGs classified as high-quality (>90% completeness and <5% contamination) or medium-quality (*≥*50% completeness and <10% contamination) assemblies based on Genome Standards Consortium criteria^65^. Genes were identified within individual genomes by Prokka v1.14.5^68^, which predicts genes using Prodigal v2.6.3^69^. The resulting genes were clustered into a comprehensive pan-genome at 95% nucleotide identity using Roary v3.13.0^70^ to generate a multi-fasta file containing gene sequences (Supplementary file 1) and a per-genome gene presence-absence table (Supplementary file 1). The pan-genome was annotated with eggNOG 5.0 using eggNOG-mapper v2^41, 71^. We used custom R scripts to parse the eggNOG output for genes predicted to encode enzymatic or transporter activities of interest (Supplementary file 1) based on gene names, KEGG numbers, KEGG reaction numbers, Clusters of Orthologous Groups (COGs), EC numbers, Transporter Classification (TC) numbers, and GO (Gene Ontology) terms, followed by manual curation of the initial search results and BLAST-based sequence review for genes with unclear annotations. We then determined the number of genomes from each species predicted to encode each gene function of interest. Since the Genome Standards Consortium criteria for high- and medium-quality genomes and MAGs allow for low-level sequence contamination within assemblies^65^, we restricted gene presence-absence analysis to gene sequences of interest that were detected in at least two genomes or MAGs from a species in order to exclude singleton contaminant sequences from our analysis. Importantly, including genomes and MAGs with completeness as low as 50% maximizes genome catalog diversity, thus increasing pan-genome size and sensitivity for detecting genes of interest within each species, but also results in a fraction of genomes and MAGs appearing to lack universally present genes due to incompleteness of assemblies.

### Maximum likelihood phylogenetic distances and phylogenetic reconstruction

Phylogenetic reconstruction of *L. iners* genomes and MAGs (Fig. 1b) was performed using assemblies with >60% completeness and <5% contamination to maximize robustness of the analysis. fetchMG v1.0^72^ was used to extract DNA sequences for each of forty single-copy universal bacterial marker genes from each genome (called using Prodigal^69^). Genome assemblies containing fewer than 10 of the 40 universal marker genes were omitted from the subsequent alignment. A phylogenetic reconstruction was then produced using ETE3 v3.1.1 (parameters: “*ete3 build -w clustalo_default-trimal-gappyout-none-none -m cog_85-alg_concat_default-fasttree_default*”)^73^, and the tree was visualized using iTOL v4^74^. An additional phylogenetic reconstruction encompassing other major FGT *Lactobacillus* species (Extended Data Fig. 2a) was performed using *L. iners* genomes and MAGs as well as genomes from *L. crispatus*, *L. jensenii*, *L. gasseri*, and *L. vaginalis*, filtered based on the same quality and completeness criteria.

### Preparation of bacterial inocula for growth experiments

For initial experiments identifying cysteine and cystine as key nutrient requirements for *L. iners* growth (Fig. 1a,c, Fig 4a, and Extended Data Fig. 1), bacteria from frozen stocks were plated on solid media, incubated for 3 days, then suspended in sterile, pre-reduced Dulbecco’s phosphate-buffered saline (PBS), adjusted to an OD600 of 0.3+/-0.05, and then inoculated into the indicated broth media formulations for measurement of growth kinetics. In subsequent growth experiments for *Lactobacillus* species, experimental bacterial inocula were prepared from liquid starter cultures in MRSQ broth with L-Cys (4 mM) that were incubated for 18-20 hours. The starter cultures were then pelleted by centrifugation for 10 min at 3716 x g, spent media was decanted, and bacteria were washed 2 times by re-resuspending in sterile, pre-reduced PBS followed by centrifugation to avoid carryover of nutrients from the original starter culture media. Washed bacteria were resuspended in PBS, adjusted to OD600 0.3 (± 0.05), then inoculated into experimental media at 3.5% (v/v), equating to bacterial titers ranging from ∼1×10^5^ to 1×10^6^ colony-forming units (C.F.U.) per mL, depending on the experimental species and strain.

### Growth kinetics quantification

We found that *Lactobacillus* growth kinetics in broth media were adversely affected by either continuous or intermittent agitation (not shown), therefore we grew broth cultures without agitation. Mono-culture growth kinetics were assessed using separate cultures prepared in parallel for each experimental timepoint to avoid repeatedly agitating a single culture by performing serial measurements. For timepoint series lasting *≤*48 hrs, broth cultures were grown in a volume of 250 μL in technical triplicate in clear 96-well flat bottom plates (Falcon), with a blank for each media condition per plate. Measurements were taken at ∼16, 20, 24, 28, ∼42, and 48 hours by removing plates from the anaerobic chamber, pipetting to re-suspend bacteria, and measuring OD600 on a Tecan Infinite*®* M1000 PRO plate reader. Due to slower growth kinetics in the HMRS broth formulation, *L. iners* growth measurements in HMRS were taken daily for up to 10 days, precluding use of 96-well plates due to evaporation at later timepoints. Bacteria were therefore incubated in low-evaporation 1.2 mL 96-well cluster tubes (Corning™) in triplicate, with separate parallel culture plates used for each experimental timepoint, then 250 μL from each culture was transferred to a clear 96-well flat-bottom Falcon plate for optical density measurement. Data from mono-culture growth experiments are depicted as median ± range for the three replicates. For each bacterial strain and media condition, figures depict representative results from 1 of *≥*2 independent experiments with distinct batches of freshly prepared media and freshly prepared bacterial input inocula unless otherwise indicated.

### Inhibitor experiments and analysis

To test mono-culture growth inhibition, bacteria were cultured in media containing inhibitors at varying concentrations as indicated, including a reference (no-inhibitor) control. Since growth kinetics differed between species and strains, inhibition was determined for each strain at the first timepoint fulfilling the European Commission on Antimicrobial Susceptibility Testing (EUCAST) criterion of “definite turbidity” in the reference control (which we defined experimentally as OD600 >0.2), unless otherwise specified. At the selected timepoint for each strain, the median OD600 value for the reference control was set to a reference value of 100%, which was used to calculate percentage growth in the other conditions. Growth experiments employing metronidazole (MTZ) used a concentration of 50 μg/mL, approximating the concentration in 0.75% intravaginal MTZ gel (a first-line treatment for BV)^21^. Data from inhibition experiments are depicted as median ± range for three technical replicates. For each bacterial strain and media condition, figures depict representative results from 1 of *≥*2 independent experiments with distinct batches of freshly prepared media and freshly prepared bacterial input inocula, unless otherwise indicated.

### Competition and mock community culture experiments

For pairwise competition between *L. iners* and *L. crispatus* strains in MRS broth ± SMC and for mock BV-like community experiments with *L. iners* (multiple strains), *L. crispatus* (multiple strains), *G. vaginalis* (ATCC 14018), *P. bivia* (ATCC 29303), and *A. vaginae* (0795_578_1_1_BHK4; see Supplementary Table 3) in S-broth ± SMC and/or MTZ, bacteria were initially prepared in individual suspensions in broth as described above for bacterial monoculture growth and inhibition experiments. Aliquots of the mono-bacterial suspensions were mixed in defined ratios, then mixtures were divided into replicate cultures for incubation (5 replicates per conditions for MRSQ experiments; 6 replicates per condition for S-broth experiments) by adding 150 μL of mixture per well into V-bottom 96-well plates (Falcon). C.F.U. titers were determined for each input mono-bacterial suspension as described above and used to calculate starting ratios within the mixed cultures. At 28 hours, cultures were harvested by centrifuging at 4700xg for 25 minutes at 4C, spent media was decanted, and pellets were frozen for later DNA extraction and analysis. Relative growth within mixed cultures was assessed by bacterial 16S rRNA gene sequencing as described below. Aliquots of mono-bacterial suspensions in the corresponding media type were cultured separately to confirm expected growth patterns (e.g. Fig. 5b and Extended Data Fig. 8a).

### Culture-independent bacterial 16S gene amplification and Illumina MiSeq sequencing

The V4 region of the bacterial 16S rRNA gene was PCR-amplified following standard protocols^4, 61, 75^. Samples were amplified using 0.5 units of Q5 High-Fidelity DNA Polymerase (NEB) in 25 μl reaction with 1X Q5 Reaction Buffer, 0.2 mM dNTPs (Sigma), 200 pM 515F primer and 200 pM barcoded 806R primer (IDT) in PCR-clean water (Invitrogen Ultra Pure DNase/RNase-Free Distilled water). A water-template negative control reaction was performed in parallel for each barcode master mix. Additional blank extraction and amplification controls performed using unique barcoded primers were also included in sequencing libraries. DNA from clinical swab samples for FGT microbiota profiling was amplified in triplicate reactions that were then combined prior to library pooling to minimize stochastic amplification biases. DNA from *in vitro* mock community experiments (for which each experimental condition had been cultured in multiple replicates) was amplified in a single reaction per replicate culture. Amplification was performed at 98°C for 30 seconds, followed by 30 cycles of 98°C for 10 s, 60°C for 30 s, and 72°C for 20 s, with a final 2 min extension at 72°C. PCR products were checked via agarose gel electrophoresis in parallel with the matching water-template control reactions to confirm successful target amplification and absence of background amplification. Gel band strength was used to semi-quantitatively estimate relative amplicon concentrations for library pooling. To prepare the sequencing libraries, 3-20 μl of individual PCR products (adjusted based on estimated relative amplicon concentration) were combined into 100 μl sub-pools and purified using an UltraClean 96 PCR Cleanup Kit (Qiagen). Blank extractions, water-template, and (for *in vitro* experiments) blank media controls were included in the sequencing libraries although they did not produce visible PCR bands. Concentration of the sub-pools were quantified using a Nanodrop (Thermo Scientific), then pooled at equal molar concentrations to assemble the final library. The pooled library was diluted and supplemented with 10% PhiX according to standard Illumina protocols, then single-end sequenced on an Illumina MiSeq using a v2 300-cycle sequencing kit with addition of custom Earth Microbiome Project sequencing primers^75^.

### Processing of Illumina MiSeq 16S gene sequences

Initial sequence demultiplexing was performed as previously described^61^, then dada2 version 1.6.0^76^ was used to filter and trim reads, infer sequences, and assign initial taxonomy employing the RDP training database rdp_train_set_16.fa.gz (obtained from https://www.mothur.org/wiki/RDP_reference_files). Taxonomic assignments were refined and extended via extensive manual review (see Supplementary Table 7 for amplicon sequence variant (ASV) taxonomy). The denoised dada2 results with final taxonomic assignment were analyzed in R using phyloseq version 1.30.0^77^ and custom R scripts (available at gitlab***).

### 16S rRNA gene-based microbiome analysis

For 16S-based microbiome profiling of clinical specimens, microbial communities were classified into four cervicotypes (CTs) as previously defined in a non-overlapping subset of participants from the FRESH cohort: CT1 includes communities with >50% relative abundance of non-*iners Lactobacillus* species (which consists almost entirely of *L. crispatus* in this population^4^); CT2 consists of communities in which *L. iners* is the most dominant taxon; CT3 consists of communities in which the genus *Gardnerella* is the most dominant taxon; and CT4 consists of communities dominated by other species, typically featuring high abundance of one or more *Prevotella* species^1^. Although *L. jensenii* and *L. gasseri* were detected in microbial communities of some FRESH study participants by both sequencing and isolation, they were not dominant in any individuals, a common finding among studies of sub-Saharan African populations^1, 4, 6, 54^ that contrasts with observations in most North American and European cohorts, where a consistent minority of women tend to have *L. gasseri*-dominant or *L. jensenii*-dominant communities^11, 16, 17, 67^. For further sequence processing and analysis, amplicon sequence variants (ASVs) that could not be defined at least to the level of taxonomic class were pruned from the dataset and one sample was excluded due to sequence read count <10,000. The remaining 142 samples were rarefied without replacement to a uniform depth of 16,603 reads (the minimum read count among remaining samples), yielding a relative abundance limit of detection (L.D.) of 6.02 x 10^-5^. ASVs were collapsed at the species or genus level as indicated for further visualization and statistical analysis. In plots displaying taxon relative abundance using a logarithmic scale, a pseudocount equal to 0.5 x L.D. was added to assist visualization of any taxon with read count 0. Correlation analysis between bacterial relative abundances and cervicovaginal metabolite concentrations was restricted to taxa detected (*≥*1 sequence read after rarefaction) in *≥*50% of samples (Extended Data Fig. 7).

### Sequence analysis for *in vitro* competition experiments and mock communities

For pairwise competitions between *L. crispatus* and *L. iners*, 16S rRNA gene sequences were generated, processed, and annotated as described above. Ratios of *L. iners* reads to *L. crispatus* reads in each sample were calculated. Significance of between-group differences for each mixture was determined by 1-way ANOVA. Significance of pairwise differences between the positive control condition (MRSQ + L-Cys without inhibitor) and each experimental condition was calculated using Dunnett’s test and all significant pairwise comparisons were plotted (see Supplementary Table 10 for numeric p-values). Results of representative competition experiments are shown in Fig. 5a.

For mock BV-like community experiments, 16S rRNA gene sequences were generated, processed, and annotated as described above. Relative abundances of each experimental strain were determined for individual replicates and displayed in Fig. 5d. To assess *L. crispatus* enrichment, read counts of all experimental strains except *L. crispatus* were summed for each replicate sample, then ratios of *L. crispatus* to the summed taxa were calculated. Significance of between-group differences for each mixture was determined by 1-way ANOVA, and significance of pairwise comparisons were calculated using Tukey’s test, with significance of selected pairwise comparisons plotted in Fig. 5e and shown in detail in Supplementary Table 11.

### Measurement and analysis of metabolites in cervicovaginal lavage fluid

Concentrations of Cys, reduced glutathione (GSH, *γ*-L-glutamyl-L-cysteinyl-glycine), and cysteinylglycine (Cys-Gly) were measured in CVL supernatants from the 143 FRESH cohort participants whose microbiome composition was concurrently profiled by 16S gene sequencing in this study. CVL supernatant samples underwent a single freeze-thaw cycle during preparation of aliquots for metabolite analysis. Metabolites of interest were quantified using ultraperformance liquid chromatography/tandem mass spectrometry (UPLC-MS/MS) by Metabolon, Inc., as part of an untargeted metabolomics dataset (Bloom, Abai & Kwon, unpublished data). To avoid bias in assay performance, measurements were performed after randomized reordering of samples by laboratory staff at Metabolon, Inc., who were blinded to microbiome composition of the donor participants. Metabolon’s method generates relative instead of absolute metabolite concentrations. For analysis, concentrations were volume-normalized and median values were adjusted to 1, then percent missingness (reflecting samples with analyte concentrations below the limit of detection, or L.D.) was calculated for each analyte. For analytes with missingness >0%, analyte limit of detection was inferred as equaling the lowest measured relative concentration in the cohort, then missing values were imputed at half the limit of detection. The Shapiro-Wilk test of normality was then performed on relative concentration values after log-transformation (not shown). In subsequent analysis of concentration differences between cervicotypes, differences were analyzed by 1-way ANOVA with post-hoc Tukey test for Cys (which had 0% missingness and was normally distributed, Fig. 3a,c) and by Kruskal-Wallis test with post-hoc Dunn’s test for GSH and Cys-Gly (which had 21.7% and 7.0% missingness respectively and were therefore not normally distributed after imputation, Extended Data Fig. 4a-d). We calculated Spearman rank-order correlations between individual metabolites and bacterial taxa at both the genus and species levels, adjusting p-values for multiple comparisons where indicated using the Bonferroni method via the R stats package function p.adjust(). The R package DescTools was used to calculate confidence intervals for the Spearman correlation coefficients (*ρ*), including Bonferroni-corrected confidence levels at (1 - 0.05/n), where n represents the number of taxa (Fig. 3e,f, Extended Data Fig. 4e,f, and Supplementary Tables 8 & 9).

### Quantification of oxidized and reduced cysteine and glutathione in MRS

Oxidized and reduced glutathione and cysteine were quantified by UPLC-MS/MS (Waters Acquity/TQ-S), according to a modification of the protocol by Sutton et al^78^. Briefly, 100 µL anaerobic broth was mixed with 90 µL buffer (100 mM ammonium bicarbonate, pH 7.4), followed by 10 µL fresh *N*-ethylmaleimide solution (25 mg/mL in ethanol), and allowed to react at room temperature for 10 minutes. Samples were then diluted 100-fold in water before analysis by UPLC-MS/MS, with a 1 µL injection volume. Separation was carried out with an Acquity BEH/C18 UPLC column (Waters 186002350), with the following gradient: 0 minutes, 2% B; 6 minutes, 98% B; 7 minutes, 98% B; 7.1 minutes, 2% B; 9 minutes, 2% B. Mobile phase A was water with 0.1% formic acid, and B was acetonitrile with 0.1% formic acid. Compounds were quantified by comparison of peak area to authentic standards processed via the same method.

### Synthesis and analysis of isotopically labeled cystine, GSSG and GSH

^13^C-labeled cystine (1,1’-^13^C_2_-L-cystine) was prepared from 1-^13^C-L-cysteine (Cambridge Isotope Labs, CLM-3852-0.5) via a modification of the procedure of Hill, Coy, and Lewandowski^79^. Briefly, 100 mg of labeled cysteine was mixed with 200 µL 10% NaOH, and 800 µL 3% hydrogen peroxide. The resulting precipitate was filtered and washed extensively with ice cold water, then finally redissolved in 1 M aqueous HCl. Complete oxidation was confirmed by UPLC-MS/MS analysis of the resulting solution, as described above.

Labeled glutathione-(1-^13^C-cysteine) was prepared enzymatically using *E. coli* GshA and GshB overexpressed and purified as N-terminal 6xHis fusions. *gshA* and *gshB* were amplified from genomic DNA of *E. coli* BL21(DE3) using Q5 polymerase (New England Biolabs) and the following primers: HIS-gshA-F, gcctggtgccgcgcggcagcATCCCGGACGTATCACAG; HIS-gshA-R, cagcttcctttcgggctttgTCAGGCGTGTTTTTCCAGCC; HIS-gshB-F gcctggtgccgcgcggcagcATCAAGCTCGGCATCGTGAT; and HIS-gshB-R, cagcttcctttcgggctttgTTACTGCTGCTGTAAACGTGC, according to the manufacturer’s instructions. The expression vector backbone pET28 was amplified from purified stock using primers: pET28-F, GCTGCCGCGCGGCACCAG; and pET28-R, CAAAGCCCGAAAGGAAGCTG. After DpnI digest and purification, expression plasmids pETgshA and pETgshB were assembled using HiFi Assembly MasterMix (NEB), and transformed into chemically competent *E. coli* DH5a. Positive colonies were verified via Sanger sequencing, and correctly assembled plasmids were transformed into *E. coli* BL21(DE3). Cells for protein expression were inoculated from overnight culture with 100-fold dilution into 4 L flasks containing 1 L LB broth, and grown at 37°C and 200 RPM until reaching OD600 ∼ 0.5, after which they were induced with 0.5 mM IPTG, and grown for a further 3 hours at 37 ⁰C. Pellets were harvested by centrifugation at 6,000 g for 10 minutes, then resuspended in ice cold 98% Buffer A (50 mM HEPES, 300 mM KCl, 10% glycerol, pH 7.5) and 2% Buffer B (Buffer A with 500 mM imidazole). Cells were lysed by three passages through an Emulsiflex C5 (Avestin) at 15,000 psi, and lysates were clarified by centrifugation at 20,000 g, 4°C for 30 minutes. Target proteins were bound to 1 mL Ni-NTA His-Bind resin (Millipore Sigma) in gravity columns, then washed with 5 mL wash buffer (90% Buffer A, 10% Buffer B) and eluted with 3 mL elution buffer (100% Buffer B). Fractions containing purified proteins were identified via SDS-PAGE, then combined and dialyzed overnight against Buffer A with 3.5k MWCO Slide-A-Lyzer Cassettes (ThermoFisher), then concentrated by centrifugation through 3k MWCO Microsep spin filters (Pall).

Glutathione (*γ*-L-glutamyl-L-cysteinyl-glycine) synthesis was carried out in 100 mM sodium phosphate buffer (pH 7.2), with 80 mM 1-^13^C-L-cysteine, 120 mM glycine, 120 mM glutamic acid, 40 mM MgCl_2_, and 100 mM ATP, using 50 mg 1-^13^C-L-cysteine, 1 mg each of GshA and GshB, incubating overnight at 37°C. The resulting glutathione-(1-^13^C-cysteine) was treated with 3% hydrogen peroxide to produce the oxidized form of glutathione (GSSG) for purification via preparative HPLC on a Thermo Scientific Dionex UltiMate 3000 HPLC system with a Thermo Scientific Hypersil GOLD aQ C18 preparative column (20 x 20 mm, 5 μm) using a gradient from 100% A (water + 0.1% formic acid) to 70% B (acetonitrile + 0.1% formic acid). Fractions containing labeled GSSG (^13^C_2_-GSSG) were pooled and lyophilized. Analysis of the purified product was conducted by ^1^H NMR (400 MHz) in the Magnetic Resonance Laboratory in the Harvard University Department of Chemistry and Chemical Biology on a Jeol J-400, as well as by LC-MS/MS as described above, and by high-resolution LC-MS using an Agilent 6530 Q-TOF (confirming expected product characteristics as detailed in HMDB: https://doi.org/10.13018/BMSE000906). This analysis confirmed the authenticity of the desired product and complete conversion of cysteine, and also revealed contamination with ADP, which was not removed by preparative HPLC (Extended Data Fig. 7). This material was used without further purification. Reduced labeled glutathione (^13^C-GSH, 4 mM) for use in experiment measuring GSH uptake was produced from the oxidized form (^13^C_2_-GSSG) by treating a stock solution of 81.6 mM labeled GSSG with TCEP at a 0.9:1 molar ratio to reduce the disulfide bonds prior to addition to MRSQ broth.

### Quantification and isotopic analysis of amino acids

For isotopic analysis of proteinogenic amino acids, *L. crispatus* (strain 233) and *L. iners* (strain F1) grown in MRS/Q broth supplemented with isotopically labeled or unlabeled substrates were harvested via centrifugation. The pellets washed 3 times with water and resuspended in 0.6 mL of 6 N aqueous HCl and heated at 100°C overnight to facilitate cell lysis, protein hydrolysis, and Cys oxidation. The hydrolysate was dried under air, then resuspended in 100 µL of 90% acetonitrile, 10% water. For determination of amino acid concentration in media, samples were prepared by diluting media 10-100 fold in 90:10 acetonitrile:water, followed by centrifugation to remove precipitated material. All amino acid samples were analyzed via UPLC-MS/MS using instrumentation described above. Separation was carried out with an Acquity BEH/Amide UPLC column (Waters 186004800) in HILIC mode, with the following gradient: 0 minutes, 90% B; 0.3 minutes, 73% B; 1 minute, 73% B; 1.5 minutes, 30% B; 2 minutes, 30% B; 2.2 minutes, 90% B; 4 minutes, 90% B. Mobile phase A was water with 0.1% formic acid, and B was acetonitrile with 0.1% formic acid. Quantification of amino acids was confirmed by comparison of peak area to authentic standards (Sigma # AAS18-5ML) processed via the same method. Isotopic distribution data were corrected for the natural abundance of ^13^C using IsoCor v2.1.3^80^ (Supplementary Tables 5 & 6).

### Statistics, software, and visualization

Data analysis, statistics, and visualization were performed in R v3.6.3, except where otherwise indicated, using packages including seqinr v4.2.5, tidyverse, v1.3.1, knitr v1.33, ggpubr v0.4.0, DescTools v0.99.41, gtools v3.8.2, gridExtra v2.3, cowplot v1.1.1, scales v1.1.1, grid v3.6.3, broom v0.7.6, e1071 v1.7.6, and table1 v1.4. All p-values are two-sided with statistical significance defined at *α* = 0.05 unless otherwise indicated.

### Data and code availability

Raw and corrected cystine and serine isotopologue measurements associated with Fig 2c,d and 4c are supplied in Supplementary Tables 5 and 6. Raw read data for genital tract 16S rRNA-gene profiling are available in the NCBI Sequence Read Archive (SRA) under BioProject PRJNA729907. Custom R code with associated data files sufficient to reproduce the analysis for pan-genome gene content analysis and for 16S-based microbiota-metabolite analysis are available as supplementary files (Supplementary Files 1 and 2 respectively). Each file contains a compressed directory with a README.txt file describing dependencies, an R Project file, an R Markdown file containing the analysis code, and associated sub-directories with the data used in the analysis. Taxonomic assignments used for amplicon sequence variants (ASVs) from bacterial 16S rRNA gene sequencing are supplied in Supplementary Table 7.

## Acknowledgments

We thank study participants for donating clinical samples used in this study; study staff at the FRESH cohort; laboratory staff at the HIV Pathogenesis Programme at UKZN for sample processing; L. Froehle for helpful discussions of analysis; D. Jenkins, M. Farcasanu, K. Jackson, L. Froehle, and J. Bramante for assay and sample assistance. Some sequencing data and experimental strains were kindly provided by J. Ravel and the Vaginal Microbiome Research Consortium (VMRC). Some strains of non-*iners* lactobacilli used for genomic analysis were a kind gift of J. Marrazzo. This work was supported by National Institutes of Health (NIH) grants NIH 1R01AI111918-01 to D.S.K., HU CFAR NIH/NAIDS 5P30AI060354-15 to S.M.B., and HU CFAR NIH/NAIDS P30-AI060354 to M.S.G., by Bill and Melinda Gates Foundation grants OPP1189208 to D.S.K. and OPP1158186 to E.P.B., and by Vincent Memorial Research Funds and a Domolky Innovation Award to C.M.M. (Massachusetts General Hospital). D.S.K. was supported by the Burroughs Wellcome Career Award for Medical Scientists. S.M.B. was supported in part by NIH grant T32 AI007387, T.N. was supported by the South African Research Chairs Initiative through the National Research Foundation and the Victor Daitz Foundation, A.B.A. was supported by funding from the Harvard Program for Research in Science and Engineering (PRISE) and the Harvard Microbial Sciences Initiative, and X.W. was supported by the Ragon Institute Summer Program Fellowship.

## Author Contributions

S.M.B. and D.S.K. conceived the overall study and guided it throughout with input from B.M.W., E.P.B., and C.M.M.; S.M.B., N.A.M., and J.K.R. performed primary bacterial isolations; S.M.B., N.A.M., J.F.F., B.M.W., A.J.M., X.W., N.C., and C.M.M. contributed to media design and production and/or bacterial growth and inhibition experiments; B.M.W. and E.P.B. synthesized labeled glutathione; B.M.W., S.M.B., N.A.M., and E.P.B. designed, performed, and/or analyzed measurements of media composition and isotopic tracing experiments; S.M.B., N.A.M., and J.X. performed nucleic acid extractions and sequencing; S.M.B. performed bacterial 16S rRNA gene sequencing analysis; M.R.H. performed genomic and metagenomic sequence analysis and assembly, genome catalog development, and phylogenetic reconstructions; S.M.B., M.R.H., F.A.H., and B.M.W. conceived and/or performed genomic pathway analysis; S.M.B. and A.B.A. performed analysis of *in vivo* metabolite data; K.L.D., M.D., T.G., X.S., T.N., N.I., S.M.B., N.X., M.S.G., and D.S.K. contributed to clinical trial design, trial performance, and/or sample acquisition and processing efforts; S.M.B., B.M.W., M.R.H., N.A.M., and D.S.K. wrote the paper, and all authors reviewed, offered input to the writing, and approved the manuscript.

---

**Extended Data Fig. 1.**
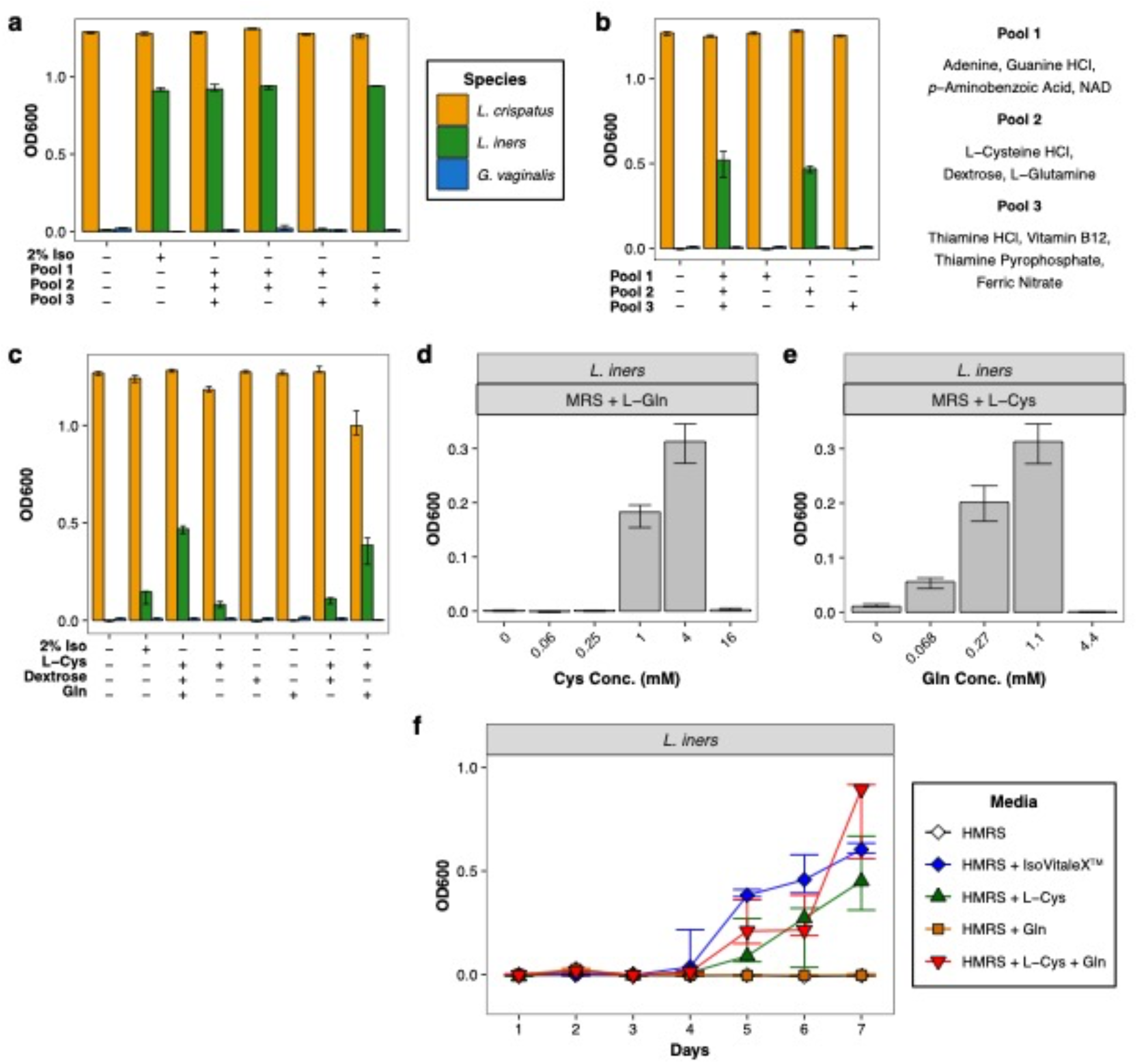
L-cysteine supplementation supports *L. iners* growth in *Lactobacillus* MRS broth, augmented by L-glutamine. (**a & b**) Growth of *L. crispatus*, *L. iners*, and *G. vaginalis* at 24 hours incubation in MRS broth (BD DIFCO™) ± supplementation with 2% IsoVitaleX*™* (“Iso”) or with the indicated sub-pools of IsoVitaleX*™* components. (**c**) Growth in MRS broth supplemented with 2% IsoVitaleX*™* or various combinations of the nutrients in “Pool 2”. (**d**) Growth of *L. iners* at 24 hours in MRS broth + L-Gln (1.1 mM) supplemented with varying concentrations of L-Cys or (**e**) in MRS broth + L-Cys (4 mM) supplemented with varying concentrations of L-Gln. Experiments in (**a-e**) all used BD DIFCO™-formulated MRS broth base. (**f**) Growth of *L. iners* in Hardy Criterion-formulated MRS (“HMRS”) broth supplemented with IsoVitaleX*™* 2% v/v, L-Cys (4 mM), and/or L-Gln (1.1 mM) produced similar results, although with a substantially longer lag phase. Each plot is from 1 of *≥*2 independent experiments per strain and media condition except (**a**), which was un-replicated.

**Extended Data Fig. 2.**
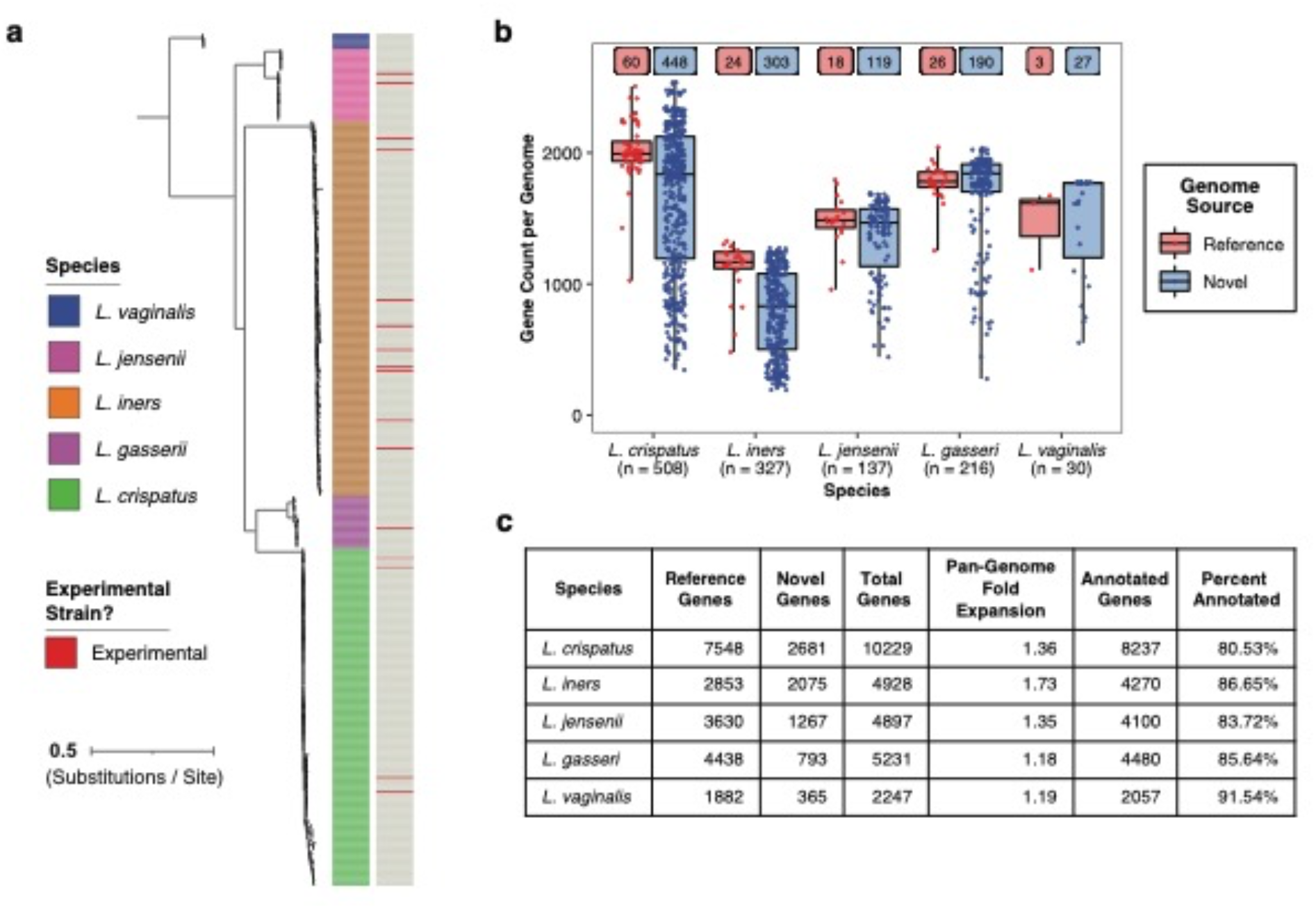
Phylogeny, gene content, pan-genome size, and novelty of *Lactobacillus* genome catalogs: **(a)** Phylogenetic tree of *L. iners* isolate genomes and MAGs as in Fig 1b, plus genomes of *L. crispatus* (n = 182), *L. jensenii* (n = 39), *L. gasseri* (n = 28), and *L. vaginalis* (n = 8). Isolates further experimentally studied in this work are indicated. The tree depicts only genome assemblies exceeding certain quality thresholds to ensure robustness of the phylogenetic reconstruction (see **Methods**); additional strains and genomes were included in other analyses. **(b)** Number of genes per genome within the *Lactobacillus* genome catalogs analyzed in Fig. 2b and 4b and Extended Data Fig. 6a. To maximize comprehensiveness of the pan-genomes, the catalogs included both high- and medium-quality genomes and MAGs with minimum estimated completeness >50% and maximum estimated contamination <10%. Gene count analysis excludes gene sequences observed in only 1 genome or MAG per species to eliminate singleton contaminating sequences within individual genome assemblies. **(c)** Total species pan-genome size, number of non-singleton genes uniquely contributed by novel isolate genomes and MAGs, degree of pan-genome expansion due to genes from novel genomes and MAGs, and number (percent) of genes within each pan-genome annotated by eggNOG.

**Extended Data Fig. 3.**
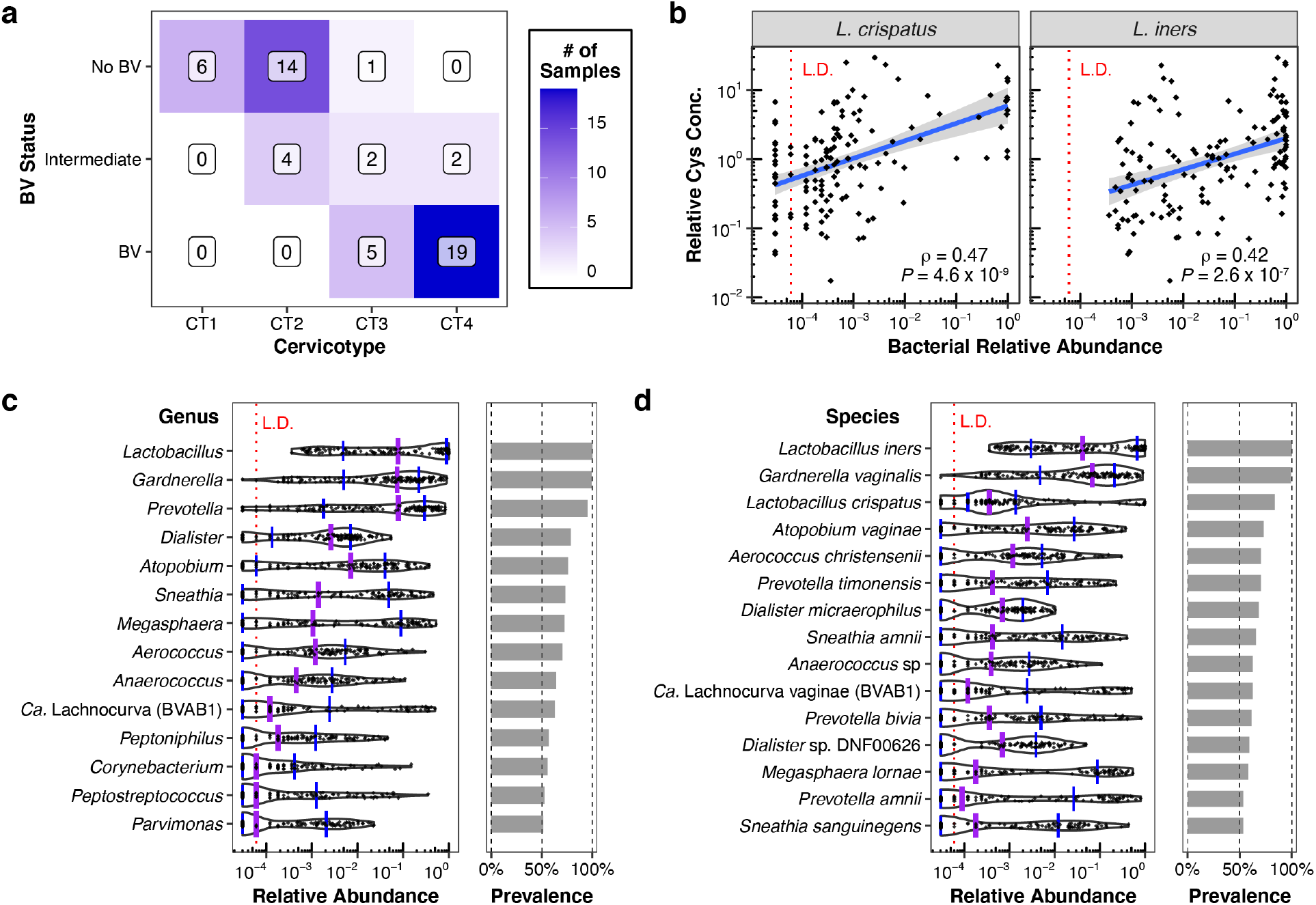
*In vivo* association of BV status and vaginal Cys concentrations with microbiota composition: **(a)** Relationship between Nugent score-based BV status^45^ and cervicotype among the 53 women depicted in Fig. 3a. BV status and cervicotype were significantly associated (p = 1.902 x 10^-11^; two-sided Fisher’s Exact Test). (**b**) Spearman correlation between relative Cys concentrations in cervicovaginal lavage (CVL) fluid and relative abundances of the species *L. iners*, and *L. crispatus* among the 142 women depicted in Fig. 3b-f, plotted and analyzed as in Fig. 3d. The red dotted line represents the bacterial limit of detection (L.D.). **(c** & **d)** Per-sample relative abundances and cohort-level prevalence (fraction of samples from the cohort in which each taxon was detected) for each genus (**c**) or species (**d**) with *≥*50% prevalence (panels correspond to main Fig. 3e and 3f, respectively). Purple and blue lines respectively represent median and interquartile range of relative abundances for each taxon.

**Extended Data Fig. 4.**
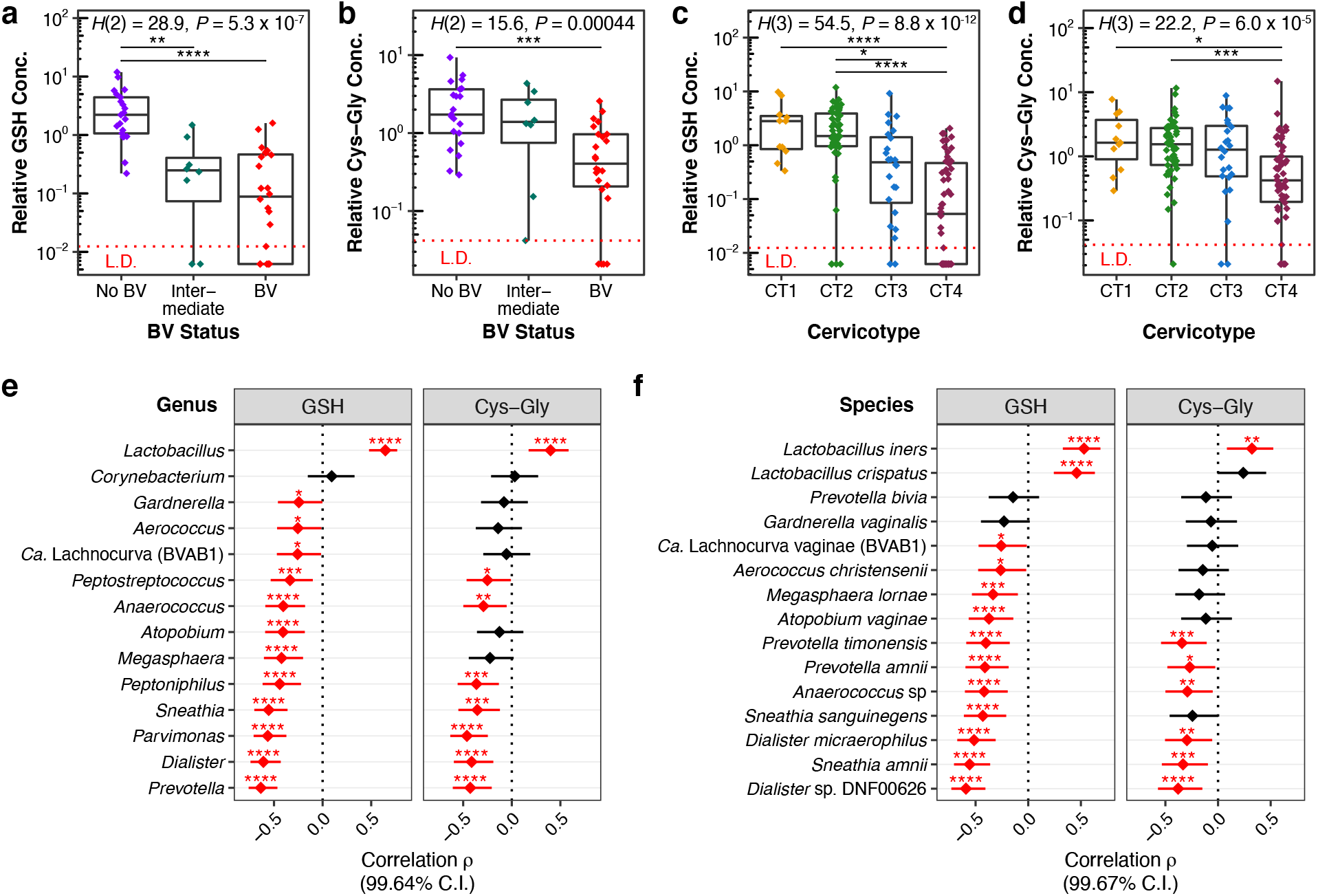
Vaginal concentrations of the Cys-containing peptides reduced glutathione (GSH) and cysteinylglycine (Cys-Gly) in cervicovaginal fluid are higher in women without BV and correlate with Lactobacillus dominance of the microbiota: **(a & b)** Relative concentrations of **(a)** GSH and **(b)** Cys-Gly by BV status in CVL fluid from the 53 women in (Fig. 3a). **(c & d)** Relative concentrations of **(c)** GSH and **(d)** Cys-Gly by cervicotype in CVL fluid from the 142 women depicted in (Fig. 3b,c). In **(a-d)** the red dotted line represents the metabolite limit of detection (L.D.). For samples in which an analyte was below the L.D., concentrations were imputed at 0.5 x L.D. Log-transformed concentrations were not normally distributed due to the imputed values, so between-group differences were determined via Kruskal-Wallis test with post-hoc Dunn’s test, adjusting for multiple comparisons using the Bonferroni method. All significant pairwise differences are displayed. (**e & f**) Forest plots depicting Spearman correlation coefficients (*ρ*) between concentrations of GSH and Cys-Gly and relative abundances of each bacterial genus **(e)** or species **(f)** detected at >50% prevalence in the cohort. P-values and confidence intervals for *ρ* were calculated and portrayed as for (Fig 3e,f) (full statistical results in Supplementary Tables **8 & 9**). Significance is depicted for adjusted p-values as * p *≤* 0.05, ** p *≤* 0.01, *** p *≤* 0.001, **** p *≤* 0.0001.

**Extended Data Fig. 5.**
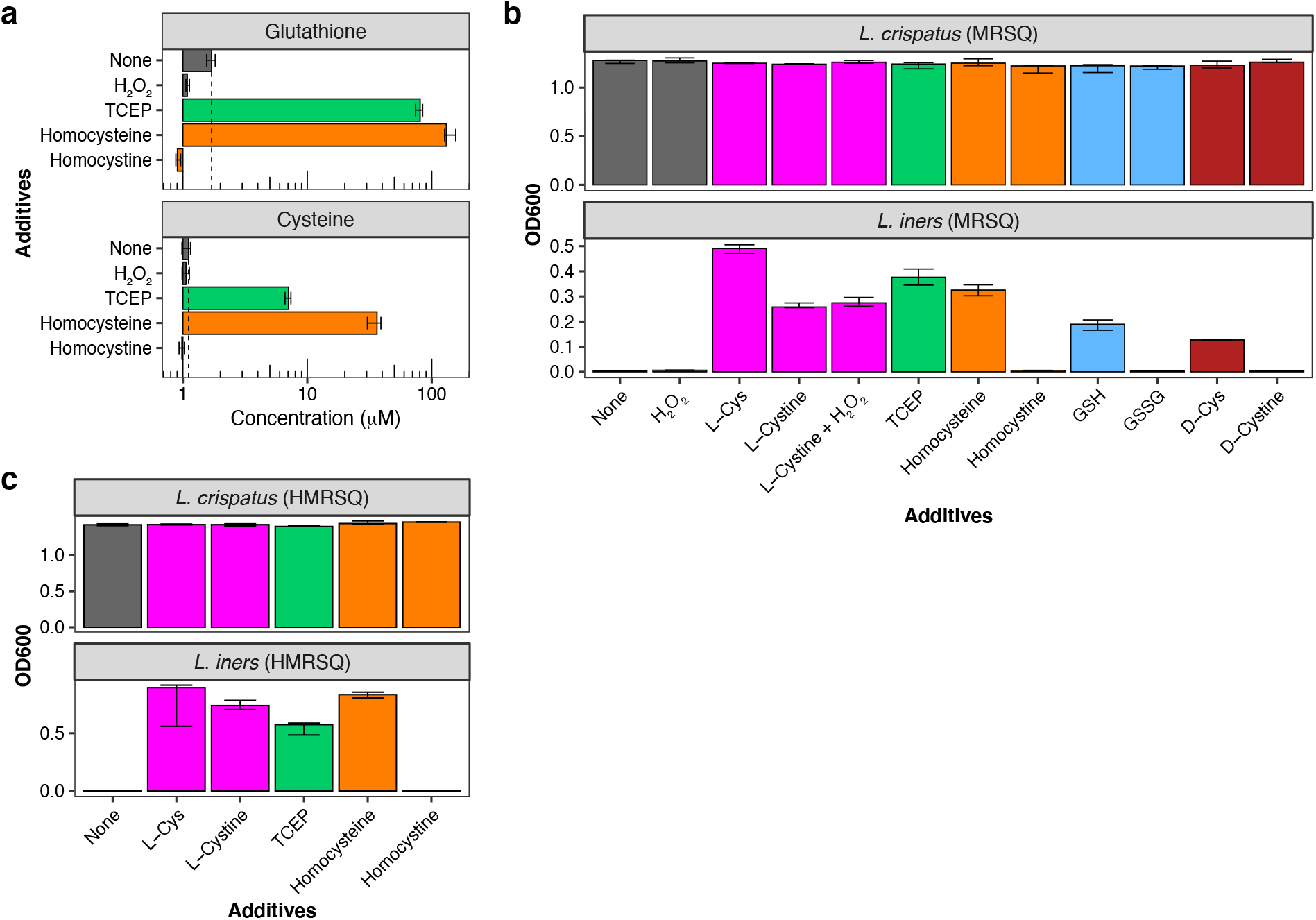
Cys and Cys-containing molecules in MRS exist primarily as mixed disulfides and addition of chemical reducing agents permits *L. iners* growth. **(a)** Concentrations of reduced Cys (baseline median concentration 1.11 μM) and glutathione (GSH; baseline median concentration 1.70 μM) in MRSQ broth supplemented with the oxidizing agent H_2_O_2_ (0.4 mM), the non-sulfur-containing reducing agents Tris(2-carboxyethyl)phosphine (TCEP) or homocysteine (each 4 mM), or homocysteine’s oxidized counterpart homocystine (2 mM). Plots depict median ± range for 3 replicates per condition**. (b)** Growth of *L. iners* and *L. crispatus* at 28 hours in MRSQ broth supplemented as indicated with the reduced thiols L-Cys, D-Cys (the non-physiological enantiomer of L-Cys), GSH, or homocysteine (each 4 mM), with their oxidized counterparts L-cystine, D-cystine, oxidized glutathione (GSSG), or homocystine (each 2 mM), or with TCEP (4 mM), H_2_O_2_ (0.4 mM), or L-cystine + H_2_O_2_. (**c**) Growth at 7 days of *L. crispatus* and *L. iners* in HMRS broth with 1.1 mM L-Gln (“HMRSQ”) supplemented as indicated with L-Cys, L-cystine, TCEP, homocysteine, or homocystine at the above concentrations. Bar coloring highlights the pairing of media conditions with each thiol-containing reducing agent and its oxidized counterpart.

**Extended Data Fig. 6.**
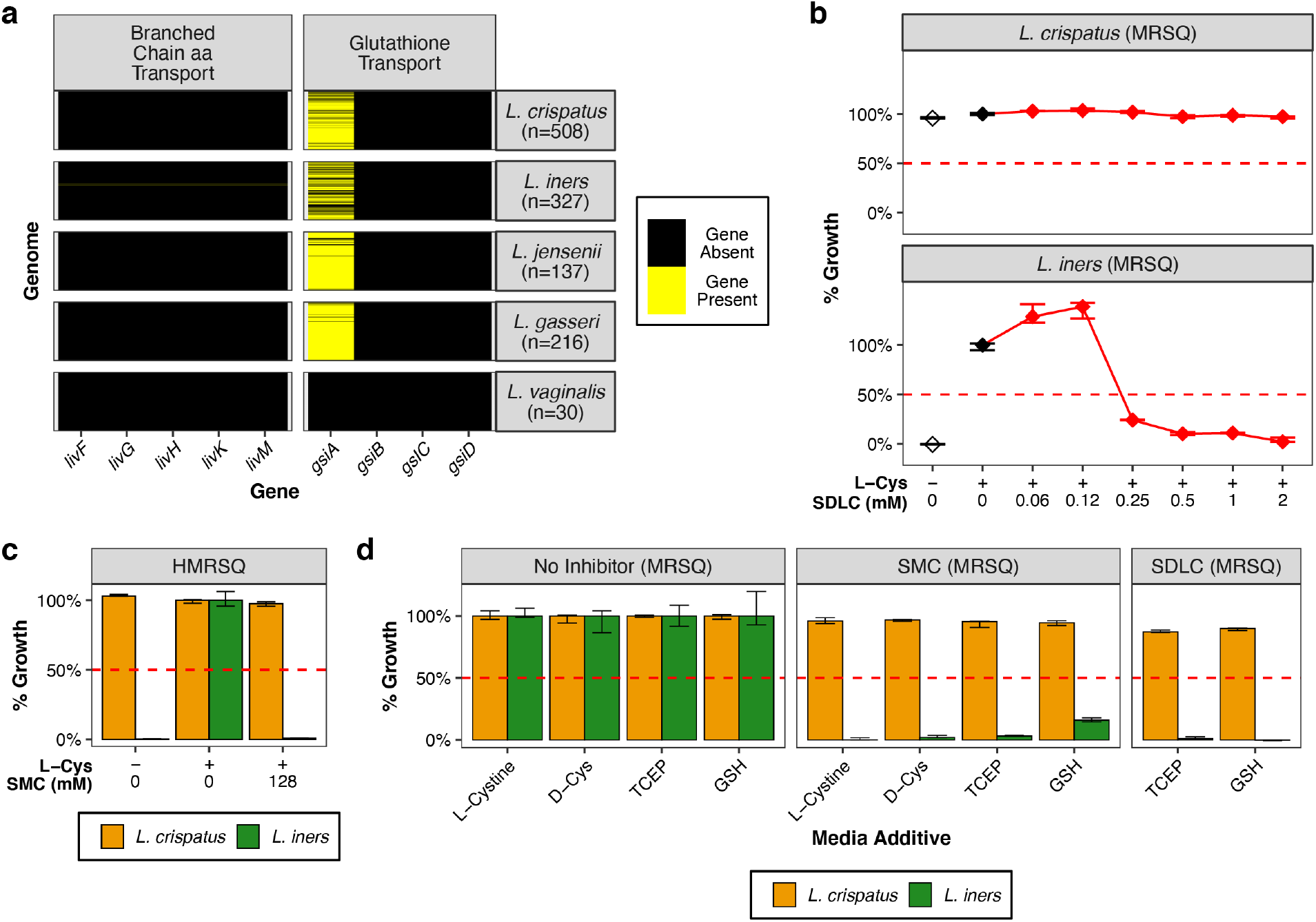
FGT *Lactobacillus* genomes lack predicted alternate Cys and GSH transporters and *L. iners* is selectively inhibited by cystine uptake inhibitors. **(a)** Predicted presence of the branched-chain amino acid transport locus *livFGHKM* (which has low-affinity Cys transport activity in *E. coli*^47^) and the glutathione transport locus *gsiABCD* in isolate genomes and MAGs of common FGT *Lactobacillus* species (n = number of genomes. Detailed statistics are in Supplementary Table 4). **(b)** Selective growth inhibition of *L. iners* in MRSQ broth with or without L-Cys (4 mM) and varying concentrations of the cystine uptake inhibitor seleno-DL-cystine (SDLC). **(c)** Growth of *L. crispatus* and *L. iners* in HMRSQ broth with or without L-Cys (4 mM) ± SMC. **(d)** Growth inhibition of *L. crispatus* and *L. iners* by SMC (128 mM) or SDLC (2 mM) in MRSQ supplemented with L-cystine (2 mM) or D-Cys, TCEP, or GSH (4 mM each). For each growth additive, percentage growth in in presence of inhibitor was calculated relative to median growth in broth containing that additive without inhibitor.

**Extended Data Fig. 7.**
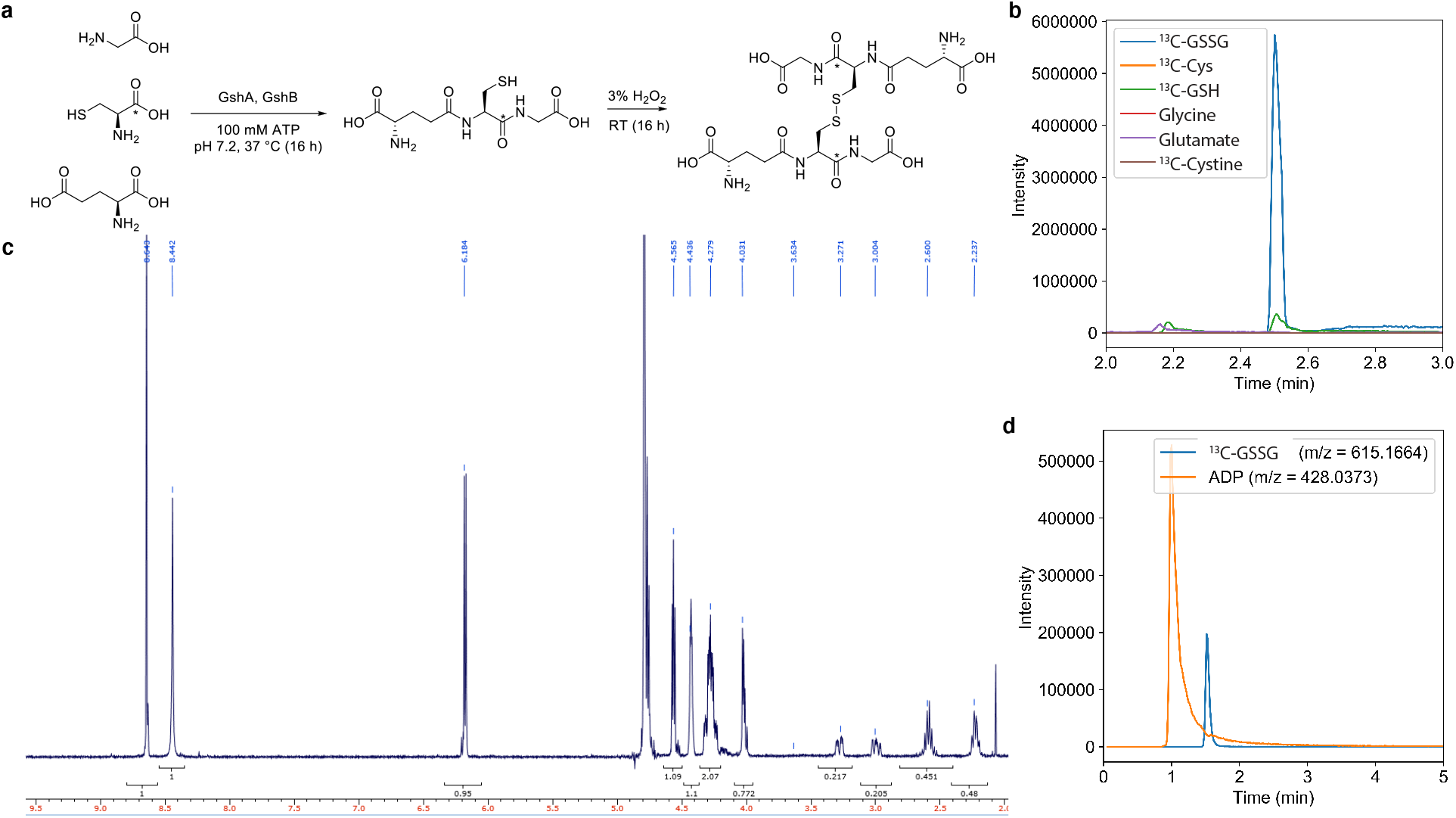
Synthesis of labeled glutathione. (**a**) Synthetic scheme for ^13^C-labeled glutathione starting from glycine, 1-^13^C-L-Cys, and L-glutamic acid to enzymatically produce ^13^C-GSH, followed by an oxidation step to produce oxidized glutathione (^13^C_2_-GSSG) for purification. (**b**) LC-MS/MS, (**c**) ^1^HNMR, and (**d**) high-resolution LC-MS analysis of purified product.

**Extended Data Fig. 8.**
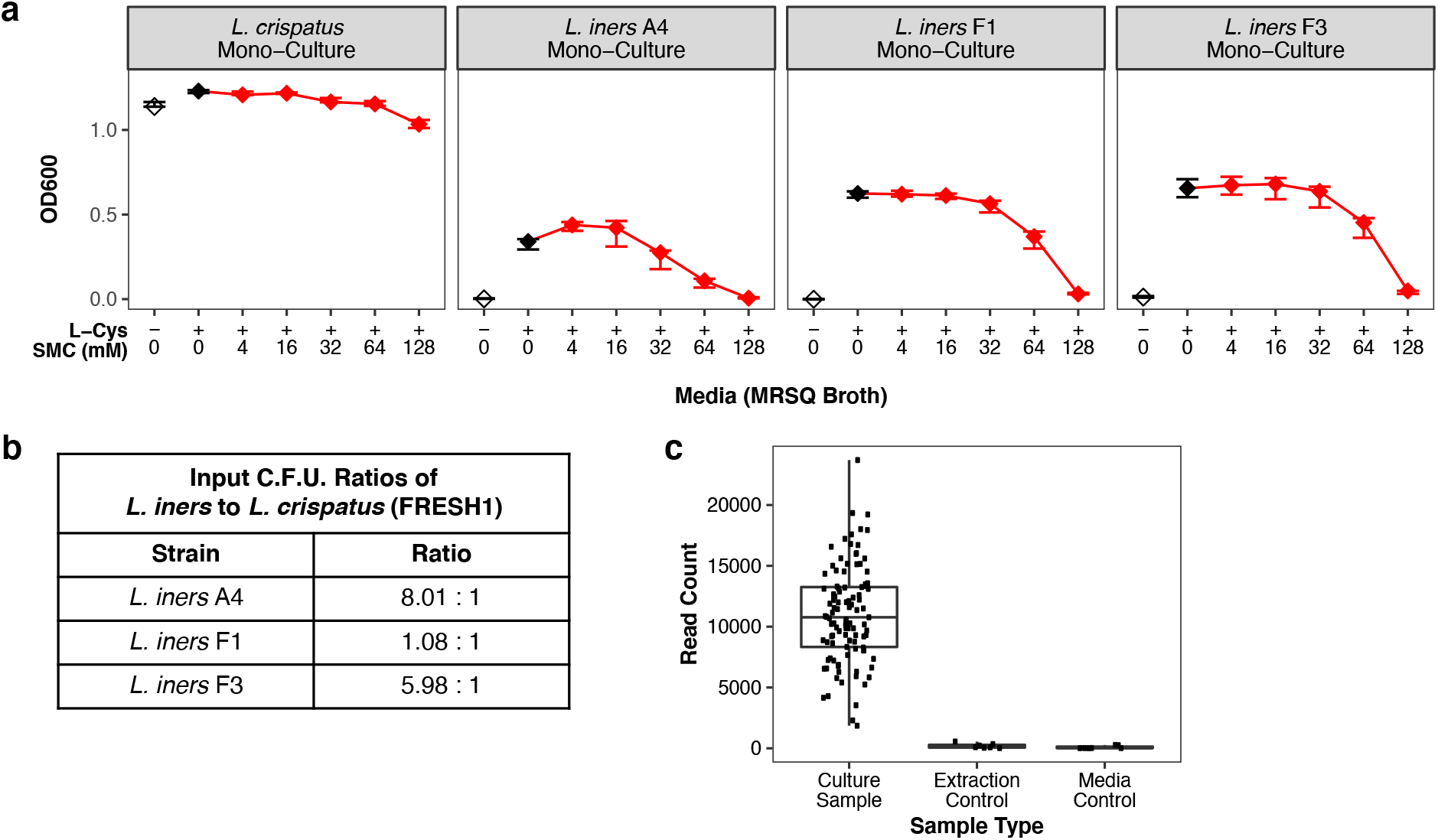
Sample preparation for *L. iners* / *L. crispatus* growth competition assays: **(a)** Mono-culture growth of *L. crispatus* (strain FRESH1) and 3 representative *L. iners* strains at 28 hrs incubation in MRSQ broth with or without L-Cys (4 mM) and varying concentrations of the cystine uptake inhibitor *S*-methyl-L-cysteine (SMC), exhibiting expected growth patterns for the respective species. The competition assays between *L. iners* and *L. crispatus* depicted in Fig. 5a were prepared by mixing the input inocula from each of these *L. iners* monocultures pairwise with the input inoculum for the *L. crispatus* monoculture at the colony-forming unit (C.F.U.) ratios shown in **(b)**. **(c)** Negligible bacterial 16S rRNA gene background amplification in blank growth media controls and extraction controls associated with the competition experiments in main Fig. 5a. (Control samples reactions were included in the sequencing library despite absence of visible PCR bands.)

**Extended Data Fig. 9.**
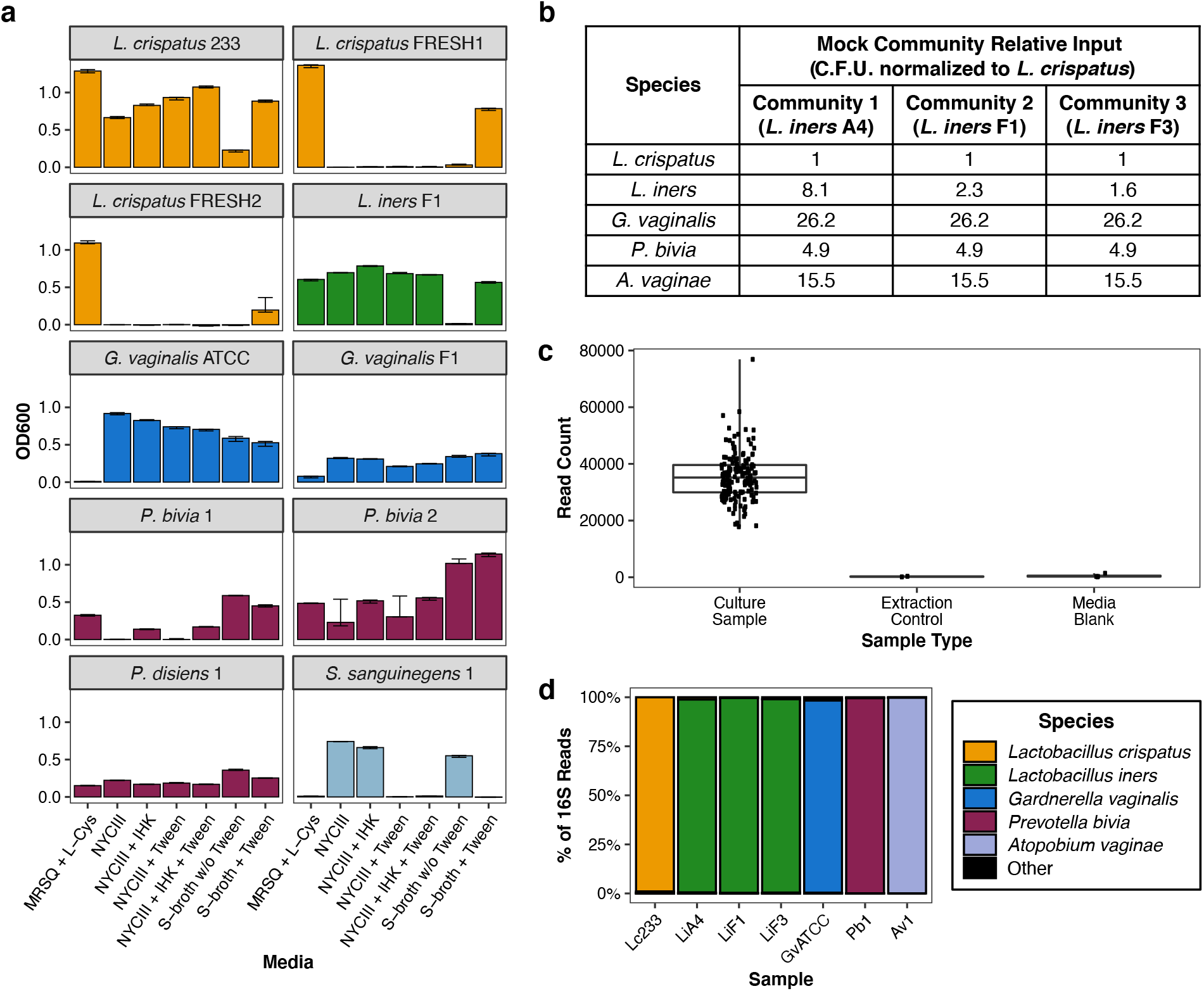
Development of “S-broth” and controls for mock BV-like community growth experiments: **(a)** Monoculture growth at 48 hours of experimental US (strain “233”) and South African (strains FRESH1 and FRESH2) *L. crispatus* strains, as well as *L. iners*, *G. vaginalis*, *Prevotella bivia*, *Prevotella disiens*, and *Sneathia sanguinegens* strains in various broth media including MRSQ broth + L-Cys (4 mM), NYCIII broth with or without 2% IsoVitaleX*™* plus 5% Vitamin K1-Hemin solution (“IHK”) and/or Tween-80 (1 g/L), and in “S-broth” (see **Methods**) with or without Tween-80 (1 g/L). S-broth + Tween was used in subsequent experiments. (Detailed strain information is in Supplementary Table 3) **(b)** Composition and C.F.U. input ratios of mock communities in main Fig. 5c,d, including the indicated strains of *L. iners*, *L. crispatus* strain 233, *G. vaginalis* strain “ATCC” (ATCC 14018), *P. bivia* strain 1, and *A. vaginae* strain 1. **(c)** Negligible bacterial 16S rRNA gene background amplification in blank growth media controls and extraction controls associated with the experiments in Fig. 5d,e. (Control samples reactions were included in the sequencing library despite absence of visible PCR bands.) **(d)** Confirmation of identity of input strains in mock BV-like communities by 16S rRNA gene sequencing of bacterial monoculture controls, prepared from the same input inocula used for the bacterial mock communities shown in Fig. 5c,d.

## Supplementary Tables

- Supplementary Table 1: Chemical and media reagents
- Supplementary Table 2: Media additive preparation
- Supplementary Table 3: Experimental bacterial strains
- Supplementary Table 4: Prevalence of predicted cysteine-related biosynthetic, metabolic, and transport genes within collections of isolate genomes and metagenomically assembled genomes (MAGs) from major cervicovaginal *Lactobacillus* species
- Supplementary Table 5: Raw cystine and serine isotopologue measurements (associated with Fig. 2c,d and 4c)
- Supplementary Table 6: Cystine and serine isotopologue measurements corrected for natural isotopic abundances using IsoCor (associated with Fig. 2c,d and 4c)
- Supplementary Table 7: 16S rRNA gene amplicon sequence variant (ASV) taxonomy assignments (associated with Fig. 3 and Extended Data Fig. 3, 4)
- Supplementary Table 8: Genus/metabolite correlation statistics associated with Fig. 3e and Extended Data Fig. 4e
- Supplementary Table 9: Species/metabolite correlation statistics associated with Fig. 3f and Extended Data Fig. 4f
- Supplementary Table 10: Dunnett’s test results for pairwise comparisons of *L. iners* / *L. crispatus* ratios in Cys-supplemented MRSQ broth treated with SMC (associated with Fig. 5a)
- Supplementary Table 11: Selected Tukey Test Results for pairwise comparison of ratios between L. crispatus and the sum of other taxa in S-Broth, comparing MTZ + SMC to other conditions (associated with Fig. 5d)

## Supplementary Files

- Supplementary Table 1: Compressed file containing directory with R analysis code and associated data files and directory structure for genome gene-content analysis.
- Supplementary Table 2: Compressed file containing directory with R analysis code and associated data files and directory structure for 16S-gene based microbiota-metabolite analysis.

