## Supplementary File 1 for "Cysteine dependence in *Lactobacillus iners* constitutes a novel therapeutic target to modify the vaginal microbiota": L_iners_Lacto_Cys_gene_analysis_for_paper_2021_06_06.Rmd

````
---
title: "FGT Lactobacillus iners gene complement analysis for paper"
author: "Seth M. Bloom"
date: "06/06/2021"
output:
  pdf_document: default
  html_document: default
  word_document: default
---
  
  
**Summary:**  
Review of FGT _Lactobacillus_ pan-genome for genes involved in biosynthesis and transport of cysteine and related compounds. Rough draft of analysis for _L. iners_ cysteine paper.  
  
  
**NOTE:** This is an update from version 6 of this code because it outputs the translated pan-genome as a multi-fasta for BLAST "sanity check" analysis.
  
    
## Figures/tables:  
  
*  Supplementary data (input):
    +  Gene presence absence table (tsv vs excel)
    +  Pan-genome multi-fasta file (fasta)
    +  Genome taxonomy table (tsv vs excel)
    +  EggNOG annotation table (tsv vs excel)
    +  Gene wishlist (tsv vs excel)  
    
*  Figures (output):
    +  Violin plot of per-genome gene counts (supplement)
    +  Heatmap (geom_rast) of per-genome gene presence absence for Cys biosynthesis pathways by species (main figure panel)
    +  Heatmap (geom_rast) of per-genome gene presence absence for cyst(e)ine and glutathione transport and GSH synthesis pathways by species (main figure panel)
    
*  Table (output):
    +  Species pan-genome size and percent annotated (supplementary)
    +  Statistics on per-species prevalence of genes of interest (supplementary)
    

Additionally, supplemental blast-based is performed, although no figures to formally include in manuscript.  
  
  
## Load packages  
  
  
```{r load-packages, echo=FALSE}
library(seqinr)
packageVersion("seqinr")
library(tidyverse)
packageVersion("tidyverse")
library(ggExtra)
packageVersion("ggExtra")
library(knitr)
packageVersion("knitr")
```
  
**Working Directory:**  
`r getwd()`  
  
### Load gene table and SGB assignments and taxonomy  
  

```{r load-pangenome-data, echo=FALSE, message=FALSE}
f_gene_pres_abs_df <- "Lacto_Genome_Data_2020_07_21/gene_presence_absence_2020_07_21.Rtab"
f_gene_pres_abs_df

f_pangenome_fasta_DNA_list <- "Lacto_Genome_Data_2020_07_21/pan_genome_reference_2020_07_21.fa"
f_pangenome_fasta_DNA_list

#Read in gene presence absence file (presence/absence of each gene in each genome)
gene_pres_abs_df <- read_tsv(f_gene_pres_abs_df)

#Read in DNA fasta files of all query genes (i.e. entire pan-genome) submitted to EggNOG
pangenome_fasta_DNA_list_sep <- read.fasta(f_pangenome_fasta_DNA_list, 
                                           seqtype = "DNA")

#Read in dataframe of headers from the above fasta file (includes gene names)
pangenome_fasta_headers <- readr::read_tsv(f_pangenome_fasta_DNA_list, col_names = c("Fasta_Header")) %>%
  dplyr::filter(str_detect(Fasta_Header, "^>"))

#Define path to write output
path_output <- "Output"
```
  

### Load Lactobacillus genome SGB taxonomy table
  
  
```{r Load-Lactobacillus-genome-taxonomy-table, echo=FALSE}
#Define file name and path
f_genome_SGB_taxonomy.FGT.Lacto <- "Lacto_Genome_Data_2020_07_21/FGT_Lacto_genome_SGB_taxonomy_2020_09_08.txt"
f_genome_SGB_taxonomy.FGT.Lacto

#Read in file
genome_SGB_taxonomy.FGT.Lacto <- readr::read_tsv(f_genome_SGB_taxonomy.FGT.Lacto)
```

  
### Reformat gene tables and merge to gene annotation data

```{r pivot-gene-table-to-long-form-and-add-gene-groups, echo=FALSE}
#Pivot gene_pres_abs_df to long form
gene_pres_abs_long.all <- gene_pres_abs_df %>% 
  pivot_longer(cols = -Gene,
               names_to = "Genome", 
               values_to = "Gene_Pres")

#Divide components of the FASTA header into the Gene_Group (before space) and
#Roary_Name (after space)
pangenome_gene_names <- pangenome_fasta_headers %>% 
  dplyr::mutate(No_Carrot = str_remove(Fasta_Header, "^>")) %>% 
  tidyr::separate(col = No_Carrot, 
                  into = c("Gene_Group", "Gene"), 
                  sep = " ")
```
  
  
Quantify the number of genes in pangenome_gene_names lacking matches in gene_pres_abs_long.all and vice versa.  
  

```{r confirm-no-missing-or-duplicate-genes, echo=FALSE}
#Quantify the number of genes in pangenome_gene_names lacking matches 
#in gene_pres_abs_long.all
pangenome_gene_names %>% 
  dplyr::anti_join(gene_pres_abs_long.all,
                   by = "Gene") %>% 
  nrow()

#Quantify the number of genes in the gene_pres_abs_long.all  with and without 
#matches in the pangenome_gene_names
gene_pres_abs_long.all %>% 
  dplyr::anti_join(pangenome_gene_names,
                   by = "Gene") %>% 
  nrow()

#Add info from pangenome_gene_names to gene_pres_abs_long.all
gene_pres_abs_long.all <- gene_pres_abs_long.all %>% 
  dplyr::left_join(pangenome_gene_names, by = "Gene")
```
  
### Reconcile genome names between SGBs and gene-presence-absence table  
  
Genome names were updated for pan-genome construction after initial assignment of genome/SGB taxonomy. This script reconciles the updated genome names in the pan-genome with the taxonomic assignments.  
  

```{r format-pan-genome-gene-names-to-match-SGB-genome-names, echo=FALSE}
#Compared to the genome names in genome_SGB_taxonomy.FGT.Lacto, some of the genomes
#in the gene_pres_abs table have had suffices including "-renamedScaffs",
#"-merged", and "-FINAL". Count the number.
gene_pres_abs_long.all$Genome %>% 
  unique() %>% 
  setdiff(genome_SGB_taxonomy.FGT.Lacto$Genome) %>% 
  length()

#Check whether removing the above suffices (in that order) removes all 
#mismatches with the genome names in genome_SGB_taxonomy.FGT.Lacto
gene_pres_abs_long.all$Genome %>% 
  unique() %>% 
  stringr::str_remove("-renamedScaffs$") %>% 
  stringr::str_remove("-merged$") %>% 
  stringr::str_remove("-FINAL$") %>% 
  setdiff(genome_SGB_taxonomy.FGT.Lacto$Genome) %>% 
  length()

#Create df genomes.renamed for genomes in gene_pres_abs_long.all
genomes.renamed <- gene_pres_abs_long.all %>% 
  dplyr::distinct(Genome) %>% 
  dplyr::mutate(Genome_Renamed = str_remove(Genome, "-renamedScaffs$") %>% 
                  str_remove("-merged$") %>% 
                  str_remove("-FINAL$"))

#Add column Genome_Rename to gene_pres_abs_long.all
gene_pres_abs_long.all_renamed <- gene_pres_abs_long.all %>% 
  dplyr::left_join(genomes.renamed,
                   by = "Genome")

#Check whether removing the suffixes from Genome causes there to be any duplicated
#Genome_Renamed names
gene_pres_abs_long.all_renamed %>% 
  dplyr::distinct(Genome, Genome_Renamed) %>% 
  dplyr::count(Genome_Renamed) %>% 
  dplyr::filter(n > 1) %>% 
  nrow()

#Are there any genomes in gene_pres_abs_long.all_renamed absent from 
#genome_SGB_taxonomy.FGT.Lacto?
gene_pres_abs_long.all_renamed$Genome_Renamed %>% 
  unique() %>% 
  setdiff(genome_SGB_taxonomy.FGT.Lacto$Genome) %>% 
  length()
```
  
  
### Summarize gene counts per genome with singletons  
  
Filter the pan-genome to remove any genomes with gene counts of 0.  
  
```{r remove-genomes-with-0-gene-count-from-gene-pres-absence-table, echo=FALSE}
#Filter out genomes with gene counts = 0
gene_pres_abs_long <- gene_pres_abs_long.all_renamed %>% 
  dplyr::group_by(Genome, Genome_Renamed) %>% 
  dplyr::filter(sum(Gene_Pres) > 0) %>% 
  dplyr::ungroup()
```
  
  
Now re-review genome gene count distribution after filtering 0-count genomes.  
  
```{r summarise-per-genome-gene-counts-with-singletons-after-filtering-0-count-genomes, echo=FALSE}
#Summarise per-genome gene counts
genome_gene_cts <- gene_pres_abs_long %>% 
  dplyr::group_by(Genome, Genome_Renamed) %>% 
  dplyr::summarise(Genome_Gene_Count = sum(Gene_Pres)) %>% 
  dplyr::ungroup()


genome_gene_cts$Genome_Gene_Count %>% 
  quantile(probs = seq(0, 1, 0.1)) %>% 
  tibble::enframe(name = "Percentile", value = "Per-Genome Gene Count") %>% 
  knitr::kable(caption = "Percentile Distribution of Per-Genome Gene Counts in Filtered Genomes (with Singletons)",
               digits = 0)

p.genome.gene.cts.with.singletons <- genome_gene_cts %>% 
  ggplot(aes(x = "Genome", y = Genome_Gene_Count)) +
  geom_boxplot(alpha = 0, outlier.size = -1, 
               width = 0.45, color = "red",
               coef = 5) +
  geom_violin(alpha = 0, color = "blue", width = 0.5) +
  geom_jitter(width = 0.1, alpha = 0.7, size = 0.5) +
  scale_y_continuous(limits = c(0, NA)) +
  labs(title = "Per-Genome Gene Count in Filtered Genomes") +
  theme(plot.title = element_text(hjust = 0.5, size = 14, face = "bold"),
        axis.title.x = element_blank(),
        axis.title.y = element_text(size = 12, face = "bold"),
        axis.text = element_text(size = 12))

p.genome.gene.cts.with.singletons
```
  
  
### Review the number of genomes per species that are reference vs novel  
  
```{r merge-SGB-and-species-information-to-genomes, echo=FALSE}
#Left_join genome_SGB_taxonomy.FGT.Lacto to genome_gene_cts by genome
genome_gene_cts_SGB_species <- genome_gene_cts %>% 
  dplyr::left_join(genome_SGB_taxonomy.FGT.Lacto,
                   by = c("Genome_Renamed" = "Genome"))
```
  

```{r identify-and-count-reference-vs-new-genomes, echo=FALSE}
#Label reference genomes
genome_reference <- genome_gene_cts_SGB_species %>% 
  dplyr::distinct(Species, Species_SGB, SGB, Genome_Renamed, Genome_Gene_Count) %>% 
  dplyr::arrange(desc(Genome_Renamed)) %>% 
  dplyr::mutate(Novelty = case_when(str_detect(Genome_Renamed, "^SAM|Lactobacillus") ~ "Reference Genome",
                                    TRUE ~ "Novel Genome")) 

#Species genomes added
genome_reference %>% 
  dplyr::group_by(Species, Novelty) %>% 
  dplyr::count(name = "Num_Genomes") %>% 
  dplyr::ungroup() %>% 
  tidyr::pivot_wider(names_from = Novelty,
                     values_from = Num_Genomes) %>% 
  dplyr::mutate(`Total Genomes` = `Novel Genome` + `Reference Genome`,
                `Fold Increase in Genome Number` = `Total Genomes` / `Reference Genome`,
                `% Increase in Genome Number` = (`Fold Increase in Genome Number` - 1) * 100,
                Species = str_replace(Species, "actobacillus_", ". "),
                Species = factor(Species,
                                 levels = c("L. crispatus",
                                            "L. iners",
                                            "L. jensenii",
                                            "L. gasseri",
                                            "L. vaginalis"))) %>% 
  dplyr::arrange(Species) %>% 
  knitr::kable(caption = "Novel genomes added by Species")

#Stacked bar plot of reference vs novel genomes by species
p.genomes.added.species.bar.stack <- genome_reference %>% 
  dplyr::group_by(Species, Novelty) %>% 
  dplyr::count(name = "Num_Genomes") %>% 
  dplyr::ungroup() %>% 
  dplyr::mutate(Species = str_replace(Species, "actobacillus_", ". "),
                Novelty = str_remove(Novelty, " Genome"),
                Species = factor(Species,
                                 levels = c("L. crispatus",
                                            "L. iners",
                                            "L. jensenii",
                                            "L. gasseri",
                                            "L. vaginalis"))) %>% 
  ggplot(aes(x = Species, y = Num_Genomes, fill = Novelty)) +
  geom_col() +
  geom_text(aes(label = Num_Genomes)) +
  labs(y = "# Genomes") +
  theme(axis.text.x = element_text(angle = 45, hjust = 1, size = 12, face = "italic"),
        axis.text.y = element_text(size = 12),
        axis.title = element_text(size = 14, face = "bold"),
        legend.title = element_blank(),
        legend.text = element_text(size = 12))

p.genomes.added.species.bar.stack
```
  
  
## Review Genome gene counts by Species with singletons (Not including in paper because excluding singletons for final analysis) 
  

```{r Review-genomes-by-species-with-singletons, echo=FALSE}
p.gene.ct.by.species.with.singletons <- genome_gene_cts_SGB_species %>%
  dplyr::add_count(Species, name = "Num_Genomes_Per_Species") %>%
  dplyr::rename(`Gene Count per Genome` = Genome_Gene_Count) %>%
  dplyr::mutate(Species = str_replace(Species,
                                      "^Lactobacillus_",
                                      "L. "),
                Species = paste0(Species,
                                 "\n(n = ",
                                 Num_Genomes_Per_Species,
                                 ")")) %>%
  ggplot(aes(x = Species, y = `Gene Count per Genome`)) +
  geom_boxplot(alpha = 0,
               outlier.size = -1,
               width = 0.45,
               color = "red",
               coef = 10) +
  geom_violin(alpha = 0, color = "blue", width = 1) +
  geom_jitter(width = 0.1, alpha = 0.5, size = 0.5) +
  scale_y_continuous(limits = c(0, NA)) +
  # labs(title = "Per-Genome Gene Count by Species") +
  theme(plot.title = element_text(hjust = 0.5, size = 14, face = "bold"),
        axis.title = element_text(size = 12, face = "bold"),
        axis.text.x = element_text(size = 12,
                                   # angle = 45,
                                   # hjust = 1,
                                   face = "italic"),
        axis.text.y = element_text(size = 12))

p.gene.ct.by.species.with.singletons

#Plot with genome novelty information
p.gene.ct.by.species.novelty.with.singletons <- genome_gene_cts_SGB_species %>% 
  dplyr::left_join(genome_reference,
                   by = c("Genome_Renamed",
                          "SGB",
                          "Species",
                          "Species_SGB",
                          "Genome_Gene_Count")) %>% 
  dplyr::mutate(Novelty = str_remove(Novelty, " Genome"),
                Novelty = factor(Novelty,
                                 levels = c("Reference", "Novel")),
                Species = str_replace(Species, 
                                      "^Lactobacillus_",
                                      "L. ")) %>% 
  dplyr::group_by(Species, Novelty) %>% 
  dplyr::add_count(name = "Num_Genomes_Per_Species_Novelty") %>% 
  dplyr::ungroup() %>% 
  dplyr::add_count(Species, name = "Num_Genomes_Per_Species") %>% 
  dplyr::rename(`Gene Count per Genome` = Genome_Gene_Count) %>% 
  dplyr::mutate(Species = str_replace(Species, 
                                      "^Lactobacillus_",
                                      "L. "),
                Species = paste0(Species, 
                                 "\n(n = ",
                                 Num_Genomes_Per_Species,
                                 ")"),
                Species = factor(Species,
                                 levels = c("L. crispatus\n(n = 508)",
                                            "L. iners\n(n = 327)",
                                            "L. jensenii\n(n = 137)",
                                            "L. gasseri\n(n = 216)",
                                            "L. vaginalis\n(n = 30)"))) %>% 
  ggplot(aes(x = Species, 
             y = `Gene Count per Genome`, 
             color = Novelty,
             fill = Novelty)) +
  geom_boxplot(outlier.size = -1, 
               color = "black",
               width = 0.8, 
               coef = 20) +
  geom_point(position = position_jitterdodge(jitter.width = 0.4, 
                                             dodge.width = 0.8), 
             size = 0.7,
             alpha = 0.9) +
  geom_label(aes(label = Num_Genomes_Per_Species_Novelty,
                y = 2700),
            position = position_dodge(width = 0.8),
            color = "black",
            show.legend = FALSE,
            size = 2.9) +
  scale_y_continuous(limits = c(0, NA)) +
  scale_fill_manual(values = c("tomato1", "dodgerblue1")) +
  scale_color_manual(values = c("red", "blue")) +
  labs(
    # title = "Per-Genome Gene Count by Species",
       fill = "Genome\nSource",
       color = "Genome\nSource") +
  theme(plot.title = element_text(hjust = 0.5, size = 14, face = "bold"),
        legend.title = element_text(size = 12, face = "bold"),
        legend.text = element_text(size = 10),
        axis.title = element_text(size = 12, face = "bold"),
        axis.text.x = element_text(size = 9, 
                                   face = "italic"),
        axis.text.y = element_text(size = 9))

p.gene.ct.by.species.novelty.with.singletons
```
  

  
### Save per-species genome gene count plots (with singletons)

```{r save-genome-gene-count-plots-with-singletons, echo=FALSE}
###File name to save
f_p.gene.ct.by.species.with.singletons <- "Gene_count_per_genome_fig_supp_w_singletons.pdf"
f_p.gene.ct.by.species.with.singletons

### Write to file
cowplot::ggsave2(filename = f_p.gene.ct.by.species.with.singletons,
                 plot = p.gene.ct.by.species.with.singletons,
                 path = path_output,
                 width = 6,
                 height = 4)

###File name to save
f_p.gene.ct.by.species.novelty.with.singletons <- "Gene_count_per_genome_novelty_fig_supp_w_singletons.pdf"
f_p.gene.ct.by.species.novelty.with.singletons

### Write to file
cowplot::ggsave2(filename = f_p.gene.ct.by.species.novelty.with.singletons,
                 plot = p.gene.ct.by.species.novelty.with.singletons,
                 path = path_output,
                 width = 6,
                 height = 4)
```
  
  
### Review Genome gene counts by Species with NO singletons (supplementary figure)  
  


```{r merge-taxonomy-and-identify-singleton-genes-per-species, echo=FALSE}
#Integrate genome taxonomy information into gene presence/absence table.
gene_pres_abs_Spec_SGB <- gene_pres_abs_long %>% 
  dplyr::left_join(genome_gene_cts_SGB_species,
                   by = c("Genome",
                          "Genome_Renamed"))

#Add column with number of times each gene is present in Species
gene_pres_Spec_SGB_times_seen <- gene_pres_abs_Spec_SGB %>% 
  dplyr::filter(Gene_Pres == 1) %>%
  dplyr::group_by(Species, Gene) %>% 
  dplyr::mutate(Num_Genomes_Present = case_when(n() == 1 ~ "Singleton Gene",
                                                TRUE ~ "Gene in Multiple Genomes")) %>% 
  dplyr::ungroup() 

#Add back information on Num_Genomes_Present to gene_pres_abs_Spec_SGB
gene_pres_abs_Spec_SGB <- gene_pres_abs_Spec_SGB %>% 
  dplyr::left_join(gene_pres_Spec_SGB_times_seen %>% 
                     dplyr::distinct(Gene, Species, Num_Genomes_Present))
```

```{r Review-genomes-by-species-without-singletons, echo=FALSE}
#DF of gene counts per genome
genome_gene_cts_SGB_species.no.singletons <- gene_pres_abs_Spec_SGB %>% 
  dplyr::filter(Num_Genomes_Present == "Gene in Multiple Genomes") %>% 
  dplyr::group_by(Genome, Genome_Renamed, Species, Species_SGB, SGB) %>% 
  dplyr::summarise(Genome_Gene_Count = sum(Gene_Pres)) %>% 
  dplyr::ungroup()
```


```{r Plot-genomes-by-species-without-singletons, echo=FALSE}
p.gene.ct.by.species.no.singletons <- genome_gene_cts_SGB_species.no.singletons %>%
  dplyr::add_count(Species, name = "Num_Genomes_Per_Species") %>%
  dplyr::rename(`Gene Count per Genome` = Genome_Gene_Count) %>%
  dplyr::mutate(Species = str_replace(Species,
                                      "^Lactobacillus_",
                                      "L. "),
                Species = paste0(Species,
                                 "\n(n = ",
                                 Num_Genomes_Per_Species,
                                 ")")) %>%
  ggplot(aes(x = Species, y = `Gene Count per Genome`)) +
  geom_boxplot(alpha = 0,
               outlier.size = -1,
               width = 0.45,
               color = "red",
               coef = 10) +
  geom_violin(alpha = 0, color = "blue", width = 1) +
  geom_jitter(width = 0.1, alpha = 0.5, size = 0.5) +
  scale_y_continuous(limits = c(0, NA)) +
  # labs(title = "Per-Genome Gene Count by Species") +
  theme(plot.title = element_text(hjust = 0.5, size = 14, face = "bold"),
        axis.title = element_text(size = 12, face = "bold"),
        axis.text.x = element_text(size = 12,
                                   # angle = 45,
                                   # hjust = 1,
                                   face = "italic"),
        axis.text.y = element_text(size = 12))

p.gene.ct.by.species.no.singletons

#Plot with genome novelty information
p.gene.ct.by.species.novelty.no.singletons <- genome_gene_cts_SGB_species.no.singletons %>% 
  dplyr::left_join(genome_reference %>% 
                     dplyr::select(Genome_Renamed,
                                   Species,
                                   Novelty),
                   by = c("Genome_Renamed",
                          "Species")) %>% 
  dplyr::mutate(Novelty = str_remove(Novelty, " Genome"),
                Novelty = factor(Novelty,
                                 levels = c("Reference", "Novel")),
                Species = str_replace(Species, 
                                      "^Lactobacillus_",
                                      "L. ")) %>% 
  dplyr::group_by(Species, Novelty) %>% 
  dplyr::add_count(name = "Num_Genomes_Per_Species_Novelty") %>% 
  dplyr::ungroup() %>% 
  dplyr::add_count(Species, name = "Num_Genomes_Per_Species") %>% 
  dplyr::rename(`Gene Count per Genome` = Genome_Gene_Count) %>% 
  dplyr::mutate(Species = str_replace(Species, 
                                      "^Lactobacillus_",
                                      "L. "),
                Species = paste0(Species, 
                                 "\n(n = ",
                                 Num_Genomes_Per_Species,
                                 ")"),
                Species = factor(Species,
                                 levels = c("L. crispatus\n(n = 508)",
                                            "L. iners\n(n = 327)",
                                            "L. jensenii\n(n = 137)",
                                            "L. gasseri\n(n = 216)",
                                            "L. vaginalis\n(n = 30)"))) %>% 
  ggplot(aes(x = Species, 
             y = `Gene Count per Genome`, 
             color = Novelty,
             fill = Novelty)) +
  geom_boxplot(outlier.size = -1, 
               color = "black",
               width = 0.8, 
               coef = 20) +
  geom_point(position = position_jitterdodge(jitter.width = 0.4, 
                                             dodge.width = 0.8), 
             size = 0.7,
             alpha = 0.9) +
  geom_label(aes(label = Num_Genomes_Per_Species_Novelty,
                y = 2700),
            position = position_dodge(width = 0.8),
            color = "black",
            show.legend = FALSE,
            size = 2.9) +
  scale_y_continuous(limits = c(0, NA)) +
  scale_fill_manual(values = c("tomato1", "dodgerblue1")) +
  scale_color_manual(values = c("red", "blue")) +
  labs(
    # title = "Per-Genome Gene Count by Species",
       fill = "Genome\nSource",
       color = "Genome\nSource") +
  theme(plot.title = element_text(hjust = 0.5, size = 14, face = "bold"),
        legend.title = element_text(size = 12, face = "bold"),
        legend.text = element_text(size = 10),
        axis.title = element_text(size = 12, face = "bold"),
        axis.text.x = element_text(size = 9, 
                                   face = "italic"),
        axis.text.y = element_text(size = 9))

p.gene.ct.by.species.novelty.no.singletons
```
  

  
### Save per-species genome gene count plots (without singletons)

```{r save-genome-gene-count-plots-without-singletons-PDF-and-RDS, echo=FALSE}
##Save PDF##

###File name to save
f.pdf_p.gene.ct.by.species.no.singletons <- "Gene_count_per_genome_fig_supp_no_singletons.pdf"
f.pdf_p.gene.ct.by.species.no.singletons

### Write to file
cowplot::ggsave2(filename = f.pdf_p.gene.ct.by.species.no.singletons,
                 plot = p.gene.ct.by.species.no.singletons,
                 path = path_output,
                 width = 6,
                 height = 4)

###File name to save
f.pdf_p.gene.ct.by.species.novelty.no.singletons <- "Gene_count_per_genome_novelty_fig_supp_no_singletons.pdf"
f.pdf_p.gene.ct.by.species.novelty.no.singletons

### Write to file
cowplot::ggsave2(filename = f.pdf_p.gene.ct.by.species.novelty.no.singletons,
                 plot = p.gene.ct.by.species.novelty.no.singletons,
                 path = path_output,
                 width = 7,
                 height = 4)

##Save RDS##

###File name to save
f.RDS_p.gene.ct.by.species.no.singletons <- "Gene_count_per_genome_fig_supp_no_singletons.RDS"
f.RDS_p.gene.ct.by.species.no.singletons

### Write to file
readr::write_rds(p.gene.ct.by.species.no.singletons,
                 file.path(path_output,
                           f.RDS_p.gene.ct.by.species.no.singletons))

###File name to save
f.RDS_p.gene.ct.by.species.novelty.no.singletons <- "Gene_count_per_genome_novelty_fig_supp_no_singletons.RDS"
f.RDS_p.gene.ct.by.species.novelty.no.singletons

### Write to file
readr::write_rds(p.gene.ct.by.species.novelty.no.singletons,
                 file.path(path_output,
                           f.RDS_p.gene.ct.by.species.novelty.no.singletons))
```


### Review # of genes shared between pan-genomes of different Species?  
  

```{r assess-per-species-gene-counts-and-shared-genes, echo=FALSE}
#What is the pan-genome size of each Species and how many are singletons?
gene_pres_Spec_SGB_times_seen %>% 
  dplyr::distinct(Species, Gene, Num_Genomes_Present) %>% 
  dplyr::count(Species, Num_Genomes_Present, name = "Pan_genome_gene_count") %>% 
  tidyr::pivot_wider(names_from = Num_Genomes_Present,
                     values_from = Pan_genome_gene_count) %>% 
  dplyr::mutate(`Total Genes` = `Singleton Gene` + `Gene in Multiple Genomes`) %>% 
  knitr::kable(caption = "Pan-genome size by Species")

#How many genes are shared between pan-genomes of different Species, including singletons?
p.species.per.gene.hist.w.singletons <- gene_pres_abs_Spec_SGB %>% 
  dplyr::filter(Gene_Pres == 1) %>% 
  dplyr::distinct(Species, Gene) %>% 
  dplyr::count(Gene, name = "Number_Lacto_Species") %>% 
  ggplot(aes(x = Number_Lacto_Species)) + 
  stat_bin(binwidth = 1) + 
  stat_bin(binwidth = 1, geom = "label", aes(label = ..count..)) +
  scale_x_continuous(breaks = c(1:7)) +
  labs(title = "Number of Distinct Lactobacillus Species\nPan-genomes Containing Each Gene (with singletons)",
       y = "Number of Genes",
       x = "Number of Lactobacillus Species") +
  theme(plot.title = element_text(hjust = 0.5, size = 14, face = "bold"),
        axis.title = element_text(size = 12, face = "bold"),
        axis.text = element_text(size = 12))

p.species.per.gene.hist.w.singletons

#How many genes are shared between pan-genomes of different Species without singletons?
p.species.per.gene.hist.no.singletons <- gene_pres_abs_Spec_SGB %>% 
  dplyr::filter(Gene_Pres == 1,
                Num_Genomes_Present == "Gene in Multiple Genomes") %>% 
  dplyr::distinct(Species, Gene) %>% 
  dplyr::count(Gene, name = "Number_Lacto_Species") %>% 
  ggplot(aes(x = Number_Lacto_Species)) + 
  stat_bin(binwidth = 1) + 
  stat_bin(binwidth = 1, geom = "label", aes(label = ..count..)) +
  scale_x_continuous(breaks = c(1:7)) +
  labs(title = "Number of Distinct Lactobacillus Species\nPan-genomes Containing Each Gene (no singletons)",
       y = "Number of Genes",
       x = "Number of Lactobacillus Species") +
  theme(plot.title = element_text(hjust = 0.5, size = 14, face = "bold"),
        axis.title = element_text(size = 12, face = "bold"),
        axis.text = element_text(size = 12))

p.species.per.gene.hist.no.singletons
```
  
```{r Calculate-species-pan-genome, echo=FALSE}
#Species pan-genome expansion with singletons
gene_pres_abs_Spec_SGB %>% 
  dplyr::filter(Gene_Pres == 1) %>% 
  dplyr::left_join(genome_reference %>% 
                     dplyr::select(Genome_Renamed, Novelty),
                   by = c("Genome_Renamed")) %>% 
  dplyr::distinct(Gene, Species, Novelty) %>% 
  dplyr::mutate(Novelty = str_remove(Novelty, " Genome"),
                Presence = 1) %>% 
  tidyr::pivot_wider(names_from = Novelty,
                     values_from = Presence) %>% 
  tidyr::replace_na(list(Novel = 0, Reference = 0)) %>% 
  dplyr::mutate(Novelty = case_when(Reference == 1 ~ "Reference Genes",
                                    Novel == 1 ~ "Novel Genes")) %>% 
  dplyr::count(Species, Novelty, name = "Num_Genes") %>% 
  tidyr::pivot_wider(names_from = Novelty,
                     values_from = Num_Genes) %>% 
  dplyr::mutate(Species = str_replace(Species, "actobacillus_", ". "),
                Species = factor(Species,
                                 levels = c("L. crispatus",
                                            "L. iners",
                                            "L. jensenii",
                                            "L. gasseri",
                                            "L. vaginalis")),
                `Total Genes` = `Novel Genes` + `Reference Genes`,
                `Fold Increase in Gene Number` = `Total Genes` / `Reference Genes`,
                `% Increase in Genes` = (`Fold Increase in Gene Number` - 1) * 100) %>% 
  dplyr::arrange(Species) %>% 
  knitr::kable(caption = "Pan-genome expansion by species (with singletons)")

#Species pan-genome expansion without singletons
pan.genome.expansion.no.singletons <- gene_pres_abs_Spec_SGB %>% 
  dplyr::filter(Gene_Pres == 1,
                Num_Genomes_Present == "Gene in Multiple Genomes") %>% 
  dplyr::left_join(genome_reference %>% 
                     dplyr::select(Genome_Renamed, Novelty),
                   by = c("Genome_Renamed")) %>% 
  dplyr::distinct(Gene, Species, Novelty) %>% 
  dplyr::mutate(Novelty = str_remove(Novelty, " Genome"),
                Presence = 1) %>% 
  tidyr::pivot_wider(names_from = Novelty,
                     values_from = Presence) %>% 
  tidyr::replace_na(list(Novel = 0, Reference = 0)) %>% 
  dplyr::mutate(Novelty = case_when(Reference == 1 ~ "Reference Genes",
                                    Novel == 1 ~ "Novel Genes")) %>% 
  dplyr::count(Species, Novelty, name = "Num_Genes") %>% 
  tidyr::pivot_wider(names_from = Novelty,
                     values_from = Num_Genes) %>% 
  dplyr::mutate(Species = str_replace(Species, "actobacillus_", ". "),
                Species = factor(Species,
                                 levels = c("L. crispatus",
                                            "L. iners",
                                            "L. jensenii",
                                            "L. gasseri",
                                            "L. vaginalis")),
                `Total Genes` = `Novel Genes` + `Reference Genes`,
                `Fold Increase in Gene Number` = `Total Genes` / `Reference Genes`,
                `% Increase in Genes` = (`Fold Increase in Gene Number` - 1) * 100) %>% 
  dplyr::arrange(Species) 

pan.genome.expansion.no.singletons %>% 
  knitr::kable(caption = "Pan-genome expansion by species (with no singletons)")

```

  
### Import EggNOG gene annotations.  
  

```{r import-EggNOG-annotations, echo=FALSE}
f_EggNOG_annotations_raw <- "eggNOG_annotations/job_MM_atwxphdb_annotations.txt"

#Read in EggNOG annotation table, skipping first 4 and final 3 lines because
#they contain non-data summary information
EggNOG_annotations_raw <- read_tsv(f_EggNOG_annotations_raw, skip = 4) %>% 
  dplyr::filter(row_number() < n() - 2) 
```
  
### Supplementary table: EggNOG annotation summary  
  
Assess gene annotation statistics and tabulate.  
  

```{r review-gene-distribution-in-EggNOG-output-and-percent-unannotated, echo=FALSE}
writeLines("\nHow many genes in the pan-genome are absent from the EggNOG output?")
pangenome_gene_names %>% 
  dplyr::anti_join(EggNOG_annotations_raw,
                   by = c("Gene_Group" = "Query")) %>% 
  nrow()

writeLines("\nAre any of the Query genes duplicated in the EggNOG output table?")
EggNOG_annotations_raw %>% 
  dplyr::pull(Query) %>% 
  duplicated() %>% 
  any()

#What is the pan-genome size of each Species and fraction annotated with singletons?
pan.genome.spec.perc.annot.with.singletons <- gene_pres_abs_Spec_SGB %>% 
  dplyr::filter(Gene_Pres == 1) %>% 
  dplyr::distinct(Species, Gene, Gene_Group) %>% 
  dplyr::mutate(Annotated = case_when(Gene_Group %in% EggNOG_annotations_raw$Query ~ "Annotated Genes",
                                      TRUE ~ "Unannotated Genes")) %>% 
  dplyr::count(Species, Annotated, name = "Pan_genome_gene_count") %>% 
  tidyr::pivot_wider(names_from = Annotated,
                     values_from = Pan_genome_gene_count) %>% 
  dplyr::mutate(`Total Genes` = `Annotated Genes` + `Unannotated Genes`,
                `Percent Annotated Genes` = `Annotated Genes` / `Total Genes`,
                `Percent Unannotated Genes` = `Unannotated Genes` / `Total Genes`,
                across(starts_with("Percent"),
                       ~ round(.x * 100, digits = 2) %>% 
                         str_pad(width = 5, side = "right", pad = "0") %>% 
                         paste0("%")),
                Species = str_replace(Species, "actobacillus_", ". "),
                Species = factor(Species,
                                 levels = c("L. crispatus",
                                            "L. iners",
                                            "L. jensenii",
                                            "L. gasseri",
                                            "L. vaginalis"))) %>% 
  dplyr::arrange(Species) 

pan.genome.spec.perc.annot.with.singletons %>% 
  knitr::kable(caption = "Pan-genome size and percent annotated by Species with singletons")

#What is the pan-genome size of each Species and fraction annotated without singletons?
pan.genome.spec.perc.annot.no.singletons <- gene_pres_abs_Spec_SGB %>% 
  dplyr::filter(Gene_Pres == 1,
                Num_Genomes_Present == "Gene in Multiple Genomes") %>% 
  dplyr::distinct(Species, Gene, Gene_Group) %>% 
  dplyr::mutate(Annotated = case_when(Gene_Group %in% EggNOG_annotations_raw$Query ~ "Annotated Genes",
                                      TRUE ~ "Unannotated Genes")) %>% 
  dplyr::count(Species, Annotated, name = "Pan_genome_gene_count") %>% 
  tidyr::pivot_wider(names_from = Annotated,
                     values_from = Pan_genome_gene_count) %>% 
  dplyr::mutate(`Total Genes` = `Annotated Genes` + `Unannotated Genes`,
                `Percent Annotated Genes` = `Annotated Genes` / `Total Genes`,
                `Percent Unannotated Genes` = `Unannotated Genes` / `Total Genes`,
                across(starts_with("Percent"),
                       ~ round(.x * 100, digits = 2) %>% 
                         str_pad(width = 5, side = "right", pad = "0") %>% 
                         paste0("%")),
                Species = str_replace(Species, "actobacillus_", ". "),
                Species = factor(Species,
                                 levels = c("L. crispatus",
                                            "L. iners",
                                            "L. jensenii",
                                            "L. gasseri",
                                            "L. vaginalis"))) %>% 
  dplyr::arrange(Species)

pan.genome.spec.perc.annot.no.singletons %>% 
  knitr::kable(caption = "Pan-genome size and percent annotated by Species with no singletons")
```


  
### Table of merged pangenome expansion and percent annotation (no singletons)  
  
  

```{r merge-pangenome-expansion-annotation-data-no-singletons, echo=FALSE}
pan.genome.expansion.annot.no.singletons <- pan.genome.expansion.no.singletons %>% 
  dplyr::select(Species, `Reference Genes`, `Novel Genes`, `Total Genes`, 
                `Fold Pan-Genome Expansion` = `Fold Increase in Gene Number`) %>% 
  dplyr::left_join(pan.genome.spec.perc.annot.no.singletons,
                   by = c("Species",
                          "Total Genes"))

pan.genome.expansion.annot.no.singletons %>% 
  knitr::kable(caption = "Species pan-genome size, degree of expansion, and percent annotated (no singletons)")
```
    
    
Write table to file.  
  

```{r save-table-of-pangenome-expansion-and-annotation-no-singletons, echo=FALSE}
f_pan.genome.expansion.annot.no.singletons <- "Lacto_species_pangenome_expansion_annotation_no_singletons.txt"
f_pan.genome.expansion.annot.no.singletons

readr::write_tsv(pan.genome.expansion.annot.no.singletons,
                 file.path(path_output, f_pan.genome.expansion.annot.no.singletons))
```

### Review and collapse seed orthologs in EggNOG output  
  
There can be multiple different gene sequences that map to the same eggNOG seed ortholog. For simplicity of identifying genes of interest within the overall eggNOG output, this step first collapses to unique seed orthologs. Subsequent steps then parse these unique seed orthologs for functional annotations of interest. Later steps then identify all genes associated with each seed ortholog of interest (sometimes just one gene, sometimes multiple) and their taxonomic associations.  
  

```{r review-seed-orthologs-in-EggNOG-output, echo=FALSE}
writeLines("\nHow many unique Seed orthologs are in the EggNOG output (with singletons)?")
EggNOG_annotations_raw %>% 
  dplyr::distinct(`Seed ortholog`) %>% 
  nrow()

#Plot mapping of Query genes to Seed orthologs
p.genes.per.Seed.Ortholog.hist.w.singletons <- EggNOG_annotations_raw %>% 
  dplyr::count(`Seed ortholog`, name = "Number_Entries") %>% 
  ggplot(aes(x = Number_Entries)) + 
  stat_bin(binwidth = 1) + 
  stat_bin(binwidth = 1, 
           geom = "label", 
           aes(label = ..count..), 
           size = 3, 
           alpha = 0.5,
           vjust = 0) +
  scale_y_log10(labels = scales::trans_format("log10", scales::math_format(10^.x))) +
  annotation_logticks(sides = "l") +
  labs(title = "Number of Query Genes Mapping to Each\nUnique Seed Ortholog (With Singletons)",
       y = "Number of Mapped Query Genes",
       x = "Number of Seed Orthologs") +
  theme(plot.title = element_text(hjust = 0.5, size = 14, face = "bold"),
        axis.title = element_text(size = 12, face = "bold"),
        axis.text = element_text(size = 12),
        panel.grid.major.y = element_blank(),
        panel.grid.minor.y = element_blank())

p.genes.per.Seed.Ortholog.hist.w.singletons

#After removing "Query", "e-value", and "score" columns, there are no 
#are there any duplicated Seed Orthologs after collapsing to unique rows
writeLines("\nAre there any duplicate Seed Orthologs after attempted deduplication?")
EggNOG_annotations_raw %>% 
  dplyr::select(-c(Query, `e-value`, score)) %>% 
  dplyr::distinct() %>% 
  dplyr::pull(`Seed ortholog`) %>% 
  duplicated() %>%
  any()
```

```{r review-annotation-stats-after-eliminating-singletons, echo=FALSE}
EggNOG_annotations_raw_no_singletons <- EggNOG_annotations_raw %>% 
  dplyr::semi_join(gene_pres_abs_Spec_SGB %>% 
                     dplyr::filter(Num_Genomes_Present == "Gene in Multiple Genomes") %>% 
                     dplyr::distinct(Gene_Group),
                   by = c("Query" = "Gene_Group"))

writeLines("\nHow many unique Seed orthologs are in the EggNOG output after eliminating singletons?")
EggNOG_annotations_raw_no_singletons %>% 
  dplyr::distinct(`Seed ortholog`) %>% 
  nrow()

#Plot mapping of Query genes to Seed orthologs
p.genes.per.Seed.Ortholog.hist.no.singletons <- EggNOG_annotations_raw_no_singletons %>% 
  dplyr::count(`Seed ortholog`, name = "Number_Entries") %>% 
  ggplot(aes(x = Number_Entries)) + 
  stat_bin(binwidth = 1) + 
  stat_bin(binwidth = 1, 
           geom = "label", 
           aes(label = ..count..), 
           size = 3, 
           alpha = 0.5,
           vjust = 0) +
  scale_y_log10(labels = scales::trans_format("log10", scales::math_format(10^.x))) +
  annotation_logticks(sides = "l") +
  labs(title = "Number of Query Genes Mapping to Each\nUnique Seed Ortholog (No Singletons)",
       y = "Number of Mapped Query Genes",
       x = "Number of Seed Orthologs") +
  theme(plot.title = element_text(hjust = 0.5, size = 14, face = "bold"),
        axis.title = element_text(size = 12, face = "bold"),
        axis.text = element_text(size = 12),
        panel.grid.major.y = element_blank(),
        panel.grid.minor.y = element_blank())

p.genes.per.Seed.Ortholog.hist.no.singletons

```


```{r identify-unique-EggNOG-OGs-no-singletons, echo=FALSE}
#Define dataframe of unique EggNOG OGs
Unique_EggNOG_OGs_no_singletons <- EggNOG_annotations_raw_no_singletons %>% 
  dplyr::select(-c(Query, `e-value`, score)) %>% 
  dplyr::distinct()
```
  
### Load table of genes/annotations of interest  
  

```{r load-genes-of-interest, echo=FALSE}
f_Cys_gene_wishlist <- "Genes_of_interest/Cys_pathway_genes_of_interest3.txt"

Cys_gene_wishlist <- read_tsv(f_Cys_gene_wishlist)

#Uniquely number each entrie in Cys_gene_wishlist
Cys_gene_wishlist <- Cys_gene_wishlist %>% 
  dplyr::mutate(Unique_gene_num = row_number()) %>% 
  dplyr::select(Unique_gene_num, everything())
```
  
### Review EggNOG output for Seed Orthologs annotated as encoding genes/functions of interest  
  
  

```{r create-list-for-subsetting-matching-EggNOG-output, echo=FALSE}
#Create an empty list of the same length as number of rows in Cys_gene_wishlist,
#each first-level list element is a list with with two elements, "gene_sought" and "NOGs"
genes_list <- map(1:nrow(Cys_gene_wishlist),
                  function(x) {x <- structure(list(gene_sought = NULL,
                                                   NOGs = NULL))})

#For each element in genes_list, add the corresponding row from 
#the Cys_gene_wishlist dataframe to the sub-element gene_sought,
#making gene_sought a dataframe
for (n in seq_along(genes_list)) {
  genes_list[[n]][["gene_sought"]] <- Cys_gene_wishlist[n, ]
  }

#Remove counting variable
rm(n)
```


The next several code chunks search through the EggNOG output to compile a coherent and consistently annotated dataframe of all the EggNOG queries ("Query") and corresponding Seed orthologs ("Seed ortholog") for the _Lactobacillus_ pan-genome that correspond to a specific gene of interest. The seed ortholog is the ortholog in the EggNOG database that is the best match for a query gene. One or more query genes can map to a single seed ortholog.  


Relevant search terms in EggNOG:  
  
* Gene name in column "Preferred_name", case insensitive
* Gene name in column "description", case insensitive
* Str_detect COG group in column "matching OGs" (looks like the search string can be in form of "COG####")
* Str_detect TC number in column "KEGG TC" (search string without TC prefix, need to be sure to escape the "." character so it is not interpreted as a special character)
* Str_detect EC number in column "EC number" (search string without EC prefix, need to be sure to escape the "." character so it is not interpreted as a special character)
* Str_detect KO number in column "KEGG KO" (search string in the form "K#####")
* Str_detect GO term in column "GO terms"
* Will not look by KEGG pathway because this is too non-specific (can encompass multiple genes within a pathway)
* Str_detect KEGG reaction number in column "KEGG reaction"
* Supplemented by manually lookingh for various full-length gene names or keywords in column "description" (e.g. "cysteine synthase" or "cystathionine", case insensitive)  
  

```{r define-function-to-search-EggNOG-genes-by-genes_wishlist, echo=FALSE}
#This function takes two dataframes as arguments:
#EggNOG_output, a multi-row dataframe of EggNOG-annotated genes or Seed Orthologs,
#and gene_sought, a single-row dataframe containing relevant annotation
#terms to query within EggNOG_output
#Output is a dataframe consisting of the the rows in EggNOG_output
#that meet one or more search criterion in gene_sought (0 or more rows)

EggNOG_gene_search <- function(EggNOG_output, gene_sought) {
  
  #Check that EggNOG_output is a dataframe
  if (!is.data.frame(EggNOG_output)) stop("EggNOG_output is not a dataframe")
  
  #Check that gene_sought is a dataframe
  if (!is.data.frame(gene_sought)) stop("gene_sought is not a dataframe")
  
  #Check that gene_sought has only 1 row
  if (!(nrow(gene_sought) == 1)) {
    stop("gene_sought row number does not equal 1")
  }
  
  EggNOG_output %>%
    dplyr::filter(
      str_detect(toupper(Preferred_name), toupper(gene_sought$All_Names)) |
        str_detect(toupper(description), toupper(gene_sought$All_Names)) |
        str_detect(`matching OGs`, gene_sought$COG) |
        str_detect(`KEGG TC`, gene_sought$TC_Numbers) |
        str_detect(`EC number`, gene_sought$EC_number) |
        str_detect(`KEGG KO`, gene_sought$KO) |
        str_detect(`GO terms`, gene_sought$GO_Terms) |
        str_detect(`KEGG reaction`, gene_sought$KEGG_Rxn_Nums)
    )
  
}
```


```{r map-unique-OGs-to-genes-of-interest, echo=FALSE}
#For each entry (first-level element) in genes_list, 
#look for gene of interest in Unique_EggNOG_OGs
#using the EggNOG_gene_search function and the annotation terms in the 
#gene_sought element, storing the output (a dataframe) in the NOGs element
for (n in seq_along(genes_list)) {
  genes_list[[n]][["NOGs"]] <- EggNOG_gene_search(EggNOG_output = Unique_EggNOG_OGs_no_singletons,
                                                  gene_sought = genes_list[[n]][["gene_sought"]])
  }

#Remove looping variable
rm(n)


p.gene.sought.OGs.hist.prefilt <- map_df(genes_list, 
                                         function(x) {tibble(Unique_gene_num = x[["gene_sought"]][["Unique_gene_num"]],
                                                             Number_Matches = nrow(x[["NOGs"]]))}) %>% 
  ggplot(aes(x = Number_Matches)) + 
  geom_histogram(binwidth = 5) +
  labs(title = "Number of Unique Seed Orthologs Mapping to\nEach Gene of Interest (Before Filtering)",
       y = "Number of Genes of Interest",
       x = "Number of Mapped Seed Orthologs") +
  theme(plot.title = element_text(hjust = 0.5, size = 14, face = "bold"),
        axis.title = element_text(size = 12, face = "bold"),
        axis.text = element_text(size = 12),
        panel.grid.major.y = element_blank(),
        panel.grid.minor.y = element_blank())

p.gene.sought.OGs.hist.prefilt
```
  
Manually filter EggNOG_gene_search output to eliminate spurious hits, filtering based on manual review of likely discrepancies.  
  

```{r filter-EggNOG-matches-to-gene-wishlist, echo=FALSE}
#Create list with same structure as genes_list to store output
#of manual filtering step
genes_list_filt <- genes_list

#Filter to retain only genes annotated as serine O-acetyltransferase KEGG KO
genes_list_filt[[1]]$NOGs <- genes_list[[1]]$NOGs %>% 
  dplyr::filter(str_detect(`KEGG KO`, genes_list[[1]]$gene_sought$KO))

#Filter to retain only genes annotated as cysteine synthase KEGG KO
genes_list_filt[[2]]$NOGs <- genes_list[[2]]$NOGs %>% 
  dplyr::filter(str_detect(`KEGG KO`, genes_list[[2]]$gene_sought$KO))

#Filter to retain only genes annotated as catalyzing O-acetyl-L-serine:thiosulfate 2-amino-2-carboxyethyltransferase KEGG reaction
genes_list_filt[[3]]$NOGs <- genes_list[[3]]$NOGs %>% 
  dplyr::filter(str_detect(`KEGG reaction`, genes_list[[3]]$gene_sought$KEGG_Rxn_Nums))

#Filter to retain only genes annotated as catalyzing O-phosphoserine sulfhydrylase KEGG reaction
genes_list_filt[[4]]$NOGs <- genes_list[[4]]$NOGs %>% 
  dplyr::filter(str_detect(`KEGG reaction`, genes_list[[4]]$gene_sought$KEGG_Rxn_Nums))

#Filter to retain only genes encoding cystathionine beta-synthase activity
genes_list_filt[[5]]$NOGs <- genes_list[[5]]$NOGs %>% 
  dplyr::filter(str_detect(`KEGG reaction`, genes_list[[5]]$gene_sought$KEGG_Rxn_Nums) |
                  str_detect(`EC number`, genes_list[[5]]$gene_sought$EC_number))

#Filter to retain only genes encoding O-acetylserine dependent cystathionine beta-synthase activity
genes_list_filt[[6]]$NOGs <- genes_list[[6]]$NOGs %>% 
  dplyr::filter(str_detect(`KEGG reaction`, genes_list[[6]]$gene_sought$KEGG_Rxn_Nums) |
                  str_detect(`EC number`, genes_list[[6]]$gene_sought$EC_number))

#Filter to retain only genes annotated as catalyzing cystathionine gamma-lyase KEGG reaction
genes_list_filt[[7]]$NOGs <- genes_list[[7]]$NOGs %>% 
  dplyr::filter(str_detect(`KEGG reaction`, genes_list[[7]]$gene_sought$KEGG_Rxn_Nums))

#Filter to retain only genes encoding adenosylhomocysteinase activity, not using EC number due to issues with parsing regular expression
genes_list_filt[[8]]$NOGs <- genes_list[[8]]$NOGs %>% 
  dplyr::filter(str_detect(`KEGG KO`, genes_list[[8]]$gene_sought$KO) |
                  str_detect(`KEGG reaction`, genes_list[[8]]$gene_sought$KEGG_Rxn_Nums))

#Not filtering results for ko:K00558 because these all seem correct

#Filter to retain only genes encoding methionine adenosyltransferase activity, not using EC number due to issues with parsing regular expression
genes_list_filt[[10]]$NOGs <- genes_list[[10]]$NOGs %>% 
  dplyr::filter(str_detect(`KEGG KO`, genes_list[[10]]$gene_sought$KO) |
                  str_detect(`KEGG reaction`, genes_list[[10]]$gene_sought$KEGG_Rxn_Nums))

#Not filtering results for ko:K07173 because these all seem correct

#Filter to retain only genes encoding adenosylhomocysteine nucleosidase activity
genes_list_filt[[12]]$NOGs <- genes_list[[12]]$NOGs %>% 
  dplyr::filter(str_detect(`KEGG KO`, genes_list[[12]]$gene_sought$KO) |
                  str_detect(`KEGG reaction`, genes_list[[12]]$gene_sought$KEGG_Rxn_Nums))

#Not filtering results for cystine reductase because there were no matches detected
#Not filtering results for GSH-cystine transhydrogenase because there were no matches detected
#Not filtering results for Cysteine racemase because there were no matches detected

#Filter to retain only genes encoding assimilatory sulfite reductase (ferredoxin) activity
genes_list_filt[[16]]$NOGs <- genes_list[[16]]$NOGs %>% 
  dplyr::filter(str_detect(`KEGG KO`, genes_list[[16]]$gene_sought$KO) |
                  str_detect(`KEGG reaction`, genes_list[[16]]$gene_sought$KEGG_Rxn_Nums))

#Not filtering results for assimilatory sulfite reductase (NADPH) because there were no matches detected
#Not filtering results for dissimilatory sulfite reductase because there were no matches detected
#Not filtering results for sulfite dehydrogenase (cytochrome) because there were no matches detected
#Not filtering results for sulfite oxidase because there were no matches detected
#Not filtering results for sulfite dehydrogenase (quinone) because there were no matches detected

#Filter to retain only genes encoding sulfur oxygenase/reductase activity
genes_list_filt[[22]]$NOGs <- genes_list[[22]]$NOGs %>% 
  dplyr::filter(str_detect(`KEGG KO`, genes_list[[22]]$gene_sought$KO) |
                  str_detect(`KEGG reaction`, genes_list[[22]]$gene_sought$KEGG_Rxn_Nums))

#Filter to retain only genes encoding homocysteine desulfhydrase activity
genes_list_filt[[23]]$NOGs <- genes_list[[23]]$NOGs %>% 
  dplyr::filter(str_detect(`KEGG KO`, genes_list[[23]]$gene_sought$KO) |
                  str_detect(`KEGG reaction`, genes_list[[23]]$gene_sought$KEGG_Rxn_Nums))

#Filter to retain only genes encoding glutathione gamma-glutamate hydrolase activity
genes_list_filt[[24]]$NOGs <- genes_list[[24]]$NOGs %>% 
  dplyr::filter(str_detect(`KEGG KO`, genes_list[[24]]$gene_sought$KO) |
                  str_detect(`KEGG reaction`, genes_list[[24]]$gene_sought$KEGG_Rxn_Nums))

#Not filtering results for glutathione-specific gamma-glutamylcyclotransferase because there were no matches detected

#Filter to retain only genes encoding gamma-glutamyltransferase activity
genes_list_filt[[26]]$NOGs <- genes_list[[26]]$NOGs %>% 
  dplyr::filter(str_detect(`KEGG KO`, genes_list[[26]]$gene_sought$KO) |
                  str_detect(`KEGG reaction`, genes_list[[26]]$gene_sought$KEGG_Rxn_Nums))

#Not filtering results for gamma-glutamylcyclotransferase because there were no matches detected

#Filter to retain only genes encoding membrane alanyl aminopeptidase activity
genes_list_filt[[28]]$NOGs <- genes_list[[28]]$NOGs %>% 
  dplyr::filter(str_detect(`KEGG KO`, genes_list[[28]]$gene_sought$KO) |
                  str_detect(`KEGG reaction`, genes_list[[28]]$gene_sought$KEGG_Rxn_Nums))

#Filter to retain only genes encoding leucyl aminopeptidase activity
genes_list_filt[[29]]$NOGs <- genes_list[[29]]$NOGs %>% 
  dplyr::filter(str_detect(`KEGG KO`, genes_list[[29]]$gene_sought$KO) |
                  str_detect(`KEGG reaction`, genes_list[[29]]$gene_sought$KEGG_Rxn_Nums))

#Filter to retain only genes encoding bis-gamma-glutamylcystine reductase activity
genes_list_filt[[30]]$NOGs <- genes_list[[30]]$NOGs %>% 
  dplyr::filter(str_detect(`KEGG KO`, genes_list[[30]]$gene_sought$KO) |
                  str_detect(`KEGG reaction`, genes_list[[30]]$gene_sought$KEGG_Rxn_Nums))

#Filter to retain only genes encoding gamma-glutamylcysteine synthetase activity, but not those encoded as gshF (bifunctional GSH synthase)
genes_list_filt[[31]]$NOGs <- genes_list[[31]]$NOGs %>% 
  dplyr::filter(str_detect(`KEGG KO`, genes_list[[31]]$gene_sought$KO) |
                  str_detect(`KEGG reaction`, genes_list[[31]]$gene_sought$KEGG_Rxn_Nums),
                Preferred_name != "gshF")

#Filter to retain only genes encoding glutathione synthase activity
genes_list_filt[[32]]$NOGs <- genes_list[[32]]$NOGs %>% 
  dplyr::filter(str_detect(`KEGG KO`, genes_list[[32]]$gene_sought$KO) |
                  str_detect(`KEGG reaction`, genes_list[[32]]$gene_sought$KEGG_Rxn_Nums))

#Not filtering results for gshF, putative glutamate--cysteine ligase because these all seem correct

#Filter to retain only genes encoding CydC (ko:K16012) and replace Preferred_name from "cydD" to "cydC" since this is miscoded in the EggNOG output
genes_list_filt[[34]]$NOGs <- genes_list[[34]]$NOGs %>% 
  dplyr::filter(str_detect(`KEGG KO`, genes_list[[34]]$gene_sought$KO) |
                  str_detect(`matching OGs`, genes_list[[34]]$gene_sought$COG)) %>% 
  dplyr::mutate(Preferred_name = str_replace(Preferred_name, "cydD", "cydC")) 

#Filter to retain only genes encoding CydD (ko:K16013)
genes_list_filt[[35]]$NOGs <- genes_list[[35]]$NOGs %>% 
  dplyr::filter(str_detect(`KEGG KO`, genes_list[[35]]$gene_sought$KO) |
                  str_detect(`matching OGs`, genes_list[[35]]$gene_sought$COG))

#Filter to retain only genes encoding CydA (ko:K00425; COG1271)
genes_list_filt[[36]]$NOGs <- genes_list[[36]]$NOGs %>% 
  dplyr::filter(str_detect(`KEGG KO`, genes_list[[36]]$gene_sought$KO) |
                  str_detect(`matching OGs`, genes_list[[36]]$gene_sought$COG))
	
#Filter to retain only genes encoding CydB (ko:K00426; COG1294)
genes_list_filt[[37]]$NOGs <- genes_list[[37]]$NOGs %>% 
  dplyr::filter(str_detect(`KEGG KO`, genes_list[[37]]$gene_sought$KO) |
                  str_detect(`matching OGs`, genes_list[[37]]$gene_sought$COG))

#Filter to retain only genes encoding TcyA by KO number (ko:K02424).
genes_list_filt[[38]]$NOGs <- genes_list[[38]]$NOGs %>% 
  dplyr::filter(str_detect(`KEGG KO`, genes_list[[38]]$gene_sought$KO))

#Filter to retain only genes encoding TcyB by KO number (ko:K10009).
genes_list_filt[[39]]$NOGs <- genes_list[[39]]$NOGs %>% 
  dplyr::filter(str_detect(`KEGG KO`, genes_list[[39]]$gene_sought$KO))

#Filter to retain only genes encoding TcyC by KO number (ko:K10010).
genes_list_filt[[40]]$NOGs <- genes_list[[40]]$NOGs %>% 
  dplyr::filter(str_detect(`KEGG KO`, genes_list[[40]]$gene_sought$KO))

#Filter to retain only genes encoding TcyJ by KO number (ko:K16956).
genes_list_filt[[41]]$NOGs <- genes_list[[41]]$NOGs %>% 
  dplyr::filter(str_detect(`KEGG KO`, genes_list[[41]]$gene_sought$KO))

#Filter to retain only genes encoding TcyK by KO number (ko:K16957),
#but not the KO number (ko:K01599), which refers to a uroporphyrinogen 
#decarboxylase adjacent to the tcyKLMN locus in L. vaginalis that is
#sometimes misannotated as TcyK (ko:K16957).
genes_list_filt[[42]]$NOGs <- genes_list[[42]]$NOGs %>% 
  dplyr::filter(str_detect(`KEGG KO`, genes_list[[42]]$gene_sought$KO),
                !str_detect(`KEGG KO`, "K01599"))

#Filter to retain only genes encoding TcyL by KO number (ko:K16958).
genes_list_filt[[43]]$NOGs <- genes_list[[43]]$NOGs %>% 
  dplyr::filter(str_detect(`KEGG KO`, genes_list[[43]]$gene_sought$KO))

#Filter to retain only genes encoding TcyM by KO number (ko:K16959).
genes_list_filt[[44]]$NOGs <- genes_list[[44]]$NOGs %>% 
  dplyr::filter(str_detect(`KEGG KO`, genes_list[[44]]$gene_sought$KO))

#Filter to retain only genes encoding TcyN by KO number (ko:K16960).
genes_list_filt[[45]]$NOGs <- genes_list[[45]]$NOGs %>% 
  dplyr::filter(str_detect(`KEGG KO`, genes_list[[45]]$gene_sought$KO))

#Not filtering results for tcyP (ko:K06956), involved in cystine transport, because result seems correct

#Filter to retain only genes encoding gsiB by KO number (ko:K13889).
genes_list_filt[[47]]$NOGs <- genes_list[[47]]$NOGs %>% 
  dplyr::filter(str_detect(`KEGG KO`, genes_list[[47]]$gene_sought$KO))

#Filter to retain only genes encoding gsiC by KO number (ko:K13890).
genes_list_filt[[48]]$NOGs <- genes_list[[48]]$NOGs %>% 
  dplyr::filter(str_detect(`KEGG KO`, genes_list[[48]]$gene_sought$KO))

#Filter to retain only genes encoding gsiD by KO number (ko:K13891).
genes_list_filt[[49]]$NOGs <- genes_list[[49]]$NOGs %>% 
  dplyr::filter(str_detect(`KEGG KO`, genes_list[[49]]$gene_sought$KO))

#Filter to retain only genes encoding gsiA by KO number (ko:K13892).
genes_list_filt[[50]]$NOGs <- genes_list[[50]]$NOGs %>% 
  dplyr::filter(str_detect(`KEGG KO`, genes_list[[50]]$gene_sought$KO))

#Filter to retain only genes encoding dppA by KO number (ko:K12368).
genes_list_filt[[51]]$NOGs <- genes_list[[51]]$NOGs %>% 
  dplyr::filter(str_detect(`KEGG KO`, genes_list[[51]]$gene_sought$KO))

#Filter to retain only genes encoding dppB by KO number (ko:K12369).
genes_list_filt[[52]]$NOGs <- genes_list[[52]]$NOGs %>% 
  dplyr::filter(str_detect(`KEGG KO`, genes_list[[52]]$gene_sought$KO))

#Filter to retain only genes encoding dppC by KO number (ko:K12370).
genes_list_filt[[53]]$NOGs <- genes_list[[53]]$NOGs %>% 
  dplyr::filter(str_detect(`KEGG KO`, genes_list[[53]]$gene_sought$KO))

#Filter to retain only genes encoding dppD by KO number (ko:K12371).
genes_list_filt[[54]]$NOGs <- genes_list[[54]]$NOGs %>% 
  dplyr::filter(str_detect(`KEGG KO`, genes_list[[54]]$gene_sought$KO))

#Filter to retain only genes encoding dppF by KO number (ko:K12372).
genes_list_filt[[55]]$NOGs <- genes_list[[55]]$NOGs %>% 
  dplyr::filter(str_detect(`KEGG KO`, genes_list[[55]]$gene_sought$KO))

#Filter to retain only genes encoding oppA by KO number (ko:K15580).
genes_list_filt[[56]]$NOGs <- genes_list[[56]]$NOGs %>% 
  dplyr::filter(str_detect(`KEGG KO`, genes_list[[56]]$gene_sought$KO))

#Filter to retain only genes encoding oppB by KO number (ko:K15581).
genes_list_filt[[57]]$NOGs <- genes_list[[57]]$NOGs %>% 
  dplyr::filter(str_detect(`KEGG KO`, genes_list[[57]]$gene_sought$KO))

#Filter to retain only genes encoding oppC by KO number (ko:K15582).
genes_list_filt[[58]]$NOGs <- genes_list[[58]]$NOGs %>% 
  dplyr::filter(str_detect(`KEGG KO`, genes_list[[58]]$gene_sought$KO))

#Filter to retain only genes encoding oppD by KO number (ko:K15583).
genes_list_filt[[59]]$NOGs <- genes_list[[59]]$NOGs %>% 
  dplyr::filter(str_detect(`KEGG KO`, genes_list[[59]]$gene_sought$KO))

#Filter to retain only genes encoding oppF by KO number (ko:K10823).
genes_list_filt[[60]]$NOGs <- genes_list[[60]]$NOGs %>% 
  dplyr::filter(str_detect(`KEGG KO`, genes_list[[60]]$gene_sought$KO))

#Filter to retain only genes encoding livK by KO number (ko:K01999).
genes_list_filt[[61]]$NOGs <- genes_list[[61]]$NOGs %>% 
  dplyr::filter(str_detect(`KEGG KO`, genes_list[[61]]$gene_sought$KO))

#Filter to retain only genes encoding livH by KO number (ko:K01997).
genes_list_filt[[62]]$NOGs <- genes_list[[62]]$NOGs %>% 
  dplyr::filter(str_detect(`KEGG KO`, genes_list[[62]]$gene_sought$KO))

#Filter to retain only genes encoding livM by KO number (ko:K01998).
genes_list_filt[[63]]$NOGs <- genes_list[[63]]$NOGs %>% 
  dplyr::filter(str_detect(`KEGG KO`, genes_list[[63]]$gene_sought$KO))

#Filter to retain only genes encoding livG by KO number (ko:K01995).
genes_list_filt[[64]]$NOGs <- genes_list[[64]]$NOGs %>% 
  dplyr::filter(str_detect(`KEGG KO`, genes_list[[64]]$gene_sought$KO))

#Filter to retain only genes encoding livF by KO number (ko:K01996).
genes_list_filt[[65]]$NOGs <- genes_list[[65]]$NOGs %>% 
  dplyr::filter(str_detect(`KEGG KO`, genes_list[[65]]$gene_sought$KO))

#Filter to retain only genes encoding cjaA homologs by KO number (ko:K02030) 
#and containing the word "binding" in the description and "cjaA" in the name.
genes_list_filt[[66]]$NOGs <- genes_list[[66]]$NOGs %>% 
  dplyr::filter(str_detect(`KEGG KO`, genes_list[[66]]$gene_sought$KO),
                Preferred_name == "cjaA",
                str_detect(description, "binding"))
```
  
### Review filtered EggNOG output for Seed Orthologs of interest  
  

```{r review-filtered-EggNOG-matches-histogram, echo=FALSE}
#Table reviewing number of hits after filtering
map_df(genes_list_filt, function(x) {tibble(Unique_gene_num = x[["gene_sought"]][["Unique_gene_num"]],
                                            Pathway = x[["gene_sought"]][["Pathway"]], 
                                            `Enzyme Name` = x[["gene_sought"]][["Enzyme_Name"]], 
                                            `Name to Plot` = x[["gene_sought"]][["Name_to_Plot"]],
                                            `Number Unique Matching Seed Orthologs` = nrow(x[["NOGs"]]))}) %>% 
  knitr::kable(caption = "Number of Unique Seed Orthologs Mapping to\nEach Gene Function of Interest (Post-Filtering)")

#Histogram reviewing number of hits after filtering
p.gene.OG.match.filt <- map_df(genes_list_filt, 
                               function(x) {tibble(Unique_gene_num = x[["gene_sought"]][["Unique_gene_num"]],
                                                   Pathway = x[["gene_sought"]][["Pathway"]],
                                                   Enzyme_Name = x[["gene_sought"]][["Enzyme_Name"]],
                                                   Name_to_Plot = x[["gene_sought"]][["Name_to_Plot"]],
                                                   Number_Matching_OGs = nrow(x[["NOGs"]]))}) %>% 
  ggplot(aes(x = Number_Matching_OGs)) + 
  stat_bin(binwidth = 1) +
  labs(title = "Number of Unique Seed Orthologs Mapping to\nEach Gene Function of Interest (Post-Filtering)",
       y = "# of Gene Functions\nof Interest",
       x = "Number of Unique\nMapped Seed Orthologs") +
  theme(plot.title = element_text(hjust = 0.5, size = 14, face = "bold"),
        axis.title = element_text(size = 12, face = "bold"),
        axis.text = element_text(size = 10)) 

p.gene.OG.match.filt

#Same as above, but faceting based on pathway of interest
p.gene.OG.match.filt.facet <- p.gene.OG.match.filt + 
  scale_x_continuous(limits = c(-0.5, NA),
                     breaks = c(0:18) * 2) +
  facet_wrap(~str_wrap(Pathway, width = 20), scales = "free_x")

p.gene.OG.match.filt.facet
```
  
  
Convert results into dataframe of unique eggNOG seed orthologs matching genes/functions of interest by adding Unique_gene_num, Pathway, and Enzyme_Name to the NOGs dataframe, then rowbinding the dfs together.  
  

```{r rowbind-filtered-EggNOG-OGs, echo=FALSE}
#Create dataframe for rowbinding the filtered EggNOG output
genes_OGs_df <- tibble()

#Add Unique_gene_num, Pathway, and Enzyme_Name to the NOGs dataframe, then rowbind the dfs together
for (n in seq_along(genes_list_filt)) {
  genes_OGs_df <- genes_OGs_df %>% 
    dplyr::bind_rows(genes_list_filt[[n]]$NOGs %>% 
                       dplyr::mutate(Unique_gene_num = genes_list_filt[[n]]$gene_sought$Unique_gene_num,
                                     Pathway = genes_list_filt[[n]]$gene_sought$Pathway,
                                     Name_to_Plot = genes_list_filt[[n]]$gene_sought$Name_to_Plot,
                                     Enzyme_Name = genes_list_filt[[n]]$gene_sought$Enzyme_Name) %>% 
                       dplyr::select(Unique_gene_num:Enzyme_Name, everything()))
  }

#remove looping variable
rm(n)
```
  
### Review Seed orthologs that match multiple genes of interest  
  
If a single Seed ortholog matches multiple genes/annotations of interest, it could indicate uncertainty or problem with the annotation, or it could indicate that one of them is a mis-annotation in the EggNOG output. Therefore manually reviewing all Seed orthologs matching multiple genes/functions.  
  

```{r review-Seed-orthologs-matching-multiple-genes-of-interest, echo=FALSE}
genes_OGs_df %>% 
  dplyr::group_by(`Seed ortholog`) %>% 
  dplyr::mutate(OG_matches = n()) %>% 
  dplyr::ungroup() %>% 
  dplyr::filter(OG_matches > 1,
                !(Unique_gene_num %in% c(28, 29))) %>%
  dplyr::select(OG_matches, Unique_gene_num:Preferred_name) %>% 
  dplyr::arrange(OG_matches, `Seed ortholog`, Unique_gene_num) %>% 
  knitr::kable(caption = "Seed orthologs matching multiple genes of interest")
```
  
  
Observations:  
  
*  Multiple OGs are dually annotated as cystathionine gamma-lyase and homocysteine desulfhydrase. These may actually be polyfunctional proteins.
*  Multiple OGs are triply mapping to gsiA, dppF, and oppF (EggNOG-annotated as oppF). These may actually be polyfunctional proteins. 
  
Check manually:  
  
*  Multiple OGs dually mapping to fliY(tcyA) and tcyB (EggNOG assigned these names as either tcyA or tcyB, but functional annotations cause them to map to both genes/functions from the "wishlist")
*  Single OG dually mapping to tcyC and tcyN (EggNOG-annotated as name tcyN)  
*  Single OG dually mapping to fliY (tcyA) and cjaA.  
  
  
To check: Identify the corresponding gene sequences in fasta form for each Seed ortholog.
They can then be manually queried by BLAST and other approaches including comparison to corresponding _Bacillus_ genes with firmer annotations.  
  
  
To review, first add in all genes that correspond to any eggNOG seed ortholog matching the "gene wish list", then tally the number of unique genes matching each gene function on the gene wish list.   
  

```{r map-individual-genes-to-EggNOG-genes-of-interest, echo=FALSE}
#Left_join annotated EggNOG genes to Seed orthologs corresponding
#to genes of interest
genes_queries_df <- genes_OGs_df %>% 
  dplyr::left_join(EggNOG_annotations_raw_no_singletons %>% 
                     dplyr::select(Query:score),
                   by = c("Seed ortholog"))

#Histogram of unique genes (sequences) for each unique gene (function of interest)
p.gene.query.match <- genes_queries_df %>% 
  dplyr::count(Unique_gene_num, Pathway, name = "Number_Matching_Query_Genes") %>% 
  ggplot(aes(x = Number_Matching_Query_Genes)) + 
  stat_bin(binwidth = 1) +
  labs(title = "Number of Unique Gene Sequences Mapping\nto Genes (Functions) of Interest (Post-Filtering)",
       y = "Number of Genes (Functions) of Interest",
       x = "Number of Unique Gene Sequences per\nGene (Function) of Interest") +
  theme(plot.title = element_text(hjust = 0.5, size = 14, face = "bold"),
        axis.title = element_text(size = 12, face = "bold"),
        axis.text = element_text(size = 10)) 

p.gene.query.match

#As above, but faceted by pathway
p.gene.query.match.facet <- p.gene.query.match + 
  scale_x_continuous(limits = c(-0.5, NA)) +
  facet_wrap(~str_wrap(Pathway, width = 20), scales = "free_x")

p.gene.query.match.facet
```
  
Incorporate the nucleic acid and amino acid gene sequences from genes of interest for review.  
  

```{r functions-to-unlist-fasta-lists, echo=FALSE}
#Define functions to parse sequences from gene query fasta files 
#(which are read in as lists) into strings for inclusion in dataframe 
#of genes of interest and for manual query by BLAST and other approaches.

unlist_DNA_fasta_list <- function(fasta_list_element) {
  #Takes an atomic vector of length one and class "SeqFastadna" with
  #attributes including "Annot", corresponding to the FASTA header,
  #and creates 2-column dataframe with columns Fasta_Header and DNA_Seq,
  #Then convert the column of seqquences to a character vector, which
  #strips its attributes and allows it to be joined to other dataframes
  df = fasta_list_element %>% 
    tibble(Fasta_Header = attributes(.)$Annot,
           DNA_Seq = .) %>% 
    dplyr::mutate(DNA_Seq = as.character(DNA_Seq))
  
  return(df)
  }

unlist_AA_fasta_list <- function(fasta_list_element) {
  #Takes an atomic vector of length one and class "SeqFastaAA" with
  #attributes including "Annot", corresponding to the FASTA header,
  #and creates 2-column dataframe with columns Fasta_Header and DNA_Seq
  df = fasta_list_element %>% 
    tibble(Fasta_Header = attributes(.)$Annot,
           AA_Seq = .) %>% 
    dplyr::mutate(AA_Seq = as.character(AA_Seq))
  
  return(df)
  }
```
  

  
Translate fasta-format DNA sequences for genes of interest into AA sequences, then unlist each gene into a single string and combine them into a dataframes.   
  

```{r subset-to-sequences-of-interest-and-translate, echo=FALSE, warning=FALSE}
#Subset pangenome_fasta_DNA_list_sep to only genes in genes_queries_df
gene_query_fasta_DNA_list_sep <- pangenome_fasta_DNA_list_sep[names(pangenome_fasta_DNA_list_sep) %in% genes_queries_df$Query] 

#Translate gene_query_fasta_DNA_list_sep to AA
gene_query_fasta_AA_list_sep <- gene_query_fasta_DNA_list_sep %>% 
  purrr::map(translate, numcode = 11)

#Assign attributes to gene_query_fasta_AA_list_sep for each sequence,
#using the name and Annot from gene_query_fasta_DNA_list_sep
#and class SeqFastaAA
for (n in names(gene_query_fasta_AA_list_sep)) {
  attributes(gene_query_fasta_AA_list_sep[[n]]) <- attributes(gene_query_fasta_DNA_list_sep[[n]])
  attributes(gene_query_fasta_AA_list_sep[[n]])$class <- "SeqFastaAA"
  }

#Remove looping variable
rm(n)

#For each gene sequence, concatenates the vector containing each of its letters individually
#into a single string DNA
gene_query_fasta_DNA_list <- gene_query_fasta_DNA_list_sep %>% 
  purrr::map(seqinr::c2s)

#Assign attributes to gene_query_fasta_DNA_list for each sequence,
#using the name and Annot from gene_query_fasta_DNA_list_sep
#and class SeqFastaAA
for (n in names(gene_query_fasta_DNA_list)) {
  attributes(gene_query_fasta_DNA_list[[n]]) <- attributes(gene_query_fasta_DNA_list_sep[[n]])
  }
#Remove looping variable
rm(n)

#Convert gene_query_fasta_DNA_list to dataframe
gene_seq_query_DNA_df <- purrr::map_df(gene_query_fasta_DNA_list, unlist_DNA_fasta_list)

#For each gene sequence, concatenates the vector containing each of its letters individually
#into a single string DNA
gene_query_fasta_AA_list <- gene_query_fasta_AA_list_sep %>% 
  purrr::map(seqinr::c2s)

#Assign attributes to gene_query_fasta_AA_list for each sequence,
#using the name and Annot from gene_query_fasta_AA_list_sep
#and class SeqFastaAA
for (n in names(gene_query_fasta_AA_list)) {
  attributes(gene_query_fasta_AA_list[[n]]) <- attributes(gene_query_fasta_AA_list_sep[[n]])
  }
#Remove looping variable
rm(n)

#Convert gene_query_fasta_AA_list to dataframe
gene_seq_query_AA_df <- purrr::map_df(gene_query_fasta_AA_list, unlist_AA_fasta_list) 

#Join dataframes to pair AA and DNA sequences for each gene
gene_seq_query_DNA_AA_df <- gene_seq_query_DNA_df %>% 
  dplyr::left_join(gene_seq_query_AA_df,
                   by = "Fasta_Header")

#Modify the dataframe to split header into columns
#Gene_Group and Gene and then
#remove trailing "*" from AA sequences
gene_seq_query_DNA_AA_df <- gene_seq_query_DNA_AA_df %>%
    dplyr::mutate(No_Carrot = str_remove(Fasta_Header, "^>"),
                  AA_Seq = str_remove(AA_Seq, "\\*$")) %>%
    dplyr::select(Fasta_Header, No_Carrot, everything()) %>%
    tidyr::separate(col = No_Carrot,
                    into = c("Gene_Group", "Gene"),
                    sep = " ")

#Add the AA and DNA sequences for each gene to genes_queries_df
genes_queries_seqs_df <- genes_queries_df %>% 
  dplyr::left_join(gene_seq_query_DNA_AA_df,
                   by = c("Query" = "Gene_Group"))
```

Now manually check gene sequences for potentially problematic seed orthologs that were annotated as matching multiple functions of interest from the "wishlist":  
  
*  Multiple OGs dually mapping to fliY(tcyA) and tcyB (some EggNOG-annotated as tcyA, some as tcyB)
  +  "1423814.HMPREF0549_0653", "1423814.HMPREF0549_0655", "575605.ACQN01000012_gene1311", "575605.ACQN01000013_gene1495"
*  Single OG dually mapping to tcyC and tcyN (EggNOG-annotated as tcyN)
  +  "1423814.HMPREF0549_0319"

To manually curate: Identify the corresponding gene sequences in fasta form for each Seed ortholog.
BLAST them against each other, against nr database and also do alignments with the Bacillus sequences to see which they match and check in IMG/M as needed.

```{r review-OGs-mapping-to-tcyA-and-tcyB, echo=FALSE}
genes_queries_seqs_df %>% 
  dplyr::filter(`Seed ortholog` %in% c("1423814.HMPREF0549_0653", "1423814.HMPREF0549_0655", "575605.ACQN01000012_gene1311", "575605.ACQN01000013_gene1495", "1423814.HMPREF0549_0319")) %>% 
  dplyr::select(Fasta_Header, `EggNOG Preferred name` = Preferred_name, AA_Seq, `Seed ortholog`, Unique_gene_num, Enzyme_Name) %>% 
  tidyr::pivot_wider(names_from = Unique_gene_num, values_from = Enzyme_Name, names_prefix = "Unique gene num ") %>% 
  dplyr::arrange(`EggNOG Preferred name`, `Seed ortholog`) %>% 
  knitr::kable(caption = "Review Genes mapping to tcyA and tcyB")
```
  
On BLASTp against each other and against the homologs from Bacillus subtilis and review in IMG/M, following findings:  
  
*  For the Seed ortholog "1423814.HMPREF0549_0319" dually mapping to tcyC and tcyN, the corresponding sequencing is from _L. vaginalis_ and review using IMG/M shows this gene is in same operon as genes annotated as tcyM, tcyL, and tcyK, suggesting it should actually be tcyN. 
*  For the sequences that were assigned to both tcyA (fliY) and tcyB, it appears that the EggNOG Preferred_name annotation is correct (i.e. for some it says tcyA and for others tcyB).
  +  Everything annotated as tcyA matches ABC transporter substrate-binding domain-containing proteins (also > Bacillus subtilis subsp. 168 | BSU_03610 | TcyA)
    *  Seed orthologs "1423814.HMPREF0549_0655", "575605.ACQN01000012_gene1311"
  +  Everything annotated as tcyB matches amino acid ABC transporter permease proteins (also > Bacillus subtilis subsp. 168 | BSU_03600 | TcyB)
    *  Seed orthologs "1423814.HMPREF0549_0653", "575605.ACQN01000013_gene1495"  
*  Single OG dually mapping to fliY (tcyA) and cjaA. The corresponding sequencing is from _L. vaginalis_ and review using IMG/M shows this gene is in same operon as another gene annotated as tcyB, suggesting it should actually be fliY (tcyA)  
  +  Seed ortholog "1423814.HMPREF0549_0528"
   
  
Now check manually:  
  
*  Multiple OGs dually mapping to tcyC and tcyN (some EggNOG-annotated as tcyC, one also annotated as tcyN)  
  
To check: Identify the corresponding gene sequences in fasta form for each Seed ortholog.  
BLAST them against each other, against nr database and also do alignments with the Bacillus sequences to see which they match.  
  
    
```{r review-OGs-mapping-to-tcyC-and-tcyN, echo=FALSE}
genes_queries_seqs_df %>% 
  dplyr::filter(Unique_gene_num %in% c(40, 45)) %>% 
  dplyr::select(Fasta_Header, `EggNOG Preferred name` = Preferred_name, AA_Seq, `Seed ortholog`, Unique_gene_num, Enzyme_Name) %>% 
  tidyr::pivot_wider(names_from = Unique_gene_num, values_from = Enzyme_Name, names_prefix = "Unique gene num ") %>% 
  dplyr::arrange(`EggNOG Preferred name`, `Seed ortholog`) %>% 
  knitr::kable(caption = "Review Genes mapping to tcyC and tcyN")
```
    
    	

```{r correct-tcy-genes-with-multiple-ortholog-mappings, echo=FALSE}
#Remove rows with tcyB genes incorrectly annotated as tcyA (fliY) and vice versa;
#Remove rows with tcyN gene incorrectly annotates as tcyC
#Remove rows with tcyA (fliY) incorrectly annotated as cjaA
genes_queries_seqs_df <- genes_queries_seqs_df %>% 
  dplyr::filter(!((`Seed ortholog` %in% c("1423814.HMPREF0549_0653", "575605.ACQN01000013_gene1495")) &
                    (Enzyme_Name == "fliY")),
                !((`Seed ortholog` %in% c("1423814.HMPREF0549_0655", "575605.ACQN01000012_gene1311")) & 
                    (Enzyme_Name == "tcyB")),
                !((`Seed ortholog` %in% c("1423814.HMPREF0549_0319")) & 
                    (Enzyme_Name == "tcyC")),
                !((`Seed ortholog` %in% c("1423814.HMPREF0549_0528")) & 
                    (Enzyme_Name == "cjaA"))
                )
```
  
  
Now merge information for curated genes of interest with taxonomy-annotated per-genome gene presence-absence table, but recoding genes that are singletons for the species as absent.  
  

```{r merge-gene-pres-abs-to-genes-of-interest-recoding-singletons-as-absent, echo=FALSE}
#Add presence absence data to genequeries dataframe
#after re-coding per-species singletons as absent
genes_queries_seq_pres_abs <- genes_queries_seqs_df %>% 
  dplyr::left_join(gene_pres_abs_Spec_SGB %>% 
                     dplyr::mutate(Gene_Pres = case_when(Num_Genomes_Present == "Singleton Gene" ~ 0,
                                                         TRUE ~ Gene_Pres)) %>% 
                     select(Genome, Gene_Pres, Gene_Group:Species_SGB),
                   by = c("Query" = "Gene_Group"))
```
  
  
  
### Review genes present in only small number genomes  
  
These may represent contaminated assemblies.  
  

```{r review-genes-present-in-only-small-number-genomes, echo=FALSE}
genes_queries_seq_pres_abs %>% 
  dplyr::group_by(Unique_gene_num, Pathway, Enzyme_Name) %>% 
  dplyr::mutate(Gene_Count = sum(Gene_Pres)) %>% 
  dplyr::ungroup() %>% 
  dplyr::filter(Gene_Pres > 0,
                Gene_Count < 20) %>% 
  dplyr::arrange(Gene_Count, Unique_gene_num) %>% 
  dplyr::select(`# of Genomes Containing Gene` = Gene_Count, 
                Unique_gene_num, 
                Pathway, 
                Enzyme_Name,
                Genome_Renamed, Species, `best tax lvl`, Fasta_Header, 
                ends_with("Seq")) %>% 
  knitr::kable(caption = "Genes present in <20 Genomes")
```
  
**low-prevalence genes (AA sequences):**  
  
  
*  >cysE Bacillus in EggNOG vs L crispatus | present in 9 genomes SAMN04498974 to SAMN04498976 and SAMN04498979 to SAMN04498980 and SAMN04498983 to SAMN04498986 | pan-genome fasta header CDJHKBKG_00251 group_27011 | these genomes are all from a single bioproject PRJNA312549 studying Lactobacillus species isolated from the ileum of commercial turkeys | Protein BLAST hits are 100% to self and 99.37% to proteins from other Lactobacillus crispatus isolates from commercial turkeys. Other best hits are ~50% to other Lactobacillus proteins. No DNA BLAST hits. | This may be real, but appears present in the pan-genome only in genomes from poultry sources
    +  MATEEKYGTNLTFYAYIKDDISRYLLMLRIRELCQYRLNNDTAGIKKLALRLLNHFLRKRIIKLGIKLGFDISINSVGPGIRIDHYGFLAINEKASVGMNCHIYGDVTIGIQNSKDKKGPKLGDNVTIGAGARILGPIRVASNTIIGANAVVIKDVDIE  
*  Genes from the putative LIV locus each detected in only two different _L. iners_ MAGs | MG_205763_bin_1 and MG_185275_bin_23 | pan-genome fasta headers DBJIOIPL_00254 braC, DBJIOIPL_00258 livH, DBJIOIPL_00257 group_35825, DBJIOIPL_00256 group_35824, DBJIOIPL_00255 livF | protein BLAST hits are ~70% - 80% identical to proteins from Clostridiales or Treponema. | No relevant DNA BLAST hits | Thus unclear if it is real or a contaminant and if a contaminant where it comes from.
  

**low-prevalence genes (DNA sequences)**  
  
    
*  >cysE Bacillus in EggNOG vs L crispatus | present in 9 genomes SAMN04498974 to SAMN04498977 and SAMN04498979 to SAMN04498980 and SAMN04498983 to SAMN04498986 | pan-genome fasta header CDJHKBKG_00251 group_27011 | these genomes are all from a single bioproject PRJNA312549 studying Lactobacillus species isolated from the ileum of commercial turkeys | Protein BLAST hits are 100% to self and 99.37% to proteins from other Lactobacillus crispatus isolates from commercial turkeys. Other best hits are ~50% to other Lactobacillus proteins. No DNA BLAST hits. | This may be real, but appears present in the pan-genome only in genomes from poultry sources genomes are all from a single bioproject PRJNA312549 studying Lactobacillus species isolated from the ileum of commercial turkeys
    +  atggcgacagaagaaaaatatggtaccaatttgacgttctatgcatacattaaggacgatattagtagatatcttcttatgttaaggataagggaactatgtcaatataggttgaataatgatactgctggaattaaaaaactggcactgagattattaaatcattttcttagaaaaagaattataaaattaggtattaaacttggattcgatatttcgattaatagcgttgggcctggtataagaatagatcattatggatttctagcaataaatgaaaaagcgtctgtaggtatgaattgccatatttatggcgatgtgactattggcatacaaaattcaaaagataaaaaaggtcctaagttgggtgataatgttaccataggggctggggcaagaattcttggaccaataagagttgcctcaaataccataattggtgccaatgcagtagttattaaagatgtagatattgagtga  
  
    

### Review cydABCD, cjaA, liv, and tcy genes in _L. iners_ genomes  
  

```{r review-cyd-genes-in-L-iners, echo=FALSE}
genes_queries_seq_pres_abs %>% 
  dplyr::group_by(Unique_gene_num, Pathway, Enzyme_Name, DNA_Seq, Species) %>% 
  dplyr::mutate(Gene_Count = sum(Gene_Pres)) %>% 
  dplyr::ungroup() %>% 
  dplyr::filter(Gene_Pres > 0,
                str_detect(Name_to_Plot, "cja|cyd|tcy|liv"),
                Species == "Lactobacillus_iners"
                ) %>% 
  dplyr::arrange(Unique_gene_num, Gene_Count, DNA_Seq, Species) %>% 
  dplyr::select(Gene_Count, Species, Genome_Renamed, Unique_gene_num, Name_to_Plot, Query) %>% 
  knitr::kable()
```
  
*  There are no cyd genes after eliminating singletons.
*  There are no cja genes
*  There are two (out of >300) MAGs that contain predicted orthologs of the liv genes  
  
  
  
### Review genes with no EggNOG matches in any species  
  

```{r construct-gene-absence-table-for-wishlist-entries-with-no-EggNOG-match, echo=FALSE}
#Create a dataframe with a unique row for each genome
gene_pres_abs_Spec_SGB_empty_wide <- gene_pres_abs_Spec_SGB %>% 
  dplyr::distinct(Genome, Genome_Gene_Count, SGB, Species, Species_SGB)

#Created columns in the dataframe for each enzyme in the the wishlist with value 0
for (x in Cys_gene_wishlist[["Enzyme_Name"]]) {
  gene_pres_abs_Spec_SGB_empty_wide[[x]] <- replicate(nrow(gene_pres_abs_Spec_SGB_empty_wide), 0)
  }

#Remove counting variable
rm(x)

#Pivot genes and their presence-absence (all 0) values to longer form
gene_pres_abs_Spec_SGB_empty_long_all <- gene_pres_abs_Spec_SGB_empty_wide %>% 
  tidyr::pivot_longer(cols = -c(Genome:Species_SGB), names_to = "Enzyme_Name", values_to = "Gene_Pres")

#Left_join gene_pres_abs_Spec_SGB_empty_long_all to Cys_gene_wishlist,
#after selecting colums, rename, and format to ensure compatibility with 
#corresponding columns in genes_queries_seq_pres_abs
genes_queries_empty_long_all <- Cys_gene_wishlist %>% 
  dplyr::select(Unique_gene_num:Enzyme_Name,
                Name_to_Plot,
                `GO terms` = GO_Terms, `EC number` = EC_number, 
                `KEGG KO` = KO, `KEGG pathway` = KEGG_Pathways, 
                `KEGG module` = KEGG_modules, `KEGG reaction` = KEGG_Rxn_Nums,
                `KEGG TC` = TC_Numbers) %>% 
  dplyr::mutate(`GO terms` = str_replace_all(`GO terms`, "\\|", ","),
                `KEGG KO` = str_replace_all(`KEGG KO`, "\\|", ",") %>% 
                  str_replace_all("K", "ko:K"),
                `KEGG pathway` = str_replace_all(`KEGG pathway`, "\\|", ","),
                `KEGG module` = str_replace_all(`KEGG module`, "\\|", ","),
                `KEGG reaction` = str_replace_all(`KEGG reaction`, "\\|", ","),
                `KEGG TC` = str_replace_all(`KEGG TC`, "\\|", ",")) %>% 
  dplyr::left_join(gene_pres_abs_Spec_SGB_empty_long_all,
                   by = "Enzyme_Name")


#Review genes without-matches in EggNOG
genes_queries_empty_long_all %>%
  dplyr::anti_join(genes_queries_seq_pres_abs,
                   by = "Enzyme_Name") %>%
  dplyr::group_by(Unique_gene_num, Pathway, Name_to_Plot, Enzyme_Name) %>% 
  dplyr::summarise(Gene_Count = sum(Gene_Pres)) %>% 
  knitr::kable(caption = "Genes without matches in EggNOG")

#Generate dataframe of empty gene_pres_absence for genes not present in EggNOG results
genes_queries_empty_long_no_EggNOG <- genes_queries_empty_long_all %>%
  dplyr::anti_join(genes_queries_seq_pres_abs,
                   by = "Enzyme_Name")

#Row_bind remaining genes_queries_empty_long_no_EggNOG to 
#genes_queries_seq_pres_abs

genes_queries_seq_pres_abs_all <- dplyr::bind_rows(genes_queries_seq_pres_abs,
                                                   genes_queries_empty_long_no_EggNOG) %>% 
  dplyr::arrange(Unique_gene_num)
```
  
  
### Review presence and absence of Cys biosynthesis pathways  
  
  
### Cysteine synthesis heatmap figure  
  

```{r plot-gene-presence-absence-Cys-synth-paper-figure, echo=FALSE}
#Create dataframe of presence-absence for selectged pathways by Species
cys.synth.gene.pres.Spec <- genes_queries_seq_pres_abs_all %>% 
  dplyr::filter(Pathway %in% c("Cysteine synthase", "Reverse transsulfuration"),
                !(Enzyme_Name %in% c("membrane alanyl aminopeptidase", "leucyl aminopeptidase")),
                !(Enzyme_Name %in% c("adenosylhomocysteinase", "DNA (cytosine-5-)-methyltransferase",
                                     "methionine adenosyltransferase", "S-ribosylhomocysteine lyase",
                                     "adenosylhomocysteine nucleosidase"))) %>% 
  dplyr::group_by(Unique_gene_num, Pathway, Name_to_Plot, Enzyme_Name, Genome, Species) %>% 
  dplyr::summarise(Gene_Pres = case_when(sum(Gene_Pres) >= 1 ~ "Present",
                                         TRUE ~ "Absent")) %>% 
  dplyr::ungroup() %>% 
  dplyr::arrange(Unique_gene_num) %>% 
  dplyr::group_by(Unique_gene_num, Pathway, Name_to_Plot, Enzyme_Name, Species) %>% 
  dplyr::mutate(Genome_Count = n()) %>% 
  dplyr::ungroup() 

#Plot gene presence-absence for selected pathways by Species
p.cys.synth.gene.pres.rast.Spec <- cys.synth.gene.pres.Spec %>% 
  dplyr::mutate(Enzyme_Name = str_replace_all(Enzyme_Name, " ", "\n"),
                Enzyme_Name = fct_inorder(Enzyme_Name),
                Name_to_Plot = str_replace(Name_to_Plot, "CBS", "cbs"),
                Name_to_Plot = fct_inorder(Name_to_Plot),
                Pathway = str_replace(Pathway, "synthase", "Synthase") %>% 
                  str_replace("transsulfuration", "Trans- Sulfuration"),
                Pathway = str_replace_all(Pathway, " ", "\n"),
                Pathway = fct_inorder(Pathway),
                Species_abbrev = str_replace(Species, "actobacillus_", ". "),
                Species_abbrev_count = paste0(Species_abbrev,
                                              "\n(n=",
                                              Genome_Count,
                                              ")"),
                Species_abbrev_count = factor(Species_abbrev_count,
                                              levels = c("L. crispatus\n(n=508)",
                                                         "L. iners\n(n=327)",
                                                         "L. jensenii\n(n=137)",
                                                         "L. gasseri\n(n=216)",
                                                         "L. vaginalis\n(n=30)")),
                Gene_Pres = paste0("Gene\n", Gene_Pres)) %>% 
  ggplot() +
  geom_raster(aes(x = Name_to_Plot, y = Genome, fill = Gene_Pres)) +
  scale_fill_manual(values = c("black", "yellow")) +
  labs(x = "Gene", fill = "Gene Presence") +
  theme(axis.text.y = element_blank(),
        axis.ticks.y = element_blank(),
        axis.text.x = element_text(size = 9, angle = 45, hjust = 1),
        strip.text.x = element_text(size = 10),
        # legend.title = element_text(hjust = 0.5, size = 12, face = "bold"),
        legend.title = element_blank(),
        legend.text = element_text(size = 10, hjust = 0.5),
        legend.position = "bottom",
        legend.background = element_rect(colour = 'black', fill = 'white', linetype='solid'),
        axis.title = element_text(size = 12, face = "bold"),
        strip.text.y.right = element_text(angle = 0, face = "italic", size = 10)) +
  guides(fill = guide_legend(title.position="top", title.hjust = 0.5)) +
  facet_grid(Species_abbrev_count ~ Pathway, scales = "free", space = "free_x")

p.cys.synth.gene.pres.rast.Spec
```
  


```{r count-gene-numbers-per-genome-for-cys-biosynthesis, echo=FALSE}
#Make dataframe of selected cysteine/cystine transport and GSH synthesis genes and 
#count number of  gene function of interest in each genome (as opposed to presence-absence)
cys.synth.gene.count.Spec <- genes_queries_seq_pres_abs_all %>% 
  dplyr::filter(Pathway %in% c("Cysteine synthase", "Reverse transsulfuration"),
                !(Enzyme_Name %in% c("membrane alanyl aminopeptidase", "leucyl aminopeptidase")),
                !(Enzyme_Name %in% c("adenosylhomocysteinase", "DNA (cytosine-5-)-methyltransferase",
                                     "methionine adenosyltransferase", "S-ribosylhomocysteine lyase",
                                     "adenosylhomocysteine nucleosidase"))) %>% 
  dplyr::group_by(Unique_gene_num, Pathway, Name_to_Plot, Enzyme_Name, Genome, Species) %>% 
  dplyr::summarise(Gene_count = sum(Gene_Pres) %>% 
                     as.character()) %>% 
  dplyr::ungroup() %>% 
  dplyr::arrange(Unique_gene_num) %>% 
  dplyr::group_by(Unique_gene_num, Pathway, Name_to_Plot, Enzyme_Name, Species) %>% 
  dplyr::mutate(Genome_Count = n()) %>% 
  dplyr::ungroup() 

#Plot heat map of above df with fill as number of genes per genome
p.cys.synth.gene.count.rast.Spec <- cys.synth.gene.count.Spec %>% 
  dplyr::mutate(Enzyme_Name = str_replace_all(Enzyme_Name, " ", "\n"),
                Enzyme_Name = fct_inorder(Enzyme_Name),
                Name_to_Plot = str_replace(Name_to_Plot, "CBS", "cbs"),
                Name_to_Plot = fct_inorder(Name_to_Plot),
                Pathway = str_replace(Pathway, " ", "\n"),
                Pathway = fct_inorder(Pathway),
                Species_abbrev = str_replace(Species, "actobacillus_", ". "),
                Species_abbrev_count = paste0(Species_abbrev,
                                              "\n(n=",
                                              Genome_Count,
                                              ")"),
                Species_abbrev_count = factor(Species_abbrev_count,
                                              levels = c("L. crispatus\n(n=508)",
                                                         "L. iners\n(n=327)",
                                                         "L. jensenii\n(n=137)",
                                                         "L. gasseri\n(n=216)",
                                                         "L. vaginalis\n(n=30)"))) %>% 
  ggplot() +
  geom_raster(aes(x = Name_to_Plot, y = Genome, fill = Gene_count)) +
  labs(x = "Gene", fill = "Gene\nCount") +
  # scale_fill_manual(values = c("black", "blue", "yellow", "red", "lightblue", "green")) +
  theme(axis.text.y = element_blank(),
        axis.ticks.y = element_blank(),
        axis.text.x = element_text(size = 12, angle = 45, hjust = 1),
        strip.text.x = element_text(size = 11, face = "bold"),
        legend.title = element_text(hjust = 0.5, size = 12, face = "bold"),
        legend.text = element_text(size = 12),
        axis.title = element_text(size = 12, face = "bold"),
        strip.text.y.right = element_text(angle = 0, face = "italic", size = 11)) +
  facet_grid(Species_abbrev_count ~ Pathway, scales = "free", space = "free_x")

p.cys.synth.gene.count.rast.Spec

#Make stacked percent bar chart of number of copies of each gene of interest in each
#genome per gene-species combination.
p.cys.synth.gene.count.bar.Spec <- cys.synth.gene.count.Spec %>% 
  dplyr::group_by(Name_to_Plot, Species, Gene_count) %>% 
  dplyr::mutate(Gene_Count_tally = n()) %>% 
  dplyr::ungroup() %>% 
  dplyr::select(-Genome) %>% 
  dplyr::distinct() %>% 
  dplyr::mutate(Enzyme_Name = str_replace_all(Enzyme_Name, " ", "\n"),
                Enzyme_Name = fct_inorder(Enzyme_Name),
                Name_to_Plot = str_replace(Name_to_Plot, "CBS", "cbs"),
                Name_to_Plot = fct_inorder(Name_to_Plot),
                Pathway = str_replace(Pathway, " ", "\n"),
                Pathway = fct_inorder(Pathway),
                Species_abbrev = str_replace(Species, "actobacillus_", ". "),
                Species_abbrev_count = paste0(Species_abbrev,
                                              "\n(n=",
                                              Genome_Count,
                                              ")"),
                Species_abbrev_count = factor(Species_abbrev_count,
                                              levels = c("L. crispatus\n(n=508)",
                                                         "L. iners\n(n=327)",
                                                         "L. jensenii\n(n=137)",
                                                         "L. gasseri\n(n=216)",
                                                         "L. vaginalis\n(n=30)"))) %>% 
  ggplot() +
  geom_col(aes(x = Name_to_Plot, y = Gene_Count_tally, fill = Gene_count),
           position = position_fill()) +
  scale_y_continuous(labels = scales::percent_format(),
                     breaks = c(0, 0.5, 1)) + 
  # scale_fill_manual(values = c("black", "blue", "yellow", "red", "lightblue", "green")) +
  labs(x = "Gene", fill = "Gene\nCount") +
  theme(axis.text.x = element_text(size = 12, angle = 45, hjust = 1),
        strip.text.x = element_text(size = 11, face = "bold"),
        legend.title = element_text(hjust = 0.5, size = 12, face = "bold"),
        legend.text = element_text(size = 12),
        axis.title = element_text(size = 12, face = "bold"),
        strip.text.y.right = element_text(angle = 0, face = "italic", size = 11)) +
  facet_grid(Species_abbrev_count ~ Pathway, scales = "free", space = "free_x")

p.cys.synth.gene.count.bar.Spec
```
  
```{r save-cys-synthesis-heatmap-as-PDF-and-RDS-no-singletons, echo=FALSE}
###File name to save for PDF
f.pdf_p.cys.synth.gene.pres.rast.Spec <- "Cys_synth_gene_heatmap_fig_v4_no_singletons.pdf"
f.pdf_p.cys.synth.gene.pres.rast.Spec

### Write PDF to file
cowplot::ggsave2(filename = f.pdf_p.cys.synth.gene.pres.rast.Spec,
                 plot = p.cys.synth.gene.pres.rast.Spec,
                 path = path_output,
                 width = 5.5,
                 height = 4.5)

###File name to save for RDS
f.RDS_p.cys.synth.gene.pres.rast.Spec <- "Cys_synth_gene_heatmap_fig_v4_no_singletons.RDS"
f.RDS_p.cys.synth.gene.pres.rast.Spec

### Write PDF to file
readr::write_rds(p.cys.synth.gene.pres.rast.Spec,
                 file.path(path_output,
                           f.RDS_p.cys.synth.gene.pres.rast.Spec))
```
  
  
### Investigate glutathione synthesis and cys/glutathione transport pathways  
  

```{r review-unique-cystine-transport-pathway-genes, echo=FALSE}
genes_queries_seq_pres_abs_all %>% 
  dplyr::filter(Pathway %in% c("Cystine Transport")) %>%
  dplyr::filter(Gene_Pres == 1) %>% 
  dplyr::group_by(Species, Unique_gene_num, Pathway, Name_to_Plot, 
                  Preferred_name, Enzyme_Name, `best tax lvl`, Fasta_Header) %>% 
  # dplyr::group_by(Species, Unique_gene_num, Fasta_Header) %>% 
  dplyr::count(name = "Num_Genes") %>% 
  dplyr::ungroup() %>% 
  dplyr::select(Num_Genes, everything()) %>% 
  dplyr::distinct() %>% 
  dplyr::arrange(Unique_gene_num, Species, desc(Num_Genes)) %>% 
  knitr::kable(caption = "Number of genes in cystine transport category by species")
```
  
```{r review-unique-cysteine-transport-pathway-genes, echo=FALSE}
genes_queries_seq_pres_abs_all %>% 
  dplyr::filter(Pathway %in% c("Cysteine Transport")) %>%
  dplyr::filter(Gene_Pres == 1) %>% 
  dplyr::group_by(Species, Unique_gene_num, Pathway, Name_to_Plot, 
                  Preferred_name, Enzyme_Name, `best tax lvl`, Fasta_Header) %>% 
  # dplyr::group_by(Species, Unique_gene_num, Fasta_Header) %>% 
  dplyr::count(name = "Num_Genes") %>% 
  dplyr::ungroup() %>% 
  dplyr::select(Num_Genes, everything()) %>% 
  dplyr::distinct() %>% 
  dplyr::arrange(Unique_gene_num, Species, desc(Num_Genes)) %>% 
  knitr::kable(caption = "Number of genes in cysteine transport category (cjaA) by species")
```
  
Notably, _L. iners_ has no predicted intact cystine transport pathway.  
  
```{r review-unique-BCAA-transport-pathway-genes, echo=FALSE}
genes_queries_seq_pres_abs_all %>% 
  dplyr::filter(Pathway %in% c("Branched Chain aa Transport")) %>%
  dplyr::filter(Gene_Pres == 1) %>% 
  dplyr::group_by(Species, Unique_gene_num, Pathway, Name_to_Plot, 
                  Preferred_name, Enzyme_Name, `best tax lvl`, Fasta_Header) %>% 
  # dplyr::group_by(Species, Unique_gene_num, Fasta_Header) %>% 
  dplyr::count(name = "Num_Genes") %>% 
  dplyr::ungroup() %>% 
  dplyr::select(Num_Genes, everything()) %>% 
  dplyr::distinct() %>% 
  dplyr::arrange(Unique_gene_num, Species, desc(Num_Genes)) %>% 
  knitr::kable(caption = "Number of genes in branched chain aa transport category by species")
```
  
_L. iners_ has two genomes annotated as possessing the LIV complex, no other species have any.  
  
  
### Figure for paper: Cys and GSH transport    
  

```{r plot-Cys-and-GSH-transport-and-gsh-synthesis-for-paper-all-combined, echo=FALSE}
#Make dataframe of selected cysteine/cystine transport and GSH transport genes,
#Also rename fliY to tcyA for consistency with literature cited in paper.
cys.transport.gene.pres.Spec <- genes_queries_seq_pres_abs_all %>% 
  dplyr::filter(Pathway %in% c("Cysteine Transport",
                               "Glutathione and cysteine transport", 
                               "Cystine Transport", 
                               "Branched Chain aa Transport",
                               "Glutathione Transport", 
                               "Cytochrome bd synthesis")) %>% 
  dplyr::group_by(Unique_gene_num, Pathway, Name_to_Plot, Enzyme_Name, Genome, Species) %>% 
  dplyr::summarise(Gene_Pres = case_when(sum(Gene_Pres) >= 1 ~ "Present",
                                         TRUE ~ "Absent")) %>% 
  dplyr::ungroup() %>% 
  dplyr::arrange(Unique_gene_num) %>% 
  dplyr::group_by(Unique_gene_num, Pathway, Name_to_Plot, Enzyme_Name, Species) %>% 
  dplyr::mutate(Genome_Count = n()) %>% 
  dplyr::ungroup() %>% 
  dplyr::mutate(Name_to_Plot = case_when(Name_to_Plot == "fliY" ~ "tcyA",
                                         TRUE ~ Name_to_Plot))

#Plot presence-absence heat map of above df
p.cys.transport.all.gene.pres.rast.Spec.fig <- cys.transport.gene.pres.Spec %>% 
  dplyr::mutate(Pathway = case_when(Pathway %in% c("Glutathione and cysteine transport", "Cytochrome bd synthesis") ~
                                      "Cys/GSH Transport\n& Redox\nHomeostasis",
                                    Pathway == "Branched Chain aa Transport" ~ 
                                      "Branched\nChain aa\nTransport",
                                    TRUE ~ str_replace_all(Pathway, " ", "\n")),
                Pathway = str_remove(Pathway, "teine"),
                Pathway = factor(Pathway, 
                                 levels = c("Cys\nTransport",
                                            "Cys/GSH Transport\n& Redox\nHomeostasis",
                                            "Cystine\nTransport",
                                            "Branched\nChain aa\nTransport",
                                            "Glutathione\nTransport")),
                Species_abbrev = str_replace(Species, "actobacillus_", ". "),
                Species_abbrev_count = paste0(Species_abbrev,
                                              "\n(n=",
                                              Genome_Count,
                                              ")"),
                Species_abbrev_count = factor(Species_abbrev_count,
                                              levels = c("L. crispatus\n(n=508)",
                                                         "L. iners\n(n=327)",
                                                         "L. jensenii\n(n=137)",
                                                         "L. gasseri\n(n=216)",
                                                         "L. vaginalis\n(n=30)")),
                Gene_Pres = paste0("Gene\n", Gene_Pres)) %>% 
  dplyr::arrange(Pathway, Name_to_Plot) %>% 
  dplyr::mutate(Enzyme_Name = str_replace_all(Enzyme_Name, " ", "\n"),
                Enzyme_Name = fct_inorder(Enzyme_Name),
                Name_to_Plot = fct_inorder(Name_to_Plot)) %>% 
  ggplot() +
  geom_raster(aes(x = Name_to_Plot, y = Genome, fill = Gene_Pres)) +
  scale_fill_manual(values = c("black", "yellow")) +
  labs(x = "Gene", fill = "Gene\nPresence") +
  theme(axis.text.y = element_blank(),
        axis.ticks.y = element_blank(),
        axis.text.x = element_text(size = 9, angle = 45, hjust = 1),
        strip.text.x = element_text(size = 10),
        # legend.title = element_text(hjust = 0.5, size = 12, face = "bold"),
        legend.title = element_blank(),
        legend.text = element_text(size = 10, hjust = 0.5),
        legend.position = "bottom",
        legend.background = element_rect(colour = 'black', fill = 'white', linetype='solid'),
        axis.title = element_text(size = 12, face = "bold"),
        strip.text.y.right = element_text(angle = 0, face = "italic", size = 10)) +
  guides(fill = guide_legend(title.position="top", title.hjust = 0.5)) +
  facet_grid(Species_abbrev_count ~ Pathway, scales = "free", space = "free_x")

p.cys.transport.all.gene.pres.rast.Spec.fig
```
  
```{r plot-Cys-and-GSH-transport-and-gsh-synthesis-for-paper-cydABCD-separated, echo=FALSE}
#Plot presence-absence heat map of cydABCD only
p.cydABCD.gene.pres.rast.Spec.fig <- cys.transport.gene.pres.Spec %>% 
  dplyr::mutate(Pathway = case_when(Pathway %in% c("Glutathione and cysteine transport", "Cytochrome bd synthesis") ~
                                      "Cys/GSH Transport\n& Redox\nHomeostasis",
                                    TRUE ~ str_replace_all(Pathway, " ", "\n")),
                Species_abbrev = str_replace(Species, "actobacillus_", ". "),
                Species_abbrev_count = paste0(Species_abbrev,
                                              "\n(n=",
                                              Genome_Count,
                                              ")"),
                Species_abbrev_count = factor(Species_abbrev_count,
                                              levels = c("L. crispatus\n(n=508)",
                                                         "L. iners\n(n=327)",
                                                         "L. jensenii\n(n=137)",
                                                         "L. gasseri\n(n=216)",
                                                         "L. vaginalis\n(n=30)")),
                Gene_Pres = paste0("Gene\n", Gene_Pres)) %>% 
  dplyr::arrange(Pathway, Name_to_Plot) %>% 
  dplyr::mutate(Enzyme_Name = str_replace_all(Enzyme_Name, " ", "\n"),
                Enzyme_Name = fct_inorder(Enzyme_Name),
                Name_to_Plot = fct_inorder(Name_to_Plot)) %>% 
  dplyr::filter(Pathway == "Cys/GSH Transport\n& Redox\nHomeostasis") %>% 
  ggplot() +
  geom_raster(aes(x = Name_to_Plot, y = Genome, fill = Gene_Pres)) +
  scale_fill_manual(values = c("black", "yellow")) +
  labs(x = "Gene", fill = "Gene\nPresence") +
  theme(axis.text.y = element_blank(),
        axis.ticks.y = element_blank(),
        axis.text.x = element_text(size = 9, angle = 45, hjust = 1),
        strip.text.x = element_text(size = 10),
        # legend.title = element_text(hjust = 0.5, size = 12, face = "bold"),
        legend.title = element_blank(),
        legend.text = element_text(size = 10, hjust = 0.5),
        # legend.position = "bottom",
        legend.background = element_rect(colour = 'black', fill = 'white', linetype='solid'),
        axis.title = element_text(size = 12, face = "bold"),
        strip.text.y.right = element_text(angle = 0, face = "italic", size = 10)) +
  guides(fill = guide_legend(keyheight = 2)) +
  facet_grid(Species_abbrev_count ~ Pathway, scales = "free", space = "free_x")

p.cydABCD.gene.pres.rast.Spec.fig

#Plot presence-absence heat map of Glutathion and cystine transport without cydABCD
p.cystine.GSH.transport.no.cydABCD.gene.pres.rast.Spec.fig <- cys.transport.gene.pres.Spec %>% 
  dplyr::mutate(Pathway = case_when(Pathway %in% c("Glutathione and cysteine transport", "Cytochrome bd synthesis") ~
                                      "Cys/GSH Transport\n& Redox\nHomeostasis",
                                    TRUE ~ str_replace_all(Pathway, " ", "\n")),
                Pathway = factor(Pathway, 
                                 levels = c("Cys/GSH Transport\n& Redox\nHomeostasis",
                                            "Cystine\nTransport",
                                            "Glutathione\nTransport")),
                Species_abbrev = str_replace(Species, "actobacillus_", ". "),
                Species_abbrev_count = paste0(Species_abbrev,
                                              "\n(n=",
                                              Genome_Count,
                                              ")"),
                Species_abbrev_count = factor(Species_abbrev_count,
                                              levels = c("L. crispatus\n(n=508)",
                                                         "L. iners\n(n=327)",
                                                         "L. jensenii\n(n=137)",
                                                         "L. gasseri\n(n=216)",
                                                         "L. vaginalis\n(n=30)")),
                Gene_Pres = paste0("Gene\n", Gene_Pres)) %>% 
  dplyr::arrange(Pathway, Name_to_Plot) %>% 
  dplyr::mutate(Enzyme_Name = str_replace_all(Enzyme_Name, " ", "\n"),
                Enzyme_Name = fct_inorder(Enzyme_Name),
                Name_to_Plot = fct_inorder(Name_to_Plot)) %>% 
  dplyr::filter(Pathway != "Cys/GSH Transport\n& Redox\nHomeostasis") %>% 
  ggplot() +
  geom_raster(aes(x = Name_to_Plot, y = Genome, fill = Gene_Pres)) +
  scale_fill_manual(values = c("black", "yellow")) +
  labs(x = "Gene", fill = "Gene\nPresence") +
  theme(axis.text.y = element_blank(),
        axis.ticks.y = element_blank(),
        axis.text.x = element_text(size = 9, angle = 45, hjust = 1),
        strip.text.x = element_text(size = 10),
        # legend.title = element_text(hjust = 0.5, size = 12, face = "bold"),
        legend.title = element_blank(),
        legend.text = element_text(size = 10, hjust = 0.5),
        # legend.position = "bottom",
        legend.background = element_rect(colour = 'black', fill = 'white', linetype='solid'),
        axis.title = element_text(size = 12, face = "bold"),
        strip.text.y.right = element_text(angle = 0, face = "italic", size = 10)) +
  guides(fill = guide_legend(keyheight = 2)) +
  facet_grid(Species_abbrev_count ~ Pathway, scales = "free", space = "free_x")

p.cystine.GSH.transport.no.cydABCD.gene.pres.rast.Spec.fig

#Cys/GSH, Cys, and cystine transport
p.cys.cystine.cydABCD.transport.all.gene.pres.rast.Spec.fig <- cys.transport.gene.pres.Spec %>% 
  dplyr::mutate(Pathway = case_when(Pathway %in% c("Glutathione and cysteine transport", "Cytochrome bd synthesis") ~
                                      "Cys/GSH Transport\n& Redox\nHomeostasis",
                                    Pathway == "Branched Chain aa Transport" ~ 
                                      "Branched\nChain aa\nTransport",
                                    Pathway == "Cysteine Transport" ~ 
                                      "Cys\nTran-\nsport",
                                    TRUE ~ str_replace_all(Pathway, " ", "\n")),
                Pathway = str_remove(Pathway, "teine"),
                Pathway = factor(Pathway, 
                                 levels = c("Cys\nTran-\nsport",
                                            "Cystine\nTransport",
                                            "Cys/GSH Transport\n& Redox\nHomeostasis",
                                            "Branched\nChain aa\nTransport",
                                            "Glutathione\nTransport")),
                Species_abbrev = str_replace(Species, "actobacillus_", ". "),
                Species_abbrev_count = paste0(Species_abbrev,
                                              "\n(n=",
                                              Genome_Count,
                                              ")"),
                Species_abbrev_count = factor(Species_abbrev_count,
                                              levels = c("L. crispatus\n(n=508)",
                                                         "L. iners\n(n=327)",
                                                         "L. jensenii\n(n=137)",
                                                         "L. gasseri\n(n=216)",
                                                         "L. vaginalis\n(n=30)")),
                Gene_Pres = paste0("Gene\n", Gene_Pres)) %>% 
  dplyr::arrange(Pathway, Name_to_Plot) %>% 
  dplyr::mutate(Enzyme_Name = str_replace_all(Enzyme_Name, " ", "\n"),
                Enzyme_Name = fct_inorder(Enzyme_Name),
                Name_to_Plot = fct_inorder(Name_to_Plot)) %>% 
  dplyr::filter(Pathway %in% c("Cys/GSH Transport\n& Redox\nHomeostasis",
                               "Cys\nTran-\nsport",
                               "Cystine\nTransport")) %>% 
  ggplot() +
  geom_raster(aes(x = Name_to_Plot, y = Genome, fill = Gene_Pres)) +
  scale_fill_manual(values = c("black", "yellow")) +
  labs(x = "Gene", fill = "Gene\nPresence") +
  theme(axis.text.y = element_blank(),
        axis.ticks.y = element_blank(),
        axis.text.x = element_text(size = 9, angle = 45, hjust = 1),
        strip.text.x = element_text(size = 10),
        # legend.title = element_text(hjust = 0.5, size = 12, face = "bold"),
        legend.title = element_blank(),
        legend.text = element_text(size = 10, hjust = 0.5),
        legend.position = "bottom",
        legend.background = element_rect(colour = 'black', fill = 'white', linetype='solid'),
        axis.title = element_text(size = 12, face = "bold"),
        strip.text.y.right = element_text(angle = 0, face = "italic", size = 10)) +
  guides(fill = guide_legend(title.position="top", title.hjust = 0.5)) +
  facet_grid(Species_abbrev_count ~ Pathway, scales = "free", space = "free_x")

p.cys.cystine.cydABCD.transport.all.gene.pres.rast.Spec.fig

#GSH and BCAA transport
p.BCAA.GSH.transport.all.gene.pres.rast.Spec.fig <- cys.transport.gene.pres.Spec %>% 
  dplyr::mutate(Pathway = case_when(Pathway %in% c("Glutathione and cysteine transport", "Cytochrome bd synthesis") ~
                                      "Cys/GSH Transport\n& Redox\nHomeostasis",
                                    Pathway == "Branched Chain aa Transport" ~ 
                                      "Branched\nChain aa\nTransport",
                                    TRUE ~ str_replace_all(Pathway, " ", "\n")),
                Pathway = str_remove(Pathway, "teine"),
                Pathway = factor(Pathway, 
                                 levels = c("Cys/GSH Transport\n& Redox\nHomeostasis",
                                            "Cys\nTransport",
                                            "Cystine\nTransport",
                                            "Branched\nChain aa\nTransport",
                                            "Glutathione\nTransport")),
                Species_abbrev = str_replace(Species, "actobacillus_", ". "),
                Species_abbrev_count = paste0(Species_abbrev,
                                              "\n(n=",
                                              Genome_Count,
                                              ")"),
                Species_abbrev_count = factor(Species_abbrev_count,
                                              levels = c("L. crispatus\n(n=508)",
                                                         "L. iners\n(n=327)",
                                                         "L. jensenii\n(n=137)",
                                                         "L. gasseri\n(n=216)",
                                                         "L. vaginalis\n(n=30)")),
                Gene_Pres = paste0("Gene\n", Gene_Pres)) %>% 
  dplyr::arrange(Pathway, Name_to_Plot) %>% 
  dplyr::mutate(Enzyme_Name = str_replace_all(Enzyme_Name, " ", "\n"),
                Enzyme_Name = fct_inorder(Enzyme_Name),
                Name_to_Plot = fct_inorder(Name_to_Plot)) %>% 
  dplyr::filter(Pathway %in% c("Branched\nChain aa\nTransport",
                               "Glutathione\nTransport")) %>% 
  ggplot() +
  geom_raster(aes(x = Name_to_Plot, y = Genome, fill = Gene_Pres)) +
  scale_fill_manual(values = c("black", "yellow")) +
  labs(x = "Gene", fill = "Gene\nPresence") +
  theme(axis.text.y = element_blank(),
        axis.ticks.y = element_blank(),
        axis.text.x = element_text(size = 9, angle = 45, hjust = 1),
        strip.text.x = element_text(size = 10),
        # legend.title = element_text(hjust = 0.5, size = 12, face = "bold"),
        legend.title = element_blank(),
        legend.text = element_text(size = 10, hjust = 0.5),
        legend.position = "bottom",
        legend.background = element_rect(colour = 'black', fill = 'white', linetype='solid'),
        axis.title = element_text(size = 12, face = "bold"),
        strip.text.y.right = element_text(angle = 0, face = "italic", size = 10)) +
  guides(fill = guide_legend(title.position="top", title.hjust = 0.5)) +
  facet_grid(Species_abbrev_count ~ Pathway, scales = "free", space = "free_x")

p.BCAA.GSH.transport.all.gene.pres.rast.Spec.fig
```
  
```{r count-gene-numbers-per-genome-for-cys-GSH-transport, echo=FALSE}
#Make dataframe of selected cysteine/cystine transport and GSH transport genes and 
#count number of  gene function of interest in each genome (as opposed to presence-absence)
#Also rename fliY to tcyA for consistency with literature cited in paper.
cys.transport.gene.count.Spec <- genes_queries_seq_pres_abs_all %>% 
  dplyr::filter(Pathway %in% c("Cysteine Transport",
                               "Glutathione and cysteine transport", 
                               "Cystine Transport", 
                               "Branched Chain aa Transport",
                               "Glutathione Transport", 
                               "Cytochrome bd synthesis")) %>% 
  dplyr::group_by(Unique_gene_num, Pathway, Name_to_Plot, Enzyme_Name, Genome, Species) %>% 
  dplyr::summarise(Gene_count = sum(Gene_Pres) %>% 
                     as.character()) %>% 
  dplyr::ungroup() %>% 
  dplyr::arrange(Unique_gene_num) %>% 
  dplyr::group_by(Unique_gene_num, Pathway, Name_to_Plot, Enzyme_Name, Species) %>% 
  dplyr::mutate(Genome_Count = n()) %>% 
  dplyr::ungroup() %>% 
  dplyr::mutate(Name_to_Plot = case_when(Name_to_Plot == "fliY" ~ "tcyA",
                                         TRUE ~ Name_to_Plot))

#Plot heat map of above df with fill as number of genes per genome
p.cys.transport.gene.count.rast.Spec <- cys.transport.gene.count.Spec %>% 
  dplyr::mutate(Pathway = case_when(Pathway %in% c("Glutathione and cysteine transport", "Cytochrome bd synthesis") ~
                                      "Cys/GSH Transport\n& Redox\nHomeostasis",
                                    Pathway == "Branched Chain aa Transport" ~ 
                                      "Branched\nChain aa\nTransport",
                                    TRUE ~ str_replace_all(Pathway, " ", "\n")),
                Pathway = str_remove(Pathway, "teine"),
                Pathway = factor(Pathway, 
                                 levels = c("Cys\nTransport",
                                            "Cystine\nTransport",
                                            "Cys/GSH Transport\n& Redox\nHomeostasis",
                                            "Branched\nChain aa\nTransport",
                                            "Glutathione\nTransport")),
                Species_abbrev = str_replace(Species, "actobacillus_", ". "),
                Species_abbrev_count = paste0(Species_abbrev,
                                              "\n(n=",
                                              Genome_Count,
                                              ")"),
                Species_abbrev_count = factor(Species_abbrev_count,
                                              levels = c("L. crispatus\n(n=508)",
                                                         "L. iners\n(n=327)",
                                                         "L. jensenii\n(n=137)",
                                                         "L. gasseri\n(n=216)",
                                                         "L. vaginalis\n(n=30)"))) %>% 
  dplyr::arrange(Pathway, Name_to_Plot) %>% 
  dplyr::mutate(Enzyme_Name = str_replace_all(Enzyme_Name, " ", "\n"),
                Enzyme_Name = fct_inorder(Enzyme_Name),
                Name_to_Plot = fct_inorder(Name_to_Plot)) %>% 
  ggplot() +
  geom_raster(aes(x = Name_to_Plot, y = Genome, fill = Gene_count)) +
  labs(x = "Gene", fill = "Gene\nCount") +
  scale_fill_manual(values = c("black", "blue", "yellow", "red", "lightblue", "green")) +
  theme(axis.text.y = element_blank(),
        axis.ticks.y = element_blank(),
        axis.text.x = element_text(size = 12, angle = 45, hjust = 1),
        strip.text.x = element_text(size = 11, face = "bold"),
        legend.title = element_text(hjust = 0.5, size = 12, face = "bold"),
        legend.text = element_text(size = 12),
        axis.title = element_text(size = 12, face = "bold"),
        strip.text.y.right = element_text(angle = 0, face = "italic", size = 11)) +
  facet_grid(Species_abbrev_count ~ Pathway, scales = "free", space = "free_x")

p.cys.transport.gene.count.rast.Spec

#Make stacked percent bar chart of number of copies of each gene of interest in each
#genome per gene-species combination.
p.cys.transport.gene.count.bar.Spec <- cys.transport.gene.count.Spec %>% 
  dplyr::group_by(Name_to_Plot, Species, Gene_count) %>% 
  dplyr::mutate(Gene_Count_tally = n()) %>% 
  dplyr::ungroup() %>% 
  dplyr::select(-Genome) %>% 
  dplyr::distinct() %>% 
  dplyr::mutate(Pathway = case_when(Pathway %in% c("Glutathione and cysteine transport", "Cytochrome bd synthesis") ~
                                      "Cys/GSH Transport\n& Redox\nHomeostasis",
                                    Pathway == "Branched Chain aa Transport" ~ 
                                      "Branched\nChain aa\nTransport",
                                    TRUE ~ str_replace_all(Pathway, " ", "\n")),
                Pathway = str_remove(Pathway, "teine"),
                Pathway = factor(Pathway, 
                                 levels = c("Cys\nTransport",
                                            "Cystine\nTransport",
                                            "Cys/GSH Transport\n& Redox\nHomeostasis",
                                            "Branched\nChain aa\nTransport",
                                            "Glutathione\nTransport")),
                Species_abbrev = str_replace(Species, "actobacillus_", ". "),
                Species_abbrev_count = paste0(Species_abbrev,
                                              "\n(n=",
                                              Genome_Count,
                                              ")"),
                Species_abbrev_count = factor(Species_abbrev_count,
                                              levels = c("L. crispatus\n(n=508)",
                                                         "L. iners\n(n=327)",
                                                         "L. jensenii\n(n=137)",
                                                         "L. gasseri\n(n=216)",
                                                         "L. vaginalis\n(n=30)"))) %>% 
  dplyr::arrange(Pathway, Name_to_Plot) %>% 
  dplyr::mutate(Enzyme_Name = str_replace_all(Enzyme_Name, " ", "\n"),
                Enzyme_Name = fct_inorder(Enzyme_Name),
                Name_to_Plot = fct_inorder(Name_to_Plot)) %>% 
  ggplot() +
  geom_col(aes(x = Name_to_Plot, y = Gene_Count_tally, fill = Gene_count),
           position = position_fill()) +
  scale_y_continuous(labels = scales::percent_format(),
                     breaks = c(0, 0.5, 1)) + 
  scale_fill_manual(values = c("black", "blue", "yellow", "red", "lightblue", "green")) +
  labs(x = "Gene", fill = "Gene\nCount") +
  theme(axis.text.x = element_text(size = 12, angle = 45, hjust = 1),
        strip.text.x = element_text(size = 11, face = "bold"),
        legend.title = element_text(hjust = 0.5, size = 12, face = "bold"),
        legend.text = element_text(size = 12),
        axis.title = element_text(size = 12, face = "bold"),
        strip.text.y.right = element_text(angle = 0, face = "italic", size = 11)) +
  facet_grid(Species_abbrev_count ~ Pathway, scales = "free", space = "free_x")

p.cys.transport.gene.count.bar.Spec
```
### Save cys-GSH transport heatmaps

```{r save-cys-GSH-transport-heatmaps-PDF, echo=FALSE}
##Save to PDF##

###File name to save
f.pdf_p.cys.transport.all.gene.pres.rast.Spec.fig<- "Cys_GSH_transport_gene_heatmap_fig_v4_no_singletons_all.pdf"
f.pdf_p.cys.transport.all.gene.pres.rast.Spec.fig

### Write to file
cowplot::ggsave2(filename = f.pdf_p.cys.transport.all.gene.pres.rast.Spec.fig,
                 plot = p.cys.transport.all.gene.pres.rast.Spec.fig,
                 path = path_output,
                 width = 7.75,
                 height = 3.8)

###File names to save
f.pdf_p.cydABCD.gene.pres.rast.Spec.fig <- "cydABCD_gene_heatmap_fig_v3_no_singletons.pdf"
f.pdf_p.cydABCD.gene.pres.rast.Spec.fig

### Write to file
cowplot::ggsave2(filename = f.pdf_p.cydABCD.gene.pres.rast.Spec.fig,
                 plot = p.cydABCD.gene.pres.rast.Spec.fig,
                 path = path_output,
                 width = 3.75,
                 height = 3.8)

###File name to save
f.pdf_p.cystine.GSH.transport.no.cydABCD.gene.pres.rast.Spec.fig <- "Cys_GSH_transport_gene_heatmap_fig_v4_no_singletons_no_cydABCD.pdf"
f.pdf_p.cystine.GSH.transport.no.cydABCD.gene.pres.rast.Spec.fig

### Write to file
cowplot::ggsave2(filename = f.pdf_p.cystine.GSH.transport.no.cydABCD.gene.pres.rast.Spec.fig,
                 plot = p.cystine.GSH.transport.no.cydABCD.gene.pres.rast.Spec.fig,
                 path = path_output,
                 width = 6,
                 height = 3.8)

###File name to save
f.pdf_p.cys.cystine.cydABCD.transport.all.gene.pres.rast.Spec.fig <- "Cys_cystine_cydABCD_transport_gene_heatmap_fig_v2_no_singletons.pdf"
f.pdf_p.cys.cystine.cydABCD.transport.all.gene.pres.rast.Spec.fig

### Write to file
cowplot::ggsave2(filename = f.pdf_p.cys.cystine.cydABCD.transport.all.gene.pres.rast.Spec.fig,
                 plot = p.cys.cystine.cydABCD.transport.all.gene.pres.rast.Spec.fig,
                 path = path_output,
                 width = 7,
                 height = 3.8)

###File name to save
f.pdf_p.BCAA.GSH.transport.all.gene.pres.rast.Spec.fig <- "BCAA_GSH_transport_gene_heatmap_fig_v2_no_singletons.pdf"
f.pdf_p.BCAA.GSH.transport.all.gene.pres.rast.Spec.fig

### Write to file
cowplot::ggsave2(filename = f.pdf_p.BCAA.GSH.transport.all.gene.pres.rast.Spec.fig,
                 plot = p.BCAA.GSH.transport.all.gene.pres.rast.Spec.fig,
                 path = path_output,
                 width = 7,
                 height = 3.8)
```
  
```{r save-cys-GSH-transport-heatmaps-RDS, echo=FALSE}
##Save to RDS##

###File name to save
f.RDS_p.cys.transport.all.gene.pres.rast.Spec.fig<- "Cys_GSH_transport_gene_heatmap_fig_v5_no_singletons_all.RDS"
f.RDS_p.cys.transport.all.gene.pres.rast.Spec.fig

### Write to file
readr::write_rds(p.cys.transport.all.gene.pres.rast.Spec.fig,
                 file.path(path_output,
                           f.RDS_p.cys.transport.all.gene.pres.rast.Spec.fig))

###File names to save
f.RDS_p.cydABCD.gene.pres.rast.Spec.fig <- "cydABCD_gene_heatmap_fig_v3_no_singletons.RDS"
f.RDS_p.cydABCD.gene.pres.rast.Spec.fig

### Write to file
readr::write_rds(p.cydABCD.gene.pres.rast.Spec.fig,
                 file.path(path_output,
                           f.RDS_p.cydABCD.gene.pres.rast.Spec.fig))

###File name to save
f.RDS_p.cystine.GSH.transport.no.cydABCD.gene.pres.rast.Spec.fig <- "Cys_GSH_transport_gene_heatmap_fig_v5_no_singletons_no_cydABCD.RDS"
f.RDS_p.cystine.GSH.transport.no.cydABCD.gene.pres.rast.Spec.fig

### Write to file
readr::write_rds(p.cystine.GSH.transport.no.cydABCD.gene.pres.rast.Spec.fig,
                 file.path(path_output,
                           f.RDS_p.cystine.GSH.transport.no.cydABCD.gene.pres.rast.Spec.fig))

###File name to save
f.RDS_p.cys.cystine.cydABCD.transport.all.gene.pres.rast.Spec.fig <- "Cys_cystine_cydABCD_transport_gene_heatmap_fig_v2_no_singletons.RDS"
f.RDS_p.cys.cystine.cydABCD.transport.all.gene.pres.rast.Spec.fig

### Write to file
readr::write_rds(p.cys.cystine.cydABCD.transport.all.gene.pres.rast.Spec.fig,
                 file.path(path_output,
                           f.RDS_p.cys.cystine.cydABCD.transport.all.gene.pres.rast.Spec.fig))

###File name to save
f.RDS_p.BCAA.GSH.transport.all.gene.pres.rast.Spec.fig <- "BCAA_GSH_transport_gene_heatmap_fig_v2_no_singletons.RDS"
f.RDS_p.BCAA.GSH.transport.all.gene.pres.rast.Spec.fig

### Write to file
readr::write_rds(p.BCAA.GSH.transport.all.gene.pres.rast.Spec.fig,
                 file.path(path_output,
                           f.RDS_p.BCAA.GSH.transport.all.gene.pres.rast.Spec.fig))
```

  
### Table summarizing gene prevalence by Species (Supplementary table)  
  
  
```{r table-summarizing-gene-prevalence-by-Species, echo=FALSE}
#Create dataframe (table) of the prevalence (per-genome presence absence)
#of each function/gene of interest within each genome
gene_prev_table_supp <- genes_queries_seq_pres_abs_all %>% 
  dplyr::filter(Pathway %in% c("Cysteine synthase", 
                               "Reverse transsulfuration", 
                               "Cysteine Transport",
                               "Branched Chain aa Transport",
                               "Glutathione and cysteine transport", 
                               "Cystine Transport", 
                               "Glutathione Transport", 
                               "Cytochrome bd synthesis"),
                !(Enzyme_Name %in% c("membrane alanyl aminopeptidase", "leucyl aminopeptidase")),
                !(Enzyme_Name %in% c("adenosylhomocysteinase", "DNA (cytosine-5-)-methyltransferase",
                                     "methionine adenosyltransferase", "S-ribosylhomocysteine lyase",
                                     "adenosylhomocysteine nucleosidase"))) %>% 
  dplyr::group_by(Unique_gene_num, Pathway, Name_to_Plot, Enzyme_Name, Genome, Species) %>% 
  dplyr::summarise(Gene_Pres = sum(Gene_Pres) >= 1) %>% 
  dplyr::ungroup() %>% 
  dplyr::group_by(Unique_gene_num, Pathway, Name_to_Plot, Enzyme_Name, Species) %>% 
  dplyr::summarise(Gene_Prevalence = sum(Gene_Pres) / n(),
                   Gene_Prevalence = signif(Gene_Prevalence * 100, digits = 2) %>% 
                     paste0("%"),
                   Species_abbrev_count = paste0(str_replace(Species, "actobacillus_", ". "), 
                                        " (n=", n(), ")"),
                   Species_abbrev_count = factor(Species_abbrev_count,
                                                 levels = c("L. crispatus (n=508)",
                                                            "L. iners (n=327)",
                                                            "L. jensenii (n=137)",
                                                            "L. gasseri (n=216)",
                                                            "L. vaginalis (n=30)"))) %>% 
  dplyr::ungroup() %>% 
  dplyr::arrange(Species_abbrev_count, Unique_gene_num) %>% 
  dplyr::select(-c(Species)) %>% 
  dplyr::distinct() %>% 
  tidyr::pivot_wider(names_from = Species_abbrev_count,
                     values_from = Gene_Prevalence)  %>% 
  dplyr::mutate(Name_to_Plot = case_when(Name_to_Plot == "fliY" ~ "tcyA",
                                         TRUE ~ Name_to_Plot))

#Review table
gene_prev_table_supp %>% 
  dplyr::rename(`Unique Gene Number` = Unique_gene_num,
                `Name to Plot` = Name_to_Plot,
                `Enzyme Name` = Enzyme_Name) %>% 
  knitr::kable(caption = "Gene Function Prevalence (Presence-Absence per Genome) by Species")
``` 
  
```{r save-species-gene-prevalence-to-file, echo=FALSE}
f_gene_prev_table_supp <- "Gene_Prevalence_per_Species_supp_v5_no_singletons.txt"
f_gene_prev_table_supp

readr::write_tsv(gene_prev_table_supp,
                 file.path(path_output, f_gene_prev_table_supp))
```


## Save relevant dfs and figures for paper as .RData file  
  
  
```{r save-data-to-RData-file, echo=FALSE}
f_L_iners_genome_files <- "L_iners_genome_analysis_files3.RData"
f_L_iners_genome_files

save(genome_gene_cts_SGB_species.no.singletons,
     genome_reference,
     pan.genome.expansion.annot.no.singletons,
     cys.synth.gene.pres.Spec,
     cys.transport.gene.pres.Spec,
     gene_prev_table_supp,
     file = file.path(path_output, f_L_iners_genome_files))
```

# BLAST sanity check  

## Export multi-fasta of all protein sequences for BLAST analysis by performing blastp of known protein sequences of interest against the pan-genome genes as targets  
  
Translate fasta-format DNA sequences for genes of interest into AA sequences, then unlist each gene into a single string and combine them into a dataframes.  
  
  
```{r translate-full-pangenome, echo=FALSE}
#Translate gene_query_fasta_DNA_list_sep to AA
pangenome_fasta_AA_list_sep <- pangenome_fasta_DNA_list_sep %>% 
  purrr::map(translate, numcode = 11)

#Assign attributes to pangenome_fasta_AA_list_sep for each sequence,
#using the name and Annot from gene_query_fasta_DNA_list_sep
#and class SeqFastaAA
for (n in names(pangenome_fasta_AA_list_sep)) {
  attributes(pangenome_fasta_AA_list_sep[[n]]) <- attributes(pangenome_fasta_DNA_list_sep[[n]])
  attributes(pangenome_fasta_AA_list_sep[[n]])$class <- "SeqFastaAA"
  }

#Remove looping variable
rm(n)

#For each gene sequence, concatenates the vector containing each of its letters individually
#into a single string DNA
pangenome_fasta_AA_list <- pangenome_fasta_AA_list_sep %>% 
  purrr::map(seqinr::c2s)

#Assign attributes to pangenome_fasta_AA_list for each sequence,
#using the name and Annot from pangenome_fasta_AA_list_sep
#and class SeqFastaAA
for (n in names(pangenome_fasta_AA_list)) {
  attributes(pangenome_fasta_AA_list[[n]]) <- attributes(pangenome_fasta_AA_list_sep[[n]])
  }
#Remove looping variable
rm(n)

#Convert pangenome_fasta_AA_list to dataframe
pangenome_AA_df <- purrr::map_df(pangenome_fasta_AA_list, unlist_AA_fasta_list) 

pangenome_AA_fasta_long <- pangenome_AA_df %>% 
  tidyr::pivot_longer(cols = everything(),
                      names_to = "Labels",
                      values_to = "Fasta") %>% 
  dplyr::select(Fasta)
```
  
  
```{r save-all-pangenome-translated-fasta, echo=FALSE}
f_pangenome_AA_fasta_long <- "Lacto_Genome_Data_2020_07_21/pan_genome_reference_2020_07_21_protein_fasta.txt"
f_pangenome_AA_fasta_long

readr::write_tsv(pangenome_AA_fasta_long,
                 f_pangenome_AA_fasta_long,
                 col_names = FALSE)
```
  
  
# Summary of BLAST query approach  
  
  
*  For blastp analysis of the above whole-pangenome multi-fasta, names and sequences were edited to remove spaces
*  A collection of query sequences for genes of interest from annotated genomes was put together, with each gene of interest represented by one or more query sequences, named (for example) CjaA.faa used to search the pan-genome.
*  BLAST was loaded onto server with command:
    +  module load engaging/ncbi-blast/2.2.30+
*  Made a BLAST database using fasta file “pan_genome” with protein sequences, using command:
    +  makeblastdb -in pan_genome.faa -out pan_genome_prot.blastdb -dbtype prot
*  Blastp performed by searching database for query proteins using blastp with commands as follows (example shown for gene CjaA):
    +  blastp -query CjaA.faa -db pan_genome_prot.blastdb -out CjaA.blastp.out -evalue 1e-8 -outfmt 6
*  The output of the blast search for each gene of interest is (for example CjaA) “CjaA.blastp.out”  
  
  
## Read in BLAST results  
  
  
```{r directory-path-to-BLAST-results, echo=FALSE}
path_blast_output <- "blast_output"
```
  
Path to subdirectory with BLAST output: `r path_blast_output`  
  
  
```{r read-in-BLAST-results, echo=FALSE}
#obtain file names of BLAST results
blastp_output <- list.files(path_blast_output) %>% 
  stringr::str_subset("blastp\\.out$")

#Review
blastp_output %>% 
  knitr::kable(col.names = "blastp output files")

#Read in list of tables with results of blastp output
blastp_list <- blastp_output %>% 
  purrr::map(~ read_tsv(file.path(path_blast_output,
                                  .x),
                        col_names = c("qseqid",
                                      "sseqid",
                                      "pident",
                                      "length",
                                      "mismatch",
                                      "gapopen",
                                      "qstart",
                                      "qend",
                                      "sstart",
                                      "send",
                                      "evalue",
                                      "bitscore")
                        ))

#Name the elements of the list according to the gene of interest
names(blastp_list) <- blastp_output %>% 
  str_remove("\\.blastp\\.out$")

#Convert blastp list to unfiltered dataframe with column for gene of interest
blastp_df_all <- blastp_list %>% 
  dplyr::bind_rows(.id = "Gene_of_Interest")
  
#Correct annotation of CydC and CydD blast query genes from B. subtilis 
#(they were mis-annotated in the source genome)
blastp_df_all <- blastp_df_all %>% 
  dplyr::mutate(Gene_of_Interest = case_when(str_detect(qseqid, "BSU_38730") ~ "CydC",
                                             str_detect(qseqid, "BSU_38740") ~ "CydD",
                                             TRUE ~ Gene_of_Interest))
```
    
In the sequences used as queries for the blastp searches, _Bacillus subtilis_ further review has indicated that the annotations of the CydD and CydC genes in that particular source genome were switched. So BSU_38730 was annotated as CydD but should be CydC and BSU_38740 was annotated as CydC but should be CydD. Manually switching these here.  
  
    
## Basic review of blastp results  
  
There are `r nrow(blastp_df_all)` total targets identified by the blastp analysis, including duplicates.  
  
There are `r blastp_df_all %>% dplyr::distinct(sseqid) %>% nrow` unique target sequences identified by blastp analysis across all query sequences. Some may have been identified by multiple queries for multiple genes of interest.  
  
  
Sanity check: How many target genes identified in BLAST output are NOT present in the overall gene-presence-absence catalog (should be 0)?    
  
  
```{r blast-targets-absent-from-pan-genome, echo=FALSE}
blastp_df_all %>% 
  dplyr::anti_join(gene_pres_abs_Spec_SGB %>% 
                     distinct(Gene_Group),
                   by = c("sseqid" = "Gene_Group")) %>% 
  nrow()
```
  
  
How many of the target genes identified in BLAST output are absent from eggNOG annotations (not annotated by eggNOG)?  
  

```{r blast-targets-absent-from-eggNOG-annotations, echo=FALSE}
blastp_df_all %>% 
  dplyr::anti_join(EggNOG_annotations_raw,
                   by = c("sseqid" = "Query")) %>% 
  nrow()
```
 


How many of the target genes identified in BLAST output are retained after filtering out per-species singletons from the pan-genome?  
  
```{r create-df-of-pangenome-genes-without-singletons}
#Eliminate per-species singletons, then deduplicate to unique genes in pan-genome
pangenome_no_singletons <- gene_pres_abs_Spec_SGB %>% 
  dplyr::filter(Num_Genomes_Present == "Gene in Multiple Genomes") %>% 
  dplyr::filter(Gene_Pres == 1) %>% 
  dplyr::distinct(Gene_Group)
```
  
  
```{r review-BLAST-hits-by-singleton-or-not, echo=FALSE}
blastp_df_all %>% 
  dplyr::distinct(sseqid) %>% 
  dplyr::mutate(`Singleton Gene` = !(sseqid %in% pangenome_no_singletons$Gene_Group)) %>% 
  dplyr::count(`Singleton Gene`,
               name = "Number of unique hits")
```


  
How many of the target genes identified in BLAST output are absent from eggNOG annotations (not annotated by eggNOG) after filtering out singletons?  
  

```{r blast-targets-absent-from-eggNOG-annotations-no-singletons, echo=FALSE}
blastp_df_all %>% 
  dplyr::filter(sseqid %in% pangenome_no_singletons$Gene_Group) %>% 
  dplyr::anti_join(EggNOG_annotations_raw,
                   by = c("sseqid" = "Query")) %>% 
  nrow()
```
  
Thus, after excluding hits to singleton genes within the pan-genome (which we are excluding from final analysis as these are either extremely rare or -- more likely -- contaminants in the assemblies), all genes found by our blastp approach also have eggNOG annotations.  
  
  
## Review overall quality of blastp hits in pangenome  
  
  
```{r plot-pident-against-evalue-all, echo=FALSE}
#For all blastp hits, plot percent identity against evalue
p_blast_pident_vs_evalue <- blastp_df_all %>% 
  ggplot(aes(x = pident, y = evalue)) +
  geom_point() +
  scale_y_log10()  

#Marginal histogram
p_blast_pident_vs_evalue.hist <- p_blast_pident_vs_evalue %>% 
  ggMarginal(type = "histogram")

p_blast_pident_vs_evalue.hist
```
  
  
  
  
## Review blastp agreement with eggNOG annotations (looking for potential false-positives from eggNOG)  
  
  
Approach:  
  
For each gene identified by eggNOG as a gene of interest:  
  
*  Does it have any matches to any of the blastp queries?
    +  If no, further sanity check by blasting it against NCBI nr db
*  If yes, do any of the blastp queries matching it have the same annotation as eggNOG?
    +  If no, see what matches it and further investigate including possibly blasting against NCBI
*  If yes, does the strongest match by evalue and/or pident have the same annotation?
    +  If yes, there are no issues.
    +  If no, further investigate.  
  
  
  
```{r eggNOG-hits-with-no-blastp-matches, echo=FALSE}
genes_queries_seq_pres_abs_all %>% 
  dplyr::filter(Gene_Pres == 1) %>% 
  dplyr::select(Species, Name_to_Plot, Query, `EC number`, `KEGG KO`, `Seed ortholog`) %>%
  dplyr::distinct() %>% 
  dplyr::mutate(Name_to_Plot = str_replace(Name_to_Plot, "fliY", "tcyA"),
                Species = str_replace_all(Species, "_", " ")) %>% 
  dplyr::filter(toupper(Name_to_Plot) %in% toupper(blastp_df_all$Gene_of_Interest),
                !is.na(Query)) %>% 
  dplyr::anti_join(blastp_df_all %>%
                     dplyr::filter(sseqid %in% 
                                     pangenome_no_singletons$Gene_Group),
                   by = c("Query" = "sseqid")) %>% 
  dplyr::arrange(Name_to_Plot, Species) %>% 
  knitr::kable(caption = "Genes from pan-genome (no singletons) identified by eggNOG that were NOT identified by BLAST analysis")
```
  
  
*  One of the genes is cysE from _L. crispatus_. Blastp analysis of this gene against the NCBI nr database is consistent with it being a serine O-acetyltransferase (cysE) as annotated. Thus does not appear to be a "false positive". Notably, this was the sequence found in isolates from poultry in a single study (see above).  
  
*  The remaining eggNOG hits not identified by BLAST are all annotated as mccB (cystathionine gamma-lyase) based on the eggNOG annotations. Blastp analysis of this gene against the nr database is consistent with it being in the methionine gamma-lyase protein family. So these could theoretically be false-positive IDs as mccB from eggNOG, but this method is inadequate to be certain. For purposes of this paper's analysis, if they were indeed false positives, it would only further strengthen our conclusion that these species lack and intact reverse transsulfuration pathway.  
   
  
  
```{r df-of-genes-found-by-both-eggNOG-and-blast, echo=FALSE}
#Create df of genes identified by eggNOG also matched by one or more blast hit
#Filter eggNOG "hits" to detected genes,
#deduplicate to sequence/Species level,
#format Name_to_Plot,
#Filter to genes portrayed in paper
#Join to blastp results by names of unique gene sequences 
#(single sequence may have been hit by multiple blastp queries)


eggNOG_blastp_common_hits <- genes_queries_seq_pres_abs_all %>% 
  dplyr::filter(Gene_Pres == 1) %>% 
  dplyr::distinct(Pathway, Name_to_Plot, Query, 
                  Species, `Seed ortholog`, AA_Seq) %>%
  dplyr::mutate(Name_to_Plot = str_replace(Name_to_Plot, "fliY", "tcyA"),
                Name_to_Plot = toupper(Name_to_Plot)) %>% 
  dplyr::filter(Name_to_Plot %in% toupper(blastp_df_all$Gene_of_Interest),
                Pathway != "Homocysteine desulfhydrase"
                ) %>% 
  dplyr::inner_join(blastp_df_all,
                   by = c("Query" = "sseqid")) %>% 
  dplyr::mutate(Gene_of_Interest = toupper(Gene_of_Interest)) %>% 
  dplyr::select(Species, Query, Name_to_Plot, Gene_of_Interest, evalue, pident, 
                `Seed ortholog`, qseqid, everything()) %>% 
  dplyr::arrange(Query, evalue)
```


```{r eggNOG-hits-with-complete-blast-agreement, echo=FALSE}
#Return sequences identified by eggNOG for which all blastp hits were for the
#same gene:
eggNOG_blastp_common_hits%>% 
  dplyr::group_by(Query) %>% 
  dplyr::filter(all(Name_to_Plot == Gene_of_Interest)) %>% 
  dplyr::ungroup() %>% 
  dplyr::arrange(Name_to_Plot) %>% 
  dplyr::group_by(`eggNOG annotation` = Name_to_Plot,
                  `blastp hit annotation` = Gene_of_Interest,
                  Query, `Seed ortholog`) %>% 
  dplyr::summarise(`Strongest blastp evalue` = min(evalue),
                   `Highest blastp percent identity` = max(pident)) %>% 
  dplyr::ungroup() %>% 
  knitr::kable(caption = "eggNOG-annotated genes for which annotations of ALL blastp hits match")
```
  
  
These results are strongly consistent with these gene annotations being "true positive" identifications by eggNOG and require no further review.  
  

```{r df-of-genes-with-complete-eggNOG-blast-disagreement, echo=FALSE}
#Return sequences identified by eggNOG for which none of the blastp hits were 
#for the same gene:
eggNOG_blastp_common_hits%>% 
  dplyr::group_by(Query) %>% 
  dplyr::filter(all(Name_to_Plot != Gene_of_Interest)) %>% 
  dplyr::ungroup() %>% 
  dplyr::arrange(Name_to_Plot) %>% 
  dplyr::select(`eggNOG annotation` = Name_to_Plot,
                `blastp hit annotation` = Gene_of_Interest,
                Query, `Seed ortholog`,
                `blastp evalue` = evalue,
                `blastp percent identity` = pident) %>% 
  knitr::kable(caption = "eggNOG-annotated genes for which there are NO blastp hits with matching annotations")
```
  
  
There is only one, where the methods disagree whether it is tcyK or tcyJ. As further addressed below, I favor it being tcyK as annotated by eggNOG (i.e., an eggNOG "true positive"), but tcyJ and tcyK likely serve similar functions when both are present in other species, so the distinction of what to call this gene may be largely academic.  
  
  

```{r genes-annotated-differently-by-eggNOG-and-blast, echo=FALSE}
# #For each gene sequence from pangenome, determine if best blastp hit 
# #(lowest evalue) is from a blast query sequence for the gene of interest
#focusing on eggNOG hits matched by at least one blastp hit
#with a differing annotation
p_blastp_matches_to_eggNOG_hits_point_mismatches <- eggNOG_blastp_common_hits %>% 
  dplyr::distinct(Query, Name_to_Plot, Gene_of_Interest, evalue, pident,
                  qseqid, AA_Seq) %>% 
  dplyr::mutate(`Agrees w blastp`= Name_to_Plot == Gene_of_Interest) %>% 
  dplyr::group_by(Query) %>% 
  dplyr::filter(!all(`Agrees w blastp`)) %>% 
  dplyr::ungroup() %>% 
  ggplot(aes(x = pident, y = evalue, color = `Agrees w blastp`)) +
  geom_point() +
  scale_y_log10() +
  scale_x_continuous(breaks = c(0:4) * 25) +
  theme(legend.position = "bottom",
        strip.text = element_text(size = 7),
        axis.text = element_text(size = 7)) +
  facet_wrap(~paste0(Name_to_Plot, "\n", Query))

p_blastp_matches_to_eggNOG_hits_point_mismatches
```
  
Conclusions from above:  
  
*  The blastp analysis largely agrees with eggNOG for the genes that were identified as being of interest via eggNOG (mostly consistent with being "true positives")
*  The only area of potential disagreement appears to be for tcyJKLMN (analyzed in more detail below)  
  
  
```{r blast-hits-for-eggNOG-annotated-TcyJKLMN, echo=FALSE}
eggNOG_blastp_common_hits %>% 
  dplyr::distinct(Species, Query, Name_to_Plot, Gene_of_Interest, evalue, 
                  pident, `Seed ortholog`,
                  qseqid, AA_Seq) %>% 
  dplyr::mutate(`Agrees\nwith\nblastp`= Name_to_Plot == Gene_of_Interest) %>% 
  dplyr::filter(str_detect(Name_to_Plot, "TCY[JKLMN]")) %>% 
  dplyr::arrange(Name_to_Plot, evalue) %>% 
  dplyr::select(Species, Query,
                `eggNOG Gene Name` = Name_to_Plot,
                `blastp Query Gene Name` = Gene_of_Interest,
                evalue, pident, 
                `eggNOG Seed ortholog` = `Seed ortholog`,
                qseqid) %>% 
  dplyr::mutate(across(.cols = c(Species, qseqid),
                       .fns = ~ str_replace_all(.x, "_", " "))) %>% 
  knitr::kable(caption = "blastp hits to genes annotated by eggNOG as tcyJKLMN")
```
  
The 4 genes annotated as tcyKLMN by eggNOG but with different or nearly different annotations based on the blastp analysis are below.  
  
`>Lactobacillus_vaginalis EGKPCKBJ_00682 TCYK 1e-14 28.24 1423814.HMPREF0549_0322  
MKISKLFTKLSVIGLVSAGSVSSATAVHAATTVNVAAPNGNPAYIVNTDNGPEGYYGELIKKVDQDLPQYKFKTTFTSQNAVFSGLQSGKYDVALNNSWYNKQRFQTYYHTRAVGLDDLRLVYNKSGKKASSLDDVAKKNLKLVPVSTDDARYSVLEDYNKSHSKKVNLKAIGDQSSADALAQVANGKYDVALYPYAAYKQVANSKNGKKLTVSKSIDQRSAYLLLHKDAKNKKLTRALNKEIKKLYDNGYLEKLTKKYLYENTFALPGAKKNYNAEFNK  
  
`>Lactobacillus_vaginalis EGKPCKBJ_00680 TCYL 1e-31 33.00 1423814.HMPREF0549_0320  
MNSQKIIGIIWRGFPNTIELLIFSFIFATLLGLLLTWVGLRKNRLAQGIAVVYLNLVRGTPPLLMLLLAYYGLPVLLKAIGINIDHWTRLMFGILGLSLGWSGYLAEAFRSAYLSVDVGQVEAARSVGMKDRTTFWQIILPQAAMLALPNIENLVVGLVKATSLVYVIGLYDMYNQASNLSNQTAGLHQLQIFILLALSYWGLVLVIEALFCLIRRRYQWLA  
  
`>Lactobacillus_vaginalis EGKPCKBJ_00681 TCYM 4e-34 33.63 1423814.HMPREF0549_0321  
VFDLNFAKDSFNQILAATPLTLIVALVATIIGLGLAILVAIAREKRIPFLSQFLAVLVSFIRGTPILVQLYVVYYGLPQLLVLMNEWGWNTNPNGLPAIIIAFSAYGLNAMANLSESIRAAYHSVDFGQYEGALSIGMTPFRAMTQIVIPQLIINLIPNFTNIFLDLIKDTALIYNIGLVEIMGQANIVSSIGFKYLETYTDALIIYIIICWIFAQLFRLVEGKLRQRYFVV  
  
`>Lactobacillus_vaginalis EGKPCKBJ_00679 TCYN 2e-85 49.59 1423814.HMPREF0549_0319  
MLLKITDLKKQFNGQQILNGIDLSINKGEVVVIIGPSGSGKTTLLRNLNFLEQADSGLMEFSGEKINLAQPTKPEIQMVRRKTAMVFQNYNLFKNKTAQENIAEGLIYGHGENAKTANQKAVQALKNLGLDKRKDYYPSQLSGGQQQRIGIARATVLNPEIILFDEPTSALDPELVGTVLKDIQRLAEQGQTMIVVTHQMWFARNVASRVLFFANGEIVESGTPAQIFDHPQEQRTKQFLNALADNY  
  
  
Review of their synteny and comparison with annotations and synteny in genome of _L. vaginalis_ ( _Limosilactobacillus vaginalis DSM 5837_ ) in IMG/M supports the eggNOG-based annotations over the blastp results. Can't definitively distinguish whether Query EGKPCKBJ_00682 should be annotated as tcyJ vs tcyK, but they are likely functionally redundant. Thus, favor continuing to classify all of these as "true positive" eggNOG results.    
   
  
## Review "hits" for each query gene -- evidence of false negatives from the eggNOG approach?   

Approach:  
  
For each blastp query of interest:  
  
*  Does it hit one or more genes in the pan-genome?
*  Are all of the genes it hits among the "correct" genes of interest identified by eggNOG?
    +  If yes, no further analysis needed (no chance of a false negative)
*  If no, review the quality of its hits in the pan-genome, assess which of those are any of the genes it hits among the "correct" genes of interest identified by eggNOG?
    +  If yes, are the hits to the "correct" genes of interest its strongest hits?
        +  If yes by a substantial margin, very encouraging. (True positive)
            +  As a sanity check, see what eggNOG called the next few strongest hits. (False positive?)
        +  If not, look at evalues and see what eggNOG called the strongest hits. (Do they look like false positives? Why was the "right" answer so weak?)
    +  If not, see what eggNOG called the strongest hits. (False positives?)
  
What is the number of total results for each query gene (function) of interest across all query sequences for that gene function (if there are multiple gene sequence queries, they may each have identified the same target/result gene)?  
  

```{r set-perc-identity-cutoff-to-review-blast-results, echo=FALSE}
blast_pident_cutoff <- 35
blast_evalue_cutoff <- 10^-30
```
  
Percent identity cutoff for some of the blastp result review is `r blast_pident_cutoff`%.  
e-valued cutoff for some of the blastp result review is `r blast_evalue_cutoff`.  


```{r review-table-summary-of-blastp-results, echo=FALSE}
#Nest on gene of interest, calculate summary statistics
blastp_df_all %>% 
  tidyr::nest(results = -Gene_of_Interest) %>% 
  dplyr::mutate(`# Query Sequences` = map_int(results,
                                              ~ .x %>% 
                                                distinct(qseqid) %>% 
                                                nrow()),
                `Total hits identified (may include duplicates)` = map_int(results,
                                                                           ~ .x %>% 
                                                                             nrow()),
                `Number of unique targets identified` = map_int(results,
                                                                ~ .x %>% 
                                                                  distinct(sseqid) %>% 
                                                                  nrow()),
                `Total hits identified (may include duplicates; no singletons)` = map_int(results,
                                                                                          ~ .x %>% 
                                                                                            dplyr::filter(sseqid %in% 
                                                                                                            pangenome_no_singletons$Gene_Group) %>% 
                                                                                            nrow()),
                `Number of unique targets identified (no singletons)` = map_int(results,
                                                                                ~ .x %>% 
                                                                                  dplyr::filter(sseqid %in% 
                                                                                                  pangenome_no_singletons$Gene_Group) %>% 
                                                                                  distinct(sseqid) %>% 
                                                                                  nrow())) %>% 
  dplyr::select(-results) %>%
  dplyr::rename(`blastp Query gene of interest` = Gene_of_Interest) %>% 
  knitr::kable(caption = "Summary of overall blastp results for genes of interest")

#Nest on gene of interest, calculate summary statistics after filtering
blastp_df_all %>% 
  dplyr::filter(pident > blast_pident_cutoff,
                evalue < blast_evalue_cutoff) %>%
  tidyr::nest(results = -Gene_of_Interest) %>% 
  dplyr::mutate(`# Query Sequences` = map_int(results,
                                              ~ .x %>% 
                                                distinct(qseqid) %>% 
                                                nrow()),
                `Total hits identified (may include duplicates)` = map_int(results,
                                                                           ~ .x %>% 
                                                                             nrow()),
                `Number of unique targets identified` = map_int(results,
                                                                ~ .x %>% 
                                                                  distinct(sseqid) %>% 
                                                                  nrow()),
                `Total hits identified (may include duplicates; no singletons)` = map_int(results,
                                                                                          ~ .x %>% 
                                                                                            dplyr::filter(sseqid %in% 
                                                                                                            pangenome_no_singletons$Gene_Group) %>% 
                                                                                            nrow()),
                `Number of unique targets identified (no singletons)` = map_int(results,
                                                                                ~ .x %>% 
                                                                                  dplyr::filter(sseqid %in% 
                                                                                                  pangenome_no_singletons$Gene_Group) %>% 
                                                                                  distinct(sseqid) %>% 
                                                                                  nrow())) %>% 
  dplyr::select(-results) %>%
  dplyr::rename(`blastp Query gene of interest` = Gene_of_Interest) %>% 
  knitr::kable(caption = paste0("Summary of overall blastp results for genes of interest, filtering for blast identity ", blast_pident_cutoff, "% and e-value <", blast_evalue_cutoff))
  
```
  
  
How many target genes within the pangenome have multiply mapped query sequences of interest after removing singletons?  
  

```{r explore-multiple-mapping, echo=FALSE}
#Histogram
blastp_df_all %>% 
  dplyr::filter(sseqid %in% 
                  pangenome_no_singletons$Gene_Group) %>% 
  dplyr::distinct(Gene_of_Interest, sseqid) %>% 
  dplyr::count(sseqid,
               name = "Num_Genes_of_Interest") %>% 
  ggplot(aes(x = Num_Genes_of_Interest)) +
  geom_histogram(binwidth = 1) +
  labs(y = "Number of genes\nwithin pangenome",
       x = "# blastp Query genes of\ninterest that map to gene in pangenome",
       title = "No singletons")

#Filtered Histogram
blastp_df_all %>% 
  dplyr::filter(pident > blast_pident_cutoff,
                evalue < blast_evalue_cutoff) %>% 
  dplyr::filter(sseqid %in% 
                  pangenome_no_singletons$Gene_Group) %>% 
  dplyr::distinct(Gene_of_Interest, sseqid) %>% 
  dplyr::count(sseqid,
               name = "Num_Genes_of_Interest") %>% 
  ggplot(aes(x = Num_Genes_of_Interest)) +
  geom_histogram(binwidth = 1) +
  labs(y = "Number of genes\nwithin pangenome",
       x = "# blastp Query genes of\ninterest that map to gene in pangenome",
       title = paste0("No singletons\nfiltering for blast identity ", blast_pident_cutoff, "% and e-value <", blast_evalue_cutoff))

#Presence-absence heat map
blastp_df_all %>% 
  dplyr::filter(sseqid %in% 
                  pangenome_no_singletons$Gene_Group) %>% 
  dplyr::distinct(Gene_of_Interest, sseqid) %>% 
  dplyr::add_count(sseqid,
               name = "Num_Genes_of_Interest") %>% 
  dplyr::arrange(desc(Num_Genes_of_Interest)) %>% 
  # dplyr::filter(Num_Genes_of_Interest > 1) %>% 
  dplyr::mutate(Present = "Hit",
                sseqid = fct_inorder(sseqid)) %>% 
  ggplot(aes(x = Gene_of_Interest, y = sseqid, fill = Present)) +
  geom_raster() +
  theme(axis.text.y = element_blank(),
        axis.text.x = element_text(angle = 45, hjust = 1)) +
  labs(y = "Gene from pangenome",
       x = "blastp Gene of Interest",
       fill = "blastp\nhit",
       title = "No singletons") +
  facet_grid(str_wrap(paste0(Num_Genes_of_Interest, " genes mapped"), 
                      width = 5) ~ ., 
             scales = "free", space = "free_y")

#Filtered presence-absence heat map
blastp_df_all %>% 
  dplyr::filter(pident > blast_pident_cutoff,
                evalue < blast_evalue_cutoff) %>%
  dplyr::filter(sseqid %in% 
                  pangenome_no_singletons$Gene_Group) %>% 
  dplyr::distinct(Gene_of_Interest, sseqid) %>% 
  dplyr::add_count(sseqid,
               name = "Num_Genes_of_Interest") %>% 
  dplyr::arrange(desc(Num_Genes_of_Interest)) %>% 
  # dplyr::filter(Num_Genes_of_Interest > 1) %>% 
  dplyr::mutate(Present = "Hit",
                sseqid = fct_inorder(sseqid)) %>% 
  ggplot(aes(x = Gene_of_Interest, y = sseqid, fill = Present)) +
  geom_raster() +
  theme(axis.text.y = element_blank(),
        axis.text.x = element_text(angle = 45, hjust = 1)) +
  labs(y = "Gene from pangenome",
       x = "blastp Gene of Interest",
       fill = "blastp\nhit",
       title = paste0("No singletons\nfiltering for blast identity ", 
                      blast_pident_cutoff, "% and e-value <", blast_evalue_cutoff)) +
  facet_grid(str_wrap(paste0(Num_Genes_of_Interest, " genes mapped"), 
                      width = 5) ~ ., 
             scales = "free", space = "free_y")
```
  
  
The multiply mapped gene are enriched for proteins with ATP-binding cassettes as part of ABC transporters, thus the vast majority of these genes are likely not functionally redundant except for the shared domains that mediate this common activity. Will explore this further below.    
  
  


```{r df-of-all-blastp-hits-with-eggNOG-annotatons-over-cutoff, echo=FALSE}
#Filter blast results by pident > blast_pident_cutoff and evalue < blast_evalue_cutoff
blastp_all_eggNOG_hits_filt <- blastp_df_all %>%
  dplyr::filter(pident > blast_pident_cutoff,
                evalue < blast_evalue_cutoff) %>%
  dplyr::mutate(Gene_of_Interest = toupper(Gene_of_Interest)) %>% 
  dplyr::inner_join(EggNOG_annotations_raw_no_singletons,
                    by = c("sseqid" = "Query")) %>% 
  dplyr::left_join(genes_queries_seq_pres_abs_all %>% 
                     dplyr::distinct(Pathway, Name_to_Plot, Query) %>%
                     dplyr::mutate(Name_to_Plot = str_replace(Name_to_Plot, "fliY", "tcyA"),
                                   Name_to_Plot = toupper(Name_to_Plot)) %>% 
                     dplyr::filter(Name_to_Plot %in% toupper(blastp_df_all$Gene_of_Interest),
                                   Pathway != "Homocysteine desulfhydrase",
                                   !is.na(Query)),
                   by = c("sseqid" = "Query")) %>% 
  dplyr::mutate(eggNOG_curated_match = Gene_of_Interest == Name_to_Plot)
```


```{r quality-all-blast-hits, echo=FALSE}
#Overall distribution
blastp_all_eggNOG_hits_filt %>% 
  dplyr::rename(`Matches\nCurated\neggNOG\nannotation` = eggNOG_curated_match) %>% 
  ggplot(aes(x = pident, 
             y = evalue, 
             color = `Matches\nCurated\neggNOG\nannotation`, 
             alpha = `Matches\nCurated\neggNOG\nannotation`)) +
  geom_point() +
  scale_y_log10() +
  labs(title = "All Species",
       x = "blastp % Identity",
       y = "blastp e-value") +
  scale_alpha_manual(values = c(1, 1),
                     na.translate = TRUE,
                     na.value = 0.2)

#Overall distribution faceted
blastp_all_eggNOG_hits_filt %>% 
  dplyr::group_by(qseqid) %>% 
  dplyr::filter(!all(eggNOG_curated_match) | 
                  is.na(all(eggNOG_curated_match))) %>%
  dplyr::ungroup() %>% 
  dplyr::rename(`Matches\nCurated\neggNOG\nannotation` = eggNOG_curated_match) %>% 
  ggplot(aes(x = pident, 
             y = evalue, 
             color = `Matches\nCurated\neggNOG\nannotation`, 
             alpha = `Matches\nCurated\neggNOG\nannotation`)) +
  geom_point() +
  scale_y_log10() +
  labs(title = "All Species",
       x = "blastp % Identity",
       y = "blastp e-value") +
  scale_alpha_manual(values = c(1, 1),
                     na.translate = TRUE,
                     na.value = 0.2) +
  facet_wrap(~paste0(Gene_of_Interest, "\n", qseqid))

```

```{r blast-hits-by-species-summary, echo=FALSE}
#Review per-species, non-singleton blastp hits for all queries by gene of interest and if they
#match the curated eggNOG annotations.
blastp_all_eggNOG_hits_filt %>% 
  dplyr::left_join(gene_pres_Spec_SGB_times_seen %>% 
                     dplyr::filter(Num_Genomes_Present == "Gene in Multiple Genomes") %>% 
                     dplyr::count(Gene_Group, Species, name = "Num_genomes"),
                   by = c("sseqid" = "Gene_Group")) %>% 
  dplyr::count(Species,
               Gene_of_Interest, 
               # qseqid, 
               # Num_genomes,
               eggNOG_curated_match,
               name = "Num_Hits") %>% 
  dplyr::mutate(Species = str_replace_all(Species, "actobacillus_", " ")) %>%
  dplyr::arrange(Species, eggNOG_curated_match) %>% 
  tidyr::pivot_wider(names_from = eggNOG_curated_match,
                     names_prefix = "match ",
                     values_from = Num_Hits,) %>% 
  dplyr::arrange(Species, Gene_of_Interest) %>% 
  knitr::kable(caption = paste0("# of blastp hits >", blast_pident_cutoff, "% identity and e-value <", blast_evalue_cutoff, "to non-singletone genes by species for each gene query of interest and whether they match curated eggNOG annotation"))
```
  
  
Conclusions:  
  
*  There are few genes at a cutoff of identify >`r blast_pident_cutoff`% and e-value <`r blast_evalue_cutoff` for which blastp identified a gene that our curated analysis in eggNOG annotated as something else.
    +  None of these for potential biosynthetic enzymes.
    +  None of these are in _L. iners_
*  For some genes of interest, there are numerous blast hits that were not identified by curated eggNOG annotations.
    +  These will require some further exploration in case they represent "false negatives" based on the eggNOG approach.
    +  The only ones that appear to have potential to fundamentally change conclusions about gene presence in some or all members of a species are:
        +  cydC in _L. iners_
        +  cydC and cydD in _L. jensenii_
        +  gsiA in _L. vaginalis_ and _L. iners_
        +  gsiC and gsiD in _L. crispatus_
        +  tcyB and tcyC in _L. crispatus_, _L. gasseri_, and _L. iners_
        +  tcyJ in _L. crispatus_
        +  tcyL in _L. crispatus_
        +  tcyN in all species (already annotated as present in _L. vaginalis_)
    +  In summary, the only intact transport mechanism that this analysis could possibly add for each species would be:
        +  _L. crispatus_:
            +  gsiCD (no gsiB)
            +  tcyBC (no tcyA)
            +  tcyLN (no tcyJ, tcyK or tcyM)
        +  _L. iners_:
            +  cydC (assuming it were separable from cydAB and cydD)
            +  tcyBC (assuming it were separable from tcyA)
        +  _L. jensenii_:
            +  cjaA
            +  cydCD
        +  _L. gasseri_:
            +  tcyBC (no tcyA)
        +  _L. vaginalis_:
            +  gsiA (but not gsiBCD)
        
  
  

```{r tally-all-blast-hits, echo=FALSE}
blastp_all_eggNOG_hits_filt %>% 
  dplyr::left_join(gene_pres_Spec_SGB_times_seen %>% 
                     dplyr::filter(Num_Genomes_Present == "Gene in Multiple Genomes") %>% 
                     dplyr::count(Gene_Group, Species, name = "Num_genomes"),
                   by = c("sseqid" = "Gene_Group")) %>% 
  dplyr::count(Species,
               Gene_of_Interest, qseqid, 
               # Num_genomes,
               eggNOG_curated_match,
               name = "Num_Hits") %>% 
  dplyr::mutate(qseqid = str_replace_all(qseqid, "_", " "),
                Species = str_replace_all(Species, "actobacillus_", " ")) %>%
  dplyr::arrange(Species, eggNOG_curated_match) %>% 
  tidyr::pivot_wider(names_from = eggNOG_curated_match,
                     names_prefix = "match ",
                     values_from = Num_Hits,) %>% 
  dplyr::arrange(Species, Gene_of_Interest) %>% 
  knitr::kable(caption = "Number of distinct gene sequences within the pan-genome matching each blastp query sequence, including whether they match genes with the same curated eggNOG annotations, match genes with conflicting annotations based on the curated eggNOG analysis, or match genes that weren't picked out by the curated eggNOG approach.")
```
  
  
## Analysis of results for blastp hits to genes from _Lactobacillus crispatus_  
  
  
```{r species-on-which-to-filter-lc, echo=FALSE}
#Specify species to filter on
filt_species <- "Lactobacillus_crispatus"
```
  
  
Filtering results for `r filt_species`.

```{r quality-all-blast-hits-L-crispatus, echo=FALSE}
#Plot quality of blastp hits by query gene
p_L_crisp_blastp_hit_quality <- blastp_all_eggNOG_hits_filt %>% 
  dplyr::left_join(gene_pres_Spec_SGB_times_seen %>% 
                     dplyr::filter(Num_Genomes_Present == "Gene in Multiple Genomes") %>% 
                     dplyr::count(Gene_Group, Species, name = "Num_genomes"),
                   by = c("sseqid" = "Gene_Group")) %>% 
  dplyr::filter(Species == filt_species) %>% 
  # dplyr::group_by(qseqid) %>% 
  # dplyr::filter(!all(eggNOG_curated_match) |
  #                 is.na(all(eggNOG_curated_match))) %>% 
  # dplyr::ungroup() %>% 
  dplyr::rename(`blastp % Identity` = pident,
                `blastp e-value` = evalue,
                `Matches\nCurated\neggNOG\nannotation` = eggNOG_curated_match,
                `#\nGenomes` = Num_genomes) %>% 
  ggplot(aes(x = `blastp % Identity`, 
             y = `blastp e-value`, 
             color = `Matches\nCurated\neggNOG\nannotation`, 
             alpha = `Matches\nCurated\neggNOG\nannotation`, 
             size = `#\nGenomes`)) +
  geom_point() +
  scale_y_log10() +
  scale_alpha_manual(values = c(1, 1),
                     na.translate = TRUE,
                     na.value = 0.3) +
  labs(title = filt_species) +
  facet_wrap(~paste0(Gene_of_Interest, "\n", qseqid))

p_L_crisp_blastp_hit_quality

#Table of quality of blastp hits by query gene
blastp_all_eggNOG_hits_filt %>% 
  dplyr::left_join(gene_pres_Spec_SGB_times_seen %>% 
                     dplyr::filter(Num_Genomes_Present == "Gene in Multiple Genomes") %>% 
                     dplyr::count(Gene_Group, Species, name = "Num_genomes"),
                   by = c("sseqid" = "Gene_Group")) %>% 
  dplyr::count(Species,
               Gene_of_Interest, 
               qseqid, 
               sseqid,
               `Seed ortholog`,
               evalue,
               pident,
               Num_genomes,
               eggNOG_curated_match,
               name = "Num_Hits") %>% 
  dplyr::filter(Species == filt_species) %>% 
  tidyr::pivot_wider(names_from = c(eggNOG_curated_match),
                     names_sep = " ",
                     names_prefix = "match ",
                     values_from = Num_Hits,
                     values_fill = 0) %>% 
  dplyr::arrange(Gene_of_Interest) %>% 
  dplyr::select(-qseqid) %>% 
  dplyr::mutate(Species = str_replace(Species, "actobacillus_", " ")) %>% 
  dplyr::rename(`blastp Gene of Interest` = Gene_of_Interest,
                `Pan-genome Gene ID` = sseqid,
                `eggNOG Seed ortholog` = `Seed ortholog`,
                `blastp e-value` = evalue,
                `blastp % identity` = pident,
                `# Genomes Containing Gene` = Num_genomes) %>% 
  knitr::kable(caption = paste0(filt_species, " pan-genome blastp hits, strength of hit, # of genomes with gene, and whether they match genes with the same curated eggNOG annotations, match genes with conflicting annotations based on the curated eggNOG analysis, or match genes that weren't picked out by the curated eggNOG approach"))
```


```{r tally-all-blast-hits-L-crisp, echo=FALSE}
#Deduplicated table of pan-genome genes not ID'ed by curated eggNOG search
#that were identified by one or more blastp query genes
blastp_all_eggNOG_hits_filt %>% 
  dplyr::left_join(gene_pres_Spec_SGB_times_seen %>% 
                     dplyr::filter(Num_Genomes_Present == "Gene in Multiple Genomes") %>% 
                     dplyr::count(Gene_Group, Species, name = "Num_genomes"),
                   by = c("sseqid" = "Gene_Group")) %>% 
  dplyr::filter(Species == filt_species) %>% 
  dplyr::arrange(Gene_of_Interest, qseqid, evalue) %>% 
  dplyr::filter(Gene_of_Interest != Name_to_Plot |
                  is.na(Name_to_Plot)) %>%
  dplyr::left_join(pangenome_AA_df %>% 
                     dplyr::mutate(Gene_ID = str_remove(Fasta_Header, " .*$") %>% 
                                     str_remove("^>")),
                   by = c("sseqid" = "Gene_ID")) %>% 
  dplyr::select(Species, Gene_of_Interest, sseqid, `Seed ortholog`, Num_genomes, 
                eggNOG_curated_match, evalue, pident, everything()) %>% 
  dplyr::distinct(Species, Gene_of_Interest, sseqid, AA_Seq) %>% 
  dplyr::add_count(sseqid, 
                   name = "# blastp Genes of Interest Hitting Gene") %>% 
  dplyr::mutate(Hit = Gene_of_Interest,
                Species = str_replace(Species, "actobacillus_", " ")) %>% 
  tidyr::pivot_wider(names_from = Gene_of_Interest,
                     values_from = Hit) %>% 
  dplyr::select(-AA_Seq, everything(), AA_Seq) %>% 
  dplyr::arrange(CJAA, 
                 CYDC, CYDD, 
                 GSIA, GSIC, GSID,
                 MCCB,
                 TCYB, TCYC, 
                 TCYN) %>% 
  dplyr::left_join(EggNOG_annotations_raw,
                   by = c("sseqid" = "Query")) %>% 
  dplyr::mutate(`KEGG KO` = str_replace_all(`KEGG KO`, ",", " ")) %>% 
  dplyr::select(Species,
                `Pan-genome Gene ID` = sseqid,
                `eggNOG Preferred name` = Preferred_name,
                `eggNOG KEGG KO` = `KEGG KO`,
                everything()) %>% 
  dplyr::select(-c(AA_Seq:description)) %>% 
  knitr::kable(caption = paste0(filt_species, " unique pan-genome genes NOT identified by curated eggNOG annotation approach that were hit by one or more blastp queries"))
  
#Fasta format of genes from above
blastp_all_eggNOG_hits_filt %>% 
  dplyr::left_join(gene_pres_Spec_SGB_times_seen %>% 
                     dplyr::filter(Num_Genomes_Present == "Gene in Multiple Genomes") %>% 
                     dplyr::count(Gene_Group, Species, name = "Num_genomes"),
                   by = c("sseqid" = "Gene_Group")) %>% 
  dplyr::filter(Species == filt_species) %>% 
  dplyr::arrange(Gene_of_Interest, qseqid, evalue) %>% 
  dplyr::filter(Gene_of_Interest != Name_to_Plot |
                  is.na(Name_to_Plot)) %>%
  dplyr::left_join(pangenome_AA_df %>% 
                     dplyr::mutate(Gene_ID = str_remove(Fasta_Header, " .*$") %>% 
                                     str_remove("^>")),
                   by = c("sseqid" = "Gene_ID")) %>% 
  dplyr::select(Species, Gene_of_Interest, sseqid, `Seed ortholog`, Num_genomes, 
                eggNOG_curated_match, evalue, pident, everything()) %>% 
  dplyr::distinct(Species, Gene_of_Interest, sseqid, AA_Seq) %>% 
  dplyr::mutate(Hit = Gene_of_Interest,
                Species = str_replace(Species, "actobacillus_", " ")) %>% 
  tidyr::pivot_wider(names_from = Gene_of_Interest,
                     values_from = Hit) %>% 
  dplyr::select(-Species, everything(), Species) %>% 
  dplyr::select(-AA_Seq, everything(), AA_Seq) %>% 
  dplyr::arrange(CJAA, 
                 CYDC, CYDD, 
                 GSIA, GSIC, GSID,
                 MCCB,
                 TCYB, TCYC, 
                 TCYN) %>% 
  tidyr::unite(col = "Fasta_Header",
               -AA_Seq,
               sep = " ",
               na.rm = TRUE) %>% 
  dplyr::mutate(Fasta_Header = paste0(">", Fasta_Header)) %>% 
  tidyr::pivot_longer(cols = everything(),
                      names_to = "Label",
                      values_to = "Fasta") %>% 
  dplyr::select(Fasta) %>% 
  knitr::kable()
# 
# #More extensive table of blastp results as above
# blastp_all_eggNOG_hits_filt %>% 
#   dplyr::left_join(gene_pres_Spec_SGB_times_seen %>% 
#                      dplyr::filter(Num_Genomes_Present == "Gene in Multiple Genomes") %>% 
#                      dplyr::count(Gene_Group, Species, name = "Num_genomes"),
#                    by = c("sseqid" = "Gene_Group")) %>% 
#   dplyr::filter(Species == filt_species) %>% 
#   dplyr::arrange(Gene_of_Interest, qseqid, evalue) %>% 
#   dplyr::filter(Gene_of_Interest != Name_to_Plot |
#                   is.na(Name_to_Plot)) %>%
#   dplyr::left_join(pangenome_AA_df %>% 
#                      dplyr::mutate(Gene_ID = str_remove(Fasta_Header, " .*$") %>% 
#                                      str_remove("^>")),
#                    by = c("sseqid" = "Gene_ID")) %>% 
#   dplyr::select(Species, Gene_of_Interest, sseqid, `Seed ortholog`, Num_genomes, 
#                 eggNOG_curated_match, evalue, pident, everything()) %>% 
#   dplyr::distinct(Species, Gene_of_Interest, sseqid, `Seed ortholog`, AA_Seq) %>% 
#   dplyr::add_count(sseqid, `Seed ortholog`, name = "# blastp Genes of Interest Hitting Gene") %>%
#   dplyr::mutate(Hit = Gene_of_Interest,
#                 Species = str_replace(Species, "actobacillus_", " ")) %>% 
#   tidyr::pivot_wider(names_from = Gene_of_Interest,
#                      values_from = Hit) %>% 
#   dplyr::select(-AA_Seq, everything(), AA_Seq) %>% 
#   dplyr::arrange(CJAA, 
#                  CYDC, CYDD, 
#                  GSIA, GSIC, GSID,
#                  MCCB,
#                  TCYB, TCYC, 
#                  TCYN) %>% 
#   dplyr::left_join(EggNOG_annotations_raw,
#                    by = c("sseqid" = "Query",
#                           "Seed ortholog" = "Seed ortholog")) %>% 
#   dplyr::select(-Species)
```
  
As noted above, the initial analysis suggests there might possibly be the following genes/transporters in _L. crispatus_ that could potentially alter conclusions in the paper if present:
*  gsiCD (no gsiB)
*  tcyBC (no tcyA)
*  tcyLN (no tcyJ, tcyK or tcyM)
  
  
Conclusions for _L. crispatus_:  
  
*  The sseqid MNKKLCJA_00906 identified by the blastp analysis as cjaA is annotated by eggNOG as encoding glnH, which is supported by blasting the sequence against the NCBI nr and swissprot database.
*  The sseqids MNKKLCJA_01040 and PAGDOOJM_00891 identified by the blastp analysis as gsiA, tcyC, and tcyN are annotated by eggNOG as encoding metN, "Part of the ABC transporter complex MetNIQ involved in methionine import. Responsible for energy coupling to the transport system". This is supported by blasting the sequence against the NCBI nr and swissprot database.
*  The sseqid EBMMIDFH_00010 identified by the blastp analysis as gsiC is annotated by eggNOG as encoding oppB, "ABC-type dipeptide oligopeptide nickel transport systems, permease components". This is supported by blasting the sequence against the NCBI nr and swissprot database, although there are weaker hits to gsiC in swissprot.
*  The sseqids LOGLDDBF_00049 and EBMMIDFH_00011 identified by the blastp analysis as gsiD are annotated by eggNOG as encoding oppC, "ABC-type dipeptide oligopeptide nickel transport systems, permease components". This is supported by blasting the sequence against the NCBI nr and swissprot database, although there are weaker hits to gsiD in swissport.
*  The sseqids MKHNNNEO_00092 and EBMMIDFH_00341 identified by the blastp analysis as tcyB and tcyL are annotated by eggNOG as encoding glnPH2, which is supported by blasting the sequence against the NCBI nr based on hits to other _Lactobacillus_ genomes. Blasting against swissprot database identifies possible hits to glnP, tcyB, and tcyL, all at fairly low identity.
*  The sseqids GDCMONNL_01174, EBMMIDFH_00493 and HGHLAHIM_00926 identified by the blastp analysis as tcyB are annotated by eggNOG as encoding glnP, which is supported by blasting the sequence against the NCBI nr based on hits to other _Lactobacillus_ genomes. Blasting against swissprot database identifies possible hits toglnP, tcyB, and tcyL, all at fairly low identity.
*  There are a large number of sseqids annotated by blastp as having hits to tcyC and tcyN, as well as a few that only had hits to one or the other. These appear (based on review of eggNOG results, blast to the NCBI nr database, and blastp to swissprot) to encompass a broad range of ATP-binding proteins associated with ABC transporters of various types including amino acid (mostly polar amino acids), polyamines, phosphates, and a few carbohydrates. Some of the polar amino acids have some degree of homology to tcyC and tcyN in swissprot, although never as the top hit.  
  
  
Thus, overall _L. crispatus_ does not have definitively annotated evidence of additional glutathione or cystine uptake mechanisms. But it could possibly have either of these activities because it does containg some predicted oligopeptide transport mechanisms with some homology to glutathione transporters and a number of transporters for polar amino acids with some homology to some of the proteins within the cystine-transport loci tcyBC and tcyLN (although lacking obvious homologs of tcyA, tcyJ, tcyK, or tcyM). However on the basis of this analysis, would not change conclusions for the manuscript.  
   
   
## Analysis of results for blastp hits to genes from _Lactobacillus iners_  
  
  
```{r species-on-which-to-filter-li, echo=FALSE}
#Specify species to filter on
filt_species <- "Lactobacillus_iners"
```
  
  
Filtering results for `r filt_species`.

```{r quality-all-blast-hits-L-iners, echo=FALSE}
#Plot quality of blastp hits by query gene
p_L_iners_blastp_hit_quality <- blastp_all_eggNOG_hits_filt %>% 
  dplyr::left_join(gene_pres_Spec_SGB_times_seen %>% 
                     dplyr::filter(Num_Genomes_Present == "Gene in Multiple Genomes") %>% 
                     dplyr::count(Gene_Group, Species, name = "Num_genomes"),
                   by = c("sseqid" = "Gene_Group")) %>% 
  dplyr::filter(Species == filt_species) %>% 
  dplyr::rename(`blastp % Identity` = pident,
                `blastp e-value` = evalue,
                `Matches\nCurated\neggNOG\nannotation` = eggNOG_curated_match,
                `#\nGenomes` = Num_genomes) %>% 
  ggplot(aes(x = `blastp % Identity`, 
             y = `blastp e-value`, 
             color = `Matches\nCurated\neggNOG\nannotation`, 
             alpha = `Matches\nCurated\neggNOG\nannotation`, 
             size = `#\nGenomes`)) +
  geom_point() +
  scale_y_log10() +
  scale_alpha_manual(values = c(1, 1),
                     na.translate = TRUE,
                     na.value = 0.3) +
  labs(title = filt_species) +
  facet_wrap(~paste0(Gene_of_Interest, "\n", qseqid))

p_L_iners_blastp_hit_quality

#Table of quality of blastp hits by query gene
blastp_all_eggNOG_hits_filt %>% 
  dplyr::left_join(gene_pres_Spec_SGB_times_seen %>% 
                     dplyr::filter(Num_Genomes_Present == "Gene in Multiple Genomes") %>% 
                     dplyr::count(Gene_Group, Species, name = "Num_genomes"),
                   by = c("sseqid" = "Gene_Group")) %>% 
  dplyr::count(Species,
               Gene_of_Interest, 
               qseqid, 
               sseqid,
               `Seed ortholog`,
               evalue,
               pident,
               Num_genomes,
               eggNOG_curated_match,
               name = "Num_Hits") %>% 
  dplyr::filter(Species == filt_species) %>% 
  tidyr::pivot_wider(names_from = c(eggNOG_curated_match),
                     names_sep = " ",
                     names_prefix = "match ",
                     values_from = Num_Hits,
                     values_fill = 0) %>% 
  dplyr::arrange(Gene_of_Interest) %>% 
  dplyr::select(-qseqid) %>% 
  dplyr::mutate(Species = str_replace(Species, "actobacillus_", " ")) %>% 
  dplyr::rename(`blastp Gene of Interest` = Gene_of_Interest,
                `Pan-genome Gene ID` = sseqid,
                `eggNOG Seed ortholog` = `Seed ortholog`,
                `blastp e-value` = evalue,
                `blastp % identity` = pident,
                `# Genomes Containing Gene` = Num_genomes) %>% 
  knitr::kable(caption = paste0(filt_species, " pan-genome blastp hits, strength of hit, # of genomes with gene, and whether they match genes with the same curated eggNOG annotations, match genes with conflicting annotations based on the curated eggNOG analysis, or match genes that weren't picked out by the curated eggNOG approach"))
```


```{r tally-all-blast-hits-L-iners, echo=FALSE}
#Deduplicated table of pan-genome genes not ID'ed by curated eggNOG search
#that were identified by one or more blastp query genes
blastp_all_eggNOG_hits_filt %>% 
  dplyr::left_join(gene_pres_Spec_SGB_times_seen %>% 
                     dplyr::filter(Num_Genomes_Present == "Gene in Multiple Genomes") %>% 
                     dplyr::count(Gene_Group, Species, name = "Num_genomes"),
                   by = c("sseqid" = "Gene_Group")) %>% 
  dplyr::filter(Species == filt_species) %>% 
  dplyr::arrange(Gene_of_Interest, qseqid, evalue) %>% 
  dplyr::filter(Gene_of_Interest != Name_to_Plot |
                  is.na(Name_to_Plot)) %>%
  dplyr::left_join(pangenome_AA_df %>% 
                     dplyr::mutate(Gene_ID = str_remove(Fasta_Header, " .*$") %>% 
                                     str_remove("^>")),
                   by = c("sseqid" = "Gene_ID")) %>% 
  dplyr::select(Species, Gene_of_Interest, sseqid, `Seed ortholog`, Num_genomes, 
                eggNOG_curated_match, evalue, pident, everything()) %>% 
  dplyr::distinct(Species, Gene_of_Interest, sseqid, AA_Seq) %>% 
  dplyr::add_count(sseqid, 
                   name = "# blastp Genes of Interest Hitting Gene") %>% 
  dplyr::mutate(Hit = Gene_of_Interest,
                Species = str_replace(Species, "actobacillus_", " ")) %>% 
  tidyr::pivot_wider(names_from = Gene_of_Interest,
                     values_from = Hit) %>% 
  dplyr::select(-AA_Seq, everything(), AA_Seq) %>% 
  dplyr::arrange(CYDC, GSIA, TCYB, TCYC, TCYN) %>% 
  dplyr::left_join(EggNOG_annotations_raw,
                   by = c("sseqid" = "Query")) %>% 
  dplyr::mutate(`KEGG KO` = str_replace_all(`KEGG KO`, ",", " ")) %>% 
  dplyr::select(Species,
                `Pan-genome Gene ID` = sseqid,
                `eggNOG Preferred name` = Preferred_name,
                `eggNOG KEGG KO` = `KEGG KO`,
                everything()) %>% 
  dplyr::select(-c(AA_Seq:description)) %>% 
  knitr::kable(caption = paste0(filt_species, " unique pan-genome genes NOT identified by curated eggNOG annotation approach that were hit by one or more blastp queries"))
  
#Fasta format of genes from above
blastp_all_eggNOG_hits_filt %>% 
  dplyr::left_join(gene_pres_Spec_SGB_times_seen %>% 
                     dplyr::filter(Num_Genomes_Present == "Gene in Multiple Genomes") %>% 
                     dplyr::count(Gene_Group, Species, name = "Num_genomes"),
                   by = c("sseqid" = "Gene_Group")) %>% 
  dplyr::filter(Species == filt_species) %>% 
  dplyr::arrange(Gene_of_Interest, qseqid, evalue) %>% 
  dplyr::filter(Gene_of_Interest != Name_to_Plot |
                  is.na(Name_to_Plot)) %>%
  dplyr::left_join(pangenome_AA_df %>% 
                     dplyr::mutate(Gene_ID = str_remove(Fasta_Header, " .*$") %>% 
                                     str_remove("^>")),
                   by = c("sseqid" = "Gene_ID")) %>% 
  dplyr::select(Species, Gene_of_Interest, sseqid, `Seed ortholog`, Num_genomes, 
                eggNOG_curated_match, evalue, pident, everything()) %>% 
  dplyr::distinct(Species, Gene_of_Interest, sseqid, AA_Seq) %>% 
  dplyr::mutate(Hit = Gene_of_Interest,
                Species = str_replace(Species, "actobacillus_", " ")) %>% 
  tidyr::pivot_wider(names_from = Gene_of_Interest,
                     values_from = Hit) %>% 
  dplyr::select(-Species, everything(), Species) %>% 
  dplyr::select(-AA_Seq, everything(), AA_Seq) %>% 
  dplyr::arrange(CYDC, GSIA, TCYB, TCYC, TCYN) %>% 
  tidyr::unite(col = "Fasta_Header",
               -AA_Seq,
               sep = " ",
               na.rm = TRUE) %>% 
  dplyr::mutate(Fasta_Header = paste0(">", Fasta_Header)) %>% 
  tidyr::pivot_longer(cols = everything(),
                      names_to = "Label",
                      values_to = "Fasta") %>% 
  dplyr::select(Fasta) %>% 
  knitr::kable()
# 
# #More extensive table of blastp results as above
# blastp_all_eggNOG_hits_filt %>% 
#   dplyr::left_join(gene_pres_Spec_SGB_times_seen %>% 
#                      dplyr::filter(Num_Genomes_Present == "Gene in Multiple Genomes") %>% 
#                      dplyr::count(Gene_Group, Species, name = "Num_genomes"),
#                    by = c("sseqid" = "Gene_Group")) %>% 
#   dplyr::filter(Species == filt_species) %>% 
#   dplyr::arrange(Gene_of_Interest, qseqid, evalue) %>% 
#   dplyr::filter(Gene_of_Interest != Name_to_Plot |
#                   is.na(Name_to_Plot)) %>%
#   dplyr::left_join(pangenome_AA_df %>% 
#                      dplyr::mutate(Gene_ID = str_remove(Fasta_Header, " .*$") %>% 
#                                      str_remove("^>")),
#                    by = c("sseqid" = "Gene_ID")) %>% 
#   dplyr::select(Species, Gene_of_Interest, sseqid, `Seed ortholog`, Num_genomes, 
#                 eggNOG_curated_match, evalue, pident, everything()) %>% 
#   dplyr::distinct(Species, Gene_of_Interest, sseqid, `Seed ortholog`, AA_Seq) %>% 
#   dplyr::add_count(sseqid, `Seed ortholog`, name = "# blastp Genes of Interest Hitting Gene") %>% 
#   dplyr::mutate(Hit = Gene_of_Interest,
#                 Species = str_replace(Species, "actobacillus_", " ")) %>% 
#   tidyr::pivot_wider(names_from = Gene_of_Interest,
#                      values_from = Hit) %>% 
#   dplyr::select(-AA_Seq, everything(), AA_Seq) %>% 
#   dplyr::arrange(CYDC, GSIA, TCYB, TCYC, TCYN) %>% 
#   dplyr::left_join(EggNOG_annotations_raw,
#                    by = c("sseqid" = "Query",
#                           "Seed ortholog" = "Seed ortholog"))
```

Conclusions for _L. iners_:  
  
*  All of the sequences identified by blastp as cydC are relatively weak hits annotated by eggNOG simply as having ABC transporter transmembrane regions. blasting the protein sequences against the NCBI nr database shows hits to non-specifically annotated ABC transport ATB-binding protein/permease from various _Lactobacillus_ species.
    +  The sseqids are NIJFCAFB_00079, AKCJOOLM_00162, POAHFJFJ_00637, BJFOJFJB_00284, EIKKBBHP_00152, JGMPPACM_00205
*  The three sequences with sseqids KAHGKJGN_00748, NGJEHHKD_00897, KAIOAAFN_00075 that the blastp analysis showed as hits to gsiA, tcyC, and tcyN are all annotated by eggNOG as metN, "Part of the ABC transporter complex MetNIQ involved in methionine import. Responsible for energy coupling to the transport system". This matches the annotations of the hits by blast against the NCBI nr database
*  The three sequences with sseqids AKCJOOLM_00480, KBPBBFLM_00700, KAIOAAFN_00192 identified by the blastp analysis as possibly gsiA are annotated by eggNOG as oppD, which is supported by blasting the sequence against the NCBI nr database.
*  The sseqid AKCJOOLM_00245 identified by the blastp analysis as tcyB is annotated by eggNOG as glnP, which is supported by blasting the sequence against the NCBI nr database.
*  The sseqid KAIOAAFN_00200 identified by the blastp analysis as tcyB is annotated by eggNOG as gluC, which is supported by blasting the sequence against the NCBI nr database.
*  The sseqids IAHDPDNI_00249, KAIOAAFN_00297, and KMMMAOFL_00779 identified by the blastp analysis as tcyC and tcyN are annotated by eggNOG as glnQ, which encodes an ATP-binding protein for a polar amino acid transporter (Asn/Gln/Glu), which is supported by blasting the sequence against the NCBI nr database.
*  The sseqids KAIOAAFN_00153 and KBPBBFLM_00127 identified by the blastp analysis as tcyC and tcyN are annotated by eggNOG as encoding lysY or lysX2, encoding an ATP-binding protein from a lysine transport system. Blast against the nr database doesn't provide clear indications.
*  The sseqids KBPBBFLM_00532 and NGJEHHKD_00956 identified by the blastp analysis as tcyC and tcyN are annotated by eggNOG as encoding pstB, "part of the ABC transporter complex PstSACB involved in phosphate import", which is supported by blasting the sequence against the NCBI nr database.
*  The sseqids MFNDNHPG_00386 and KAIOAAFN_00289 identified by the blastp analysis as tcyC and tcyN (relatively weaker hits) are annotated by eggNOG as encoding a non-specific putative ABC transport system ATP-binding protein. Blast against the NCBI nr database is similarly non-specific; blastp against the swissprot database suggests this may be orthologous to an ATP-binding protein for a lipoprotein-binding system.
*  The sseqid KAIOAAFN_00049 identified by the blastp analysis as tcyC and tcyN is annotated by eggNOG as encoding potA, "Part of the ABC transporter complex PotABCD involved in spermidine putrescine import. Responsible for energy coupling to the transport system." This is supported by blasting the sequence against the NCBI nr and swissprot database.
*  The sseqid NIJFCAFB_00198 identified by the blastp analysis as tcyC is annotated by eggNOG as encoding a non-specific putative ABC transport system ATP-binding protein. Blast against the NCBI nr database is similarly non-specific; blastp against the swissprot database suggests this may be orthologous to an ATP-binding protein for a lipoprotein-binding system.
*  The sseqid KBPBBFLM_00533 identified by the blastp analysis as tcyN is annotated by eggNOG as encoding pstB, "part of the ABC transporter complex PstSACB involved in phosphate import", which is supported by blasting the sequence against the NCBI nr and swissprot database.
*  The sseqids KBPBBFLM_00779, KAIOAAFN_00542 identified by the blastp analysis as tcyN are annotated by eggNOG as encoding an inositol-phosphate and/or sugar transport protein (e.g. malK). Blast against the NCBI nr database is similarly non-specific; blastp against the swissprot database is is consistent with this potentially being malK or another carbohydrate transporter.
*  The sseqids AKCJOOLM_00528, KBPBBFLM_00036, KAIOAAFN_00027 identified by the blastp analysis as tcyN are annotated by eggNOG as encoding ecfA, "ATP-binding (A) component of a common energy-coupling factor (ECF) ABC-transporter complex. Unlike classic ABC transporters this ECF transporter provides the energy necessary to transport a number of different substrates". This is supported by blasting the sequence against the NCBI nr and swissprot database.  
  
  
This, this analysis identifies some ATP-binding proteins in ABC transport systems with unknown substrates or with likely alternate substrates including polar amino acids (transporters for which have some similarity to cystine transporters), but nothing definitively consistent with cysteine, cystine, or glutathione transporters in _L. iners_.  
  
  
## Analysis of results for blastp hits to genes from _Lactobacillus jensenii_  
  
  
```{r species-on-which-to-filter-lj, echo=FALSE}
#Specify species to filter on
filt_species <- "Lactobacillus_jensenii"
```
  
  
Filtering results for `r filt_species`.

```{r quality-all-blast-hits-L-jensenii, echo=FALSE}
#Plot quality of blastp hits by query gene
p_L_jensenii_blastp_hit_quality <- blastp_all_eggNOG_hits_filt %>% 
  dplyr::left_join(gene_pres_Spec_SGB_times_seen %>% 
                     dplyr::filter(Num_Genomes_Present == "Gene in Multiple Genomes") %>% 
                     dplyr::count(Gene_Group, Species, name = "Num_genomes"),
                   by = c("sseqid" = "Gene_Group")) %>% 
  dplyr::filter(Species == filt_species) %>% 
  # dplyr::group_by(qseqid) %>% 
  # dplyr::filter(!all(eggNOG_curated_match) |
  #                 is.na(all(eggNOG_curated_match))) %>% 
  # dplyr::ungroup() %>% 
  dplyr::rename(`blastp % Identity` = pident,
                `blastp e-value` = evalue,
                `Matches\nCurated\neggNOG\nannotation` = eggNOG_curated_match,
                `#\nGenomes` = Num_genomes) %>% 
  ggplot(aes(x = `blastp % Identity`, 
             y = `blastp e-value`, 
             color = `Matches\nCurated\neggNOG\nannotation`, 
             alpha = `Matches\nCurated\neggNOG\nannotation`, 
             size = `#\nGenomes`)) +
  geom_point() +
  scale_y_log10() +
  scale_alpha_manual(values = c(1, 1),
                     na.translate = TRUE,
                     na.value = 0.3) +
  labs(title = filt_species) +
  facet_wrap(~paste0(Gene_of_Interest, "\n", qseqid))

p_L_jensenii_blastp_hit_quality

#Table of quality of blastp hits by query gene
blastp_all_eggNOG_hits_filt %>% 
  dplyr::left_join(gene_pres_Spec_SGB_times_seen %>% 
                     dplyr::filter(Num_Genomes_Present == "Gene in Multiple Genomes") %>% 
                     dplyr::count(Gene_Group, Species, name = "Num_genomes"),
                   by = c("sseqid" = "Gene_Group")) %>% 
  dplyr::count(Species,
               Gene_of_Interest, 
               qseqid, 
               sseqid,
               `Seed ortholog`,
               evalue,
               pident,
               Num_genomes,
               eggNOG_curated_match,
               name = "Num_Hits") %>% 
  dplyr::filter(Species == filt_species) %>% 
  tidyr::pivot_wider(names_from = c(eggNOG_curated_match),
                     names_sep = " ",
                     names_prefix = "match ",
                     values_from = Num_Hits,
                     values_fill = 0) %>% 
  dplyr::arrange(Gene_of_Interest) %>% 
  dplyr::select(-qseqid) %>% 
  dplyr::mutate(Species = str_replace(Species, "actobacillus_", " ")) %>% 
  dplyr::rename(`blastp Gene of Interest` = Gene_of_Interest,
                `Pan-genome Gene ID` = sseqid,
                `eggNOG Seed ortholog` = `Seed ortholog`,
                `blastp e-value` = evalue,
                `blastp % identity` = pident,
                `# Genomes Containing Gene` = Num_genomes) %>% 
  knitr::kable(caption = paste0(filt_species, " pan-genome blastp hits, strength of hit, # of genomes with gene, and whether they match genes with the same curated eggNOG annotations, match genes with conflicting annotations based on the curated eggNOG analysis, or match genes that weren't picked out by the curated eggNOG approach"))
```


```{r tally-all-blast-hits-L-jensenii, echo=FALSE}
#Deduplicated table of pan-genome genes not ID'ed by curated eggNOG search
#that were identified by one or more blastp query genes
blastp_all_eggNOG_hits_filt %>% 
  dplyr::left_join(gene_pres_Spec_SGB_times_seen %>% 
                     dplyr::filter(Num_Genomes_Present == "Gene in Multiple Genomes") %>% 
                     dplyr::count(Gene_Group, Species, name = "Num_genomes"),
                   by = c("sseqid" = "Gene_Group")) %>% 
  dplyr::filter(Species == filt_species) %>% 
  dplyr::arrange(Gene_of_Interest, qseqid, evalue) %>% 
  dplyr::filter(Gene_of_Interest != Name_to_Plot |
                  is.na(Name_to_Plot)) %>%
  dplyr::left_join(pangenome_AA_df %>% 
                     dplyr::mutate(Gene_ID = str_remove(Fasta_Header, " .*$") %>% 
                                     str_remove("^>")),
                   by = c("sseqid" = "Gene_ID")) %>% 
  dplyr::select(Species, Gene_of_Interest, sseqid, `Seed ortholog`, Num_genomes, 
                eggNOG_curated_match, evalue, pident, everything()) %>% 
  dplyr::distinct(Species, Gene_of_Interest, sseqid, AA_Seq) %>% 
  dplyr::add_count(sseqid, 
                   name = "# blastp Genes of Interest Hitting Gene") %>% 
  dplyr::mutate(Hit = Gene_of_Interest,
                Species = str_replace(Species, "actobacillus_", " ")) %>% 
  tidyr::pivot_wider(names_from = Gene_of_Interest,
                     values_from = Hit) %>% 
  dplyr::select(-AA_Seq, everything(), AA_Seq) %>% 
  dplyr::arrange(CYDC, CYDD, 
                 GSIA,
                 TCYB, TCYC, 
                 TCYN) %>% 
  dplyr::left_join(EggNOG_annotations_raw,
                   by = c("sseqid" = "Query")) %>% 
  dplyr::mutate(`KEGG KO` = str_replace_all(`KEGG KO`, ",", " ")) %>% 
  dplyr::select(Species,
                `Pan-genome Gene ID` = sseqid,
                `eggNOG Preferred name` = Preferred_name,
                `eggNOG KEGG KO` = `KEGG KO`,
                everything()) %>% 
  dplyr::select(-c(AA_Seq:description)) %>% 
  knitr::kable(caption = paste0(filt_species, " unique pan-genome genes NOT identified by curated eggNOG annotation approach that were hit by one or more blastp queries"))
  
#Fasta format of genes from above
blastp_all_eggNOG_hits_filt %>% 
  dplyr::left_join(gene_pres_Spec_SGB_times_seen %>% 
                     dplyr::filter(Num_Genomes_Present == "Gene in Multiple Genomes") %>% 
                     dplyr::count(Gene_Group, Species, name = "Num_genomes"),
                   by = c("sseqid" = "Gene_Group")) %>% 
  dplyr::filter(Species == filt_species) %>% 
  dplyr::arrange(Gene_of_Interest, qseqid, evalue) %>% 
  dplyr::filter(Gene_of_Interest != Name_to_Plot |
                  is.na(Name_to_Plot)) %>%
  dplyr::left_join(pangenome_AA_df %>% 
                     dplyr::mutate(Gene_ID = str_remove(Fasta_Header, " .*$") %>% 
                                     str_remove("^>")),
                   by = c("sseqid" = "Gene_ID")) %>% 
  dplyr::select(Species, Gene_of_Interest, sseqid, `Seed ortholog`, Num_genomes, 
                eggNOG_curated_match, evalue, pident, everything()) %>% 
  dplyr::distinct(Species, Gene_of_Interest, sseqid, AA_Seq) %>% 
  dplyr::mutate(Hit = Gene_of_Interest,
                Species = str_replace(Species, "actobacillus_", " ")) %>% 
  tidyr::pivot_wider(names_from = Gene_of_Interest,
                     values_from = Hit) %>% 
  dplyr::select(-Species, everything(), Species) %>% 
  dplyr::select(-AA_Seq, everything(), AA_Seq) %>% 
  dplyr::arrange(CYDC, CYDD, 
                 GSIA,
                 TCYB, TCYC, 
                 TCYN) %>% 
  tidyr::unite(col = "Fasta_Header",
               -AA_Seq,
               sep = " ",
               na.rm = TRUE) %>% 
  dplyr::mutate(Fasta_Header = paste0(">", Fasta_Header)) %>% 
  tidyr::pivot_longer(cols = everything(),
                      names_to = "Label",
                      values_to = "Fasta") %>% 
  dplyr::select(Fasta) %>% 
  knitr::kable()
# 
# #More extensive table of blastp results as above
# blastp_all_eggNOG_hits_filt %>%
#   dplyr::left_join(gene_pres_Spec_SGB_times_seen %>%
#                      dplyr::filter(Num_Genomes_Present == "Gene in Multiple Genomes") %>%
#                      dplyr::count(Gene_Group, Species, name = "Num_genomes"),
#                    by = c("sseqid" = "Gene_Group")) %>%
#   dplyr::filter(Species == filt_species) %>%
#   dplyr::arrange(Gene_of_Interest, qseqid, evalue) %>%
#   dplyr::filter(Gene_of_Interest != Name_to_Plot |
#                   is.na(Name_to_Plot)) %>%
#   dplyr::left_join(pangenome_AA_df %>%
#                      dplyr::mutate(Gene_ID = str_remove(Fasta_Header, " .*$") %>%
#                                      str_remove("^>")),
#                    by = c("sseqid" = "Gene_ID")) %>%
#   dplyr::select(Species, Gene_of_Interest, sseqid, `Seed ortholog`, Num_genomes,
#                 eggNOG_curated_match, evalue, pident, everything()) %>%
#   dplyr::distinct(Species, Gene_of_Interest, sseqid, `Seed ortholog`, AA_Seq) %>%
#   dplyr::add_count(sseqid, `Seed ortholog`, name = "# blastp Genes of Interest Hitting Gene") %>%
#   dplyr::mutate(Hit = Gene_of_Interest,
#                 Species = str_replace(Species, "actobacillus_", " ")) %>%
#   tidyr::pivot_wider(names_from = Gene_of_Interest,
#                      values_from = Hit) %>%
#   dplyr::select(-AA_Seq, everything(), AA_Seq) %>%
#   dplyr::arrange(CYDC, CYDD, 
#                  GSIA,
#                  TCYB, TCYC, 
#                  TCYN) %>% 
#   dplyr::left_join(EggNOG_annotations_raw,
#                    by = c("sseqid" = "Query",
#                           "Seed ortholog" = "Seed ortholog")) %>%
#   dplyr::select(-Species)
```


For _L. jensenii_, the only gene on this list that would potentially change some conclusions for the manuscript would be addition of further cydC or cydD genes. For the genes that were identified as potentially being these genes via our blastp analysis, review via both review of eggNOG annotations and blastp against the NCBI nr and swissprot databases indicate that none are likely to be true cydCD orthologs. Thus there is nothing via this analysis that appears to significantly change conclusions from the manuscript for gene content of _L. jensenii_.   
  
  
  
## Analysis of results for blastp hits to genes from _Lactobacillus gasseri_  
  
  
```{r species-on-which-to-filter-lg, echo=FALSE}
#Specify species to filter on
filt_species <- "Lactobacillus_gasseri"
```
  
  
Filtering results for `r filt_species`.

```{r quality-all-blast-hits-L-gasseri, echo=FALSE}
#Plot quality of blastp hits by query gene
p_L_gasseri_blastp_hit_quality <- blastp_all_eggNOG_hits_filt %>% 
  dplyr::left_join(gene_pres_Spec_SGB_times_seen %>% 
                     dplyr::filter(Num_Genomes_Present == "Gene in Multiple Genomes") %>% 
                     dplyr::count(Gene_Group, Species, name = "Num_genomes"),
                   by = c("sseqid" = "Gene_Group")) %>% 
  dplyr::filter(Species == filt_species) %>% 
  # dplyr::group_by(qseqid) %>% 
  # dplyr::filter(!all(eggNOG_curated_match) |
  #                 is.na(all(eggNOG_curated_match))) %>% 
  # dplyr::ungroup() %>% 
  dplyr::rename(`blastp % Identity` = pident,
                `blastp e-value` = evalue,
                `Matches\nCurated\neggNOG\nannotation` = eggNOG_curated_match,
                `#\nGenomes` = Num_genomes) %>% 
  ggplot(aes(x = `blastp % Identity`, 
             y = `blastp e-value`, 
             color = `Matches\nCurated\neggNOG\nannotation`, 
             alpha = `Matches\nCurated\neggNOG\nannotation`, 
             size = `#\nGenomes`)) +
  geom_point() +
  scale_y_log10() +
  scale_alpha_manual(values = c(1, 1),
                     na.translate = TRUE,
                     na.value = 0.3) +
  labs(title = filt_species) +
  facet_wrap(~paste0(Gene_of_Interest, "\n", qseqid))

p_L_gasseri_blastp_hit_quality

#Table of quality of blastp hits by query gene
blastp_all_eggNOG_hits_filt %>% 
  dplyr::left_join(gene_pres_Spec_SGB_times_seen %>% 
                     dplyr::filter(Num_Genomes_Present == "Gene in Multiple Genomes") %>% 
                     dplyr::count(Gene_Group, Species, name = "Num_genomes"),
                   by = c("sseqid" = "Gene_Group")) %>% 
  dplyr::count(Species,
               Gene_of_Interest, 
               qseqid, 
               sseqid,
               `Seed ortholog`,
               evalue,
               pident,
               Num_genomes,
               eggNOG_curated_match,
               name = "Num_Hits") %>% 
  dplyr::filter(Species == filt_species) %>% 
  tidyr::pivot_wider(names_from = c(eggNOG_curated_match),
                     names_sep = " ",
                     names_prefix = "match ",
                     values_from = Num_Hits,
                     values_fill = 0) %>% 
  dplyr::arrange(Gene_of_Interest) %>% 
  dplyr::select(-qseqid) %>% 
  dplyr::mutate(Species = str_replace(Species, "actobacillus_", " ")) %>% 
  dplyr::rename(`blastp Gene of Interest` = Gene_of_Interest,
                `Pan-genome Gene ID` = sseqid,
                `eggNOG Seed ortholog` = `Seed ortholog`,
                `blastp e-value` = evalue,
                `blastp % identity` = pident,
                `# Genomes Containing Gene` = Num_genomes) %>% 
  knitr::kable(caption = paste0(filt_species, " pan-genome blastp hits, strength of hit, # of genomes with gene, and whether they match genes with the same curated eggNOG annotations, match genes with conflicting annotations based on the curated eggNOG analysis, or match genes that weren't picked out by the curated eggNOG approach"))
```


```{r tally-all-blast-hits-L-gasseri, echo=FALSE}
#Deduplicated table of pan-genome genes not ID'ed by curated eggNOG search
#that were identified by one or more blastp query genes
blastp_all_eggNOG_hits_filt %>% 
  dplyr::left_join(gene_pres_Spec_SGB_times_seen %>% 
                     dplyr::filter(Num_Genomes_Present == "Gene in Multiple Genomes") %>% 
                     dplyr::count(Gene_Group, Species, name = "Num_genomes"),
                   by = c("sseqid" = "Gene_Group")) %>% 
  dplyr::filter(Species == filt_species) %>% 
  dplyr::arrange(Gene_of_Interest, qseqid, evalue) %>% 
  dplyr::filter(Gene_of_Interest != Name_to_Plot |
                  is.na(Name_to_Plot)) %>%
  dplyr::left_join(pangenome_AA_df %>% 
                     dplyr::mutate(Gene_ID = str_remove(Fasta_Header, " .*$") %>% 
                                     str_remove("^>")),
                   by = c("sseqid" = "Gene_ID")) %>% 
  dplyr::select(Species, Gene_of_Interest, sseqid, `Seed ortholog`, Num_genomes, 
                eggNOG_curated_match, evalue, pident, everything()) %>% 
  dplyr::distinct(Species, Gene_of_Interest, sseqid, AA_Seq) %>% 
  dplyr::add_count(sseqid, 
                   name = "# blastp Genes of Interest Hitting Gene") %>% 
  dplyr::mutate(Hit = Gene_of_Interest,
                Species = str_replace(Species, "actobacillus_", " ")) %>% 
  tidyr::pivot_wider(names_from = Gene_of_Interest,
                     values_from = Hit) %>% 
  dplyr::select(-AA_Seq, everything(), AA_Seq) %>% 
  dplyr::arrange(CYDC, 
                 GSIA,
                 TCYB, TCYC, 
                 TCYN) %>% 
  dplyr::left_join(EggNOG_annotations_raw,
                   by = c("sseqid" = "Query")) %>% 
  dplyr::mutate(`KEGG KO` = str_replace_all(`KEGG KO`, ",", " ")) %>% 
  dplyr::select(Species,
                `Pan-genome Gene ID` = sseqid,
                `eggNOG Preferred name` = Preferred_name,
                `eggNOG KEGG KO` = `KEGG KO`,
                everything()) %>% 
  dplyr::select(-c(AA_Seq:description)) %>% 
  knitr::kable(caption = paste0(filt_species, " unique pan-genome genes NOT identified by curated eggNOG annotation approach that were hit by one or more blastp queries"))
  
#Fasta format of genes from above
blastp_all_eggNOG_hits_filt %>% 
  dplyr::left_join(gene_pres_Spec_SGB_times_seen %>% 
                     dplyr::filter(Num_Genomes_Present == "Gene in Multiple Genomes") %>% 
                     dplyr::count(Gene_Group, Species, name = "Num_genomes"),
                   by = c("sseqid" = "Gene_Group")) %>% 
  dplyr::filter(Species == filt_species) %>% 
  dplyr::arrange(Gene_of_Interest, qseqid, evalue) %>% 
  dplyr::filter(Gene_of_Interest != Name_to_Plot |
                  is.na(Name_to_Plot)) %>%
  dplyr::left_join(pangenome_AA_df %>% 
                     dplyr::mutate(Gene_ID = str_remove(Fasta_Header, " .*$") %>% 
                                     str_remove("^>")),
                   by = c("sseqid" = "Gene_ID")) %>% 
  dplyr::select(Species, Gene_of_Interest, sseqid, `Seed ortholog`, Num_genomes, 
                eggNOG_curated_match, evalue, pident, everything()) %>% 
  dplyr::distinct(Species, Gene_of_Interest, sseqid, AA_Seq) %>% 
  dplyr::mutate(Hit = Gene_of_Interest,
                Species = str_replace(Species, "actobacillus_", " ")) %>% 
  tidyr::pivot_wider(names_from = Gene_of_Interest,
                     values_from = Hit) %>% 
  dplyr::select(-Species, everything(), Species) %>% 
  dplyr::select(-AA_Seq, everything(), AA_Seq) %>% 
  dplyr::arrange(CYDC, 
                 GSIA,
                 TCYB, TCYC, 
                 TCYN) %>% 
  tidyr::unite(col = "Fasta_Header",
               -AA_Seq,
               sep = " ",
               na.rm = TRUE) %>% 
  dplyr::mutate(Fasta_Header = paste0(">", Fasta_Header)) %>% 
  tidyr::pivot_longer(cols = everything(),
                      names_to = "Label",
                      values_to = "Fasta") %>% 
  dplyr::select(Fasta) %>% 
  knitr::kable()
# 
# #More extensive table of blastp results as above
# blastp_all_eggNOG_hits_filt %>%
#   dplyr::left_join(gene_pres_Spec_SGB_times_seen %>%
#                      dplyr::filter(Num_Genomes_Present == "Gene in Multiple Genomes") %>%
#                      dplyr::count(Gene_Group, Species, name = "Num_genomes"),
#                    by = c("sseqid" = "Gene_Group")) %>%
#   dplyr::filter(Species == filt_species) %>%
#   dplyr::arrange(Gene_of_Interest, qseqid, evalue) %>%
#   dplyr::filter(Gene_of_Interest != Name_to_Plot |
#                   is.na(Name_to_Plot)) %>%
#   dplyr::left_join(pangenome_AA_df %>%
#                      dplyr::mutate(Gene_ID = str_remove(Fasta_Header, " .*$") %>%
#                                      str_remove("^>")),
#                    by = c("sseqid" = "Gene_ID")) %>%
#   dplyr::select(Species, Gene_of_Interest, sseqid, `Seed ortholog`, Num_genomes,
#                 eggNOG_curated_match, evalue, pident, everything()) %>%
#   dplyr::distinct(Species, Gene_of_Interest, sseqid, `Seed ortholog`, AA_Seq) %>%
#   dplyr::add_count(sseqid, `Seed ortholog`, name = "# blastp Genes of Interest Hitting Gene") %>%
#   dplyr::mutate(Hit = Gene_of_Interest,
#                 Species = str_replace(Species, "actobacillus_", " ")) %>%
#   tidyr::pivot_wider(names_from = Gene_of_Interest,
#                      values_from = Hit) %>%
#   dplyr::select(-AA_Seq, everything(), AA_Seq) %>%
#   dplyr::arrange(CYDC, 
#                  GSIA,
#                  TCYB, TCYC, 
#                  TCYN) %>% 
#   dplyr::left_join(EggNOG_annotations_raw,
#                    by = c("sseqid" = "Query",
#                           "Seed ortholog" = "Seed ortholog")) %>%
#   dplyr::select(-Species)
```


For _L. gasseri_, the only genes on this list that would potentially change some conclusions for the manuscript would be cystine transport tcyB, tcyC, or tcyN genes. For the genes that were identified as potentially being these genes via our blastp analysis, review via both review of eggNOG annotations and blastp against the NCBI nr and swissprot databases indicate that none are likely to be true orthologs of these genes -- most are annotated as transporters for other substrates including other amino acids or other substrates. Thus there is nothing via this analysis that appears to significantly change conclusions from the manuscript for gene content of _L. gasseri_.    
  
  
## Analysis of results for blastp hits to genes from _Lactobacillus vaginalis_  
  
  
```{r species-on-which-to-filter-lv, echo=FALSE}
#Specify species to filter on
filt_species <- "Lactobacillus_vaginalis"
```
  
  
Filtering results for `r filt_species`.

```{r quality-all-blast-hits-L-vaginalis, echo=FALSE}
#Plot quality of blastp hits by query gene
p_L_vaginalis_blastp_hit_quality <- blastp_all_eggNOG_hits_filt %>% 
  dplyr::left_join(gene_pres_Spec_SGB_times_seen %>% 
                     dplyr::filter(Num_Genomes_Present == "Gene in Multiple Genomes") %>% 
                     dplyr::count(Gene_Group, Species, name = "Num_genomes"),
                   by = c("sseqid" = "Gene_Group")) %>% 
  dplyr::filter(Species == filt_species) %>% 
  # dplyr::group_by(qseqid) %>% 
  # dplyr::filter(!all(eggNOG_curated_match) |
  #                 is.na(all(eggNOG_curated_match))) %>% 
  # dplyr::ungroup() %>% 
  dplyr::rename(`blastp % Identity` = pident,
                `blastp e-value` = evalue,
                `Matches\nCurated\neggNOG\nannotation` = eggNOG_curated_match,
                `#\nGenomes` = Num_genomes) %>% 
  ggplot(aes(x = `blastp % Identity`, 
             y = `blastp e-value`, 
             color = `Matches\nCurated\neggNOG\nannotation`, 
             alpha = `Matches\nCurated\neggNOG\nannotation`, 
             size = `#\nGenomes`)) +
  geom_point() +
  scale_y_log10() +
  scale_alpha_manual(values = c(1, 1),
                     na.translate = TRUE,
                     na.value = 0.3) +
  labs(title = filt_species) +
  facet_wrap(~paste0(Gene_of_Interest, "\n", qseqid))

p_L_vaginalis_blastp_hit_quality

#Table of quality of blastp hits by query gene
blastp_all_eggNOG_hits_filt %>% 
  dplyr::left_join(gene_pres_Spec_SGB_times_seen %>% 
                     dplyr::filter(Num_Genomes_Present == "Gene in Multiple Genomes") %>% 
                     dplyr::count(Gene_Group, Species, name = "Num_genomes"),
                   by = c("sseqid" = "Gene_Group")) %>% 
  dplyr::count(Species,
               Gene_of_Interest, 
               qseqid, 
               sseqid,
               `Seed ortholog`,
               evalue,
               pident,
               Num_genomes,
               eggNOG_curated_match,
               name = "Num_Hits") %>% 
  dplyr::filter(Species == filt_species) %>% 
  tidyr::pivot_wider(names_from = c(eggNOG_curated_match),
                     names_sep = " ",
                     names_prefix = "match ",
                     values_from = Num_Hits,
                     values_fill = 0) %>% 
  dplyr::arrange(Gene_of_Interest) %>% 
  dplyr::select(-qseqid) %>% 
  dplyr::mutate(Species = str_replace(Species, "actobacillus_", " ")) %>% 
  dplyr::rename(`blastp Gene of Interest` = Gene_of_Interest,
                `Pan-genome Gene ID` = sseqid,
                `eggNOG Seed ortholog` = `Seed ortholog`,
                `blastp e-value` = evalue,
                `blastp % identity` = pident,
                `# Genomes Containing Gene` = Num_genomes) %>% 
  knitr::kable(caption = paste0(filt_species, " pan-genome blastp hits, strength of hit, # of genomes with gene, and whether they match genes with the same curated eggNOG annotations, match genes with conflicting annotations based on the curated eggNOG analysis, or match genes that weren't picked out by the curated eggNOG approach"))
```


```{r tally-all-blast-hits-L-vaginalis, echo=FALSE}
#Deduplicated table of pan-genome genes not ID'ed by curated eggNOG search
#that were identified by one or more blastp query genes
blastp_all_eggNOG_hits_filt %>% 
  dplyr::left_join(gene_pres_Spec_SGB_times_seen %>% 
                     dplyr::filter(Num_Genomes_Present == "Gene in Multiple Genomes") %>% 
                     dplyr::count(Gene_Group, Species, name = "Num_genomes"),
                   by = c("sseqid" = "Gene_Group")) %>% 
  dplyr::filter(Species == filt_species) %>% 
  dplyr::arrange(Gene_of_Interest, qseqid, evalue) %>% 
  dplyr::filter(Gene_of_Interest != Name_to_Plot |
                  is.na(Name_to_Plot)) %>%
  dplyr::left_join(pangenome_AA_df %>% 
                     dplyr::mutate(Gene_ID = str_remove(Fasta_Header, " .*$") %>% 
                                     str_remove("^>")),
                   by = c("sseqid" = "Gene_ID")) %>% 
  dplyr::select(Species, Gene_of_Interest, sseqid, `Seed ortholog`, Num_genomes, 
                eggNOG_curated_match, evalue, pident, everything()) %>% 
  dplyr::distinct(Species, Gene_of_Interest, sseqid, AA_Seq) %>% 
  dplyr::add_count(sseqid, 
                   name = "# blastp Genes of Interest Hitting Gene") %>% 
  dplyr::mutate(Hit = Gene_of_Interest,
                Species = str_replace(Species, "actobacillus_", " ")) %>% 
  tidyr::pivot_wider(names_from = Gene_of_Interest,
                     values_from = Hit) %>% 
  dplyr::select(-AA_Seq, everything(), AA_Seq) %>% 
  dplyr::arrange(GSIA,
                 MCCB,
                 TCYB, TCYC, 
                 TCYL,
                 TCYN) %>% 
  dplyr::left_join(EggNOG_annotations_raw,
                   by = c("sseqid" = "Query")) %>% 
  dplyr::mutate(`KEGG KO` = str_replace_all(`KEGG KO`, ",", " ")) %>% 
  dplyr::select(Species,
                `Pan-genome Gene ID` = sseqid,
                `eggNOG Preferred name` = Preferred_name,
                `eggNOG KEGG KO` = `KEGG KO`,
                everything()) %>% 
  dplyr::select(-c(AA_Seq:description)) %>% 
  knitr::kable(caption = paste0(filt_species, " unique pan-genome genes NOT identified by curated eggNOG annotation approach that were hit by one or more blastp queries"))
  
#Fasta format of genes from above
blastp_all_eggNOG_hits_filt %>% 
  dplyr::left_join(gene_pres_Spec_SGB_times_seen %>% 
                     dplyr::filter(Num_Genomes_Present == "Gene in Multiple Genomes") %>% 
                     dplyr::count(Gene_Group, Species, name = "Num_genomes"),
                   by = c("sseqid" = "Gene_Group")) %>% 
  dplyr::filter(Species == filt_species) %>% 
  dplyr::arrange(Gene_of_Interest, qseqid, evalue) %>% 
  dplyr::filter(Gene_of_Interest != Name_to_Plot |
                  is.na(Name_to_Plot)) %>%
  dplyr::left_join(pangenome_AA_df %>% 
                     dplyr::mutate(Gene_ID = str_remove(Fasta_Header, " .*$") %>% 
                                     str_remove("^>")),
                   by = c("sseqid" = "Gene_ID")) %>% 
  dplyr::select(Species, Gene_of_Interest, sseqid, `Seed ortholog`, Num_genomes, 
                eggNOG_curated_match, evalue, pident, everything()) %>% 
  dplyr::distinct(Species, Gene_of_Interest, sseqid, AA_Seq) %>% 
  dplyr::mutate(Hit = Gene_of_Interest,
                Species = str_replace(Species, "actobacillus_", " ")) %>% 
  tidyr::pivot_wider(names_from = Gene_of_Interest,
                     values_from = Hit) %>% 
  dplyr::select(-Species, everything(), Species) %>% 
  dplyr::select(-AA_Seq, everything(), AA_Seq) %>% 
  dplyr::arrange(GSIA,
                 MCCB,
                 TCYB, TCYC, 
                 TCYL,
                 TCYN) %>% 
  tidyr::unite(col = "Fasta_Header",
               -AA_Seq,
               sep = " ",
               na.rm = TRUE) %>% 
  dplyr::mutate(Fasta_Header = paste0(">", Fasta_Header)) %>% 
  tidyr::pivot_longer(cols = everything(),
                      names_to = "Label",
                      values_to = "Fasta") %>% 
  dplyr::select(Fasta) %>% 
  knitr::kable()

# #More extensive table of blastp results as above
# blastp_all_eggNOG_hits_filt %>%
#   dplyr::left_join(gene_pres_Spec_SGB_times_seen %>%
#                      dplyr::filter(Num_Genomes_Present == "Gene in Multiple Genomes") %>%
#                      dplyr::count(Gene_Group, Species, name = "Num_genomes"),
#                    by = c("sseqid" = "Gene_Group")) %>%
#   dplyr::filter(Species == filt_species) %>%
#   dplyr::arrange(Gene_of_Interest, qseqid, evalue) %>%
#   dplyr::filter(Gene_of_Interest != Name_to_Plot |
#                   is.na(Name_to_Plot)) %>%
#   dplyr::left_join(pangenome_AA_df %>%
#                      dplyr::mutate(Gene_ID = str_remove(Fasta_Header, " .*$") %>%
#                                      str_remove("^>")),
#                    by = c("sseqid" = "Gene_ID")) %>%
#   dplyr::select(Species, Gene_of_Interest, sseqid, `Seed ortholog`, Num_genomes,
#                 eggNOG_curated_match, evalue, pident, everything()) %>%
#   dplyr::distinct(Species, Gene_of_Interest, sseqid, `Seed ortholog`, AA_Seq) %>%
#   dplyr::add_count(sseqid, `Seed ortholog`, name = "# blastp Genes of Interest Hitting Gene") %>%
#   dplyr::mutate(Hit = Gene_of_Interest,
#                 Species = str_replace(Species, "actobacillus_", " ")) %>%
#   tidyr::pivot_wider(names_from = Gene_of_Interest,
#                      values_from = Hit) %>%
#   dplyr::select(-AA_Seq, everything(), AA_Seq) %>%
#   dplyr::arrange(GSIA,
#                  MCCB,
#                  TCYB, TCYC, 
#                  TCYL,
#                  TCYN) %>% 
#   dplyr::left_join(EggNOG_annotations_raw,
#                    by = c("sseqid" = "Query",
#                           "Seed ortholog" = "Seed ortholog")) %>%
#   dplyr::select(-Species)
```


For _L. vaginalis_, the only genes on this list that would potentially change some conclusions for the manuscript would be the glutathione transport genes in gsiABCD; there are potential gsiA orthologs. Review of these via both review of eggNOG annotations and blastp against the NCBI nr and swissprot databases indicate that these are methionine transport proteins. Thus there is nothing via this analysis that appears to significantly change conclusions from the manuscript for gene content of _L. vaginalis_.  


## Session information  
  
```{r session-info, echo=FALSE}
sessionInfo()
```
````
