## Supplementary File 1 for "Cysteine dependence in *Lactobacillus iners* constitutes a novel therapeutic target to modify the vaginal microbiota": README_L_iners_Genome_Analysis.txt

README file for cysteine-related gene analysis in Bloom et al manuscript on Lactobacillus iners. Contains information on code, analysis files, and directory structure to reproduce the analysis, as well as required dependencies. Additional details on the analysis are contained within the R Markdown analysis file itself.


Directory contains the following files:
* Genome_analysis_for_L_iners_paper_2021_06_06.Rproj
  - (R Project file to be opened in R Studio to perform analysis)
* L_iners_Lacto_Cys_gene_analysis_for_paper_2021_06_06.Rmd
  - (R Markdown file containing analysis code
* This README.txt file: README_L_iners_Genome_Analysis.txt


Also contains the following subdirectories with their contents:

* Lacto_Genome_Data_2020_07_21, which contains the following files:
  - gene_presence_absence_2020_07_21.Rtab
    + A tab-delimited file of gene presence and absence within Lactobacillus genomes, with genomes in columns and genes in rows, where coding of 1 indicates that gene is present in the genome assembly and 0 indicates that it is not present
  - pan_genome_reference_2020_07_21.fa
    + A multi-fasta file of DNA sequences for genes in the Lactobacillus pan-genome, corresponding to the genes in the gene presence-absence table
  - FGT_Lacto_genome_SGB_taxonomy_2020_09_08.txt
    + A tab-delimited file of taxonomic assignments for the Lactobacillus genomes, containing 4 columns with the operational SGB assignment, Species assignment, Genome name, and taxonomically informed Species/SGB assignment
    + NOTE: Some of the genome names in the gene presence-absence table have additional suffixes that prevent directly matching them with the taxonomy table. These are reconciled within R markdown analysis script.


* eggNOG_annotations, which contains the following file:
  - job_MM_atwxphdb_annotations.txt
    + A tab-delimited file of eggNOG output for the genes in the Lactobacillus pan-genome. Not all genes in the overall pan-genome were annotated by eggNOG.


* Genes_of_interest, which contains the following files:
  - Cys_pathway_genes_of_interest3.txt
    + A tab-delimited file for genes of interest in Lactobacillus genome analysis with various associated annotations and details


* blast_output, which contains the following files with output of blastp analysis of query genes against the Lactobacillus pan-genome in form of tab-delimited files:
  - CjaA.blastp.out
  - CydA.blastp.out
  - CydB.blastp.out
  - CydC.blastp.out
  - CydD.blastp.out
  - CysE.blastp.out
  - CysK.blastp.out
  - GsiA.blastp.out
  - GsiB.blastp.out
  - GsiC.blastp.out
  - GsiD.blastp.out
  - LivF.blastp.out
  - LivG.blastp.out
  - LivH.blastp.out
  - LivK.blastp.out
  - LivM.blastp.out
  - MccA.blastp.out
  - MccB.blastp.out
  - TcyA.blastp.out
  - TcyB.blastp.out
  - TcyC.blastp.out
  - TcyJ.blastp.out
  - TcyK.blastp.out
  - TcyL.blastp.out
  - TcyM.blastp.out
  - TcyN.blastp.out
  - TcyP.blastp.out
  - NOTE: The files do not include column names. The column names in order are:
    + qseqid (ID of the query sequence)
    + sseqid (ID of the subject gene identified within the pan-genome)
    + pident (percent identity)
    + length (length of hit)
    + mismatch
    + gapopen
    + qstart (start position of blast match in query sequence)
    + qend (end position of blast match in query sequence)
    + sstart (start position of blast match in subject sequence)
    + send (end position of blast match in subject sequence)
    + evalue (blastp e-value)
    + bitscore


* Output, empty directory into which figures, tables, and other files are saved.

The dependencies for the analysis are as follows:

## R version 3.6.3 (2020-02-29)
## Platform: x86_64-apple-darwin15.6.0 (64-bit)
## Running under: macOS  10.16
## 
## Matrix products: default
## BLAS:   /Library/Frameworks/R.framework/Versions/3.6/Resources/lib/libRblas.0.dylib
## LAPACK: /Library/Frameworks/R.framework/Versions/3.6/Resources/lib/libRlapack.dylib
## 
## locale:
## [1] en_US.UTF-8/en_US.UTF-8/en_US.UTF-8/C/en_US.UTF-8/en_US.UTF-8
## 
## attached base packages:
## [1] stats     graphics  grDevices utils     datasets  methods   base     
## 
## other attached packages:
##  [1] knitr_1.33      ggExtra_0.9     forcats_0.5.1   stringr_1.4.0  
##  [5] dplyr_1.0.5     purrr_0.3.4     readr_1.4.0     tidyr_1.1.3    
##  [9] tibble_3.1.1    ggplot2_3.3.3   tidyverse_1.3.1 seqinr_4.2-5   
## 
## loaded via a namespace (and not attached):
##  [1] Rcpp_1.0.6        lubridate_1.7.10  prettyunits_1.1.1 assertthat_0.2.1 
##  [5] digest_0.6.27     utf8_1.2.1        mime_0.10         R6_2.5.0         
##  [9] cellranger_1.1.0  backports_1.2.1   reprex_2.0.0      evaluate_0.14    
## [13] highr_0.9         httr_1.4.2        pillar_1.6.0      rlang_0.4.10     
## [17] progress_1.2.2    readxl_1.3.1      rstudioapi_0.13   miniUI_0.1.1.1   
## [21] jquerylib_0.1.4   rmarkdown_2.7     labeling_0.4.2    munsell_0.5.0    
## [25] shiny_1.6.0       broom_0.7.6       compiler_3.6.3    httpuv_1.6.0     
## [29] modelr_0.1.8      xfun_0.22         pkgconfig_2.0.3   htmltools_0.5.1.1
## [33] tidyselect_1.1.0  fansi_0.4.2       crayon_1.4.1      dbplyr_2.1.1     
## [37] withr_2.4.2       later_1.2.0       MASS_7.3-53.1     grid_3.6.3       
## [41] xtable_1.8-4      jsonlite_1.7.2    gtable_0.3.0      lifecycle_1.0.0  
## [45] DBI_1.1.1         magrittr_2.0.1    scales_1.1.1      cli_2.5.0        
## [49] stringi_1.5.3     farver_2.1.0      promises_1.2.0.1  fs_1.5.0         
## [53] xml2_1.3.2        bslib_0.2.4       ellipsis_0.3.1    generics_0.1.0   
## [57] vctrs_0.3.7       cowplot_1.1.1     tools_3.6.3       ade4_1.7-16      
## [61] glue_1.4.2        hms_1.0.0         fastmap_1.1.0     yaml_2.2.1       
## [65] colorspace_2.0-0  rvest_1.0.0       haven_2.4.1       sass_0.3.1
