## Supplementary File 2 for "Cysteine dependence in *Lactobacillus iners* constitutes a novel therapeutic target to modify the vaginal microbiota": L_iners_Cysteine_analysis_2021_06_10_v5.Rmd

````
---
title: "Cleaned 16S figure for cysteine analysis, v5"
author: "Seth M. Bloom"
date: "06/10/2021"
output:
  html_document: default
  word_document: default
---


### Introduction

This markdown file explores cysteine concentration in the FGT environment in relation to different bacterial taxa, testing the hypothesis that cysteine and cysteine-containing metabolites have higher concentrations in samples associated with high _Lactobacillus_ relative abundance, including both _L. crispatus_ and _L. iners_.  
  
**NOTE:**  
  
*  Final version for initial manuscript submission.
  

### Load packages  
  

```{r load-packages, echo=FALSE}
library(phyloseq); packageVersion("phyloseq")
library(DescTools); packageVersion("DescTools")
library(e1071); packageVersion("e1071")
library(tidyverse); packageVersion("tidyverse")
library(knitr); packageVersion("knitr")
library(ggpubr); packageVersion("ggpubr")
library(cowplot); packageVersion("cowplot")
library(table1); packageVersion("table1")
```

```{r global-theme-settings, echo=FALSE}
knitr::opts_chunk$set(fig.width=8, 
                      fig.height=8,
                      echo = FALSE)

# Set global theming
theme_set(theme_bw(base_size = 10,
                   base_family = "Helvetica"))
```

### Load 16S Data  
  

```{r load-16S-data, echo=FALSE}
f_ps0.raw <- "Data/FRESH_data/ps0_FRESH_16S_BV_for_cys_metabolomics_2021_03_07_v3.RDS"
f_ps0.raw

ps0.raw <- readRDS(f_ps0.raw)

ps0.raw
```

  
### Process and rarefy 16S data  
  
```{r prune-NA-taxa, echo=FALSE}
#Prune out taxa in ps0 that are NA at Class level
ps0.filt <- prune_taxa(!is.na(as.character(tax_table(ps0.raw)[, "Class"])),
                       ps0.raw)
```

```{r prune-ps0, echo=FALSE}
#Plot read count distribution
p.Readcount.histo.unfiltered <- ps0.filt %>% 
  sample_sums() %>% 
  qplot(bins = 60) +
  scale_x_log10(limits = c(1, NA),
                breaks = 10^c(0:5)) +
  labs(x = "16S Read Count",
       y = "# of Samples",
       title = "Read Counts of 16S-Sequenced Samples (Unfiltered)") +
  theme(plot.title = element_text(hjust = 0.5, face = "bold", size = 14),
        axis.title = element_text(size = 12, face = "bold"),
        axis.text = element_text(size = 11, color = "black"))

p.Readcount.histo.unfiltered


#Review minimum read counts in the dataset
ps0.filt %>% 
  sample_sums() %>% 
  sort() %>% 
  tibble::enframe(name = "Sequence ID",
                  value = "Read Count") %>% 
  head() %>% 
  knitr::kable(caption = "Samples with lowest read counts in cohort")
```

## Prune and rarefy phyloseq object

First exclude samples with read count <10,000.  
  

```{r rarefy-ps0, echo=FALSE}
#Drop samples with <10,000 reads
ps0.rare <- prune_samples(sample_sums(ps0.filt) >= 10000,
                          ps0.filt)

#Prune taxa to remove 0 counts
ps0.rare <- prune_taxa(taxa_sums(ps0.rare) > 0,
                       ps0.rare)

#Rarefy reads to total read count of sample with lowest total count
ps0.rare <- rarefy_even_depth(ps0.rare, 
                              sample.size = min(sample_sums(ps0.rare)), 
                              rngseed = 100, 
                              replace = FALSE)

#Review
ps0.rare
```
  
  
The above step rarefied (down-sampled) the data to a uniform read depth of `r unique(sample_sums(ps0.rare))`, resulting in a limit of detection of 1 / `r unique(sample_sums(ps0.rare))` or `r 1 / unique(sample_sums(ps0.rare))`.  

## Adjust taxonomy and create columns of Genus an binomial Genus-species names  
  
  

```{r create-taxa-table-with-Genus-and-Gen_Spec-names, echo=FALSE}
#Extract ASV taxonomy table
ASV.tax.table <- ps0.rare %>% 
  tax_table() %>% 
  as.data.frame(stringsAsFactors = FALSE) %>% 
  tibble::rownames_to_column("ASV")

#Manually curate to assign species names to key ASVs
#There are two Prevotella species that cannot be easily classified
#and which are being denoted sp_1 and sp_2 here
ASV.tax.table.curated <- ASV.tax.table %>%
  dplyr::mutate(Species = case_when(
    ASV %in% c("GCAAGCGTTGTCCGGAATTATTGGGCGTAAAGCGCGCGCAGGCGGTCTCTTAAGTCCATCTTAGAAGTGCGGGGCTTAACCCCGTGAGGGGATGGAAACTGGGAGACTGGAGTATCGGAGAGGAAAGTGGAATTCCTAGTGTAGCGGTGAAATGCGTAGATATTAGGAAGAACACCGGTGGCGAAGGCGACTTTCTGGACGAAAACTGACGCTGAGGCGC",
               "GCAAGCGTTGTCCGGAATTATTGGGCGTAAAGCGCGCGCAGGCGGTCTCTTAAGTCCATCTTAGAAGTGCGGGGCTTAACCCCGTGATGGGATGGAAACTGGGAGACTGGAGTATCGGAGAGGAAAGTGGAATTCCTAGTGTAGCGGTGAAATGCGTAGATATTAGGAAGAACACCGGTGGCGAAGGCGACTTTCTGGACGAAAACTGACGCTGAGGCGC") ~ "micraerophilus",
    ASV == "TCAGGCGTTATCCGGATTTATTGGGTTTAAAGGGAGCGTAGGCTGTTTGTTAAGCGTGTTGTGAAATGTAAGAGCTCAACTTTTAGATTGCAGCGCGAACTGGCAGACTTGAGTGCGCACAACGTAGGCGGAATTCATGGTGTAGCGGTGAAATGCTTAGATATCATGACGAACTCCGATTGCGAAGGCAGCTTACGGGAGCGCAACTGACGCTAAAGCT" ~ "amnii",
    ASV == "CCGGGCGTTATCCGGATTTATTGGGTTTAAAGGGAGCGCAGGCCGCCCGATAAGCGTGTTGTGAAATGTACCGGCTCAACCGGTGAGTTGCAGCGCGAACTGTCAGGCTTGAGTGCACGGTAAGCAGGCGGAATTCATGGTGTAGCGGTGAAATGCTTAGATATCATGAGGAACTCCGATTGCGAAGGCAGCTTGCTGCAGTGCGACTGACGCTTAGGCT" ~ "sp_1",
    ASV == "CCAGGCGTTATCCGGATTTATTGGGTTTAAAGGGAGTGTAGGCTGTTAGATAAGCGTGTTGTGAAATATTCTAGCTTAACTAGAATCGTGCAGCGCGAACTGTTTAACTTGAGTTCGTTCAACGTAGGCGGAATTTATGGTGTAGCGGTGAAATGCTTAGATATCATAAAGAACTCCGATTGCGAAGGCAGCTTACGGGGTCGTAACTGACGCTAAAGCT" ~ "sp_2",
    TRUE ~ Species))

#Create columns Species_Name (binomial name) and Genus_Name
ASV.tax.table.names <- ASV.tax.table.curated %>%
  dplyr::mutate(
    Species_Name = case_when(
      Species == "BVAB2" ~ Species,
      is.na(Order) ~ paste0(Class, "_sp"),
      is.na(Family) ~ paste0(Order, "_sp"),
      is.na(Genus) ~ paste0(Family, "_sp"),
      is.na(Species) ~ paste0(Genus, "_sp"),
      TRUE ~ paste0(Genus, "_", Species)
    ),
    Species_Name = str_replace_all(Species_Name, " ", "_"),
    Genus_Name = case_when(
      Genus == "BVAB2" ~ Genus,
      Genus == "Candidatus Lachnocurva" ~ "Candidatus_Lachnocurva_(BVAB1)",
      is.na(Order) ~ paste0(Class, "_gen"),
      is.na(Family) ~ paste0(Order, "_gen"),
      is.na(Genus) ~ paste0(Family, "_gen"),
      TRUE ~ Genus
    ),
    Genus_Name = str_replace_all(Genus_Name, " ", "_"),
  )
```

## Create data objects with data collapsed as species or genus level

```{r extract-count-data-from-rarefied-ps0, echo=FALSE}
#Extract ASV count table
ASV.count.table.wide <- otu_table(ps0.rare) %>% 
  as.data.frame(stringAsFactors = FALSE) %>% 
  tibble::rownames_to_column("Seq_ID")

#Pivot ASV count table to long form
ASV.count.table.long <- ASV.count.table.wide %>% 
  tidyr::pivot_longer(-Seq_ID, names_to = "ASV", values_to = "Read_Count")
```


```{r extract-16S-metadata, echo=FALSE}
#Extract 16S ps0.rare metadata
meta.16S.2017 <- sample_data(ps0.rare) %>% 
  data.frame(stringsAsFactors = FALSE) %>% 
  tibble::rownames_to_column("Seq_ID") %>%
  dplyr::rename(Visit_Code = VisitCode) %>% 
  dplyr::mutate(BV_Interpretation = str_replace(BV_Interpretation, "_", " ") %>% 
                  str_replace("Intermediate", "Inter-\nmediate"),
                BV_Interpretation = factor(BV_Interpretation,
                                           levels = c("No BV",
                                                      "Inter-\nmediate",
                                                      "BV")))

```

```{r merge-taxonomy-and-count-data-and-tax-collapse, echo=FALSE}
#Merge ASV taxonomy table with ASV count table (long form)
ASV.tax.count.table <- ASV.tax.table.names %>% 
  dplyr::left_join(ASV.count.table.long,
                   by = "ASV")

### SPECIES ###

#Collapse ASV.tax.count.table at Species_Name level
Species.tax.count.table.long <- ASV.tax.count.table %>% 
  dplyr::group_by(Species_Name, Seq_ID) %>% 
  dplyr::summarise(Read_Count = sum(Read_Count)) %>% 
  dplyr::ungroup()

#Convert Species counts to relative abundances, pseudocounts, pseudocount-based
#relative abundance, and calculate total reads and limit of detection
Species.ra.table.long <- Species.tax.count.table.long %>% 
  dplyr::group_by(Seq_ID) %>% 
  dplyr::mutate(Rel_Abund = Read_Count / sum(Read_Count),
                Pseudo_Count = case_when(Read_Count == 0 ~ Read_Count + 0.5,
                                         TRUE ~ Read_Count),
                Pseudo_Rel_Abund = Pseudo_Count / sum(Read_Count),
                Samp_Total_Reads = sum(Read_Count),
                Microb_LoD = 1 / Samp_Total_Reads) %>% 
  dplyr::ungroup() %>% 
  dplyr::arrange(desc(Rel_Abund))

#Join relevant metadata to Species relative abundance table (long)
Species.abund.CT.long <- meta.16S.2017 %>% 
  dplyr::left_join(Species.ra.table.long,
                   by = "Seq_ID")

#Calculate cohort-level stats for cumulative relative abundance and prevalence
Species.abund.CT.long.sumstats <- Species.abund.CT.long %>% 
  dplyr::group_by(Species_Name) %>% 
  dplyr::mutate(Cohort_RA = sum(Rel_Abund) / n(),
                Cohort_Prev = sum(Rel_Abund > 0) / n(),
                Prev_50 = Cohort_Prev > 0.5) %>% 
  dplyr::ungroup()


### GENUS ###

#Collapse ASV.tax.count.table at Genus_Name level
Genus.tax.count.table.long <- ASV.tax.count.table %>% 
  dplyr::group_by(Genus_Name, Seq_ID) %>% 
  dplyr::summarise(Read_Count = sum(Read_Count)) %>% 
  dplyr::ungroup()

#Convert Genus counts to relative abundance
Genus.ra.table.long <- Genus.tax.count.table.long %>% 
  dplyr::group_by(Seq_ID) %>% 
  dplyr::mutate(Rel_Abund = Read_Count / sum(Read_Count),
                Pseudo_Count = case_when(Read_Count == 0 ~ Read_Count + 0.5,
                                         TRUE ~ Read_Count),
                Pseudo_Rel_Abund = Pseudo_Count / sum(Read_Count),
                Samp_Total_Reads = sum(Read_Count),
                Microb_LoD = 1 / Samp_Total_Reads) %>% 
  dplyr::ungroup() %>% 
  dplyr::arrange(desc(Rel_Abund))

#Join relevant metadata to Species relative abundance table (long)
Genus.abund.CT.long <- meta.16S.2017 %>% 
  dplyr::left_join(Genus.ra.table.long,
                   by = "Seq_ID")

#Calculate cohort-level stats for cumulative relative abundance and prevalence
Genus.abund.CT.long.sumstats <- Genus.abund.CT.long %>% 
  dplyr::group_by(Genus_Name) %>% 
  dplyr::mutate(Cohort_RA = sum(Rel_Abund) / n(),
                Cohort_Prev = sum(Rel_Abund > 0) / n(),
                Prev_50 = Cohort_Prev > 0.5) %>% 
  dplyr::ungroup()

# Pivot Genus.ra.table.long to wide form
Genus.ra.table.wide <- Genus.ra.table.long %>%
  tidyr::pivot_wider(names_from = Genus_Name,
                     values_from = Rel_Abund)
```


```{r prepare-data-for-species-relative-abundance-plot, echo=FALSE}
#Review cohort stats for most abundant species
Species.abund.CT.long.sumstats %>% 
  dplyr::distinct(Species_Name, 
                  `Cumulative Relative Abundance Within Cohort` = Cohort_RA, 
                  `Prevalence Within Cohort` = Cohort_Prev, 
                  `>50% Prevalence` = Prev_50) %>% 
  dplyr::arrange(desc(`Cumulative Relative Abundance Within Cohort`), 
                 desc(`Prevalence Within Cohort`)) %>% 
  dplyr::mutate(Species_Name = str_replace_all(Species_Name, "_", " ")) %>% 
  head(n = 30) %>% 
  knitr::kable(caption = "Cohort abundance and prevalence of abundant species")

#Collapse key taxa to genus level for relative abundance plot
Taxon.ra.table.long <- Species.ra.table.long %>% 
  dplyr::mutate(Taxon = case_when(str_detect(Species_Name, "Gardnerella") ~ "Gardnerella",
                                  str_detect(Species_Name, "Prevotella") ~ "Prevotella",
                                  str_detect(Species_Name, "Atopobium") ~ "Atopobium",
                                  str_detect(Species_Name, "Veillonella") ~ "Veillonella",
                                  str_detect(Species_Name, "Megasphaera") ~ "Megasphaera",
                                  str_detect(Species_Name, "Peptoniphilus") ~ "Peptoniphilus",
                                  str_detect(Species_Name, "Peptostreptococcus") ~ "Peptostreptococcus",
                                  str_detect(Species_Name, "Anaerococcus") ~ "Anaerococcus",
                                  str_detect(Species_Name, "Parvimonas") ~ "Parvimonas",
                                  str_detect(Species_Name, "Dialister") ~ "Dialister",
                                  str_detect(Species_Name, "Corynebacterium") ~ "Corynebacterium",
                                  str_detect(Species_Name, "^Mycoplasma") ~ "Mycoplasma",
                                  str_detect(Species_Name, "Clostridiales") ~ "Clostridiales",
                                  str_detect(Species_Name, "Fusobacterium") ~ "Fusobacterium",
                                  str_detect(Species_Name, "Streptococcus") ~ "Streptococcus",
                                  str_detect(Species_Name, "Mobiluncus") ~ "Mobiluncus",
                                  str_detect(Species_Name, "Gemella") ~ "Gemella",
                                  TRUE ~ Species_Name)) %>% 
  dplyr::group_by(Seq_ID, Taxon) %>% 
  dplyr::summarise(Rel_Abund = sum(Rel_Abund)) %>% 
  dplyr::ungroup()

#Calculate cohort summary statistics for collapsed taxa
Taxon.ra.table.long.sumstats <- Taxon.ra.table.long %>% 
  dplyr::group_by(Taxon) %>% 
  dplyr::mutate(Cohort_RA = sum(Rel_Abund) / n(),
                Cohort_Prev = sum(Rel_Abund > 0) / n(),
                Prev_50 = Cohort_Prev > 0.5) %>% 
  dplyr::ungroup()

#Join 16S collapsed taxon summary stats to 16S metadata
Taxon.abund.CT.long.sumstats <- meta.16S.2017 %>% 
  dplyr::left_join(Taxon.ra.table.long.sumstats,
                   by = "Seq_ID")

#Review cohort stats for most abundant taxa after collapsing
Taxon.abund.CT.long.sumstats %>% 
  dplyr::distinct(Taxon, 
                  `Cumulative Relative Abundance Within Cohort` = Cohort_RA, 
                  `Prevalence Within Cohort` = Cohort_Prev, 
                  `>50% Prevalence` = Prev_50) %>% 
  dplyr::arrange(desc(`Cumulative Relative Abundance Within Cohort`), 
                 desc(`Prevalence Within Cohort`)) %>% 
  dplyr::mutate(Taxon = str_replace_all(Taxon, "_", " ")) %>% 
  head(n = 30) %>% 
  knitr::kable(caption = "Cohort abundance and prevalence of dominant collapsed taxa")

## Collapse rare taxa to "Other" (keeping top 17 taxa)
Taxcollapse.abund.CT.long <- Taxon.abund.CT.long.sumstats %>% 
  dplyr::group_by(Taxon) %>% 
  dplyr::mutate(Taxon_Collapse = case_when(Cohort_RA > 0.0045 ~ Taxon,
                                           TRUE ~ "Other")) %>% 
  dplyr::ungroup() %>% 
  dplyr::group_by(Seq_ID, Cervicotype, Taxon_Collapse, BV_Interpretation) %>% 
  dplyr::summarise(Rel_Abund = sum(Rel_Abund)) %>% 
  dplyr::ungroup() 

#Review taxa for relative abundance plot
Taxcollapse.abund.CT.long %>% 
  dplyr::group_by(Taxon_Collapse) %>% 
  dplyr::summarise(`Cumulative Relative Abundance Within Cohort` = sum(Rel_Abund) / n(),
                   `Prevalence Within Cohort` = sum(Rel_Abund > 0) / n(),
                   `>50% Prevalence` = `Prevalence Within Cohort` > 0.5) %>% 
  dplyr::ungroup() %>% 
  dplyr::arrange(desc(`Cumulative Relative Abundance Within Cohort`), 
                 desc(`Prevalence Within Cohort`)) %>% 
  dplyr::mutate(Taxon_Collapse = str_replace_all(Taxon_Collapse, "_", " "),
                Taxon_Collapse = fct_inorder(Taxon_Collapse),
                Taxon_Collapse = fct_relevel(Taxon_Collapse, "Other", after = Inf)) %>% 
  dplyr::arrange(Taxon_Collapse) %>% 
  dplyr::rename(Taxon = Taxon_Collapse) %>% 
  knitr::kable(caption = "Cohort abundance and prevalence of collapsed taxa to plot")
```


```{r arrange-sample-variables-for-species-relative-abundance-plot, echo=FALSE}
#Widen data to perform calculations needed to arrange samples for plotting
Taxcollapse.abund.CT.wide <- Taxcollapse.abund.CT.long %>% 
  dplyr::arrange(Cervicotype, desc(Rel_Abund)) %>% 
  tidyr::pivot_wider(names_from = Taxon_Collapse,
                     values_from = Rel_Abund)

#Set seed for reproducibility in next step
set.seed(100)

#Arrange samples for plotting, by descending summed 
#L. crispatus and L. iners genus relative abundance for all CTs 
#Calculate number of samples per CT, and remove unneeded final columns
Taxcollapse.abund.CT.samp.order.wide <- Taxcollapse.abund.CT.wide %>% 
  dplyr::mutate(Lact_Sum = Lactobacillus_crispatus + Lactobacillus_iners) %>% 
  dplyr::arrange(Cervicotype, desc(Lact_Sum)) %>% 
  dplyr::mutate(Seq_ID = fct_inorder(Seq_ID)) %>% 
  dplyr::group_by(Cervicotype) %>% 
  dplyr::mutate(Samps_per_CT = n()) %>% 
  dplyr::ungroup() %>% 
  dplyr::select(-Lact_Sum) %>% 
  dplyr::select(Seq_ID, Cervicotype, BV_Interpretation, starts_with("Samps"), everything())

#pivot data back to long form with factored sample IDs
Taxcollapse.abund.CT.samp.order.long <- Taxcollapse.abund.CT.samp.order.wide %>% 
  tidyr::pivot_longer(-c(Seq_ID, Cervicotype, BV_Interpretation, starts_with("Samps")),
                      names_to = "Taxon_Collapse",
                      values_to = "Rel_Abund")

#Arrange and factor taxa in desired order for plotting
Taxcollapse.abund.CT.long.sumstats.fct <- Taxcollapse.abund.CT.samp.order.long %>% 
  dplyr::group_by(Taxon_Collapse) %>%
  dplyr::mutate(Cohort_RA = sum(Rel_Abund),
                Cohort_Prev = sum(Rel_Abund > 0) / n(),
                Prev_50 = Cohort_Prev > 0.5) %>%
  dplyr::ungroup() %>% 
  dplyr::arrange(desc(Cohort_RA)) %>% 
  dplyr::mutate(Taxon_Collapse = str_replace_all(Taxon_Collapse, "_", " "),
                Taxon_Collapse = str_replace(Taxon_Collapse, "ndidatus", "."),
                Taxon_Collapse = fct_inorder(Taxon_Collapse),
                Taxon_Collapse = fct_relevel(Taxon_Collapse,
                                             "Lactobacillus crispatus",
                                             "Lactobacillus iners",
                                             "Gardnerella",
                                             "Prevotella"),
                Taxon_Collapse = fct_relevel(Taxon_Collapse,
                                             "Other",
                                             after = Inf))
```


### Relative abundance stacked bar plot.  
  

```{r taxon-relative-abundance-plot, echo=FALSE}
#Relative abundance bar plot
p.tax.ra.bar <- Taxcollapse.abund.CT.long.sumstats.fct %>%
  dplyr::mutate(Facet_labels = paste0(Cervicotype, "\n(", Samps_per_CT, ")")) %>%
  ggplot(aes(x = Seq_ID, y = Rel_Abund, fill = Taxon_Collapse)) +
  geom_col(position = position_fill(reverse = TRUE), width = 1) +
  scale_y_continuous(labels = scales::percent_format(),
                     limits = c(0, 1)) +
  scale_fill_manual(values = c("orange2",
                               "forestgreen",
                               "dodgerblue3",
                               "violetred4",
                               "#4FC3F7",
                               "burlywood2",
                               "#9FA8DA",
                               "tomato",
                               "lightskyblue3",
                               "#FDD835",
                               "maroon3",
                               "plum1",
                               "darkolivegreen3",
                               "#004D40",
                               "#CD93D8",
                               "slateblue3",
                               "snow3",
                               "grey48")) +
  labs(x = "Participant",
       y = "Relative Abundance") +
  geom_rug(aes(color = Cervicotype),
           sides = "t",
           size = 4,
           length = unit(0.01, "npc")) +
  scale_color_manual(guide = "none",
                     values = c("orange2",
                                "forestgreen",
                                "dodgerblue3",
                                "violetred4")) +
  theme_minimal() +
  guides(fill = guide_legend(keyheight = 0.65)) +
  theme(panel.spacing.x = unit(0.2, "line"),
        axis.ticks = element_blank(),
        axis.text.x = element_blank(),
        panel.grid = element_blank(),
        legend.title = element_blank(),
        legend.text = element_text(face = "italic", size = 9.5),
        axis.text.y = element_text(size = 11, color = "black"),
        axis.title = element_text(size = 12, face = "bold"),
        strip.text = element_text(size = 11)) +
  facet_grid(~Facet_labels, scales = "free_x", space = "free_x")

p.tax.ra.bar
```
  
### Relative abundance heat map  
   
```{r taxon-relative-abundance-heatmap-horizontal, echo=FALSE}
#Relative abundance bar plot
p.tax.ra.heatmap.horizontal <- Taxcollapse.abund.CT.long.sumstats.fct %>% 
  dplyr::mutate(Facet_labels = paste0(Cervicotype, "\n(n=", Samps_per_CT, ")"),
                Taxon_Collapse = fct_rev(Taxon_Collapse)) %>% 
  ggplot(aes(x = Seq_ID, y = Taxon_Collapse, fill = Rel_Abund)) +
  geom_raster() +
  scale_fill_gradient(labels = scales::percent_format(),
                      low = "white",
                      high = "red3") +
  scale_y_discrete(expand = expansion(add = c(0, 1.5))) +
  geom_rug(aes(color = Cervicotype), 
           sides = "t", 
           size = 4,
           length = unit(0.01, "npc"),
           outside = TRUE,
           show.legend = FALSE) +
  scale_color_manual(guide = "none",
                     values = c("orange2", 
                                "forestgreen", 
                                "dodgerblue3", 
                                "violetred4")) +
  labs(x = "Participant",
       fill = "Bacterial\nRelative\nAbundance") +
  theme(panel.spacing.x = unit(0, "lines"),
        panel.border = element_rect(size = 0.25),
        # strip.background = element_rect(size = 0, fill = "white"),
        axis.ticks = element_blank(),
        axis.text.x = element_blank(),
        panel.grid = element_blank(),
        legend.title = element_text(size = 8.5, hjust = 0.5, face = "bold"),
        legend.text = element_text(size = 7.5, vjust = 0),
        legend.background = element_rect(color = "black", size = 0.5),
        axis.text.y = element_text(size = 7.5, face = "italic", color = "black"),
        axis.title.x = element_text(size = 8.5, face = "bold"),
        axis.title.y = element_blank(),
        strip.text = element_text(size = 8.5, face = "bold")) +
  guides(fill = guide_colorbar(barheight = 4)) +
  facet_grid(~Facet_labels, scales = "free_x", space = "free_x")
  
p.tax.ra.heatmap.horizontal
```
  
  
### Panel B: Relative abundance heat map vertical  
  

```{r taxon-relative-abundance-heatmap-vertical, echo=FALSE}
#Relative abundance bar plot
p.tax.ra.heatmap.vertical <- Taxcollapse.abund.CT.long.sumstats.fct %>% 
  dplyr::mutate(Facet_labels = paste0(Cervicotype, "\n(n=", Samps_per_CT, ")")) %>% 
  ggplot(aes(y = Seq_ID, x = Taxon_Collapse, fill = Rel_Abund)) +
  geom_raster() +
  scale_fill_gradient(labels = scales::percent_format(),
                      low = "white",
                      high = "red3") +
  scale_y_discrete(expand = expansion(add = c(0, 1.5))) +
  geom_rug(aes(color = Cervicotype), 
           sides = "r", 
           size = 4,
           length = unit(0.01, "npc"),
           outside = TRUE,
           show.legend = FALSE) +
  scale_color_manual(guide = "none",
                     values = c("orange2", 
                                "forestgreen", 
                                "dodgerblue3", 
                                "violetred4")) +
  labs(y = "Participant",
       fill = "Bacterial\nRelative\nAbundance") +
  theme(panel.spacing.y = unit(0, "lines"),
        panel.border = element_rect(size = 0.25),
        # strip.background = element_rect(size = 0, fill = "white"),
        axis.ticks = element_blank(),
        axis.text.y = element_blank(),
        panel.grid = element_blank(),
        legend.position = "left",
        legend.title = element_text(size = 8.5, hjust = 0.5, face = "bold"),
        legend.text = element_text(size = 7.5, vjust = 0),
        legend.background = element_rect(color = "black", size = 0.5),
        axis.text.x = element_text(size = 7.5, face = "italic", 
                                   angle = 35, hjust = 1, color = "black"),
        axis.title.y = element_text(size = 8.5, face = "bold"),
        axis.title.x = element_blank(),
        strip.text.y = element_text(size = 8.5, angle = 0)) +
  guides(fill = guide_colorbar(barheight = 4)) +
  facet_grid(Facet_labels~., scales = "free_y", space = "free_y")
  
p.tax.ra.heatmap.vertical
```
  
### Load Metabolomics Data 

```{r load-data, echo=FALSE}
#load in cysteine metabolomics data 
f_cysteine_data.unprocessed <- "Data/FRESH_data/cysteine_metabolomics_data_2020_09_03.txt"
f_cysteine_data.unprocessed

cysteine_data.unprocessed <- read_tsv(f_cysteine_data.unprocessed)
```

### Process Metabolomics Data 

[a couple lines about processing steps and citing Gabudza source]

filter out biochemicals with greater than 70% missingness across samples, adjust raw relative concentration value by volume of sample extracted, normalize volume-adjsuted values by median, then impute missing values with 0.5 * minimum of median-centered volume-adjusted values. intial filter step not performed becaue 


```{r process-metabolomics-data, echo=FALSE}
#exclude sample excluded from 16S date due to low read count
cysteine_data.unprocessed.low16s.excluded  <- cysteine_data.unprocessed %>%
  dplyr::semi_join(meta.16S.2017, by = c("Subject_ID", 
                                         "Collection_Date", 
                                         "Visit_Code"))

# Calculate volume-adjusted concentrations (not all samples had equal input volumes)
cysteine_data.voladjusted <- cysteine_data.unprocessed.low16s.excluded %>%
  tidyr::pivot_longer( c(cysteine:`glutathione, reduced (GSH)`),   
                       names_to = "Biochemical", 
                       values_to = "Concentration") %>%
  dplyr::group_by(Biochemical) %>%
  dplyr::mutate(Concentration_voladj = Concentration / Volume_Extracted) %>%
  dplyr::ungroup()

# Calculate median-centered concentration for each metabolite (after excluding missing data) 
# and inferred limit of detection based on minimum measured value
cysteine_data.centered <- cysteine_data.voladjusted %>%
  dplyr::group_by(Biochemical) %>%
  dplyr::mutate(Concentration_norm = (Concentration_voladj / median(Concentration_voladj, 
                                                                    na.rm = TRUE))) %>%
  dplyr::mutate(metab_LoD = min(Concentration_norm, na.rm = TRUE)) %>%
  dplyr::ungroup() 

# NA values represent samples below the limit of detection. Impute the missing values
# as half of minimum of the calculated limit of detection 
cysteine_data.imputed <- cysteine_data.centered %>%
  dplyr::group_by(Biochemical) %>%
  dplyr::mutate(Conc_norm_minimp = dplyr::coalesce(Concentration_norm, 
                                                   0.5 * min(Concentration_norm, 
                                                             na.rm = TRUE))) %>%
  dplyr::ungroup() 

#retain only columns needed for analysis, including metab_lod and the centered 
#concentrations with imputed values, keep in long form
cysteine_data.processed.long <- cysteine_data.imputed %>%
  dplyr::select(Group_Metab:Visit_Code, Biochemical, 
                Concentration = Conc_norm_minimp, metab_LoD)

#retain only columns needed for analysis, pivot to wide form
cysteine_data.processed <- cysteine_data.imputed %>%
  dplyr::select(Group_Metab:Visit_Code, Biochemical, Conc_norm_minimp, metab_LoD) %>%
  tidyr::pivot_wider(names_from = Biochemical, values_from = c(Conc_norm_minimp, metab_LoD)) %>% 
  dplyr::rename_with(function(x) {str_remove(x, "Conc_norm_minimp_")}, starts_with("Conc_norm_minimp_")) %>% 
  dplyr::rename_with(function(x) {str_remove(x, "metab_LoD_") %>% paste0("_LoD")},
                     starts_with("metab_LoD_"))
```  
  
### Supplementary table: For each analyte, percent of samples with missing data, and result of test for normality
    

```{r missingness-table, echo=FALSE}
#generate table describing missingness for each metabolite
metabolite_missingness_table <- cysteine_data.unprocessed %>%
  tidyr::pivot_longer( c(cysteine:`glutathione, reduced (GSH)`), names_to = "Biochemical", values_to = "Concentration") %>%
  dplyr::group_by(Biochemical) %>%
  dplyr::mutate(`Percent Missing` = (sum(is.na(Concentration))/ n()) * 100 ) %>%
  dplyr::ungroup() %>% 
  dplyr::distinct(Biochemical, `Percent Missing`) %>%
  knitr::kable(caption = "Fraction Missing for Each Analyte")

metabolite_missingness_table
```
  
### Check cysteine concentration (log10) normality  
  

```{r normality-cys, echo=FALSE}
#test normality of cysteine 
ggdensity(log10(cysteine_data.processed$cysteine))
ggqqplot(log10(cysteine_data.processed$cysteine))
shapiro.test(log10(cysteine_data.processed$cysteine))
```
  
### Check cysteinylglycine concentration (log10) normality  
  

```{r normality-cysgly, echo=FALSE}
#test normality of cysteinylglycine 
ggdensity(log10(cysteine_data.processed$cysteinylglycine))
ggqqplot(log10(cysteine_data.processed$cysteinylglycine))
shapiro.test(log10(cysteine_data.processed$cysteinylglycine))
```
  
### Check GSH concentration (log10) normality  
  

```{r normality-gsh, echo=FALSE}
#test normality of GSH 
ggdensity(log10(cysteine_data.processed$`glutathione, reduced (GSH)`))
ggqqplot(log10(cysteine_data.processed$`glutathione, reduced (GSH)`))
shapiro.test(log10(cysteine_data.processed$`glutathione, reduced (GSH)`))
```
  
  
### Nugent plots  
  
```{r review-samples-with-Nugent-testing-by-testing-regimen, echo=FALSE}

meta.16S.2017 %>% 
  dplyr::mutate(BV_Testing_Regimen = case_when(Collection_Date <= as.Date("2017-09-14") ~ "Tested Under Sporadic Nugent Testing Protocol",
                                               TRUE ~ "Tested Under Universal Nugent Testing Protocol")) %>% 
  dplyr::count(BV_Interpretation, BV_Testing_Regimen, name = "Num_Samps") %>% 
  tidyr::pivot_wider(names_from = BV_Testing_Regimen,
                     values_from = Num_Samps) %>% 
  knitr::kable(caption = "Number of samples with Nugent scoring by Nugent testing regimen protocol")


```

```{r merge metabolomics-data-16s-BV-data, echo=FALSE}
#merge metabolomics with 16S 
cysteine_metab_16s <- meta.16S.2017 %>%
  dplyr::left_join(cysteine_data.processed, by = c("Subject_ID", 
                                                   "Collection_Date", 
                                                   "Visit_Code"))

#merge metabolomics.long with 16S data
cysteine_metab_16s.long <- meta.16S.2017 %>%
  dplyr::left_join(cysteine_data.processed.long, by = c("Subject_ID", 
                                                   "Collection_Date", 
                                                   "Visit_Code"))
```
  
  
### Panel A: Cysteine concentrations by BV box plot
  
For analysis of metabolite concentrations in BV, in order to remove any potential bias from samples collected under the sporadic Nugent testing protocol, restricting BV-related analysis to samples collected under the universal Nugent scoring protocol.     
  

```{r filter-cysteine-metab-dfs-to-Universal-Nugent-samples, echo=FALSE}
#Filtering data to samples collected under universal Nugent testing protocol
cysteine_metab_16s_univ_Nugent <- cysteine_metab_16s %>%
  dplyr::filter(Collection_Date > as.Date("2017-09-14"))

cysteine_metab_16s.long_univ_Nugent <- cysteine_metab_16s.long %>%
  dplyr::filter(Collection_Date > as.Date("2017-09-14"))
```


```{r cys-BV-box-plot-parameterized, echo=FALSE}
#create box plot - paramterized 
p.cysteine.BV.boxplot <- cysteine_metab_16s_univ_Nugent %>% 
  dplyr::rename(`BV Status` = BV_Interpretation) %>% 
  ggplot(aes(x = `BV Status`, y = cysteine)) +
  geom_boxplot(coef = 5,
               fatten = 1) +
  geom_jitter(aes(color = `BV Status`),
              width = 0.1, size = 1.25, shape = 18,
              show.legend = FALSE) +
  scale_y_log10(labels = scales::trans_format("log10", scales::math_format(10^.x)),
                expand = expansion(mult = c(0.05, 0.09))) +
  scale_color_manual(values = c(hsv(h = 0.75, s = 1, v = 1),
                                hsv(h = 0.48, s = 1, v = 0.45),
                                hsv(h = 0, s = 1, v = 1))) +
  annotation_logticks(sides = "l",
                      long = unit(0.15, "cm"),
                      mid = unit(0.075, "cm"),
                      short = unit(0.037, "cm")) +
  stat_compare_means(method = "anova",
                     label.y.npc = 0.02,
                     label.x.npc = 0.3,
                     label.sep = "\n",
                     size = 3,
                     hjust = 0.5) +
  labs(y = "Relative Cys Conc.") +
  theme(axis.title = element_text(size = 8.5, face = "bold"),
        axis.text.x = element_text(size = 7.5, color = "black"),
        axis.text.y = element_text(size = 7.5, color = "black"),
        legend.text = element_text(size = 7.5),
        legend.title = element_text(size = 8.5, hjust = 0.5, face = "bold"),
        panel.grid = element_blank())

p.cysteine.BV.boxplot
```


```{r cys-BV-stats, echo=FALSE}
#anova
cysteine.BV.anova <- aov(log10(cysteine) ~ BV_Interpretation,
                         data = cysteine_metab_16s_univ_Nugent)
summary(cysteine.BV.anova)

#post-hoc Tukey test
cysteine.BV.tukey <- TukeyHSD(cysteine.BV.anova)
cysteine.BV.tukey

#Kruskal-Wallis
cysteine.BV.kruskal <- kruskal.test(cysteine ~ BV_Interpretation,
                                    data = cysteine_metab_16s_univ_Nugent)
cysteine.BV.kruskal

#Dunn's test
cysteine.BV.dunns <- DescTools::DunnTest(cysteine ~ BV_Interpretation,
                                         data = cysteine_metab_16s_univ_Nugent,
                                         method = "bonferroni")
cysteine.BV.dunns
```
  
Use of 1-way ANOVA on log-transformed concentrations with post-hoc Tukey test vs Kruskal-Wallis test with post-hoc Dunn's Test (Bonferroni method) yield equivalent results for Cys. Will use the parametric tests for Cys because its concentrations are normally distributed. Will use non-parametric for GSH and Cys-Gly since they are not.  
  

```{r plot-cysteine-BV-tukey-comparisons, echo=FALSE}
#Create dataframe of post-hoc Tukey test for cysteine cervicotype analysis for plotting
cysteine.BV.Tukey.df <- cysteine.BV.tukey$BV_Interpretation %>% 
  as.data.frame() %>% 
  dplyr::select(P_adjust = `p adj`, everything()) %>% 
  tibble::rownames_to_column("Groups") %>%
  dplyr::mutate(group1 = case_when(str_detect(Groups,
                                              "^Inter-\nmediate") ~ "Inter-\nmediate",
                                   TRUE ~ str_extract(Groups, "[^-]+")),
                group2 = case_when(str_detect(Groups,
                                              "Inter-\nmediate$") ~ "Inter-\nmediate",
                                   TRUE ~ str_remove(Groups, "^Inter-\nmediate-")%>% 
                                     str_replace(".+-(.+)", "\\1"))) %>% 
  dplyr::mutate(Sigsymbol_adj = case_when(P_adjust <= 0.0001 ~ "****",
                                          P_adjust <= 0.001 ~ "***",
                                          P_adjust <= 0.01 ~ "**",
                                          P_adjust <= 0.05 ~ "*",
                                          TRUE ~ "n.s."))

#Plote all significant pairwise post-hoc comparisons
p.cysteine.BV.boxplot.Tukey <- p.cysteine.BV.boxplot +
  stat_pvalue_manual(cysteine.BV.Tukey.df %>% 
                       dplyr::filter(P_adjust <= 0.05),
                     label = "Sigsymbol_adj",
                     y.position = 1.7,
                     step.increase = -0.08,
                     tip.length = 0,
                     size = 3.5)

p.cysteine.BV.boxplot.Tukey
```
  
### Supplementary Figure 4A and 4B: CysGly and GSH concentration box plots vs Nugent 
  
Formatted as for cysteine plot above.

Statistics: as above.  


```{r cysgly-BV-box-plot-nonparameterized, echo=FALSE}

#use long form 
p.cysgly.BV.boxplot <- cysteine_metab_16s.long_univ_Nugent %>%
  dplyr::filter(Biochemical == "cysteinylglycine") %>%
  dplyr::rename(`BV Status` = BV_Interpretation) %>% 
  ggplot(aes(x = `BV Status`, y = Concentration)) +
  geom_boxplot(coef = 5,
               fatten = 1) +
  geom_jitter(aes(color = `BV Status`),
              width = 0.1, size = 1.25, shape = 18,
              show.legend = FALSE) +
  scale_y_log10(labels = scales::trans_format("log10", scales::math_format(10^.x)),
                expand = expansion(mult = c(0.05, 0.17))) +
  scale_color_manual(values = c(hsv(h = 0.75, s = 1, v = 1),
                                hsv(h = 0.48, s = 1, v = 0.45),
                                hsv(h = 0, s = 1, v = 1))) +
  annotation_logticks(sides = "l",
                      long = unit(0.15, "cm"),
                      mid = unit(0.075, "cm"),
                      short = unit(0.037, "cm")) +
  stat_compare_means(method = "kruskal",
                     label.y.npc = "top",
                     label.x.npc = "middle",
                     label.sep = ", ",
                     size = 3,
                     hjust = 0.5) +
  annotate("text",
           label = "L.D.",
           x = 1,
           y = 0.025,
           color = "red",
           size = 2.8) +
  labs(y = "Relative Cys-Gly Conc.") +
  theme(axis.title = element_text(size = 8.5, face = "bold"),
        axis.text.x = element_text(size = 7.5, color = "black"),
        axis.text.y = element_text(size = 7.5, color = "black"),
        legend.text = element_text(size = 7.5),
        legend.title = element_text(size = 8.5, hjust = 0.5, face = "bold"),
        panel.grid = element_blank()) +
  geom_hline(aes(yintercept = metab_LoD), linetype = "dotted",
           color = "red")

p.cysgly.BV.boxplot
```


  
```{r cysgly-BV-stats-nonparametric, echo=FALSE}
#Kruskal-Wallis
cysgly.BV.kruskal <- kruskal.test(cysteinylglycine ~ BV_Interpretation,
                                  data = cysteine_metab_16s_univ_Nugent)
cysgly.BV.kruskal

#Dunns
cysgly.BV.dunns <- DescTools::DunnTest(cysteinylglycine ~ BV_Interpretation,
                                       data = cysteine_metab_16s_univ_Nugent,
                                       method = "bonferroni")
cysgly.BV.dunns
```
  
```{r plot-CysGly-BV-Dunns-comparisons, echo=FALSE}
#Create dataframe of post-hoc Tukey test for cysteinylglycine cervicotype analysis for plotting
cysgly.BV.dunns.df <- cysgly.BV.dunns[[1]] %>% 
  as.data.frame() %>% 
  dplyr::select(P_adjust = `pval`, everything()) %>% 
  tibble::rownames_to_column("Groups") %>%
  dplyr::mutate(group1 = case_when(str_detect(Groups,
                                              "^Inter-\nmediate") ~ "Inter-\nmediate",
                                   TRUE ~ str_extract(Groups, "[^-]+")),
                group2 = case_when(str_detect(Groups,
                                              "Inter-\nmediate$") ~ "Inter-\nmediate",
                                   TRUE ~ str_remove(Groups, "^Inter-\nmediate-")%>% 
                                     str_replace(".+-(.+)", "\\1"))) %>% 
  dplyr::mutate(Sigsymbol_adj = case_when(P_adjust <= 0.0001 ~ "****",
                                          P_adjust <= 0.001 ~ "***",
                                          P_adjust <= 0.01 ~ "**",
                                          P_adjust <= 0.05 ~ "*",
                                          TRUE ~ "n.s."))
  
#Plot all significant pairwise comparisons
p.cysgly.BV.boxplot.Dunn <- p.cysgly.BV.boxplot + 
  stat_pvalue_manual(cysgly.BV.dunns.df %>% 
                       dplyr::filter(P_adjust <= 0.05),
                     label = "Sigsymbol_adj",
                     y.position = 1.1,
                     step.increase = -0.07,
                     tip.length = 0,
                     size = 3.5)

p.cysgly.BV.boxplot.Dunn
```

```{r gsh-BV-box-plot-nonparametric, echo=FALSE}

p.gsh.BV.boxplot <- cysteine_metab_16s.long_univ_Nugent %>%
  dplyr::filter(Biochemical == "glutathione, reduced (GSH)") %>%
  dplyr::rename(`BV Status` = BV_Interpretation) %>% 
  ggplot(aes(x = `BV Status`, y = Concentration)) +
  geom_boxplot(coef = 5,
               fatten = 1) +
  geom_jitter(aes(color = `BV Status`),
              width = 0.1, size = 1.25, shape = 18,
              show.legend = FALSE) +
  scale_y_log10(labels = scales::trans_format("log10", scales::math_format(10^.x)),
                expand = expansion(mult = c(0.05, 0.17))) +
  scale_color_manual(values = c(hsv(h = 0.75, s = 1, v = 1),
                                hsv(h = 0.48, s = 1, v = 0.45),
                                hsv(h = 0, s = 1, v = 1))) +
  annotation_logticks(sides = "l",
                      long = unit(0.15, "cm"),
                      mid = unit(0.075, "cm"),
                      short = unit(0.037, "cm")) +
  stat_compare_means(method = "kruskal.test",
                     label.y.npc = "top",
                     label.x.npc = "middle",
                     label.sep = ", ",
                     size = 3,
                     hjust = 0.5) +
  labs(y = "Relative GSH Conc.") +
  annotate("text",
           label = "L.D.",
           x = 1,
           y = 0.007,
           color = "red",
           size = 2.8) +
  theme(axis.title = element_text(size = 8.5, face = "bold"),
        axis.text.x = element_text(size = 7.5, color = "black"),
        axis.text.y = element_text(size = 7.5, color = "black"),
        legend.text = element_text(size = 7.5),
        legend.title = element_text(size = 8.5, hjust = 0.5, face = "bold"),
        panel.grid = element_blank()) +
  geom_hline(aes(yintercept = metab_LoD), linetype = "dotted",
           color = "red")

p.gsh.BV.boxplot
```

```{r gsh-BV-stats-nonparameterized, echo=FALSE}

#Kruskal Wallis
gsh.BV.kruskal <- kruskal.test(`glutathione, reduced (GSH)` ~ BV_Interpretation,
                               data = cysteine_metab_16s_univ_Nugent)
gsh.BV.kruskal

#Dunns
gsh.BV.dunns <-  DescTools::DunnTest(`glutathione, reduced (GSH)` ~ BV_Interpretation,
                                     data = cysteine_metab_16s_univ_Nugent,
                                     method = "bonferroni")
gsh.BV.dunns
```

```{r plot-GSH-BV-Dunns-comparisons, echo=FALSE}
#Create dataframe of post-hoc Tukey test for cysteinylglycine cervicotype analysis for plotting
gsh.BV.dunns.df <- gsh.BV.dunns[[1]] %>% 
  as.data.frame() %>% 
  dplyr::select(P_adjust = `pval`, everything()) %>% 
  tibble::rownames_to_column("Groups") %>%
  dplyr::mutate(group1 = case_when(str_detect(Groups,
                                              "^Inter-\nmediate") ~ "Inter-\nmediate",
                                   TRUE ~ str_extract(Groups, "[^-]+")),
                group2 = case_when(str_detect(Groups,
                                              "Inter-\nmediate$") ~ "Inter-\nmediate",
                                   TRUE ~ str_remove(Groups, "^Inter-\nmediate-")%>% 
                                     str_replace(".+-(.+)", "\\1"))) %>% 
  dplyr::mutate(Sigsymbol_adj = case_when(P_adjust <= 0.0001 ~ "****",
                                          P_adjust <= 0.001 ~ "***",
                                          P_adjust <= 0.01 ~ "**",
                                          P_adjust <= 0.05 ~ "*",
                                          TRUE ~ "n.s."))
  
#Plot all significant pairwise comparisons
p.gsh.BV.boxplot.Dunn <- p.gsh.BV.boxplot + 
  stat_pvalue_manual(gsh.BV.dunns.df %>% 
                       dplyr::filter(P_adjust <= 0.05),
                     label = "Sigsymbol_adj",
                     y.position = 1.5,
                     step.increase = -0.07,
                     tip.length = 0,
                     size = 3.5)

p.gsh.BV.boxplot.Dunn
```
  
### Supplementary Figure 3A: Nugent-Cervicotype relationship  
  
```{r review-BV-CT-interaction, echo=FALSE}
#Summarize number of samples by BV status and CT
BV_CT_sum_non_missing <- cysteine_metab_16s_univ_Nugent %>% 
  dplyr::count(Cervicotype, BV_Interpretation, name = "# of\nSamples") %>% 
  tidyr::complete(Cervicotype, BV_Interpretation,
                  fill = list(`# of\nSamples` = 0)) 

p.BV.CT.tally.heatmap <- BV_CT_sum_non_missing %>% 
  dplyr::mutate(`BV Status` = str_remove(BV_Interpretation, "-\n")) %>% 
  ggplot(aes(x = Cervicotype, y = `BV Status`, fill = `# of\nSamples`)) +
  geom_raster() +
  scale_fill_gradient(low = "white",
                      high = "blue3") +
  geom_label(aes(label = `# of\nSamples`),
             fill = "white",
             alpha = 0.75,
             size = 3) +
  theme(
        panel.grid = element_blank(),
        legend.title = element_text(size = 8.5, hjust = 0.5, face = "bold"),
        legend.text = element_text(size = 7.5, vjust = 0),
        legend.background = element_rect(color = "black", size = 0.5),
        axis.text = element_text(size = 7.5, color = "black"),
        axis.title = element_text(size = 8.5, face = "bold")) 

p.BV.CT.tally.heatmap
```
  
Calculate Fisher's Exact Test to confirm non-random association of CT vs BV status.  
  
```{r fisher-exact-test-BV-CT-interaction, echo=FALSE}
#Fisher test to confirm non-random relationship between CT and Nugent results
BV_CT_sum_non_missing %>% 
  tidyr::pivot_wider(values_from = `# of\nSamples`,
                     names_from = Cervicotype) %>% 
  tibble::column_to_rownames("BV_Interpretation") %>% 
  as.matrix() %>% 
  fisher.test()
```

  
### Panel C: Cysteine concentration box plot with cysteine concentration vs cervicotype box plot
 


```{r cys-CT-box-plot-parameterized, echo=FALSE}
#create box plot - paramterized 
p.cysteine.CT.boxplot <- cysteine_metab_16s %>%
  ggplot(aes(x = Cervicotype, y = cysteine)) +
  geom_boxplot(coef = 5,
               fatten = 1) +
  geom_jitter(aes(color = Cervicotype),
              width = 0.1, size = 1.25, shape = 18,
              show.legend = FALSE) +
  scale_y_log10(labels = scales::trans_format("log10", scales::math_format(10^.x)),
                expand = expansion(mult = c(0.05, 0.09))) +
  scale_color_manual(values = c("orange2",
                                "forestgreen",
                                "dodgerblue3",
                                "violetred4")) +
  annotation_logticks(sides = "l",
                      long = unit(0.15, "cm"),
                      mid = unit(0.075, "cm"),
                      short = unit(0.037, "cm")) +
  stat_compare_means(method = "anova",
                     label.y.npc = 0.02,
                     label.x.npc = 0.3,
                     label.sep = "\n",
                     size = 3,
                     hjust = 0.5) +
  labs(y = "Relative Cys Conc.") +
  theme(axis.title = element_text(size = 8.5, face = "bold"),
        axis.text = element_text(size = 7.5, color = "black"),
        legend.text = element_text(size = 7.5),
        legend.title = element_text(size = 8.5, hjust = 0.5, face = "bold"),
        panel.grid = element_blank())

p.cysteine.CT.boxplot
```


```{r cys-CT-stats, echo=FALSE}
#anova
cysteine.CT.anova <- summary(aov(lm(log10(cysteine_metab_16s$cysteine) ~ cysteine_metab_16s$Cervicotype)))
cysteine.CT.anova

#post-hoc Tukey test
cysteine.CT.tukey <- TukeyHSD(aov(lm(log10(cysteine_metab_16s$cysteine) ~ cysteine_metab_16s$Cervicotype)))
cysteine.CT.tukey

#Kruskal-Wallis
cysteine.CT.kruskal <- kruskal.test(log10(cysteine_metab_16s$cysteine), as.factor(cysteine_metab_16s$Cervicotype))
cysteine.CT.kruskal

#Dunn's test
cysteine.CT.dunns <- DescTools::DunnTest(log10(cysteine_metab_16s$cysteine), 
                                         as.factor(cysteine_metab_16s$Cervicotype),
                                         method = "bonferroni")
cysteine.CT.dunns
```

```{r plot-cysteine-CT-tukey-comparisons, echo=FALSE}
#Create dataframe of post-hoc Tukey test for cysteine cervicotype analysis for plotting
cysteine.CT.Tukey.df <- cysteine.CT.tukey$`cysteine_metab_16s$Cervicotype` %>% 
  as.data.frame() %>% 
  dplyr::select(P_adjust = `p adj`, everything()) %>% 
  tibble::rownames_to_column("Groups") %>%
  tidyr::separate(col = Groups,
                  into = c("group1", "group2")) %>% 
  dplyr::mutate(Sigsymbol_adj = case_when(P_adjust <= 0.0001 ~ "****",
                                          P_adjust <= 0.001 ~ "***",
                                          P_adjust <= 0.01 ~ "**",
                                          P_adjust <= 0.05 ~ "*",
                                          TRUE ~ "n.s."))

#Plote all significant pairwise post-hoc comparisons
p.cysteine.CT.boxplot.Tukey <- p.cysteine.CT.boxplot +
  stat_pvalue_manual(cysteine.CT.Tukey.df %>% 
                       dplyr::filter(P_adjust <= 0.05),
                     label = "Sigsymbol_adj",
                     y.position = 2.5,
                     step.increase = -0.08,
                     tip.length = 0,
                     size = 3.5)

p.cysteine.CT.boxplot.Tukey
```

  


### Supplementary Figures 4C and 4D: CysGly and GSH concentration box plots vs cervicotype 
  
Formatted as for cysteine plot above.

Statistics: as above.  


```{r cysgly-ct-box-plot-nonparameterized, echo=FALSE}

#use long form 
p.cysgly.CT.boxplot <- cysteine_metab_16s.long %>%
  dplyr::filter(Biochemical == "cysteinylglycine") %>%
  ggplot(aes(x = Cervicotype, y = Concentration)) +
  geom_boxplot(coef = 5,
               fatten = 1) +
  geom_jitter(aes(color = Cervicotype),
              width = 0.1, size = 1.25, shape = 18,
              show.legend = FALSE) +
  scale_y_log10(labels = scales::trans_format("log10", scales::math_format(10^.x)),
                expand = expansion(mult = c(0.05, 0.17))) +
  scale_color_manual(values = c("orange2",
                                "forestgreen",
                                "dodgerblue3",
                                "violetred4")) +
  annotation_logticks(sides = "l",
                      long = unit(0.15, "cm"),
                      mid = unit(0.075, "cm"),
                      short = unit(0.037, "cm")) +
  stat_compare_means(method = "kruskal",
                     label.y.npc = "top",
                     label.x.npc = "middle",
                     label.sep = ", ",
                     size = 3,
                     hjust = 0.5) +
  annotate("text",
           label = "L.D.",
           x = 1,
           y = 0.025,
           color = "red",
           size = 2.8) +
  labs(y = "Relative Cys-Gly Conc.") +
  theme(axis.title = element_text(size = 8.5, face = "bold"),
        axis.text = element_text(size = 7.5, color = "black"),
        legend.text = element_text(size = 7.5),
        legend.title = element_text(size = 8.5, hjust = 0.5, face = "bold"),
        panel.grid = element_blank()) +
  geom_hline(aes(yintercept = metab_LoD), linetype = "dotted",
           color = "red")

p.cysgly.CT.boxplot
```


  
```{r cysgly-ct-stats-nonparametric, echo=FALSE}
#Kruskal-Wallis
cysgly.CT.kruskal <- kruskal.test(cysteine_metab_16s$cysteinylglycine, as.factor(cysteine_metab_16s$Cervicotype))
cysgly.CT.kruskal

#Dunns
cysgly.CT.dunns <- DescTools::DunnTest(cysteine_metab_16s$cysteinylglycine, 
                                       as.factor(cysteine_metab_16s$Cervicotype),
                                       method = "bonferroni")
cysgly.CT.dunns
```
  
```{r plot-CysGly-CT-Dunns-comparisons, echo=FALSE}
#Create dataframe of post-hoc Tukey test for cysteinylglycine cervicotype analysis for plotting
cysgly.CT.dunns.df <- cysgly.CT.dunns[[1]] %>% 
  as.data.frame() %>% 
  dplyr::select(P_adjust = `pval`, everything()) %>% 
  tibble::rownames_to_column("Groups") %>%
  tidyr::separate(col = Groups,
                  into = c("group1", "group2")) %>% 
  dplyr::mutate(Sigsymbol_adj = case_when(P_adjust <= 0.0001 ~ "****",
                                          P_adjust <= 0.001 ~ "***",
                                          P_adjust <= 0.01 ~ "**",
                                          P_adjust <= 0.05 ~ "*",
                                          TRUE ~ "n.s."))
  
#Plot all significant pairwise comparisons
p.cysgly.CT.boxplot.Dunn <- p.cysgly.CT.boxplot + 
  stat_pvalue_manual(cysgly.CT.dunns.df %>% 
                       dplyr::filter(P_adjust <= 0.05),
                     label = "Sigsymbol_adj",
                     y.position = 1.7,
                     step.increase = -0.07,
                     tip.length = 0,
                     size = 3.5)

p.cysgly.CT.boxplot.Dunn
```

```{r gsh-CT-box-plot-nonparametric, echo=FALSE}

p.gsh.CT.boxplot <- cysteine_metab_16s.long %>%
  dplyr::filter(Biochemical == "glutathione, reduced (GSH)") %>%
  ggplot(aes(x = Cervicotype, y = Concentration)) +
  geom_boxplot(coef = 5,
               fatten = 1) +
  geom_jitter(aes(color = Cervicotype),
              width = 0.1, size = 1.25, shape = 18,
              show.legend = FALSE) +
  scale_y_log10(labels = scales::trans_format("log10", scales::math_format(10^.x)),
                expand = expansion(mult = c(0.05, 0.17))) +
  scale_color_manual(values = c("orange2",
                                "forestgreen",
                                "dodgerblue3",
                                "violetred4")) +
  annotation_logticks(sides = "l",
                      long = unit(0.15, "cm"),
                      mid = unit(0.075, "cm"),
                      short = unit(0.037, "cm")) +
  stat_compare_means(method = "kruskal.test",
                     label.y.npc = "top",
                     label.x.npc = "middle",
                     label.sep = ", ",
                     size = 3,
                     hjust = 0.5) +
  labs(y = "Relative GSH Conc.") +
  annotate("text",
           label = "L.D.",
           x = 1,
           y = 0.007,
           color = "red",
           size = 2.8) +
  theme(axis.title = element_text(size = 8.5, face = "bold"),
        axis.text = element_text(size = 7.5, color = "black"),
        panel.grid = element_blank()) +
  geom_hline(aes(yintercept = metab_LoD), linetype = "dotted",
           color = "red")

p.gsh.CT.boxplot
```

```{r gsh-ct-stats-nonparameterized, echo=FALSE}

#Kruskal Wallis
gsh.CT.kruskal <- kruskal.test(cysteine_metab_16s$`glutathione, reduced (GSH)`, 
                               as.factor(cysteine_metab_16s$Cervicotype))
gsh.CT.kruskal

#Dunns
gsh.CT.dunns <-  DescTools::DunnTest(cysteine_metab_16s$`glutathione, reduced (GSH)`, 
                                     as.factor(cysteine_metab_16s$Cervicotype),
                                     method = "bonferroni")
gsh.CT.dunns
```

```{r plot-GSH-CT-Dunns-comparisons, echo=FALSE}
#Create dataframe of post-hoc Tukey test for cysteinylglycine cervicotype analysis for plotting
gsh.CT.dunns.df <- gsh.CT.dunns[[1]] %>% 
  as.data.frame() %>% 
  dplyr::select(P_adjust = `pval`, everything()) %>% 
  tibble::rownames_to_column("Groups") %>%
  tidyr::separate(col = Groups,
                  into = c("group1", "group2")) %>% 
  dplyr::mutate(Sigsymbol_adj = case_when(P_adjust <= 0.0001 ~ "****",
                                          P_adjust <= 0.001 ~ "***",
                                          P_adjust <= 0.01 ~ "**",
                                          P_adjust <= 0.05 ~ "*",
                                          TRUE ~ "n.s."))
  
#Plot all significant pairwise comparisons
p.gsh.CT.boxplot.Dunn <- p.gsh.CT.boxplot + 
  stat_pvalue_manual(gsh.CT.dunns.df %>% 
                       dplyr::filter(P_adjust <= 0.05),
                     label = "Sigsymbol_adj",
                     y.position = 1.82,
                     step.increase = -0.07,
                     tip.length = 0,
                     size = 3.5)

p.gsh.CT.boxplot.Dunn
```
  
  
### Panel D: Point plots of correlation between log-transformed relative abundance (with pseudocount) of Lactobacillus, Gardnerella, and Prevotella (all Genus level) against log-transformed cysteine concentration  
  

```{r correlation-plots-of-selected-genera-cysteine, echo=FALSE}
#Horizontal
p.cysteine.pseudocount.genuslevel.correlationplot.horizontal <- Genus.abund.CT.long.sumstats %>% 
  dplyr::filter(Genus_Name %in% c("Lactobacillus",
                                  "Gardnerella",
                                  "Prevotella")) %>%
  dplyr::mutate(Genus_Name = factor(Genus_Name,
                                    levels = c("Lactobacillus",
                                  "Gardnerella",
                                  "Prevotella"))) %>%
  dplyr::left_join(cysteine_data.processed, by = c("Subject_ID", "Collection_Date", "Visit_Code")) %>%
  ggplot(aes(x = Pseudo_Rel_Abund,
             y = cysteine # (Relative/imputed),
             )) +
  geom_vline(aes(xintercept = Microb_LoD), 
               linetype = "dotted",
               color = "red") +
  geom_smooth( method='lm' , se = TRUE) +
  geom_point(size = 1, shape = 18) +
  scale_x_log10(labels = scales::trans_format("log10", scales::math_format(10^.x)),
                expand = expansion(mult = c(0.075, 0.05))) +
  scale_y_log10(labels = scales::trans_format("log10", scales::math_format(10^.x)),
                expand = expansion(mult = c(0.1, 0.05))) +
  annotation_logticks(sides = "bl",
                      long = unit(0.15, "cm"),
                      mid = unit(0.075, "cm"),
                      short = unit(0.037, "cm")) +
  stat_cor(method = "spearman",
           aes(label = ..r.label..),
           cor.coef.name = "rho",
           label.y.npc = 0.11,
           label.x.npc = 0.5,
           hjust = 0.5,
           size = 3) +
  stat_cor(method = "spearman",
           aes(label = ..p.label..),
           cor.coef.name = "rho",
           label.y.npc = 0.045,
           label.x.npc = 0.5,
           hjust = 0.5,
           size = 3) +
  annotate("text",
           label = "L.D.",
           x = 0.0001,
           y = 25,
           color = "red",
           size = 2.8,
           hjust = 0) +
  facet_wrap(~Genus_Name) +
  labs(y = "Relative Cys Conc.", 
       x = "Bacterial Relative Abundance") +
   theme(axis.text = element_text(size = 7.5, color = "black"),
        axis.title = element_text(size = 8.5, face = "bold"),
        strip.text = element_text(face = "italic", size = 8.5),
        panel.grid = element_blank())


p.cysteine.pseudocount.genuslevel.correlationplot.horizontal
  
#Vertical
p.cysteine.pseudocount.genuslevel.correlationplot.vertical <- Genus.abund.CT.long.sumstats %>% 
  dplyr::filter(Genus_Name %in% c("Lactobacillus",
                                  "Gardnerella",
                                  "Prevotella")) %>%
  dplyr::mutate(Genus_Name = factor(Genus_Name,
                                    levels = c("Lactobacillus",
                                  "Gardnerella",
                                  "Prevotella"))) %>%
  dplyr::left_join(cysteine_data.processed, by = c("Subject_ID", "Collection_Date", "Visit_Code")) %>%
  ggplot(aes(x = Pseudo_Rel_Abund,
             y = cysteine # (Relative/imputed),
             )) +
  geom_vline(aes(xintercept = Microb_LoD), 
               linetype = "dotted",
               color = "red") +
  geom_smooth( method='lm' , se = TRUE) +
  geom_point(size = 1, shape = 18) +
  scale_x_log10(labels = scales::trans_format("log10", scales::math_format(10^.x)),
                expand = expansion(mult = c(0.075, 0.05))) +
  scale_y_log10(labels = scales::trans_format("log10", scales::math_format(10^.x)),
                expand = expansion(mult = c(0.1, 0.05))) +
  annotation_logticks(sides = "bl",
                      long = unit(0.15, "cm"),
                      mid = unit(0.075, "cm"),
                      short = unit(0.037, "cm")) +
  stat_cor(method = "spearman",
           aes(label = ..r.label..),
           cor.coef.name = "rho",
           label.y.npc = 0.11,
           label.x.npc = 0.5,
           hjust = 0.5,
           size = 3) +
  stat_cor(method = "spearman",
           aes(label = ..p.label..),
           cor.coef.name = "rho",
           label.y.npc = 0.045,
           label.x.npc = 0.5,
           hjust = 0.5,
           size = 3) +
  annotate("text",
           label = "L.D.",
           x = 0.0001,
           y = 25,
           color = "red",
           size = 2.8,
           hjust = 0) +
  facet_grid(Genus_Name~.) +
  labs(y = "Relative Cys Conc.", 
       x = "Bacterial Relative Abundance") +
   theme(axis.text = element_text(size = 7.5, color = "black"),
        axis.title = element_text(size = 8.5, face = "bold"),
        strip.text = element_text(face = "italic", size = 8.5),
        panel.grid = element_blank())


p.cysteine.pseudocount.genuslevel.correlationplot.vertical
```


### Not shown (for now): Point plots of correlation between log-transformed relative abundance (with pseudocount) of Lactobacillus, Gardnerella, and Prevotella (all Genus level) against log-transformed cysteinylglycine and GSH concentrations

```{r correlation-plots-of-selected-genera-cysgly, echo=FALSE}

p.cysgly.pseudocount.genuslevel.correlationplot <- Genus.abund.CT.long.sumstats %>% 
  dplyr::filter(Genus_Name %in% c("Lactobacillus",
                                  "Gardnerella",
                                  "Prevotella")) %>%
  dplyr::mutate(Genus_Name = factor(Genus_Name,
                                    levels = c("Lactobacillus",
                                  "Gardnerella",
                                  "Prevotella"))) %>%
  # dplyr::rename(Visit_Code = VisitCode) %>%
  dplyr::left_join(cysteine_data.processed, 
                   by = c("Subject_ID", "Collection_Date", "Visit_Code")) %>%
  ggplot(aes(x = Pseudo_Rel_Abund,
             y = cysteinylglycine # (Relative/imputed),
             )) +
  geom_vline(aes(xintercept = Microb_LoD), 
               linetype = "dotted",
               color = "red") +
  geom_hline(aes(yintercept = cysteinylglycine_LoD), 
               linetype = "dotted",
               color = "red") +
  geom_smooth( method='lm' , se = TRUE) +
  geom_point(size = 1, shape = 18) +
  scale_x_log10(labels = scales::trans_format("log10", scales::math_format(10^.x)),
                expand = expansion(mult = c(0.05))) +
  scale_y_log10(labels = scales::trans_format("log10", scales::math_format(10^.x)),
                expand = expansion(mult = c(0.05))) +
  annotation_logticks(sides = "bl",
                      long = unit(0.15, "cm"),
                      mid = unit(0.075, "cm"),
                      short = unit(0.037, "cm")) +
  stat_cor(method = "spearman",
           aes(label = ..r.label..),
           cor.coef.name = "rho",
           label.y.npc = 0.19,
           label.x.npc = 0.5,
           hjust = 0.5,
           size = 3) +
  stat_cor(method = "spearman",
           aes(label = ..p.label..),
           cor.coef.name = "rho",
           label.y.npc = 0.15,
           label.x.npc = 0.5,
           hjust = 0.5,
           size = 3) +
  annotate("text",
           label = "L.D.",
           x = 0.0001,
           y = 10,
           color = "red",
           size = 2.8,
           hjust = 0) +
  facet_wrap(~Genus_Name) +
  labs(y = "Relative Cys-Gly Conc.", 
       x = "Bacterial Relative Abundance") +
   theme(axis.text = element_text(size = 7.5, color = "black"),
        axis.title = element_text(size = 8.5, face = "bold"),
        strip.text = element_text(face = "italic", size = 8.5),
        panel.grid = element_blank())

p.cysgly.pseudocount.genuslevel.correlationplot
```
  
```{r correlation-plots-of-selected-genera-gsh, echo=FALSE}

p.gsh.pseudocount.genuslevel.correlationplot <- Genus.abund.CT.long.sumstats %>% 
  dplyr::filter(Genus_Name %in% c("Lactobacillus",
                                  "Gardnerella",
                                  "Prevotella")) %>%
  dplyr::mutate(Genus_Name = factor(Genus_Name,
                                    levels = c("Lactobacillus",
                                  "Gardnerella",
                                  "Prevotella"))) %>%
  dplyr::left_join(cysteine_data.processed, by = c("Subject_ID", "Collection_Date", "Visit_Code")) %>%
  ggplot(aes(x = Pseudo_Rel_Abund,
             y = `glutathione, reduced (GSH)` # (Relative/imputed),
             )) +
  geom_vline(aes(xintercept = Microb_LoD), 
               linetype = "dotted",
               color = "red") +
  geom_hline(aes(yintercept = `glutathione, reduced (GSH)_LoD`), 
               linetype = "dotted",
               color = "red") +
  geom_smooth( method='lm' , se = TRUE) +
  geom_point(size = 1, shape = 18) +
  scale_x_log10(labels = scales::trans_format("log10", scales::math_format(10^.x)),
                expand = expansion(mult = c(0.05))) +
  scale_y_log10(labels = scales::trans_format("log10", scales::math_format(10^.x)),
                expand = expansion(mult = c(0.05))) +
  annotation_logticks(sides = "bl",
                      long = unit(0.15, "cm"),
                      mid = unit(0.075, "cm"),
                      short = unit(0.037, "cm")) +
  stat_cor(method = "spearman",
           aes(label = ..r.label..),
           cor.coef.name = "rho",
           label.y.npc = 0.13,
           label.x.npc = 0.5,
           hjust = 0.5,
           size = 3) +
  stat_cor(method = "spearman",
           aes(label = ..p.label..),
           cor.coef.name = "rho",
           label.y.npc = 0.07,
           label.x.npc = 0.5,
           hjust = 0.5,
           size = 3) +
  annotate("text",
           label = "L.D.",
           x = 0.00008,
           y = 10,
           color = "red",
           size = 2.8,
           hjust = 0) +
  facet_wrap(~Genus_Name) +
  labs(y = "Relative GSH Conc.", 
       x = "Bacterial Relative Abundance") +
   theme(axis.text = element_text(size = 7.5, color = "black"),
        axis.title = element_text(size = 8.5, face = "bold"),
        strip.text = element_text(face = "italic", size = 8.5),
        panel.grid = element_blank())

p.gsh.pseudocount.genuslevel.correlationplot
```


### Supplementary figure 3b: Point plots of correlation between log-transformed relative abundance (with pseudocount) of Lactobacillus crispatus and Lactobacillus iners (Species level) against log-transformed cysteine, cysteinylglycine, and GSH concentration

```{r correlation-plots-of-cysteine-L-iners-and-crispatus, echo=FALSE}

p.cysteine.pseudocount.lacto.specieslevel.correlationplot <- Species.abund.CT.long.sumstats %>% 
  dplyr::filter(Species_Name %in% c("Lactobacillus_crispatus",
                                    "Lactobacillus_iners")) %>%
  dplyr::mutate(Species_Name = str_replace(Species_Name, "actobacillus_", ". ")) %>% 
  dplyr::left_join(cysteine_data.processed, by = c("Subject_ID", "Collection_Date", "Visit_Code")) %>%
  ggplot(aes(x = Pseudo_Rel_Abund,
             y = cysteine #(Relative/imputed),
             )) +
  geom_vline(aes(xintercept = Microb_LoD), 
               linetype = "dotted",
               color = "red") +
  geom_smooth( method='lm' , se = TRUE) +
  geom_point(size = 1, shape = 18) +
  scale_x_log10(labels = scales::trans_format("log10", scales::math_format(10^.x)),
                expand = expansion(mult = c(0.1, 0.05))) +
  scale_y_log10(labels = scales::trans_format("log10", scales::math_format(10^.x)),
                expand = expansion(mult = c(0.1, 0.05))) +
  annotation_logticks(sides = "bl",
                      long = unit(0.15, "cm"),
                      mid = unit(0.075, "cm"),
                      short = unit(0.037, "cm")) +
  stat_cor(method = "spearman",
           aes(label = ..r.label..),
           cor.coef.name = "rho",
           label.y.npc = 0.15,
           label.x.npc = 0.6,
           hjust = 0.5,
           size = 3) +
  stat_cor(method = "spearman",
           aes(label = ..p.label..),
           cor.coef.name = "rho",
           label.y.npc = 0.07,
           label.x.npc = 0.6,
           hjust = 0.5,
           size = 3) +
  annotate("text",
           label = "L.D.",
           x = 0.0001,
           y = 20,
           color = "red",
           size = 2.8,
           hjust = 0) +
  facet_wrap(~Species_Name) +
  labs(y = "Relative Cys Conc.", 
       x = "Bacterial Relative Abundance") +
   theme(axis.text = element_text(size = 7.5, color = "black"),
        axis.title = element_text(size = 8.5, face = "bold"),
        strip.text = element_text(face = "italic", size = 8.5),
        panel.grid = element_blank())

p.cysteine.pseudocount.lacto.specieslevel.correlationplot
```
  
  
### Not shown (for now): point plots of L. iners and L. crispatus against GSH and Cys-Gly  
  
  
```{r correlation-plots-of-cysgly-L-iners-and-crispatus, echo=FALSE}

p.cysgly.pseudocount.lacto.specieslevel.correlationplot <- Species.abund.CT.long.sumstats %>% 
  dplyr::filter(Species_Name %in% c("Lactobacillus_crispatus",
                                    "Lactobacillus_iners")) %>%
  dplyr::mutate(Species_Name = str_replace(Species_Name, "actobacillus_", ". ")) %>% 
  dplyr::left_join(cysteine_data.processed, by = c("Subject_ID", "Collection_Date", "Visit_Code")) %>%
  ggplot(aes(x = Pseudo_Rel_Abund,
             y = cysteinylglycine #(Relative/imputed),
             )) +
  geom_vline(aes(xintercept = Microb_LoD), 
               linetype = "dotted",
               color = "red") +
  geom_hline(aes(yintercept = `cysteinylglycine_LoD`), 
               linetype = "dotted",
               color = "red") +
  geom_smooth( method='lm' , se = TRUE) +
  geom_point(size = 1, shape = 18) +
  scale_x_log10(labels = scales::trans_format("log10", scales::math_format(10^.x)),
                expand = expansion(mult = c(0.05))) +
  scale_y_log10(labels = scales::trans_format("log10", scales::math_format(10^.x)),
                expand = expansion(mult = c(0.05))) +
  annotation_logticks(sides = "bl",
                      long = unit(0.15, "cm"),
                      mid = unit(0.075, "cm"),
                      short = unit(0.037, "cm")) +
  stat_cor(method = "spearman",
           aes(label = ..r.label..),
           cor.coef.name = "rho",
           label.y.npc = 0.2,
           label.x.npc = 0.6,
           hjust = 0.5,
           size = 3) +
  stat_cor(method = "spearman",
           aes(label = ..p.label..),
           cor.coef.name = "rho",
           label.y.npc = 0.15,
           label.x.npc = 0.6,
           hjust = 0.5,
           size = 3) +
  annotate("text",
           label = "L.D.",
           x = 0.0001,
           y = 10,
           color = "red",
           size = 2.8,
           hjust = 0) +
  facet_wrap(~Species_Name) +
  labs(y = "Relative Cys-Gly Conc.", 
       x = "Bacterial Relative Abundance") +
   theme(axis.text = element_text(size = 7.5, color = "black"),
        axis.title = element_text(size = 8.5, face = "bold"),
        strip.text = element_text(face = "italic", size = 8.5),
        panel.grid = element_blank())

p.cysgly.pseudocount.lacto.specieslevel.correlationplot
```

```{r correlation-plots-of-gsh-L-iners-and-crispatus, echo=FALSE}

p.gsh.pseudocount.lacto.specieslevel.correlationplot <- Species.abund.CT.long.sumstats %>% 
  dplyr::filter(Species_Name %in% c("Lactobacillus_crispatus",
                                    "Lactobacillus_iners")) %>%
  dplyr::mutate(Species_Name = str_replace(Species_Name, "_", " ")) %>% 
  dplyr::left_join(cysteine_data.processed, by = c("Subject_ID", "Collection_Date", "Visit_Code")) %>%
  ggplot(aes(x = Pseudo_Rel_Abund,
             y = `glutathione, reduced (GSH)` #(Relative/imputed),
             )) +
  geom_vline(aes(xintercept = Microb_LoD), 
               linetype = "dotted",
               color = "red") +
  geom_hline(aes(yintercept = `glutathione, reduced (GSH)_LoD`), 
               linetype = "dotted",
               color = "red") +
  geom_smooth( method='lm' , se = TRUE) +
  geom_point(size = 1, shape = 18) +
  scale_x_log10(labels = scales::trans_format("log10", scales::math_format(10^.x)),
                expand = expansion(mult = c(0.05))) +
  scale_y_log10(labels = scales::trans_format("log10", scales::math_format(10^.x)),
                expand = expansion(mult = c(0.05))) +
  annotation_logticks(sides = "bl",
                      long = unit(0.15, "cm"),
                      mid = unit(0.075, "cm"),
                      short = unit(0.037, "cm")) +
  stat_cor(method = "spearman",
           aes(label = ..r.label..),
           cor.coef.name = "rho",
           label.y.npc = 0.2,
           label.x.npc = 0.6,
           hjust = 0.5,
           size = 3) +
  stat_cor(method = "spearman",
           aes(label = ..p.label..),
           cor.coef.name = "rho",
           label.y.npc = 0.15,
           label.x.npc = 0.6,
           hjust = 0.5,
           size = 3) +
  annotate("text",
           label = "L.D.",
           x = 0.0001,
           y = 10,
           color = "red",
           size = 2.8,
           hjust = 0) +
  facet_wrap(~Species_Name) +
  labs(y = "Relative GSH Conc.", 
       x = "Bacterial Relative Abundance") +
   theme(axis.text = element_text(size = 7.5, color = "black"),
        axis.title = element_text(size = 8.5, face = "bold"),
        strip.text = element_text(face = "italic", size = 8.5),
        panel.grid = element_blank())

p.gsh.pseudocount.lacto.specieslevel.correlationplot
```
  
### Supplementary Figures 3C and 3D: Summary statistics of genus or of species with >50% prevalence (table and/or box/violin plot)
  
  
Species distribution.  
  
  

```{r filter-Genus-to-Prev-50-and-review-taxa-distributions, echo=FALSE}
#Calculate Prevalence within cohort,
#Filter on Prevalence,
#Calculate Rel_Abund summary statistics
Genus.ra.table.p50.stats <- Genus.abund.CT.long.sumstats %>% 
  dplyr::filter(Prev_50) %>% 
  dplyr::group_by(Genus_Name) %>% 
  dplyr::mutate(Median_RA = median(Rel_Abund),
                Percentile_5th_RA = quantile(Rel_Abund, probs = 0.05),
                Lower_Quartile_RA = quantile(Rel_Abund, probs = 0.25),
                Upper_Quartile_RA = quantile(Rel_Abund, probs = 0.75),
                Percentile_95th_RA = quantile(Rel_Abund, probs = 0.95),
                Median_Pseudo_RA = median(Pseudo_Rel_Abund),
                Percentile_5th_Pseudo_RA = quantile(Pseudo_Rel_Abund, probs = 0.05),
                Lower_Quartile_Pseudo_RA = quantile(Pseudo_Rel_Abund, probs = 0.25),
                Upper_Quartile_Pseudo_RA = quantile(Pseudo_Rel_Abund, probs = 0.75),
                Percentile_95th_Pseudo_RA = quantile(Pseudo_Rel_Abund, probs = 0.95)) %>% 
  dplyr::ungroup() %>% 
  dplyr::arrange(desc(Cohort_Prev), Genus_Name, Seq_ID)

#Review summary table
Genus.ra.table.p50.stats %>% 
  dplyr::distinct(Genus = Genus_Name, 
                  `Prevalance in Cohort` = Cohort_Prev, 
                  `Relative Abundance in Entire Cohort` = Cohort_RA, 
                  `Per-Sample Relative Abundance, 5th Percentile` = Percentile_5th_RA, 
                  `Per-Sample Relative Abundance, 25th Percentile` = Lower_Quartile_RA, 
                  `Per-Sample Relative Abundance, Median` = Median_RA, 
                  `Per-Sample Relative Abundance, 75th Percentile` = Upper_Quartile_RA, 
                  `Per-Sample Relative Abundance, 95th Percentile` = Percentile_95th_RA) %>% 
  dplyr::mutate(across(-Genus, function(x) {paste0(signif(x * 100, 2), "%")})) %>% 
  knitr::kable(caption = "Prevalence and Relative Abundance of Genera with >=50% Prevalence")

### Plot taxon relative abundances, violin plot with crossbars for median and IQR and with LoD
p.Genus.p50.ra.violin <- Genus.ra.table.p50.stats %>% 
  dplyr::mutate(Genus_Name = str_replace(Genus_Name, "ndidatus", "."),
                Genus_Name = str_replace_all(Genus_Name, "_", " "),
                Genus = fct_inorder(Genus_Name),
                Genus = fct_rev(Genus)) %>% 
  ggplot(aes(y = Genus, x = Pseudo_Rel_Abund)) + 
  geom_vline(aes(xintercept = Microb_LoD), linetype = "dotted",
           color = "red") +
  geom_violin(fill = NA) +
  geom_errorbar(aes(xmin = Lower_Quartile_Pseudo_RA,
                    xmax = Lower_Quartile_Pseudo_RA),
                width = 0.7, color = "blue",
                size = 0.5) +
  geom_errorbar(aes(xmin = Median_Pseudo_RA, 
                    xmax = Median_Pseudo_RA),
                width = 0.9, color = "purple",
                size = 1) +
  geom_errorbar(aes(xmin = Upper_Quartile_Pseudo_RA, 
                    xmax = Upper_Quartile_Pseudo_RA),
                width = 0.9, color = "blue",
                size = 0.5) +
  geom_jitter(width = 0, height = 0.1, 
              alpha = 0.8, size = 0.5, shape = 18) +
  annotate("text",
           label = "L.D.",
            x = 0.00008,
            y = 15,
           hjust = 0,
           color = "red",
           size = 2.8) +
  scale_x_log10(labels = scales::trans_format("log10", scales::math_format(10^.x))) +
  scale_y_discrete(expand = expansion(add = c(1, 1.7))) +
  annotation_logticks(sides = "b",
                      long = unit(0.15, "cm"),
                      mid = unit(0.075, "cm"),
                      short = unit(0.037, "cm")) +
  labs(x = "Relative Abundance",
       tag = "Genus") +
  theme(axis.text.y = element_text(size = 7.5, face = "italic", color = "black"),
        axis.text.x = element_text(hjust = 1, size = 7.5, color = "black"),
        axis.title.x = element_text(size = 8.5, face = "bold"),
        axis.title.y = element_blank(),
        plot.tag.position = c(0.4, 0.97),
        plot.tag = element_text(face = "bold",
                                size = 8.5,
                                hjust = 1,
                                vjust = 0.5),
        panel.grid = element_blank())

p.Genus.p50.ra.violin

#Plot taxon prevalences
p.Genus.p50.prev.col <- Genus.ra.table.p50.stats %>% 
  dplyr::mutate(Genus_Name = str_replace(Genus_Name, "ndidatus", "."),
                Genus_Name = str_replace_all(Genus_Name, "_", " "),
                Genus = fct_inorder(Genus_Name),
                Genus = fct_rev(Genus)) %>% 
  dplyr::distinct(Genus, Cohort_Prev, Cohort_RA) %>% 
  ggplot(aes(x = Cohort_Prev, y = Genus)) +
  geom_col(width = 0.6, alpha = 0.6) +
  scale_x_continuous(labels = scales::percent_format(),
                     breaks = c(0:2) / 2) +
  scale_y_discrete(expand = expansion(add = c(1, 1.7))) +
  labs(x = "Prevalence") +
  theme(axis.text.y = element_blank(),
        axis.text.x = element_text(
          # angle = 45, hjust = 1, 
                                   size = 7.5, color = "black"),
        axis.title.x = element_text(size = 8.5, face = "bold"),
        axis.title.y = element_blank(),
        axis.ticks.y = element_blank(),
        panel.grid.major.y = element_blank(),
        panel.grid.minor.y = element_blank(),
        panel.grid.major.x = element_line(linetype = "dashed",
                                          colour = "black",
                                          size = 0.25),
        panel.grid.minor.x = element_blank())

p.Genus.p50.prev.col

#Align rel abundance and prevalence plots
p.Genus.p50.ra.prev <- cowplot::plot_grid(p.Genus.p50.ra.violin,
                                          p.Genus.p50.prev.col,
                                          rel_widths = c(1, 0.3),
                                          align = "h",
                                          axis = "b"
                                          )

p.Genus.p50.ra.prev
```
  
Species distribution.  
  
  
```{r filter-Species-to-Prev-50-and-review-taxa-distributions, echo=FALSE}
#Calculate Prevalence within cohort,
#Filter on Prevalence,
#Calculate Rel_Abund summary statistics
Species.ra.table.p50.stats <- Species.abund.CT.long.sumstats %>% 
  dplyr::filter(Prev_50) %>% 
  dplyr::group_by(Species_Name) %>% 
  dplyr::mutate(Median_RA = median(Rel_Abund),
                Percentile_5th_RA = quantile(Rel_Abund, probs = 0.05),
                Lower_Quartile_RA = quantile(Rel_Abund, probs = 0.25),
                Upper_Quartile_RA = quantile(Rel_Abund, probs = 0.75),
                Percentile_95th_RA = quantile(Rel_Abund, probs = 0.95),
                Median_Pseudo_RA = median(Pseudo_Rel_Abund),
                Percentile_5th_Pseudo_RA = quantile(Pseudo_Rel_Abund, probs = 0.05),
                Lower_Quartile_Pseudo_RA = quantile(Pseudo_Rel_Abund, probs = 0.25),
                Upper_Quartile_Pseudo_RA = quantile(Pseudo_Rel_Abund, probs = 0.75),
                Percentile_95th_Pseudo_RA = quantile(Pseudo_Rel_Abund, probs = 0.95)) %>% 
  dplyr::ungroup() %>% 
  dplyr::arrange(desc(Cohort_Prev), Species_Name, Seq_ID)

#Review summary table
Species.ra.table.p50.stats %>% 
  dplyr::distinct(Species = Species_Name, 
                  `Prevalance in Cohort` = Cohort_Prev, 
                  `Relative Abundance in Entire Cohort` = Cohort_RA, 
                  `Per-Sample Relative Abundance, 5th Percentile` = Percentile_5th_RA, 
                  `Per-Sample Relative Abundance, 25th Percentile` = Lower_Quartile_RA, 
                  `Per-Sample Relative Abundance, Median` = Median_RA, 
                  `Per-Sample Relative Abundance, 75th Percentile` = Upper_Quartile_RA, 
                  `Per-Sample Relative Abundance, 95th Percentile` = Percentile_95th_RA) %>% 
  dplyr::mutate(across(-Species, function(x) {paste0(signif(x * 100, 2), "%")})) %>% 
  knitr::kable(caption = "Prevalence and Relative Abundance of Species with >=50% Prevalence")

### Plot taxon relative abundances, violin plot with crossbars for median and IQR and with LoD
p.Species.p50.ra.violin <- Species.ra.table.p50.stats %>% 
  dplyr::mutate(Species_Name = str_replace(Species_Name, "ndidatus", "."),
                Species_Name = str_replace_all(Species_Name, "_", " "),
                Species = fct_inorder(Species_Name),
                Species = fct_rev(Species)) %>% 
  ggplot(aes(y = Species, x = Pseudo_Rel_Abund)) + 
  geom_vline(aes(xintercept = Microb_LoD), linetype = "dotted",
           color = "red") +
  geom_violin(fill = NA) +
  geom_errorbar(aes(xmin = Lower_Quartile_Pseudo_RA,
                    xmax = Lower_Quartile_Pseudo_RA),
                width = 0.7, color = "blue",
                size = 0.5) +
  geom_errorbar(aes(xmin = Median_Pseudo_RA, 
                    xmax = Median_Pseudo_RA),
                width = 0.9, color = "purple",
                size = 1) +
  geom_errorbar(aes(xmin = Upper_Quartile_Pseudo_RA, 
                    xmax = Upper_Quartile_Pseudo_RA),
                width = 0.9, color = "blue",
                size = 0.5) +
  geom_jitter(width = 0, height = 0.1, 
              alpha = 0.8, size = 0.5, shape = 18) +
  annotate("text",
           label = "L.D.",
            x = 0.00008,
            y = 16,
           hjust = 0,
           color = "red",
           size = 2.8) +
  scale_x_log10(labels = scales::trans_format("log10", scales::math_format(10^.x))) +
  scale_y_discrete(expand = expansion(add = c(1, 1.7))) +
  annotation_logticks(sides = "b",
                      long = unit(0.15, "cm"),
                      mid = unit(0.075, "cm"),
                      short = unit(0.037, "cm")) +
  labs(x = "Relative Abundance",
       tag = "Species") +
  theme(axis.text.y = element_text(size = 7.5, face = "italic", color = "black"),
        axis.text.x = element_text(hjust = 1, size = 7.5, color = "black"),
        axis.title.x = element_text(size = 8.5, face = "bold"),
        axis.title.y = element_blank(),
        plot.tag.position = c(0.44, 0.973),
        plot.tag = element_text(face = "bold",
                                size = 8.5,
                                hjust = 1,
                                vjust = 0.5),
        panel.grid = element_blank())

p.Species.p50.ra.violin

#Plot taxon prevalences
p.Species.p50.prev.col <- Species.ra.table.p50.stats %>% 
  dplyr::mutate(Species_Name = str_replace(Species_Name, "ndidatus", "."),
                Species_Name = str_replace_all(Species_Name, "_", " "),
                Species = fct_inorder(Species_Name),
                Species = fct_rev(Species)) %>% 
  dplyr::distinct(Species, Cohort_Prev, Cohort_RA) %>% 
  ggplot(aes(x = Cohort_Prev, y = Species)) +
  geom_col(width = 0.6, alpha = 0.6) +
  scale_x_continuous(labels = scales::percent_format(),
                     breaks = c(0:2) / 2) +
  scale_y_discrete(expand = expansion(add = c(1, 1.7))) +
  labs(x = "Prevalence") +
  theme(axis.text.y = element_blank(),
        axis.text.x = element_text(
          # angle = 45, hjust = 1, 
                                   size = 7.5, 
                                   color = "black"),
        axis.title.x = element_text(size = 8.5, face = "bold"),
        axis.title.y = element_blank(),
        axis.ticks.y = element_blank(),
        panel.grid.major.y = element_blank(),
        panel.grid.minor.y = element_blank(),
        panel.grid.major.x = element_line(linetype = "dashed",
                                          colour = "black",
                                          size = 0.25),
        panel.grid.minor.x = element_blank())

p.Species.p50.prev.col

#Align rel abundance and prevalence plots
p.Species.p50.ra.prev <- cowplot::plot_grid(p.Species.p50.ra.violin,
                                          p.Species.p50.prev.col,
                                          rel_widths = c(1, 0.27),
                                          align = "h",
                                          axis = "b"
                                          )

p.Species.p50.ra.prev
```

  
  
###  Panel e: Cys correlation Forest plot at Genus level 


```{r Genus-Metabolite-correlation-calcs, echo=FALSE, warning=FALSE}
# merge and filter
cysteine_data_genus <- Genus.abund.CT.long %>%
  dplyr::group_by(Genus_Name) %>% 
  dplyr::mutate(Prevalence = sum(Rel_Abund > 0) / n()) %>% 
  dplyr::filter(Prevalence >= 0.5) %>%
  dplyr::ungroup() %>%
  dplyr::right_join(cysteine_data.processed, 
                    by = c("Subject_ID", 
                          "Collection_Date",
                          "Visit_Code"))

#NEST DATA
cysteine_data_genus.nest <- cysteine_data_genus %>%
  dplyr::select(cysteine, cysteinylglycine, `glutathione, reduced (GSH)`, Genus_Name, Rel_Abund) %>% 
  tidyr::pivot_longer(cols = c(cysteine, cysteinylglycine, `glutathione, reduced (GSH)`),
                      names_to = "Metabolite",
                      values_to = "Concentration") %>% 
  dplyr::mutate(Metabolite = case_when(Metabolite == "cysteine" ~ "Cys",
                                       Metabolite == "cysteinylglycine" ~ "Cys-Gly",
                                       Metabolite == "glutathione, reduced (GSH)" ~ "GSH"),
                Metabolite = factor(Metabolite,
                                    levels = c("Cys",
                                               "GSH",
                                               "Cys-Gly"))) %>% 
  tidyr::nest(data = c(Rel_Abund, Concentration)) %>% 
  dplyr::arrange(Metabolite)
  
#Spearman correlation and associated statistics
cysteine_genus_cor.CI <- cysteine_data_genus.nest %>% 
  dplyr::group_by(Metabolite) %>% 
  dplyr::mutate(Cor_Spear = map(data,
                                ~ cor.test(~ Rel_Abund + Concentration,
                                           data = .x,
                                           method = "spearman")),
                Rho = map_dbl(Cor_Spear,
                              ~ .x$estimate),
                Rho_MinCI_95 = map_dbl(data,
                                       ~ SpearmanRho(x = .x$Rel_Abund,
                                                      y = .x$Concentration,
                                                      conf.level = 0.95)[2]),
                Rho_MaxCI_95 = map_dbl(data,
                                       ~ SpearmanRho(x = .x$Rel_Abund,
                                                      y = .x$Concentration,
                                                      conf.level = 0.95)[3]),
                Conf_level_adjusted = (1 - 0.05/n()),
                Rho_MinCI_adj = map_dbl(data,
                                       ~ SpearmanRho(x = .x$Rel_Abund,
                                                      y = .x$Concentration,
                                                      conf.level = Conf_level_adjusted)[2]),
                Rho_MaxCI_adj = map_dbl(data,
                                       ~ SpearmanRho(x = .x$Rel_Abund,
                                                      y = .x$Concentration,
                                                      conf.level = Conf_level_adjusted)[3]),
                P_value = map_dbl(Cor_Spear,
                                  ~ .x$p.value),
                P_adjust = p.adjust(P_value,
                                    method = "bonferroni"),
                Sigsymbol = case_when(P_value <= 0.0001 ~ "****",
                                      P_value <= 0.001 ~ "***",
                                      P_value <= 0.01 ~ "**",
                                      P_value <= 0.05 ~ "*",
                                      TRUE ~ "n.s."),
                Sigsymbol_adj = case_when(P_adjust <= 0.0001 ~ "****",
                                          P_adjust <= 0.001 ~ "***",
                                          P_adjust <= 0.01 ~ "**",
                                          P_adjust <= 0.05 ~ "*",
                                          TRUE ~ "n.s.")) %>% 
  dplyr::ungroup() %>% 
  dplyr::arrange(Metabolite, Rho)
```


```{r Genus-Metabolite-correlation-plot-all, echo=FALSE, warning=FALSE}
# plot 
p.Cys.Metab.Genus.corr.forest.all <- cysteine_genus_cor.CI %>% 
  dplyr::mutate(Genus_Name = str_replace_all(Genus_Name, "_", " "),
                Genus_Name = str_replace(Genus_Name, "ndidatus", "."),
                Genus_Name = fct_inorder(Genus_Name),
                Sigsymbol_adj = case_when(Sigsymbol_adj == "n.s." ~ " ",
                                          TRUE ~ Sigsymbol_adj),
                StatSignif = P_adjust < 0.05,
                CI_label = paste0("(",
                                  round(Conf_level_adjusted * 100, 2),
                                  "% C.I.)")) %>%
  {ggplot(., aes(x = Rho, y = Genus_Name, color = StatSignif)) +
  geom_vline(xintercept = 0, linetype = "dotted") +
  geom_pointrange(aes(xmin = Rho_MinCI_adj,
                      xmax = Rho_MaxCI_adj),
                  size = 0.5,
                  shape = 18) +
  geom_text(aes(label = Sigsymbol_adj), 
            hjust = 0.5,
            vjust = 0.2,
            size = 3.5) +
  scale_x_continuous(expand = expansion(mult = 0.1)) +
  scale_y_discrete(expand = expansion(add = c(0.5, 1))) +
  scale_color_manual(values = c("black", "red")) +
  labs(x = bquote("Correlation"~
                  rho~.(.$CI_label)),
       tag = "Genus") +
  theme(axis.title.x = element_text(size = 8.5, face = "bold"),
        axis.title.y = element_blank(),
        axis.text.y = element_text(size = 7.5, 
                                   face = "italic", 
                                   color = "black"),
        axis.text.x = element_text(size = 7.5,
                                   angle = 45,
                                   hjust = 1, 
                                   color = "black"
                                   ),
        strip.text = element_text(size = 8.5),
        panel.grid.major.x = element_blank(),
        panel.grid.minor.x = element_blank(),
        panel.grid.major.y = element_line(size = 0.25),
        legend.position = "none",
        plot.tag.position = c(0.31, 0.94),
        plot.tag = element_text(face = "bold",
                                size = 8.5,
                                hjust = 1,
                                vjust = 0.5)) +
  facet_wrap(~Metabolite)}

p.Cys.Metab.Genus.corr.forest.all
```

```{r Genus-correlation-plot-cys-only, echo=FALSE, warning=FALSE}
# plot 
p.Cys.Genus.corr.forest <- cysteine_genus_cor.CI %>% 
  dplyr::filter(Metabolite == "Cys") %>% 
  dplyr::mutate(Genus_Name = str_replace_all(Genus_Name, "_", " "),
                Genus_Name = str_replace(Genus_Name, "ndidatus", "."),
                Genus_Name = fct_inorder(Genus_Name),
                Sigsymbol_adj = case_when(Sigsymbol_adj == "n.s." ~ " ",
                                          TRUE ~ Sigsymbol_adj),
                StatSignif = P_adjust < 0.05,
                CI_label = paste0("(",
                                  round(Conf_level_adjusted * 100, 2),
                                  "% C.I.)")) %>%
  {ggplot(., aes(x = Rho, y = Genus_Name, color = StatSignif)) +
  geom_vline(xintercept = 0, linetype = "dotted") +
  geom_pointrange(aes(xmin = Rho_MinCI_adj,
                      xmax = Rho_MaxCI_adj),
                  size = 0.5,
                  shape = 18) +
  geom_text(aes(label = Sigsymbol_adj), 
            hjust = 0.5,
            vjust = 0.2,
            size = 3.5) +
  scale_x_continuous(expand = expansion(mult = 0.1)) +
  scale_y_discrete(expand = expansion(add = c(0.5, 1))) +
  scale_color_manual(values = c("black", "red")) +
  labs(x = bquote(atop("Correlation"~
                  rho,
                  .(.$CI_label))),
       tag = "Genus") +
  theme(axis.title.x = element_text(size = 8.5, face = "bold"),
        axis.title.y = element_blank(),
        axis.text.y = element_text(size = 7.5, 
                                   face = "italic", 
                                   color = "black"),
        axis.text.x = element_text(size = 7.5,
                                   angle = 45,
                                   hjust = 1, 
                                   color = "black"
                                   ),
        strip.text = element_text(size = 8.5),
        panel.grid.major.x = element_blank(),
        panel.grid.minor.x = element_blank(),
        panel.grid.major.y = element_line(size = 0.25),
        legend.position = "none",
        plot.tag.position = c(0.31, 0.94),
        plot.tag = element_text(face = "bold",
                                size = 8.5,
                                hjust = 1,
                                vjust = 0.5)) +
  facet_wrap(~Metabolite)}

p.Cys.Genus.corr.forest
```
  
  
### Supplementary figure 4e: GSH and Cys-Gly correlation Forest plot at Genus level  
  

```{r Genus-GSH-CysGly-correlation-plot-only, echo=FALSE, warning=FALSE}
# plot 
p.metab.Genus.corr.forest <- cysteine_genus_cor.CI %>% 
  dplyr::filter(Metabolite %in% c("GSH", "Cys-Gly")) %>% 
  dplyr::mutate(Genus_Name = str_replace_all(Genus_Name, "_", " "),
                Genus_Name = str_replace(Genus_Name, "ndidatus", "."),
                Genus_Name = fct_inorder(Genus_Name),
                Sigsymbol_adj = case_when(Sigsymbol_adj == "n.s." ~ " ",
                                          TRUE ~ Sigsymbol_adj),
                StatSignif = P_adjust < 0.05,
                CI_label = paste0("(",
                                  round(Conf_level_adjusted * 100, 2),
                                  "% C.I.)")) %>%
  {ggplot(., aes(x = Rho, y = Genus_Name, color = StatSignif)) +
  geom_vline(xintercept = 0, linetype = "dotted") +
  geom_pointrange(aes(xmin = Rho_MinCI_adj,
                      xmax = Rho_MaxCI_adj),
                  size = 0.5,
                  shape = 18) +
  geom_text(aes(label = Sigsymbol_adj), 
            hjust = 0.5,
            vjust = 0.2,
            size = 3.5) +
  scale_x_continuous(expand = expansion(mult = 0.1)) +
  scale_y_discrete(expand = expansion(add = c(0.5, 1))) +
  scale_color_manual(values = c("black", "red")) +
  labs(x = bquote(atop("Correlation"~
                  rho,
                  .(.$CI_label))),
       tag = "Genus") +
  theme(axis.title.x = element_text(size = 8.5, face = "bold"),
        axis.title.y = element_blank(),
        axis.text.y = element_text(size = 7.5, 
                                   face = "italic", 
                                   color = "black"),
        axis.text.x = element_text(size = 7.5,
                                   angle = 45,
                                   hjust = 1, 
                                   color = "black"
                                   ),
        strip.text = element_text(size = 8.5),
        panel.grid.major.x = element_blank(),
        panel.grid.minor.x = element_blank(),
        panel.grid.major.y = element_line(size = 0.25),
        legend.position = "none",
        plot.tag.position = c(0.31, 0.94),
        plot.tag = element_text(face = "bold",
                                size = 8.5,
                                hjust = 1,
                                vjust = 0.5)) +
  facet_wrap(~Metabolite)}

p.metab.Genus.corr.forest
```
  
###  Panel 3F: Cys correlation Forest plot at Genus level 


```{r Species-Metabolite-correlation-calcs, echo=FALSE, warning=FALSE}
# filter to taxa with >=50% prevalence in the cohort, then join with
# metabolite data
cysteine_data_species <- Species.abund.CT.long %>%
  dplyr::group_by(Species_Name) %>% 
  dplyr::mutate(Prevalence = sum(Rel_Abund > 0) / n()) %>% 
  dplyr::filter(Prevalence >= 0.5) %>%
  dplyr::ungroup() %>%
  dplyr::right_join(cysteine_data.processed, 
                    by = c("Subject_ID", 
                          "Collection_Date",
                          "Visit_Code"))

#subset to relevant columns, pivot metabolite data to long form,
#then nest concentration and abundance data for each taxon-metabolite combination
cysteine_data_species.nest <- cysteine_data_species %>%
  dplyr::select(cysteine, cysteinylglycine, `glutathione, reduced (GSH)`, Species_Name, Rel_Abund) %>% 
  tidyr::pivot_longer(cols = c(cysteine, cysteinylglycine, `glutathione, reduced (GSH)`),
                      names_to = "Metabolite",
                      values_to = "Concentration") %>% 
  dplyr::mutate(Metabolite = case_when(Metabolite == "cysteine" ~ "Cys",
                                       Metabolite == "cysteinylglycine" ~ "Cys-Gly",
                                       Metabolite == "glutathione, reduced (GSH)" ~ "GSH"),
                Metabolite = factor(Metabolite,
                                    levels = c("Cys",
                                               "GSH",
                                               "Cys-Gly"))) %>% 
  tidyr::nest(data = c(Rel_Abund, Concentration)) %>% 
  dplyr::arrange(Metabolite)
  
#For each taxon-metabolite concentration, calculate Spearman correlation
#and create separate columns in the dataframe for the correlation coefficient (rho),
#95% CI for rho, p-value fro the correlation, bonferroni-adjusted confidence level
#for n multiple comparisons (1 - alpha/n), bonferroni-adjusted p-value,
#bonferroni-adjusted CI, and significance symbols for p-value and adjusted p-value
cysteine_species_cor.CI <- cysteine_data_species.nest %>% 
  dplyr::group_by(Metabolite) %>% 
  dplyr::mutate(Cor_Spear = map(data,
                                ~ cor.test(~ Rel_Abund + Concentration,
                                           data = .x,
                                           method = "spearman")),
                Rho = map_dbl(Cor_Spear,
                              ~ .x$estimate),
                Rho_MinCI_95 = map_dbl(data,
                                       ~ SpearmanRho(x = .x$Rel_Abund,
                                                      y = .x$Concentration,
                                                      conf.level = 0.95)[2]),
                Rho_MaxCI_95 = map_dbl(data,
                                       ~ SpearmanRho(x = .x$Rel_Abund,
                                                      y = .x$Concentration,
                                                      conf.level = 0.95)[3]),
                Conf_level_adjusted = (1 - 0.05/n()),
                Rho_MinCI_adj = map_dbl(data,
                                       ~ SpearmanRho(x = .x$Rel_Abund,
                                                      y = .x$Concentration,
                                                      conf.level = Conf_level_adjusted)[2]),
                Rho_MaxCI_adj = map_dbl(data,
                                       ~ SpearmanRho(x = .x$Rel_Abund,
                                                      y = .x$Concentration,
                                                      conf.level = Conf_level_adjusted)[3]),
                P_value = map_dbl(Cor_Spear,
                                  ~ .x$p.value),
                P_adjust = p.adjust(P_value,
                                    method = "bonferroni"),
                Sigsymbol = case_when(P_value <= 0.0001 ~ "****",
                                      P_value <= 0.001 ~ "***",
                                      P_value <= 0.01 ~ "**",
                                      P_value <= 0.05 ~ "*",
                                      TRUE ~ "n.s."),
                Sigsymbol_adj = case_when(P_adjust <= 0.0001 ~ "****",
                                          P_adjust <= 0.001 ~ "***",
                                          P_adjust <= 0.01 ~ "**",
                                          P_adjust <= 0.05 ~ "*",
                                          TRUE ~ "n.s.")) %>% 
  dplyr::ungroup() %>% 
  dplyr::arrange(Metabolite, Rho)
```


```{r Species-Metabolite-correlation-plot-all, echo=FALSE, warning=FALSE}
# forest plot of Spearman correlation results with bonferroni-adjusted
# p-value and confidence interval
p.Cys.Metab.Species.corr.forest.all <- cysteine_species_cor.CI %>% 
  dplyr::mutate(Genus_Name = str_replace_all(Species_Name, "_", " "),
                Genus_Name = str_replace(Genus_Name, "ndidatus", "."),
                Genus_Name = fct_inorder(Genus_Name),
                Sigsymbol_adj = case_when(Sigsymbol_adj == "n.s." ~ " ",
                                          TRUE ~ Sigsymbol_adj),
                StatSignif = P_adjust < 0.05,
                CI_label = paste0("(",
                                  round(Conf_level_adjusted * 100, 2),
                                  "% C.I.)")) %>%
  {ggplot(., aes(x = Rho, y = Genus_Name, color = StatSignif)) +
  geom_vline(xintercept = 0, linetype = "dotted") +
  geom_pointrange(aes(xmin = Rho_MinCI_adj,
                      xmax = Rho_MaxCI_adj),
                  size = 0.5,
                  shape = 18) +
  geom_text(aes(label = Sigsymbol_adj), 
            hjust = 0.5,
            vjust = 0.2,
            size = 3.5) +
  scale_x_continuous(expand = expansion(mult = 0.1)) +
  scale_y_discrete(expand = expansion(add = c(0.5, 1))) +
  scale_color_manual(values = c("black", "red")) +
  labs(x = bquote("Correlation"~
                  rho~.(.$CI_label)),
       tag = "Species") +
  theme(axis.title.x = element_text(size = 8.5, face = "bold"),
        axis.title.y = element_blank(),
        axis.text.y = element_text(size = 7.5, 
                                   face = "italic", 
                                   color = "black"),
        axis.text.x = element_text(size = 7.5,
                                   angle = 45,
                                   hjust = 1, 
                                   color = "black"
                                   ),
        strip.text = element_text(size = 8.5),
        panel.grid.major.x = element_blank(),
        panel.grid.minor.x = element_blank(),
        panel.grid.major.y = element_line(size = 0.25),
        legend.position = "none",
        plot.tag.position = c(0.37, 0.94),
        plot.tag = element_text(face = "bold",
                                size = 8.5,
                                hjust = 1,
                                vjust = 0.5)) +
  facet_wrap(~Metabolite)}

p.Cys.Metab.Species.corr.forest.all
```


```{r Species-Cys-correlation-plot-only, echo=FALSE, warning=FALSE}
# forest plot of Spearman correlation results with bonferroni-adjusted
# p-value and confidence interval
p.Cys.Species.corr.forest <- cysteine_species_cor.CI %>% 
  dplyr::filter(Metabolite == "Cys") %>% 
  dplyr::mutate(Genus_Name = str_replace_all(Species_Name, "_", " "),
                Genus_Name = str_replace(Genus_Name, "ndidatus", "."),
                Genus_Name = fct_inorder(Genus_Name),
                Sigsymbol_adj = case_when(Sigsymbol_adj == "n.s." ~ " ",
                                          TRUE ~ Sigsymbol_adj),
                StatSignif = P_adjust < 0.05,
                CI_label = paste0("(",
                                  round(Conf_level_adjusted * 100, 2),
                                  "% C.I.)")) %>%
  {ggplot(., aes(x = Rho, y = Genus_Name, color = StatSignif)) +
  geom_vline(xintercept = 0, linetype = "dotted") +
  geom_pointrange(aes(xmin = Rho_MinCI_adj,
                      xmax = Rho_MaxCI_adj),
                  size = 0.5,
                  shape = 18) +
  geom_text(aes(label = Sigsymbol_adj), 
            hjust = 0.5,
            vjust = 0.2,
            size = 3.5) +
  scale_x_continuous(expand = expansion(mult = 0.1)) +
  scale_y_discrete(expand = expansion(add = c(0.5, 1))) +
  scale_color_manual(values = c("black", "red")) +
  labs(x = bquote(atop("Correlation"~rho,
                       .(.$CI_label))),
       tag = "Species") +
  theme(axis.title.x = element_text(size = 8.5, face = "bold"),
        axis.title.y = element_blank(),
        axis.text.y = element_text(size = 7.5, 
                                   face = "italic", 
                                   color = "black"),
        axis.text.x = element_text(size = 7.5,
                                   angle = 45,
                                   hjust = 1, 
                                   color = "black"
                                   ),
        strip.text = element_text(size = 8.5),
        panel.grid.major.x = element_blank(),
        panel.grid.minor.x = element_blank(),
        panel.grid.major.y = element_line(size = 0.25),
        legend.position = "none",
        plot.tag.position = c(0.37, 0.94),
        plot.tag = element_text(face = "bold",
                                size = 8.5,
                                hjust = 1,
                                vjust = 0.5)) +
  facet_wrap(~Metabolite)}

p.Cys.Species.corr.forest
```
  
  
### Supplementary figure 4f: GSH and Cys-Gly correlation Forest plot at Species level  
  

```{r Species-GSH-CysGly-correlation-plot-only, echo=FALSE, warning=FALSE}
# forest plot of Spearman correlation results with bonferroni-adjusted
# p-value and confidence interval
p.metab.Species.corr.forest <- cysteine_species_cor.CI %>% 
  dplyr::filter(Metabolite %in% c("GSH", "Cys-Gly")) %>% 
  dplyr::mutate(Genus_Name = str_replace_all(Species_Name, "_", " "),
                Genus_Name = str_replace(Genus_Name, "ndidatus", "."),
                Genus_Name = fct_inorder(Genus_Name),
                Sigsymbol_adj = case_when(Sigsymbol_adj == "n.s." ~ " ",
                                          TRUE ~ Sigsymbol_adj),
                StatSignif = P_adjust < 0.05,
                CI_label = paste0("(",
                                  round(Conf_level_adjusted * 100, 2),
                                  "% C.I.)")) %>%
  {ggplot(., aes(x = Rho, y = Genus_Name, color = StatSignif)) +
  geom_vline(xintercept = 0, linetype = "dotted") +
  geom_pointrange(aes(xmin = Rho_MinCI_adj,
                      xmax = Rho_MaxCI_adj),
                  size = 0.5,
                  shape = 18) +
  geom_text(aes(label = Sigsymbol_adj), 
            hjust = 0.5,
            vjust = 0.2,
            size = 3.5) +
  scale_x_continuous(expand = expansion(mult = 0.1)) +
  scale_y_discrete(expand = expansion(add = c(0.5, 1))) +
  scale_color_manual(values = c("black", "red")) +
  labs(x = bquote(atop("Correlation"~rho,
                       .(.$CI_label))),
       tag = "Species") +
  theme(axis.title.x = element_text(size = 8.5, face = "bold"),
        axis.title.y = element_blank(),
        axis.text.y = element_text(size = 7.5, 
                                   face = "italic", 
                                   color = "black"),
        axis.text.x = element_text(size = 7.5,
                                   angle = 45,
                                   hjust = 1, 
                                   color = "black"
                                   ),
        strip.text = element_text(size = 8.5),
        panel.grid.major.x = element_blank(),
        panel.grid.minor.x = element_blank(),
        panel.grid.major.y = element_line(size = 0.25),
        legend.position = "none",
        plot.tag.position = c(0.37, 0.94),
        plot.tag = element_text(face = "bold",
                                size = 8.5,
                                hjust = 1,
                                vjust = 0.5)) +
  facet_wrap(~Metabolite)}

p.metab.Species.corr.forest
```
  
###  Supplementary: Associated table of statistics. 

```{r make-supplementary-table-of-stats-for-forest-plot, echo=FALSE}
#Species
cysteine_species_cor.CI.supptable <- cysteine_species_cor.CI %>% 
  dplyr::mutate(Taxon_Level = "Species",
                Metabolite = str_remove(Metabolite, "-\n") %>% 
                  str_replace("\n", " ")) %>% 
  dplyr::select(Taxon_Level, Taxon_Name = Species_Name, everything()) %>% 
  dplyr::select(-c(data, Cor_Spear, starts_with("Sigsymbol"))) 

#Genus
cysteine_genus_cor.CI.supptable <- cysteine_genus_cor.CI %>% 
  dplyr::mutate(Taxon_Level = "Genus",
                Metabolite = str_remove(Metabolite, "-\n") %>% 
                  str_replace("\n", " ")) %>% 
  dplyr::select(Taxon_Level, Taxon_Name = Genus_Name, everything()) %>% 
  dplyr::select(-c(data, Cor_Spear, starts_with("Sigsymbol"))) 

#Combine species and genus into single table
Tax.cor.CI.supptable <- dplyr::bind_rows(cysteine_genus_cor.CI.supptable,
                                         cysteine_species_cor.CI.supptable)

#Review table
Tax.cor.CI.supptable %>% 
  dplyr::mutate(Taxon_Name = str_replace_all(Taxon_Name, "_", " ")) %>% 
  knitr::kable(caption = "Statistics for forest plots of taxa relative abundance against metabolite concentrentrations")
  
```


### Make figures  
  
  
### Define output directory  
  
```{r define-output-directory, echo=FALSE}
path_output <- "Output"

path_output
```


### Main figure  
  
  
```{r make-main-figure, echo=FALSE}
p.metab.main.figure.v2 <- cowplot::plot_grid(
  plot_grid(p.cysteine.BV.boxplot.Tukey,
            p.cysteine.CT.boxplot.Tukey,
            p.cysteine.pseudocount.genuslevel.correlationplot.vertical,
            ncol = 1,
            rel_heights = c(1, 1.1, 2.5),
            scale = 0.95,
            labels = c("a", "c", "d"),
            align = "v",
            axis = "l"),
  plot_grid(
    plot_grid(p.tax.ra.heatmap.vertical,
              labels = "b"),
    plot_grid(p.Cys.Genus.corr.forest,
              p.Cys.Species.corr.forest,
              labels = c("e", "f"),
              rel_widths = c(0.85, 1),
              align = "h",
              axis = "b",
              nrow = 1),
    rel_heights = c(1.8, 1),
    ncol = 1),
  rel_widths = c(1, 2.4),
  nrow = 1
  )

p.metab.main.figure.v2
```


```{r save-main-figure-v2, echo=FALSE}
###File name to save
f_p.metab.main.figure.v2 <- "L_iners_Figure_3_plots_v3.pdf"
f_p.metab.main.figure.v2

### Write to file
cowplot::ggsave2(filename = f_p.metab.main.figure.v2,
                 plot = p.metab.main.figure.v2,
                 path = path_output,
                 width = 8.5,
                 height = 8.5)
```

  
### Supplementary figure 3 on BV-CT relationship, Cys-Lacto correlation plots, and relative abundances and prevalences of prevalent taxa shown in correlation forest plots.  
  

```{r 1st-supp-fig-v2, echo=FALSE}
p.metab.supp.Fig.1st <- cowplot::plot_grid(
  plot_grid(p.BV.CT.tally.heatmap,
            # p.gsh.CT.boxplot.Dunn, 
            # p.cysgly.CT.boxplot.Dunn,
            p.cysteine.pseudocount.lacto.specieslevel.correlationplot,
            labels = c("a", "b"),
            scale = 0.95,
            nrow = 1,
            rel_widths = c(1, 1),
            axis = "l"),
  plot_grid(p.Genus.p50.ra.prev,
            p.Species.p50.ra.prev,
            labels = c("c", "d"),
            rel_widths = c(0.90, 1),
            align = "h",
            axis = "tb",
            scale = 0.95,
            nrow = 1),
  ncol = 1,
  rel_heights = c(0.7, 1))
  
p.metab.supp.Fig.1st
```


```{r save-1st-supp-fig-v2, echo=FALSE}
###File name to save
f_p.metab.supp.Fig.1st <- "L_iners_Supp_Figure_3_aw_3_plots_v4.pdf"
f_p.metab.supp.Fig.1st

### Write to file
cowplot::ggsave2(filename = f_p.metab.supp.Fig.1st,
                 plot = p.metab.supp.Fig.1st,
                 path = path_output,
                 width = 8.5,
                 height = 6)
```
  
  
### Second supplementary figure (4): GSH and Cys-Gly concs with BV, CT, and Forest plots  
  
  
```{r save-2nd-supp-fig-v1, echo=FALSE}
p.metab.supp.Fig.GSH.CysGly <- cowplot::plot_grid(
  plot_grid(p.gsh.BV.boxplot.Dunn,
            p.cysgly.BV.boxplot.Dunn,
            p.gsh.CT.boxplot.Dunn, 
            p.cysgly.CT.boxplot.Dunn,
            labels = c("a", "b", "c", "d"),
            scale = 0.95,
            nrow = 1,
            align = "h",
            axis = "tb",
            rel_widths = c(1, 1, 1, 1)),
  plot_grid(p.metab.Genus.corr.forest,
            p.metab.Species.corr.forest,
            labels = c("e", "f"),
            rel_widths = c(0.90, 1),
            align = "h",
            axis = "tb",
            scale = 0.95,
            nrow = 1),
  ncol = 1,
  rel_heights = c(0.65, 1))
  
p.metab.supp.Fig.GSH.CysGly
```


```{r supp-fig-v1, echo=FALSE}
###File name to save
f_p.metab.supp.Fig.GSH.CysGly <- "L_iners_Supp_Figure_4_GSH_CysGly_aw_3_plots_v2.pdf"
f_p.metab.supp.Fig.GSH.CysGly

### Write to file
cowplot::ggsave2(filename = f_p.metab.supp.Fig.GSH.CysGly,
                 plot = p.metab.supp.Fig.GSH.CysGly,
                 path = path_output,
                 width = 8.5,
                 height = 6)
```
  
  
### Write table of ASV taxonomy to TSV file  
  
```{r write-ASV-taxonomy-table-to-file, echo=FALSE}
###File name to save
f_ASV.tax.table.names <- "ASV_taxonomy_assignments_v1.txt"
f_ASV.tax.table.names

#Write to TSV file
readr::write_tsv(ASV.tax.table.names,
                 file.path(path_output, f_ASV.tax.table.names))
```
  
### Write table of taxa-metabolite correlations from forest plots to TSV file  
  

```{r write-correlations-stats-table-to-file, echo=FALSE}
###File name to save
f_Tax.cor.CI.supptable <- "Taxa_metab_correlations_stats_supp_v3.txt"
f_Tax.cor.CI.supptable

#Write to TSV file
readr::write_tsv(Tax.cor.CI.supptable,
                 file.path(path_output, f_Tax.cor.CI.supptable))
```

### Session information  
  
```{r session-info, echo=FALSE}
sessionInfo()
```
````
