## Supplementary File 2 for "Cysteine dependence in *Lactobacillus iners* constitutes a novel therapeutic target to modify the vaginal microbiota": README_L_iners_16S_BV_Metab_Analysis.txt

README file for analysis of correlations between BV, 16S microbiota composition, and cysteine-related metabolites in Bloom et al manuscript on Lactobacillus iners. Contains information on code, analysis files, and directory structure to reproduce the analysis, as well as required dependencies. Additional details on the analysis are contained within the R Markdown analysis file itself.


Directory contains the following files:
* FRESH_16S_metabolite_analysis_for_L_iners_paper_2021_06_10.Rproj
  - (R Project file to be opened in R Studio to perform analysis)
* L_iners_Cysteine_analysis_2021_06_10_v5.Rmd
  - (R Markdown file containing analysis code)
* This README.txt file: README_L_iners_16S_BV_Metab_Analysis.txt


Also contains the following subdirectories with their contents:

* Data/FRESH_data, which contains the following two files:
  + ps0_FRESH_16S_BV_for_cys_metabolomics_2021_03_07_v3.RDS
    + Phyloseq object containing the requisite DADA2-processed 16S data with metadata including BV status and cervicotype
  + cysteine_metabolomics_data_2020_09_03.txt
    + Tab-delimited file containing metabolite concentrations for the samples.


* Output, empty directory into which figures, tables, and other files are saved.

The dependencies for the analysis are as follows:


## R version 3.6.3 (2020-02-29)
## Platform: x86_64-apple-darwin15.6.0 (64-bit)
## Running under: macOS  10.16
## 
## Matrix products: default
## BLAS:   /Library/Frameworks/R.framework/Versions/3.6/Resources/lib/libRblas.0.dylib
## LAPACK: /Library/Frameworks/R.framework/Versions/3.6/Resources/lib/libRlapack.dylib
## 
## locale:
## [1] en_US.UTF-8/en_US.UTF-8/en_US.UTF-8/C/en_US.UTF-8/en_US.UTF-8
## 
## attached base packages:
## [1] stats     graphics  grDevices utils     datasets  methods   base     
## 
## other attached packages:
##  [1] table1_1.4        cowplot_1.1.1     ggpubr_0.4.0      knitr_1.33       
##  [5] forcats_0.5.1     stringr_1.4.0     dplyr_1.0.5       purrr_0.3.4      
##  [9] readr_1.4.0       tidyr_1.1.3       tibble_3.1.1      ggplot2_3.3.3    
## [13] tidyverse_1.3.1   e1071_1.7-6       DescTools_0.99.41 phyloseq_1.30.0  
## 
## loaded via a namespace (and not attached):
##   [1] colorspace_2.0-0    ggsignif_0.6.1      ellipsis_0.3.1     
##   [4] class_7.3-18        rio_0.5.26          XVector_0.26.0     
##   [7] fs_1.5.0            gld_2.6.2           rstudioapi_0.13    
##  [10] proxy_0.4-25        farver_2.1.0        fansi_0.4.2        
##  [13] mvtnorm_1.1-1       lubridate_1.7.10    xml2_1.3.2         
##  [16] codetools_0.2-18    splines_3.6.3       rootSolve_1.8.2.1  
##  [19] ade4_1.7-16         Formula_1.2-4       jsonlite_1.7.2     
##  [22] broom_0.7.6         cluster_2.1.2       dbplyr_2.1.1       
##  [25] compiler_3.6.3      httr_1.4.2          backports_1.2.1    
##  [28] assertthat_0.2.1    Matrix_1.3-2        cli_2.5.0          
##  [31] htmltools_0.5.1.1   prettyunits_1.1.1   tools_3.6.3        
##  [34] igraph_1.2.6        gtable_0.3.0        glue_1.4.2         
##  [37] lmom_2.8            reshape2_1.4.4      Rcpp_1.0.6         
##  [40] carData_3.0-4       Biobase_2.46.0      cellranger_1.1.0   
##  [43] jquerylib_0.1.4     vctrs_0.3.7         Biostrings_2.54.0  
##  [46] multtest_2.42.0     ape_5.5             nlme_3.1-152       
##  [49] iterators_1.0.13    xfun_0.22           openxlsx_4.2.3     
##  [52] rvest_1.0.0         lifecycle_1.0.0     rstatix_0.7.0      
##  [55] zlibbioc_1.32.0     MASS_7.3-53.1       scales_1.1.1       
##  [58] hms_1.0.0           parallel_3.6.3      biomformat_1.14.0  
##  [61] rhdf5_2.30.1        expm_0.999-6        yaml_2.2.1         
##  [64] curl_4.3            Exact_2.1           sass_0.3.1         
##  [67] stringi_1.5.3       highr_0.9           S4Vectors_0.24.4   
##  [70] foreach_1.5.1       permute_0.9-5       BiocGenerics_0.32.0
##  [73] boot_1.3-27         zip_2.1.1           rlang_0.4.10       
##  [76] pkgconfig_2.0.3     evaluate_0.14       lattice_0.20-41    
##  [79] Rhdf5lib_1.8.0      labeling_0.4.2      tidyselect_1.1.0   
##  [82] plyr_1.8.6          magrittr_2.0.1      R6_2.5.0           
##  [85] IRanges_2.20.2      generics_0.1.0      DBI_1.1.1          
##  [88] pillar_1.6.0        haven_2.4.1         foreign_0.8-76     
##  [91] withr_2.4.2         mgcv_1.8-35         abind_1.4-5        
##  [94] survival_3.2-11     modelr_0.1.8        crayon_1.4.1       
##  [97] car_3.0-10          utf8_1.2.1          rmarkdown_2.7      
## [100] progress_1.2.2      grid_3.6.3          readxl_1.3.1       
## [103] data.table_1.13.6   vegan_2.5-7         reprex_2.0.0       
## [106] digest_0.6.27       stats4_3.6.3        munsell_0.5.0      
## [109] bslib_0.2.4
